# Folds induced by multiple parallel or antiparallel double-helices: (pseudo)knotting of single-stranded RNA

**DOI:** 10.1101/2021.03.12.435210

**Authors:** Stephen T. Hyde

## Abstract

We develop tools to explore and catalogue the topologies of knotted or pseudoknotted circular folds due to secondary and tertiary interactions within a closed loop of RNA which generate multiple double-helices due (for example) to strand complementarity. The fold topology is captured by a ‘contracted fold’ which merges helices separated by bulges and removes hairpin loops. Contracted folds are either trivial or pseudoknotted. Strand folding is characterised by a rigid-vertex ‘polarised strand graph’, whose vertices correspond to double-helices and edges correspond to strands joining those helices. Each vertex has a plumbline whose polarisation direction defines the helical axis. That polarised graph has a corresponding circular ribbon diagram and canonical alphanumeric fold label. Key features of the ‘fully-flagged’ fold are the arrangement of complementary domains along the strand, described by a numerical bare fold label, and a pair of binary ‘flags’: a parity flag that specifies the twist in each helix (even or odd half-twists), and an orientation flag that characterises each double-helix as parallel or antiparallel. A simple algorithm is presented to translate an arbitrary fold label into a polarised strand graph. Any embedding of the graph in 3-space is an admissible fold geometry; the simplest embeddings minimise the number of edge-crossings in a planar graph drawing. If that number is zero, the fold lies in one of two classes: (a)-type ‘relaxed’ folds, which contain conventional junctions and (b)-type folds whose junctions are described as meso-junctions in H. Wang and N.C. Seeman, *Biochem*, vol. 34, pp920-929. (c)-type folds induce polarised strand graphs with edge-crossings, regardless of the planar graph drawing. Canonical fold labelling allows us to sort and enumerate all ‘semi-flagged’ folds with up to six contracted double-helices as windings around the edges of a graph-like fold skeleton, whose cyclomatic number - the ‘fold genus’ - ranges from 0 – 3, resulting in a pair of duplexed strands along each skeletal edge. Those semi-flagged folds admit both even and odd double-helical twists. Appending specific parity flags to those semi-flagged folds gives fully-flagged (a)-type folds, which are also enumerated up to genus-3 cases. We focus on all-antiparallel folds, characteristic of conventional ssRNA and enumerate all distinct (a), (b) and (c)-type folds with up to five double-helices. Those circular folds lead to pseudoknotted folds for linear ssRNA strands. We describe all linear folds derived from (a) or (b)-type circular folds with up to four contracted double-helices, whose simplest cases correspond to so-called *H*, *K* and *L* pseudoknotted folds, detected in ssRNA. Fold knotting is explored in detail, via constructions of so-called antifolds and isomorphic folds. We also tabulate fold knottings for (a) and (b)-type folds whose embeddings minimise the number of edge-crossings and outline the procedure for (c)-type folds. The inverse construction - from a specific knot to a suitable nucleotide sequence - results in a hierarchy of knots. A number of specific alternating knots with up to 10 crossings emerge as favoured fold designs for ssRNA, since they are readily constructed as (a)-type all-antiparallel folds.

## INTRODUCTION

Tangling at the biomolecular scale of various flavours, from unknotted tangles to ‘pseudoknotted’ an *bona fide* knotted configurations, is both confounding and fascinating. The specificity of Watson-Crick base-pairing in DNA and RNA strands can lead to specific tangles *in vivo*, as well as attractive design goals in synthetic DNA and RNA science (1,2). The synthesis of DNA and RNA knots were significant milestones in the then-nascent field of DNA nanotechnology (1, 3–6) and subsequent advances in RNA nanotechnology have revived interest (7). From a biological perspective, it is clear that tangling of DNA and RNA is an active phenomenon *in vivo*. A number of distinct topoisomerase enzymes, which switch strand crossings and therefore tangling, have been detected in a variety of organisms. The ubiquity of both DNA and RNA topoisomerases (8–10) suggests that both biopolymers are prone to non-trivial entanglements *in vivo*. The tangling of DNA or RNA may have multiple origins, from unwinding of double-helices during replication, to secondary and tertiary interactions within or between strands. Known structures exhibit a variety of entanglements, though well-characterised structures observed to date are possibly all unknotted (11). That observation is perhaps unsurprising, given the key role of mRNA in protein transcription, where the maintenance of unknotted strands may be crucial for biological function. On the other hand, the inherent stickiness of pseudoknots in viral RNA allows for programmed ribosomal frameshifting, common to human coronaviruses (12). Pseudoknotted “ring knot” folds have been detected in flaviviruses, including Dengue, West Nile and Murray Valley Encephalitis viruses (13, 14) and more recently in dianthoviruses (15). The fold is also present in the frameshifting region of the SARS-Cov and SARS-Cov-2 viral genomes(16) and is common to exoribonuclease resistant ssRNA (xrRNA), whose topology is very likely responsible for their resistance to ribonuclease breakdown (17, 18). Evidently, increasingly sophisticated structural studies are suggesting increasingly complex fold topologies.

Here we address one simple driver for topological complexity, namely, the duplexing of a DNA or RNA strand induced by some arrangement of complementary sequences along its length. We focus on pseudoknotted and knotted structural topologies, and build a taxonomy of simplest cases, based on the topological complexity of the folded strand. Explicit topologies are deduced for ‘contracted’ folds, which ignore simpler secondary structural features, focussing on more deeply tangled, pseudoknotted folds. An oriented (rigid-vertex) strand graph, whose vertices are double-helices equipped with a local orientation, can be constructed for an arbitrary arrangement of self-complementary nucleotides along a strand. We draw up a catalog of simpler folds, most of which combine parallel and antiparallel double-helices, and distinguish simpler ‘relaxed’ folds, from others. All-antiparallel folds, most relevant to RNA and DNA, are described in detail. The analysis also allows the design of nucleotide sequences along a strand to induce a specific knot via duplexing alone, which suggests that specific alternating knots (with ten crossings or less) are more simply assembled by duplexing of DNA or RNA than others.

The concepts introduced here: rigid-vertex graphs, circular and linear diagrams and fold topology, have been used in this context in various guises by others. Our constructions are however different, focussed squarely on (re)constructive algorithms to allow explicit spatial realisations of fold topologies from those concepts. The utility of graph descriptions of RNA structure has been demonstrated by others, see for example (19) and references therein, though our graphs are exclusively degree-4 structures, in contrast to that work. Rigid-vertex graphs of even degree were recognised almost thirty years ago as relevant to folding of RNA (20). Our graphs differ in detail, since they include polarisation directions at each vertex. The arc diagrams found (for example) in that work, as well as many other more recent descriptions of folding (e.g. related Nussinov diagrams), resemble our construction of linear and circular fold diagrams. We extend those arc folding diagrams to what we call ‘circular diagrams’, allowing parallel or antiparallel double-helices. Our topological classification of fold by genus turns out to be related (though not identical in detail) to earlier fundamental studies, which were motivated by the search for numerically efficient algorithms to explore pseudoknotted folds in RNA (21–24). Those papers were concerned with the thermodynamics and statistics of forming pseudoknotted RNA, in order to arrive at likely binary interactions between nucleotides in an RNA strand leading to secondary and tertiary structures. Here we assume that those binary interactions are explicitly known, and determine possible strand topologies consistent with those interactions.

The classical Watson-Crick double-helix of (duplexed) DNA is induced by complementary nucleotides within a pair of DNA strands. A very similar duplexed structure is realised in regions of a single strand of RNA (ssRNA) with complementary nucleotides, where the string folds back onto itself, by self-duplexing. In the first part of the paper, we will assume that all folds emerge by duplexing of a closed single-stranded loop, allowing the most general folds, containing both parallel and antiparallel double-helices. The simpler cases of those folds will be cataloged by topological genus and fold type. With those results in place, the second part of the paper will focus on folds containing exclusively all-antiparallel duplexes, allowing a deeper study of those folds, which are more likely to be found *in vivo* and realised synthetically.

## METHODS

### Arbitrary folds

Consider first a simple representation of a double-helix formed by a single closed loop, formed for example by fusing 5′ and 3′ ends of a fragment of ssRNA. We will model the unfolded loop by a circle which represents the perimeter of a ‘circular diagram’ and adopt the convention that a closed walk from the 5′ to 3′ end is represented by a clockwise traverse of the circle, usually starting at 12 o’clock. A closed double-helix is encoded in this circular diagram by a ribbon which connects the complementary domains on the string, forming a bridge anchored at two ‘footings’, each occupying the nucleotides contributing to that double-helix. The width of both footings, assumed to be equal, and given by the ribbon width, is determined by the number of duplexed nucleotides: typically, a single full-turn of a DNA or RNA helix is produced by just over 10 nucleotides on each local strand. The number of half-twists in the double-helix, which we denote *t*, is therefore approximately equal to the number of nucleotides in each local strand divided by 5. That helix may be left- or right-handed; we adopt the convention that right-and left-handed twists have t > 0 and *t* < 0 respectively. Circular diagrams encoding a single double-helix with t half-twists, where t is either even or odd, are illustrated in the two left-hand columns in Fig. 1. If the strands in the duplex are antiparallel, *a* is complementary to *b* and *c* to *d*, so the directed footing *ad* is duplexed to *bc*, as in the top row of Figs. 1(a,b). In contrast, if the strands are parallel, *a* is complementary to *c* and *b* to *d*, so the footing *ad* is duplexed to *cb* rather than *bc*. The ribbon encoding that parallel double-helix is therefore twisted as in the top row of Figs. 1(c,d). To avoid any confusion between the twisted or untwisted ribbons and the unrelated twist in the double-helix *t*, we call an untwisted ribbon ‘annular’ and a twisted ribbon ‘moebius’ (since those terms describe the respective ribbon topologies of their ends are glued.) Antiparallel and parallel double-helices are therefore coded within circular ribbon diagrams by annular and moebius ribbons respectively. (For brevity, we will call circular ribbon diagrams simply ‘circular diagrams’.)

**Figure 1:**
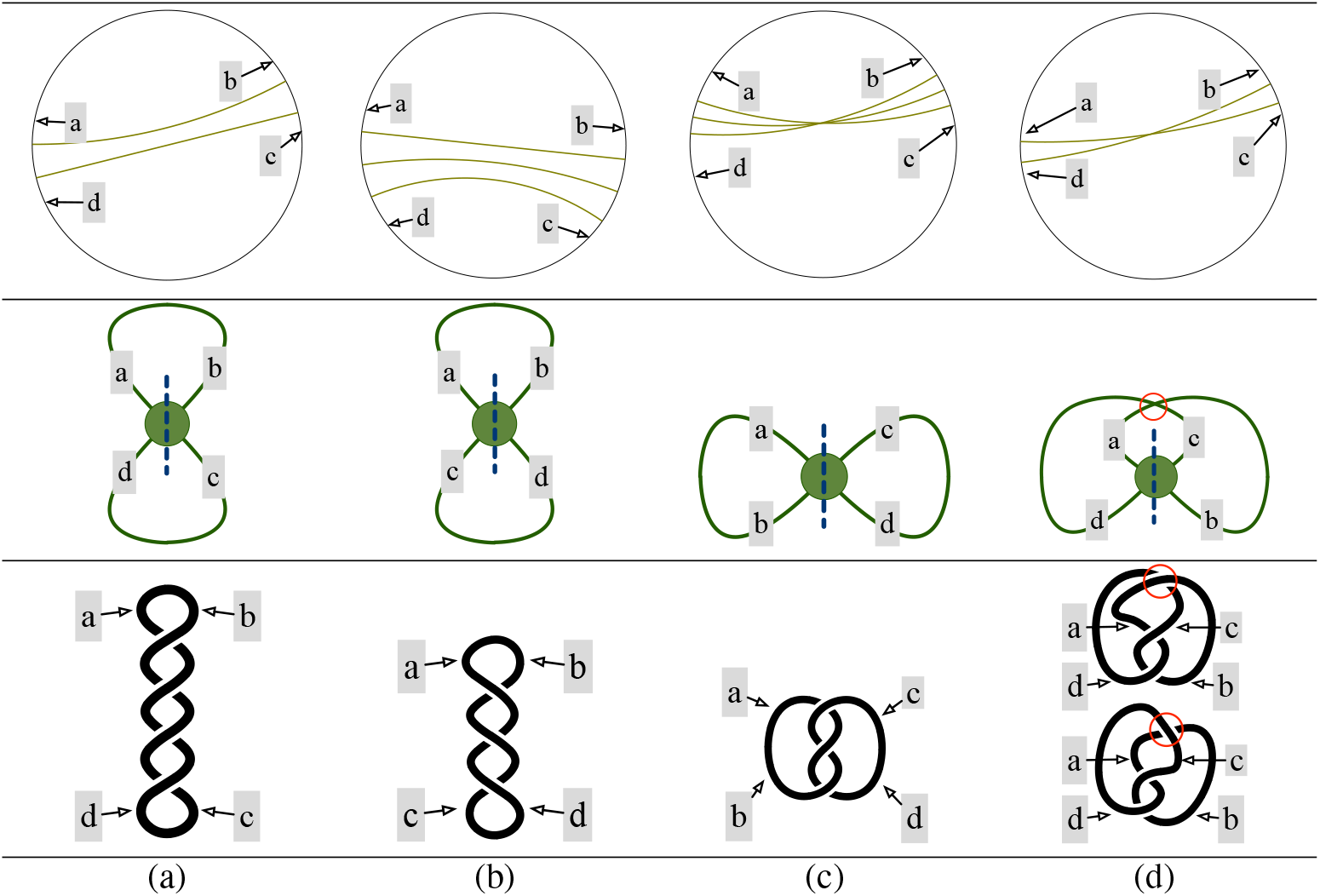
Single-ribbon folds. (Top row) Circular diagrams for single-ribbon folds. Ribbons are represented by pairs or triples of ruled lines. The perimeter circle represents a single-stranded loop studded with nucleotides, spanned by a ribbon representing a duplex with complementary strand domains at each foot of the ribbon. (a,b) An annular ribbon, encoding an antiparallel double-helix; (c,d) a moebius ribbon, encoding a parallel double-helix. The number of rulings in each ribbon is either two (a,d) or three (b,c), indicating ribbons encoding duplexes whose number of half-twists, *t*, is even or odd respectively.) (Middle row) Oriented (rigid-vertex) strand graphs for each fold, with vertical plumblines. The essential edge-crossing in the polarised strand graph for fold (d) is marked by a red circle. (Bottom row) Typical strand windings for each fold, with (a) *t* = 4, (b) 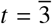, (c) *t* = 3 and (d) *t* = 2. The edge crossing in the polarised strand graph for fold (d) allows a pair of simplest embeddings, depending on the essential edge-crossing circled in red.

The four folds induced by these single-ribbon diagrams can be compactly portrayed by the ‘polarised strand graphs’ shown in the second row in Fig. 1. The double helix is replaced by a single vertex (marked by the central green dot) and unduplexed portions of the loop, outside the footings *ab* and *cd*, are replaced by curved graph edges. The topology of those strand graphs are identical for all four folds, despite their structural differences, which depend on the ribbon type, as well as (even or odd) twist parity of t: all four graphs contain one vertex (*V*) and two edges (*a, b*), (*c, d*), whose ends are incident to that vertex. That graph topology can be encoded by the edge-vertex pair denoted *V*: (*ab, cd*). To distinguish the four distinct folds, we append additional ‘orientation’ and ‘parity’ flags to the strand graph, characterising the ribbon type (annular/moebius) and twist parity (even/odd) respectively. The cyclic arrangement of all four strands emanating from the double-helices, at sites *a, b, c* and *d* depend on the helix parity and parallelism, as shown in the second row of Fig 1. Cyclic orderings of labels around the vertex are *abcd, abdc, acbd* and *acdb* for annular-even, annular-odd, moebius-even and moebius-odd diagrams respectively, since those orderings correspond to the clockwise sequence of sites on the folded strand surrounding the relevant double-helix. Those letters label the graph edges, as shown in the third row of Fig 1. That cyclic ordering of edge labels is crucial, since it encodes the orientation and parity flags. We therefore demand it is retained in all graph embeddings. It may be embedded as in Fig. 1 with clockwise orientation, or, if the vertex is drawn ‘face-down’ (assuming the graph embedding in Fig. 1 is face-up) reversed to give anticlockwise ordering. In addition, the orientation of the strand at *a* need not be *NW* as in Fig. 1, any rotated embedding, moving *a* to e.g. *SE* orientation is possible, provided the orientation of the other three other outgoing strand fragments (*b, c, d*) is rotated likewise. Those requirements imply any cyclic or reverse cyclic permutation of the cyclic label is allowed: for example, *abcd* in Fig. 1(a) can be permuted to give the cycles *bcda, cdab, dabc, dcba, adcb, badc* and *cbad*. That permutation constraint is equivalent to modelling the strand graphs as degree-4 ‘rigid-vertex’ graphs ((25)).

The cyclic edge-ordering may induce an edge-crossing in a planar drawing of the graph embedding. For example, a single edge-crossing is essential in any planar drawing of the strand graph corresponding to the even-parity moebius-ribbon fold in Fig. 1(d); the other cases in Fig. 1(a-c) allow the polarised strand graph to be crossing-free. Since the polarised strand graph topology is indifferent to the embedding of this edge-crossing, two distinct embeddings of the graph are possible for any edge-crossing, depending on the choice of upper and lower strands in the embedding, as shown in Fig. 2. Therefore any ‘edge-crossing’ in a planar drawing of the strand graph is indicated by a ‘shadow drawing’ of the crossed edges (circled in red in Figs. 1(d) and 2), which allows for either strand to cross on top of the other. Those alternative embeddings affect the spatial tangling of the strand, as shown in the windings drawn in the bottom row of Fig. 1(d).

**Figure 2:**
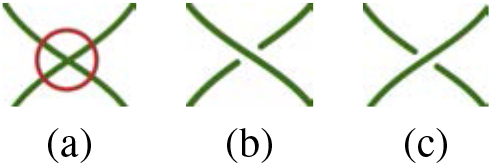
(a) Schematic ‘shadow drawing’ of an edge-crossing in an embedding of a strand graph. (b,c) Alternative embeddings of the crossing.

Thus far, the rigid-vertex strand graph is ‘unpolarised’. A ‘polarised’ (rigid-vertex) strand graph is formed by inserting a ‘plumbline’ through each vertex in the unpolarised strand graph, whose orientation defines of the axis of the double-helix corresponding to that vertex. The complete ‘rigid-vertex polarised strand graphs’ (called for brevity ‘polarised strand graphs’) are shown in the second row of Fig. 1: they include vertical plumblines through the graph vertex. Any embedding of a polarised strand graph leads to a corresponding embedding of the strand fold, including duplexing. The fold embedding is recovered from the polarised strand graph by replacing its vertex with a double-helix of appropriate (even or odd) twist, extended along the plumbline, as shown in the bottom row of Fig. 1. It is helpful to view the plumbline as a local *NS* axis, defining the poles of the vertex: if the vertex is rotated (preserving its rigid-vertex ordering) the plumbline follows accordingly. Note that it is not oriented (*NS* = *SN*), so flipping the vertex about horizontal (*NS*) or vertical (*EW*) axes does not reorient the plumbline, although it swaps edge-ordering.

### Circular diagrams and fold graphs

The construction of polarised strand graphs can be extended to include folds with multiple parallel or antiparallel duplexes (double-helices), each contributing a ribbon to the circular diagram. A fold with *n* duplexes has a circular diagram containing *n* ribbons. That diagram induces a polarised strand strand graph with *n* degree-4 vertices and 2*n* edges (*A, B*,…,), analogous to the single-vertex graphs in the second row of Fig. 1. The polarised strand graph is built from the circular diagram as follows. As above, each ribbon maps to a vertex of the graph and the perimeter strand fragments between ribbons footings map to graph edges and the graph topology can be determined from a cyclic walk around the perimeter. For example, the circular diagram in Fig. 3(a) corresponds to a polarised strand graph with 12 edges (labelled *A - L*) and 6 vertices (labelled 1 - 6). The strand graph for this fold has edge-vertex data *A*: (1, 2), *B*: (2, 3), *C*: (3, 1), *D*: (1, 4), *E*: (4, 5), *F*: (5, 2), *G*: (2, 4), *H*: (4, 6), *I*: (6, 3), *J*: (3, 5), *K*: (5, 6), *L*: (6, 1), which induces a graph topology - possibly with invalid rigid-vertex ordering. One embedding of that graph is shown in Fig. 3(b). (In fact, that embedding respects the rigid-vertex ordering, as we show next.) A polarised strand graph containing multiple vertices is built from its multi-ribbon circular diagram in stages. First, build a (sub)graph for each vertex, with four edges radiating from that vertex. The ordering of edges around each vertex is found from the one-ribbon algorithm, described above: namely *abcd* if the vertex corresponds to an annular ribbon of even parity, *abdc* for an odd-parity annular ribbon, *acdb* for an odd-parity moebius ribbon and *acbd* for an even-parity moebius ribbon (Figs. 1(a,b,c,d) respectively). Those labels (*a, b, c, d*) map to the labels of edges incident to each ribbon *i* = 1 – 6 in the circular diagram in Fig. 3(a). The orientation and parity flags can be read from the diagram for each ribbon *i*: it is either annular or moebius with even or odd parity (2 or 3 rulings respectively). The mapping can be found by overlaying the relevant one-ribbon template in Figs. 1(a,b,c,d) whose flags match those of ribbon *i* so that the single ribbon in the template aligns with ribbon *i*. With that alignment in place, the four edges *a, b, c, d* in the template overlay four edges incident to ribbon *i*. For example, ribbon 4 in Fig. 3(a) is incident to edges with labels D, *E, G* and *H*. The ribbon is roughly horizontal, as are the template ribbons in Fig. 3, so edges can be matched as *a* → *H, b* → *D, c* → *E, d* → *G*. Alternatively, the circular diagram in Fig. 3(a) can be rotated *ca*. 180°, or flipped etc., giving different edge mappings, related by cyclic or reverse cyclic permutations. All the resulting mappings are also allowed, since they will lead to identical rigid-vertex ordering in the subgraph. That ordering is equivalent to that of the relevant single-vertex graph in Figs. 1, namely either *abcd, abdc, acdb* or *acbd*, depending on the orientation and parity (annular and even, annular and odd, etc.) flags. For example, ribbon 4 in Fig. 3(a) is an odd-parity moebius ribbon, corresponding to the pattern in Figs. 1(c). The relevant (clockwise) cyclic label for this flagged duplex is *cdba*, which maps to *EGDH*. The (clockwise) cyclic ordering around vertex 4 is therefore *E, G, D, H* (which is in fact that in Fig. 3(b)). The analogous mapping is done for the remaining 5 ribbons in Fig. 3(a), leading to 6 star-like rigid-vertex subgraphs, whose edges are labelled letters *A – L* in (clockwise!) order around their central vertex *i*. (Remaining vertices 1, 6 are moebius ribbons of odd parity, 2, 3, 5 are annular, even parity ribbons.) Since each edge *A – L* adjoins two ribbons in the circular diagram, with labels *i, j*, any edge label is common to two subgraphs. (In general, *i* and *j* may share the same label, as in Fig. 1, although they differ in Fig. 3(a).) The complete (unpolarised) rigid-vertex strand graph is constructed by gluing the star-like subgraphs together along those common edges. That construction induces rigid-vertex ordering corresponding to the ordering in Fig. 3(c), so that drawing is therefore a faithful embedding of the unpolarised strand graph of the fold whose circular diagram is that in Fig. 3(a). (Note that this embedding has uncrossed strand graph edges, other embeddings may be crossed.) Lastly, the polarised graph is formed by inserting plumblines joining the *N, S* poles at each vertex. For example, the plumbline for ribbons with flags moebius and odd (Fig. 1(c)) passes between the edges *a, c* and *b, d*. Therefore the plumbline through vertex 4 runs between edges *H, E* and *D, G*, as shown in Fig. 3(c). The analogous matching of plumblines for all vertices 1 - 6 leads to the polarised strand graph shown in Fig. 1(c).

**Figure 3:**
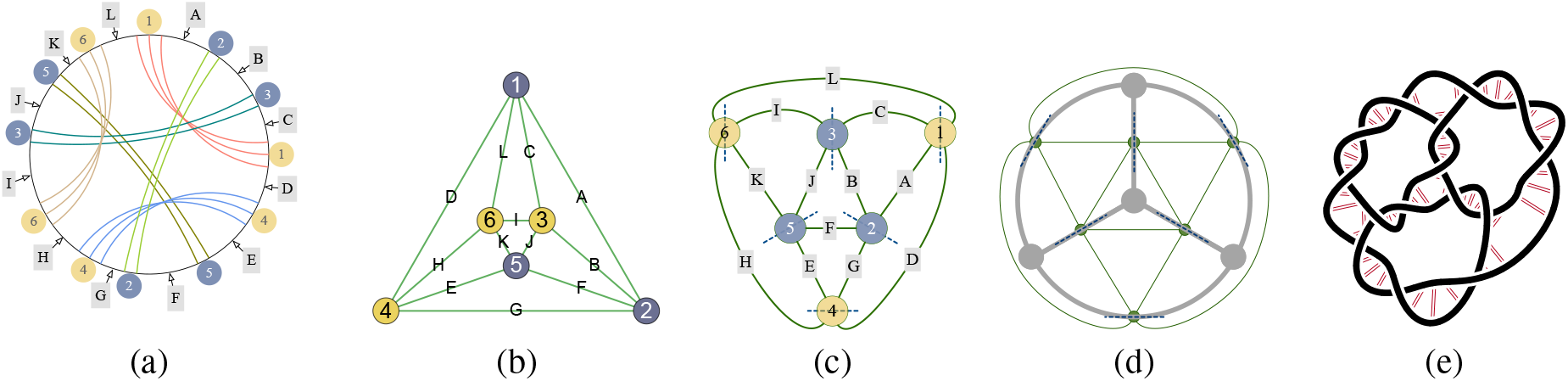
(a) Circular diagram of a fold, whose ribbons are labelled 1 – 6. Annular and moebius ribbons are labelled 2,3,5 and 1,4,6 respectively (grey and yellow circular labels) and unduplexed strand domains between ribbons are labelled *A* – *L*. (b) An embedding of the polarised strand graph whose vertices are labelled 1 - 6 corresponding to the labels in (a). (c) The polarised strand graph, whose plumblines are marked by dashed axes through each vertex. (d) The skeleton induced by this polarised strand graph. (e) A specific strand winding containing 6 double helices with 2 or 3 half-twists.

Conversely, a circular diagram can be built from a polarised strand graph, provided each vertex is flagged by its twist parity, e.g. Fig. 4(a). First, label each vertex (in any way) 1, 2, 3,…. Next, form an Euler cycle *L* around the graph which traverses each edge once, passing through each vertex (i) twice. If the twist parity at vertex *i* is even, the cycle passes through *i* without crossing the plumbline; if *i* has odd parity, it crosses the plumbline (and itself). Thus, the polarised strand graph embedding whose twist parities are labelled *o* and *e* (odd or even) is shown in Fig. 4(a). The parities imply a directed cycle described by the polygon *L* = 162351346542 shown in Fig. 4(b). *L* also encodes the cycle around the circular perimeter in the circular diagram corresponding to the polarised strand graph, giving the ordering of labels 1 - 6 around that perimeter (e.g. 162351346542 in Fig. 4(c)). Each pair of indices *i* locate the footers of ribbon i, determining the arrangement of all the ribbons in the circular diagram. The orientation of each ribbon (annular or moebius) is then found by ‘marking’ the cycle as follows. Set out from any vertex along the cycle L, and mark (by an arrow, or a dot) each strand on exiting a vertex until the cycle *L* is completed (e.g. red dots in Fig. 4(b)). The resulting pair of markings adjacent to each vertex in the strand graph define the orientation of that vertex, which depend on their locations along the *NS* plumbline. If both marks are located at the same (*N* or S) pole, the duplex is parallel and the associated ribbon is moebius, if they are antipodal the duplex is antiparallel, giving an annular ribbon. For example, ribbons 1 - 6 in Fig. 4(b), are annular, moebius, moebius, moebius, annular and annular respectively, giving the circular diagram shown in Fig. 4(c).

**Figure 4:**
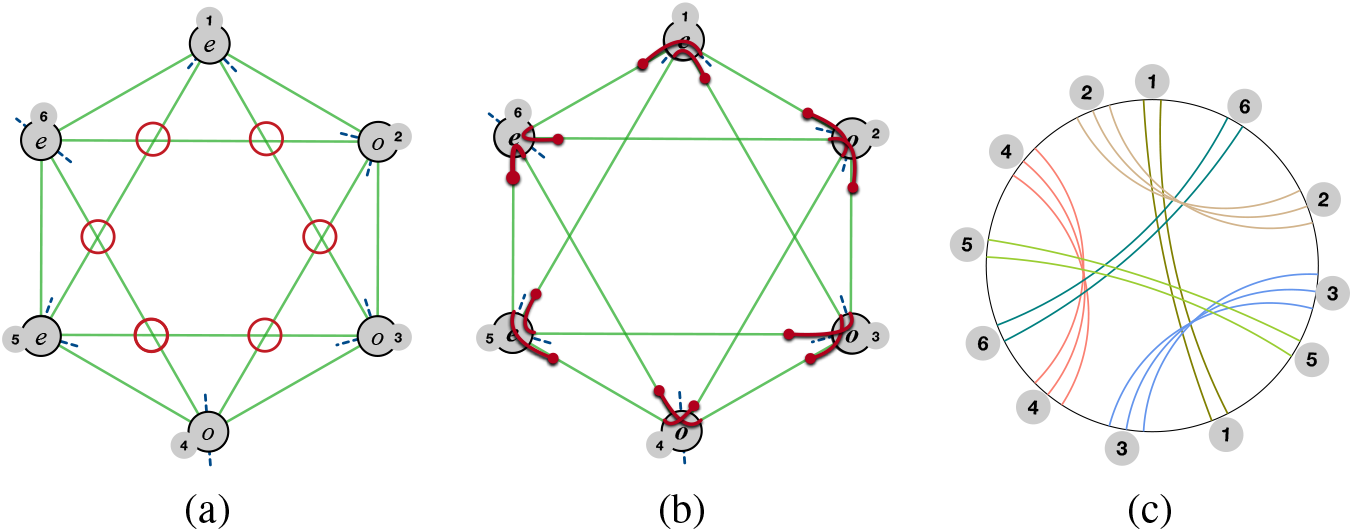
(a) An planar embedding of a polarised strand graph whose vertices (1 - 6) are flagged by twist parity (e,o) on each vertex with a number of edge-crossings (circled in red). (b) A cycle in the graph with label, *L* = 162351346542, formed by extending strand edges through vertices of the graph. Entry and exit sites at each vertex are marked by red arcs, dotted (or arrowed) at one end consistent with a directed walk around the cycle. The dot configuration determines whether vertices induce parallel or antiparallel double-helices. (c) The corresponding polarised strand graph, whose perimeter is labelled by the same cycle 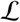.

### Contracted folds

An infinite variety of circular diagrams is possible, with any combination of annular and moebius ribbons distributed within the interior of the circular perimeter. We ignore geometric details of the fold, focussing rather on the fold topology. That means that distances around the perimeter between ribbon footers and ribbon widths are irrelevant. Ignoring those metric details allows the circular diagram to be characterised by the label 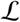. A catalog of sorted folds can will be built from these labels below. To limit the size of that catalog, we further simplify specific folds, forming ‘contracted’ folds.

Contraction can be most simply explained by example. Multiple co-axial double-helices, connected in series and bridged by unduplexed regions, known in the jargon of RNA assemblies as ‘bulges’ (26), can be collapsed into a single double-helix without affecting the fold topology. The circular diagrams and polarised strand graphs in Fig. 1 therefore also describe ‘contracted’ versions of those ‘expanded folds’, formed by collapsing series of all-parallel or all antiparallel double-helices separated by bulges into a single duplex. Circular diagrams of these expanded folds consisting of antiparallel duplexes in series contain nested annular ribbons, whereas those for parallel duplexes contain fans of crossed moebius ribbons, illustrated in Fig. 5.

**Figure 5:**
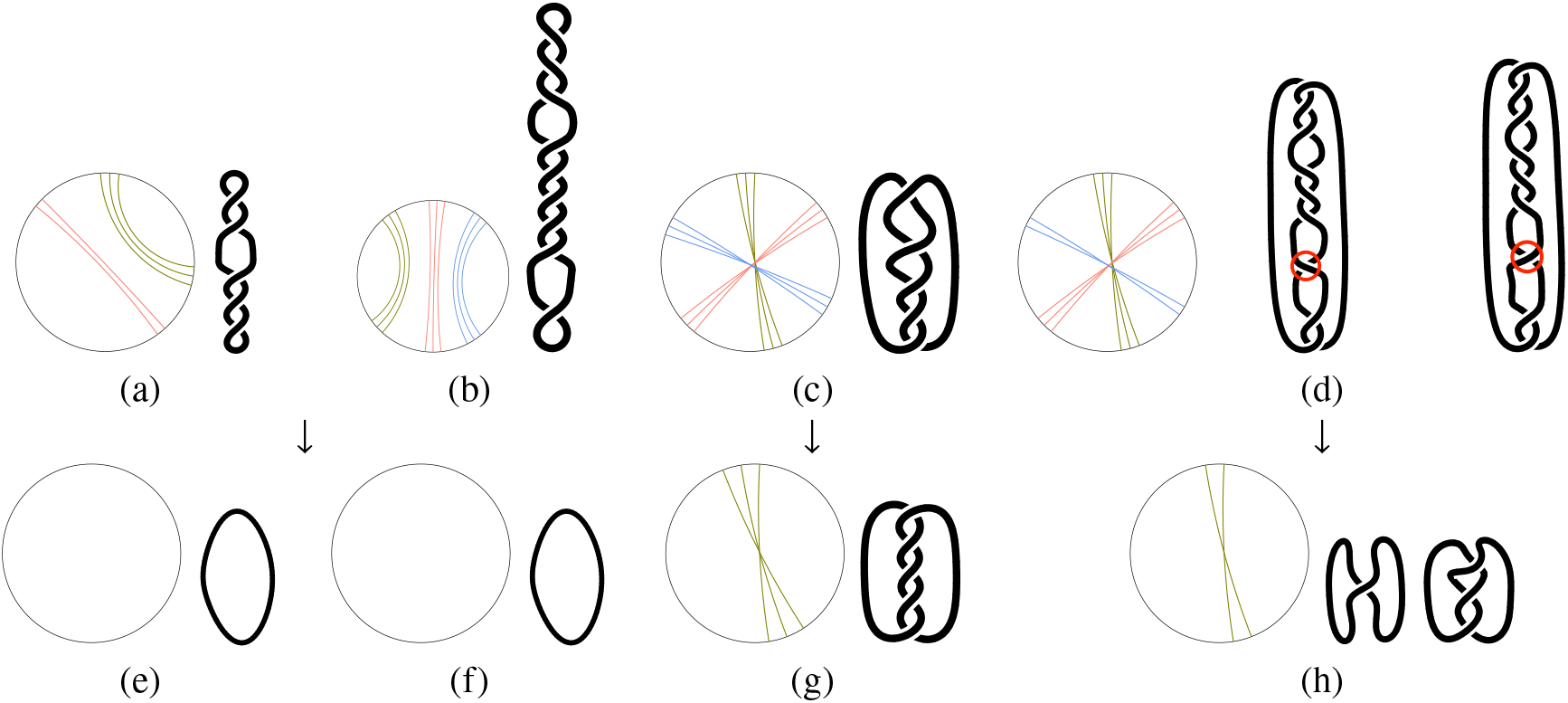
(Upper row) Circular diagrams and strand windings of folds including ‘bulges’ separating coaxial double-helices. (a) nested annular ribbons of odd and even parities associated with a strand winding containing a pair of antiparallel double-helices with twist 2 and 3 (equal to the number of half-twists) separated by a bulge; (b) nested odd-parity annular ribbons and an associated antiparallel strand winding with twists 3, 5 and 1; (c) a fan of three moebius ribbons all of odd parity, and an associated strand winding with twists 1, 1 and 3; (d) a fan including two odd and one even parity moebius ribbon and associated fold with three parallel duplexes of twists 3, 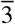 and 2. (Lower row) Contracted circular diagrams and strand windings. (e,f) Contracted folds from (a,b): the trivial (un)fold. (g,h) Contracted folds from (c,d) respectively. The fold in (d) induces an edge-crossing in the strand winding, regardless of its embedding (circled in red, *cf*. Fig.1(d)), allowing alternative contracted folds.

Contracted folds are formed by collapsing those expanded folds, easily done within the circular diagram directly. First, nested annular ribbons are merged, giving a single annular ribbon whose twist is the sum of those of the nested ribbons. (This contraction of nested annular ribbons corresponds to the formation of ‘reduced’ diagrams, introduced by Bon et al. (21, 23).) A second contraction rule holds for moebius ribbons: ‘fans’ of crossed (intersecting) moebius ribbons are closed to a single moebius ribbon whose twist is the sum of the component moebius ribbons. Both contraction operations can be done by appropriate sliding of ribbon footers around the perimeter of the circular diagram until footers of expanded ribbons merge. If those motions are blocked by intervening footers of other ribbons, contraction is impossible. A third rule is useful in that it removes an infinite variety of relatively uninteresting folds, whose circular diagrams include ‘isolated’ annular ribbons, whose pair of footers can be merged with each other, resulting in an annular ribbon bridging an arbitrarily small section of the perimeter. Those isolated annular ribbons effect local twisting of the strand only: the associated strand windings contain small twisted hairpin-looped regions. Any fold can be intercalated with an arbitrary number of those loops, without inducing new fold topologies. We therefore delete all isolated annular ribbons from contracted circular diagrams. On the other hand, isolated moebius ribbons generally induce non-trivial windings, since their deletion or insertion into a circular diagram induces a new fold that can only be effected by cutting the loop and refolding. Contracted circular diagrams and their associated folds result by applying those three rules to a circular diagram. If the contracted folding diagram is non-empty the resulting fold is nontrivially tangled; otherwise, the fold is referred to as the trivial unfold. Some examples of contractions are illustrated in Fig. 6.

**Figure 6:**
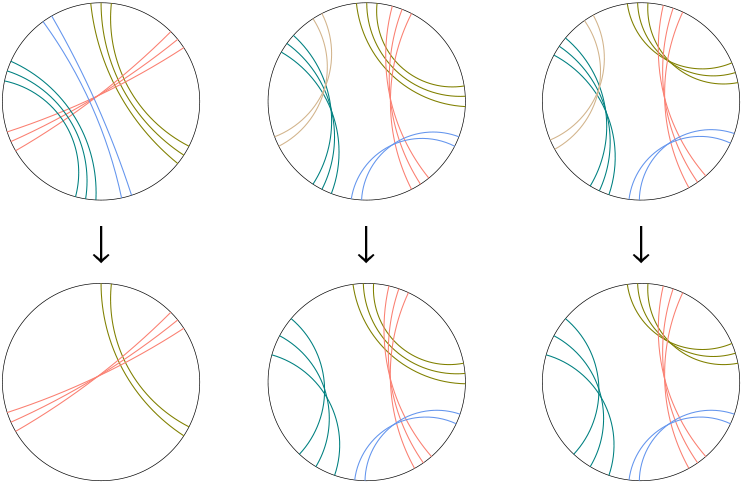
Circular diagrams for (upper row)) uncontracted and (lower row) contracted fully-flagged folds.

We shall see that contraction leads to a useful and manageable classification of fold topologies,. Note, however, that our topological approach ignores structural features of ssRNA folds, including hairpin strands, bulges, stem-loops and multibranched loops (2, 26). With the exception of so-called pseudoknotted folds, most currently known ssRNA folds are, from this topological perspective, trivial unfolds. However, that structural coarse-graining allows us to access topological aspects of folding and knotting, extending the reach of fold classification.

### Labelling folds

A useful alphanumeric label for a fold of the form 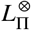, where *L* is a digit string and 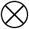 and Π denote characters appended to the string, can be deduced from its circular diagram as follows. First, index ribbons in the diagram by digits *i* = 1, 2 … and label the ribbon footers accordingly, giving a label *L* (introduced above) for the perimeter cycle around the circular diagram. For example, *L* = 123145246356 for the circular diagram in Fig. 3(a) and 124564315326 for the diagram in Fig. 4(c). Since the ribbon labelling is arbitrary, many distinct labels *L*_1_, *L*_2_,… encode the same fold. The canonical label 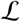 is that whose associated number *L_j_* is as the smallest. In practice, that amounts to assigning ribbon index ribbon 1 to the ribbon with the shortest span between its footings, and sequentially numbering successive ribbons in either a clockwise or an anticlockwise sense, such that the span of ribbon 2 is the next shortest, and so on. Often that ribbon numbering is not unique due to symmetry of the circular diagram, nevertheless a single canonical label 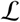 results. For example, the fold in Fig. 3 has canonical label 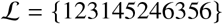. Likewise, the fold described by the polarised strand graph embedding in Fig. 4(a) has 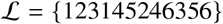.

Conversely, a digit string *L* with distinct digits *i* = 1, 2,…, *n* such that each digit is present twice somewhere in the string encodes a circular diagram, whose *n* ribbons are indexed 1, 2,… *n*. Any new label *L* induced by exchanging ribbon labels (e.g. *L* = 121332, 131223, 212331 etc.) gives the same fold. Further, normal or reverse cyclic permutation of *L* leaves the fold unchanged. The canonical label 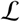 is the smallest number among all possible digit strings induced by exchanging ribbon labels, plus cyclic and reverse cyclic permutations. Thus, 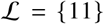 is the sole one-ribbon unflagged label, whereas there are 2 canonical labels with two ribbons, namely 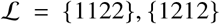, and 5 with three ribbons 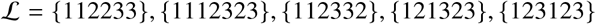 and so on. (For convenience, we include the canonical label {0}, denoting the trivial unfold with 0 ribbons.) The number of distinct canonical unflagged labels for folds with up to 5 ribbons are given in Table 1, along with the smallest and largest labels. All distinct unflagged canonical labels with up to *n* ≤ 5 are collected in Supporting Information (Table 7).

**Table 1:** Distinct unflagged canonical fold labels, 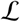 containing up to 5 duplexes.

| $n$ | *nmbr of labels* | *smallest $\mathcal{L}$* | *largest $\mathcal{L}$* |
| --- | --- | --- | --- |
| 0 | 1 | {0} | {0} |
| 1 | 1 | {11} | {11} |
| 2 | 2 | {1122} | {1212} |
| 3 | 5 | {112233} | {123123} |
| 4 | 17 | {11223344} | {12341234} |
| 5 | 79 | {1122334455} | {1234512345} |

**Table 2:** Contracted(*) one-colored genus-g semiflagged folds 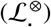, where *g* lies between 0 - 3. Folds are portrayed by their circular diagrams, canonical semiflagged fold labels, polarised strand graphs (whose vertices are labelled by ribbon index *i* within the fold label 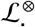, *i* = 1, 2,… *i*) and (one-colored) skeletons (whose edges are also labelled by *i*). All of the genus-2 and −3 folds contain three-way junctions only. (The genus-1 fold is junction-free.) These semi-flagged folds are ‘fully-duplexed’, with duplexed strands winding each edge of the skeleton, giving 3(*g* – 1) contracted double-helices in total. (*The strand graph and skeleton of the trivial unfold {0} shown in the first row are uncontracted; they vanish under contraction.)Plumblines marking the orientation of double-helical axe are drawn on each strand graph vertex by broken grey lines. The plumblines in oriented strand graphs are marked by dotted lines through vertices. Graph vertices are colored blue-gray and mustard for vertices hosting antiparallel and parallel double-helices respectively.

| | *diagram* | $\mathcal{L}_i^\otimes$ | *genus* | *strand graph\** | *skeleton\** |
| --- | --- | --- | --- | --- | --- |
| 1 | | $\{0\}$ | 0 | | |
| 2 | | $\{\bar{1}1\}$ | 1 | | |
| 3 | | $\{\bar{1}12\bar{3}32\}$ | 2 | | |
| 4 | | $\{\bar{1}21\bar{3}23\}$ | 2 | | |
| 5 | | $\{123123\}$ | 2 | | |
| 6 | | $\{\bar{1}123435\bar{6}6542\}$ | 3 | | |
| 7 | | $\{\bar{1}1234435\bar{6}652\}$ | 3 | | |
| 8 | | $\{\bar{1}12345364562\}$ | 3 | | |

| | *diagram* | $\mathcal{L}_\bullet^\otimes$ | *genus* | *strand graph* * | *skeleton* * |
| --- | --- | --- | --- | --- | --- |
| 9 | | $\{\bar{1}1234\bar{5}3\bar{6}4652\}$ | 3 | | |
| 10 | | $\{\bar{1}1234\bar{5}365462\}$ | 3 | | |
| 11 | | $\{\bar{1}2134\bar{5}23\bar{6}465\}$ | 3 | | |
| 12 | | $\{\bar{1}21345326456\}$ | 3 | | |
| 13 | | $\{\bar{1}2134\bar{5}326546\}$ | 3 | | |
| 14 | | $\{\bar{1}2314\bar{5}246\bar{3}56\}$ | 3 | | |
| 15 | | $\{\bar{1}23145364256\}$ | 3 | | |
| 16 | | $\{\bar{1}2314\bar{5}3\bar{6}4265\}$ | 3 | | |

**Table 3:** Extension of Table 2 to one-colored semi-flagged genus-4 folds with Y-junctions, all of whose duplexes are antiparallel. (Data courtesy of Neave Taylor.)

| | diagram | $\mathcal{L}_i^\otimes$ | genus | strand graph* | skeleton* |
| --- | --- | --- | --- | --- | --- |
| 17 |  | {123145235678649789} | 4 |  |  |
| 18 |  | {123145623678497589} | 4 |  |  |
| 19 |  | {123145647823895697} | 4 |  |  |
| 20 |  | {123145672378495869} | 4 |  |  |
| 21 |  | {123415637859724689} | 4 |  |  |

**Table 4:**
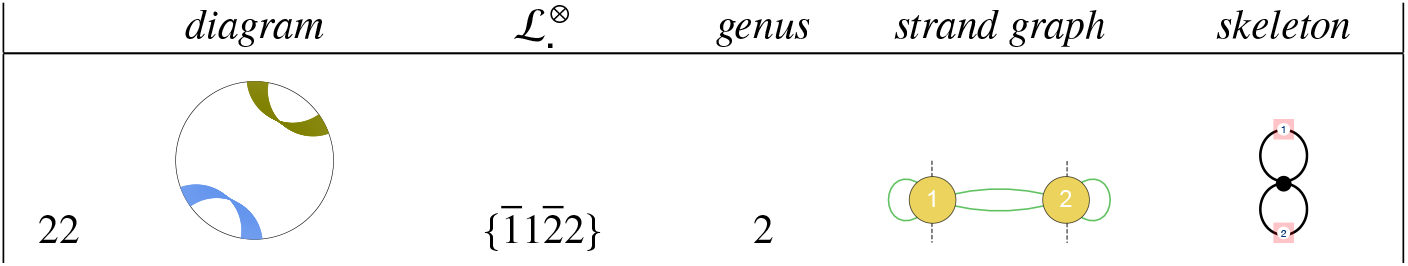
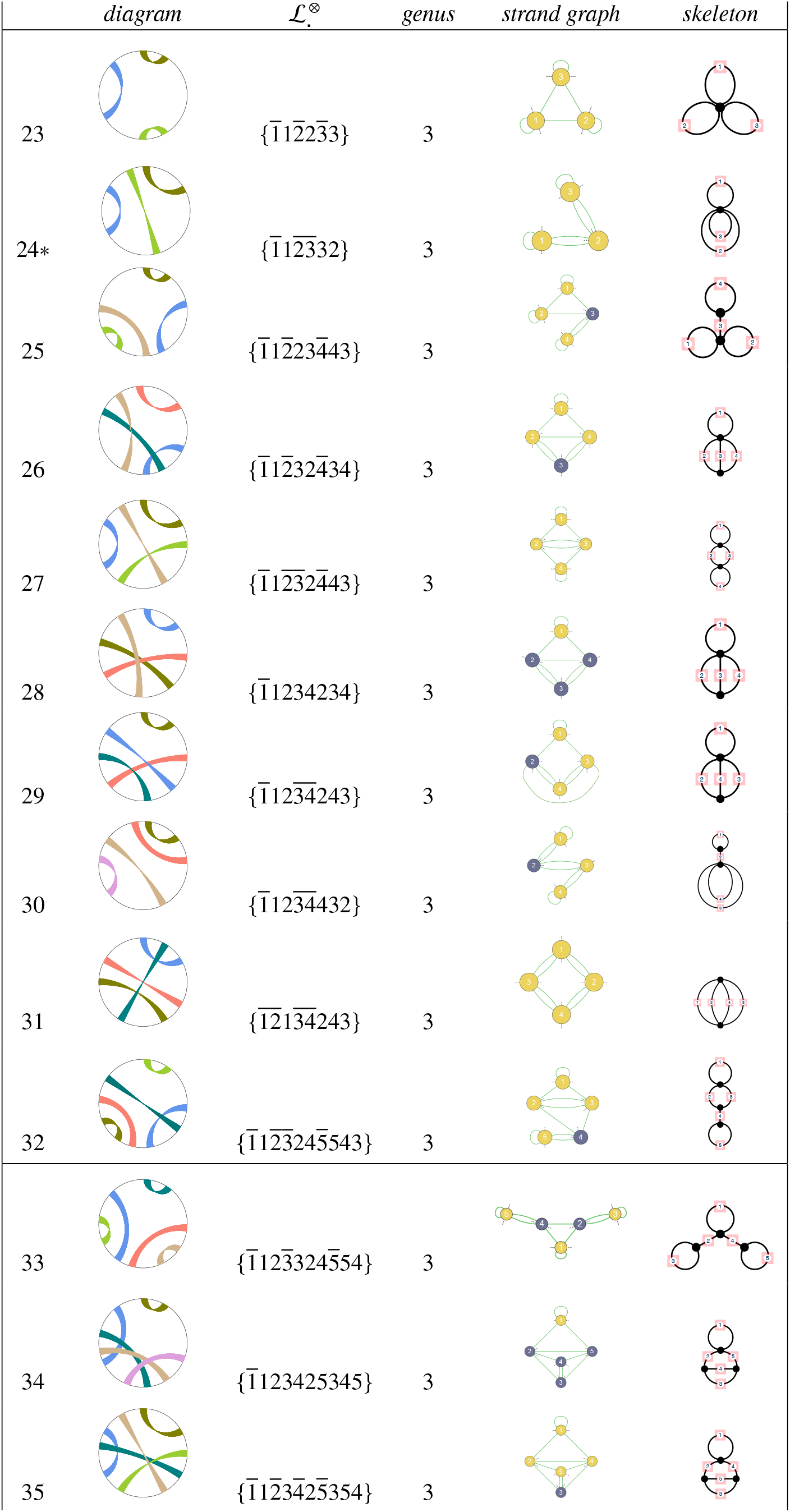
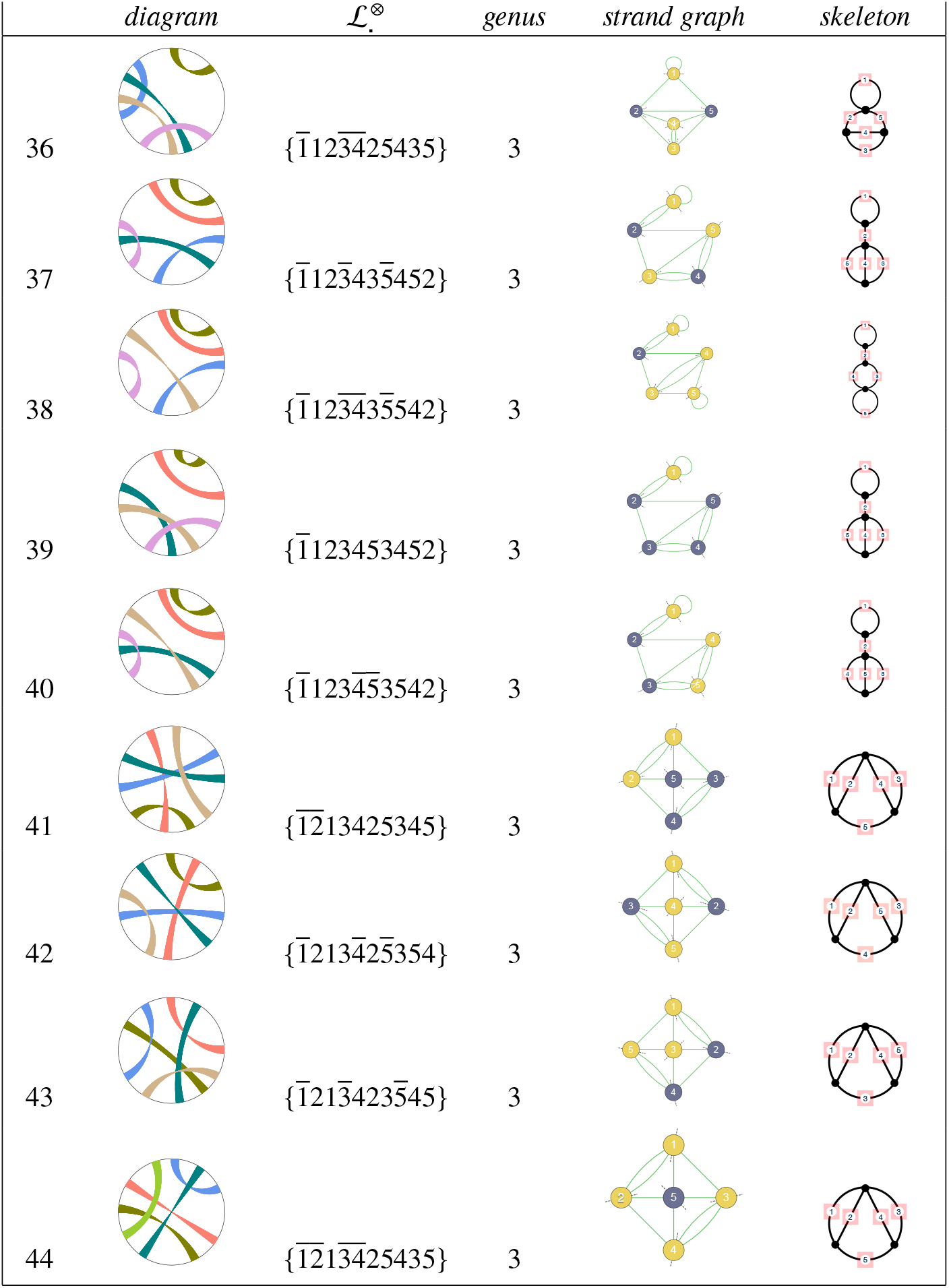
Circular diagrams and semi-flagged fold labels 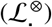 for contracted one-colored genus-g folds containing four-way (and higher) junctions, where g lies between 0 - 3. The folds are fully-duplexed, with a single duplex along each edge of their skeleton, like the folds in Table 2. The total number of contracted duplexes is < 3(*g* – 1), due to their higher-order junctions. (*This fold has the same unflagged label as the 112332 fold in Table 2. However, since it has unique semiflagged and flagged labels it is a different fold.)

**Table 5:** Classification of folds via minimum crossing numbers of one- or two-colored embeddings of their polarised strand graphs. Low-genus simplest folds, of type (a), are listed in Tables 2 and 3.

| *Fold features* | $N_{\times}$ (one-colored fold) | $N_{\times}$ (two-colored fold) |
| --- | --- | --- |
| (a) *Junctions folds* ds | 0 | $> 0$ |
| (b) *Meso-junction folds* ss+ds | $> 0$ | 0 |
| (c) *Crossed* (no junctions); ss+ds | $> 0$ | $> 0$ |

**Table 6:** Distribution of distinct semi-flagged folds whose duplexes are odd parity only 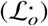 between folds whose polarised graph embeddings necessarily contain edge-crossings #(*N*_×_ > 0), and folds allowing uncrossed embeddings #(*N*_×_ = 0), for folds containing up to 5 duplexes (*cf*. Table 1).

| $n$ | $\#(N_x > 0)$ | $\#(N_x = 0)$ | $\mathcal{L}_o (N_x = 0)$ |
| --- | --- | --- | --- |
| 1 | 0 | 1 | $\{1_o 1\}$ |
| 2 | 1 | 1 | $\{1_o 1 2_o 2\}$ |
| 3 | 2 | 3 | $\{1_o 1 2_o 2 3_o 3\}, \{1_o 1 2_o 3_o 3 2\}, \{1_o 2_o 3_o 1 2 3\}$ |
| 4 | 12 | 5 | $\{1_o 1 2_o 2 3_o 3 4_o 4\}, \{1_o 1 2_o 2 3_o 4_o 4 3\}, \{1_o 1 2_o 3_o 4_o 2 3 4\}, \{1_o 1 2_o 3_o 4_o 4 3 2\}, \{1_o 2_o 3_o 1 4_o 3 2 4\}$ |
| 5 | 64 | 15 | $\{1_o 1 2_o 2 3_o 3 4_o 4 5_o 5\}, \{1_o 1 2_o 2 3_o 3 4_o 5_o 5 4\}, \{1_o 1 2_o 2 3_o 4_o 4 5_o 5 3\}, \{1_o 1 2_o 2 3_o 4_o 5_o 3 4 5\},$<br>$\{1_o 1 2_o 2 3_o 4_o 5_o 5 4 3\}, \{1_o 1 2_o 3_o 3 2 4_o 5_o 5 4\}, \{1_o 1 2_o 3_o 3 4_o 5_o 2 4 5\}, \{1_o 1 2_o 3_o 4_o 2 5_o 4 3 5\},$<br>$\{1_o 1 2_o 3_o 4_o 2 5_o 5 3 4\}, \{1_o 1 2_o 3_o 4_o 5_o 3 2 5 4\}, \{1_o 1 2_o 3_o 4_o 5_o 3 4 5 2\}, \{1_o 1 2_o 3_o 4_o 5_o 5 2 3 4\},$<br>$\{1_o 1 2_o 3_o 4_o 5_o 5 4 3 2\}, \{1_o 2_o 3_o 1 4_o 5_o 2 3 5 4\}, \{1_o 2_o 3_o 4_o 5_o 1 2 3 4 5\}$ |

**Table 7:**
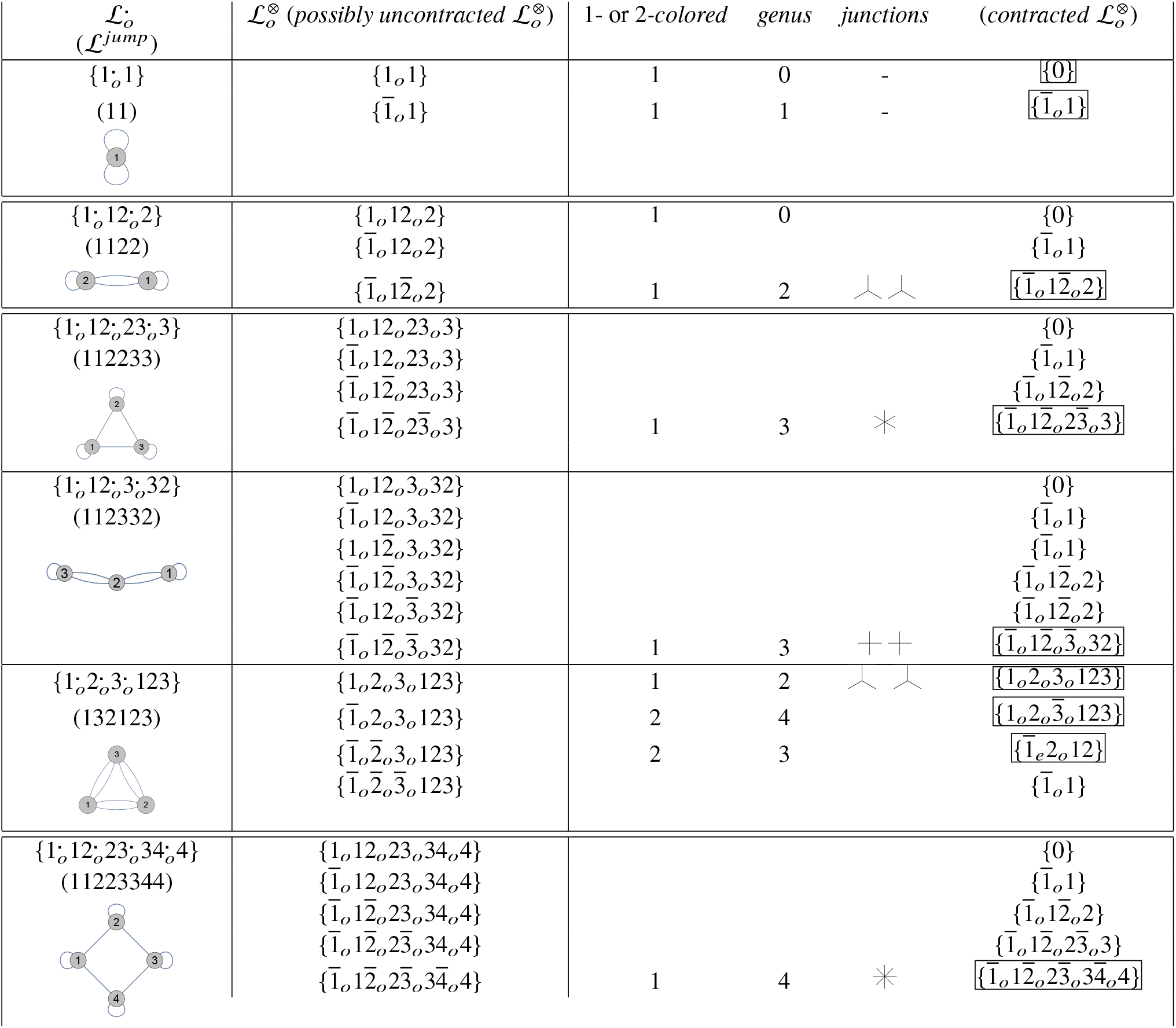
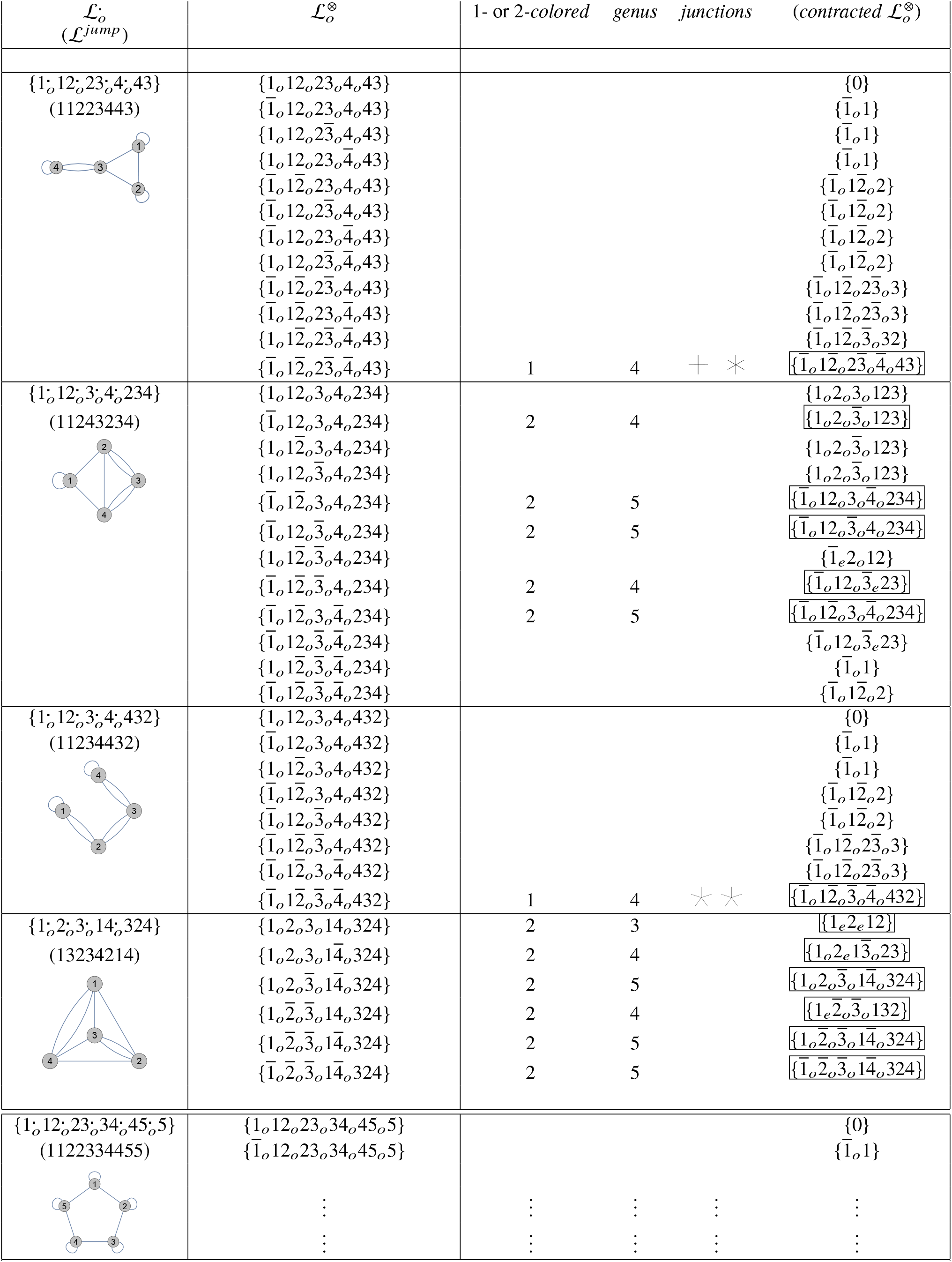
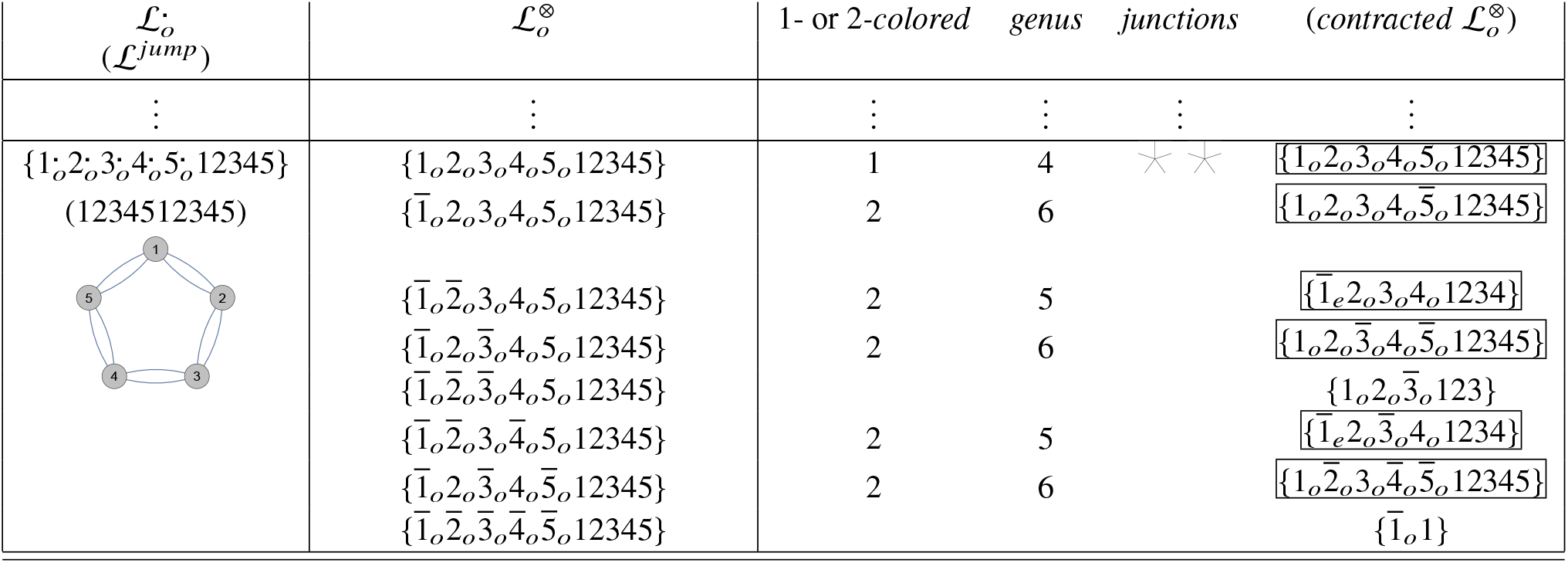
All semi-flagged folds with odd parities 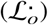 containing up to four ribbons, admitting planar unpolarised strand graph embeddings without edge-crossings, illustrated in the left-hand column. “Jump” paths within the fold, described in the text, are listed as 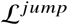. The fully-flagged folds 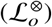 are generated by appending either annular or moebius orientation flags to 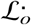, giving contracted canonical fold labels listed in the right-hand columns. Their crossing-free embeddings are wound on either one- or two-colored skeletons, whose genus and type are listed. One-colored skeletons induce folds with conventional three-, four-, five-, six- or eight-way junctions, denoted by graphics stars with the appropriate number of rays. Many folds are duplicates; the first occurrence of each fold is <u>boxed</u>. (Symmetrically equivalent folds to those tabulated - evident from the unpolarised graph symmetry - are ignored.)

The canonical label 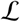 is ‘unflagged’, since it does not distinguish moebius from annular ribbons, nor does it encode the double-helical twists associated with each ribbon. Two additional flags are therefore needed to specify a generic duplexed fold: an ‘orientation’ flag 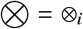, which specifies whether a duplex is parallel or antiparallel, and a ‘parity’ flag Π = *π_i_*, which is either even or odd, depending on the number of half-twists in the duplex^1^. Both flags are appended as superscripts and subscripts to the first occurrence of each digit *i* in the canonical bare label 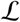 only, giving a canonical ‘fully-flagged’ fold label, 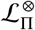. The subscripted flags, Π = *π*_1_*π*_2_ …, specify twist parities for each *i*. That flag is either even (*i_e_*) or odd (*i_o_*), depending on the number of half twists in the duplex. If ribbon *i* is annular, the digit *i* is left as is; if it is a moebius ribbon, the digit is flagged with a ‘superscript’ 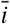. Thus, the one-duplex folds in Fig. 1(a,b,c,d) have fully-flagged fold labels 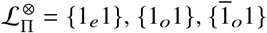 and 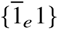 respectively.

It is useful to group folds by increasingly specific labels. The largest group contains all folds with common unflagged (bare) label 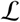, which we also denote 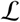, where the “.” super and subscripts indicate that both (orientation and parity) flags remain unspecified. That group includes two overlapping ‘semi-flagged’ labels: 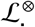 and 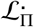, which collect folds with common bare label and orientation flags, or bare label and parity flags respectively. Thus, the folds in Figs. 1(a,b,c,d) have semiflagged labels 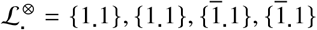 and 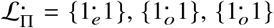 and 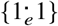 respectively. If both flags are specified, a smaller group of folds, with common fully-flagged label, 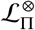, are possible. 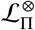 includes an infinite number of specific folds sharing the same twist parities Π = *π_i_* but different twist integers *t_i_* (odd or even as per the parity). Each fold in that group has a fully-flagged fold label with specific twists *t_i_* in each duplex *i*, denoted 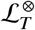. For example, 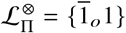 includes the folds 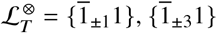, etc. and 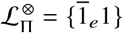 includes the folds 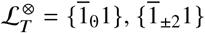, etc. Note that 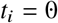^2^ ‘untwisted’ duplexes are distinct from unduplexed and untwisted double strands, since the former though untwisted, are duplexed by (typically less than 5) hydrogen-bonded nucleotides, whereas the latter, though untwisted, are also unbonded. Folds flagged by different twists 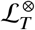 and 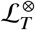, share common label 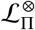, provided the parities of twists *T* and *T′*, match.

The contraction operation depends on the ribbon orientations in a fold’s circular diagram since the contraction rules are specific to annular and moebius ribbons. Therefore unflagged folds 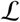 and semi-flagged folds 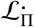 are neither contracted nor uncontracted. If orientation flags are filled, i.e. semi-flagged folds 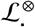 and fully-flagged folds, 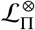, the fold can be classified as contracted or uncontracted. Appending orientation flags to unflagged labels 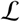 leads to uncontracted or contracted folds, following the trio of contraction rules listed above. For example, if a digit *i* is repeated in adjacent positions within the string, 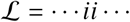 and it is annular, it is deleted. Similarly if *i, j, k*,… are all annular ribbons, palindromic subsequences 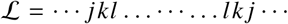 contract to 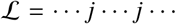 whereas if they are all moebius ribbons repeated subsequences 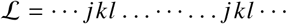 contract to 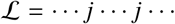.

The bare label 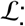 can be sorted by magnitude, giving a convenient rule for sorting unflagged folds. Additional rules for naming and sorting semi- and fully-flagged folds are given in the Supporting Information. To aid the reading of circular diagrams and their associated fold labels, distinct ribbons in the circular diagrams are generally colored differently and numbered 1, 2… clockwise around the perimeter of the diagram from ribbon 1, which has one footing located at 12 o’clock. We style the circular diagrams for semi-flagged and fully-flagged folds slightly differently. Ribbons in circular diagrams of semi-flagged folds are drawn as thick uncrossed or crossed bands, denoting annular and moebius ribbons respectively. (In the case of fully-flagged folds, each ribbon is denoted by (crossed or uncrossed) rulings. Even- and odd-parity flags are coded by two or three like-colored rulings respectively (e.g. Fig. 1, top row).)

#### Distinguishing folds

We define folds to be equivalent if and only if they share a common canonical fully-flagged fold label 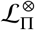. Equivalent folds therefore share the same polarised strand graph. Since that graph is dependent on twist parities (*π_i_*) rather than magnitudes (*t_i_*), duplexed folds with different labels 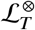 are classified as equivalent, provided all parity flags *π*_1_ *π*_2_… associated with their twists *T* = *t*_1_*t*_2_… are the same. Further, the definition allows any graph embedding of the polarised strand graph so that equivalent folds may have different number of crossed edges in the graph embedding. Thus, the embeddings in Fig. 3(c) and Fig. 4(a) describe the same fold.

Related folds with the same unflagged canonical label, 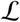, but different fully-flagged labels 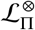 share common strand graph topology, disregarding plumbline-orientation (polarisation) and rigid-vertex ordering at each vertex. However, their polarised (rigid-vertex) strand graphs necessarily differ since they have distinct canonical fully-flagged labels. For example, the three-ribbon (contracted) folds in Fig. 7(a-d) have unflagged label 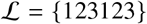 and strand graph topology shown in Fig. 7(e). But they have distinct flagged canonical labels 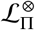 (listed in Fig. 7(a-d)) and polarised strand graphs, shown in Fig. 7(f-i). Further, folds with different unflagged canonical labels may also have the same strand graph topology, however they have different polarised strand graphs, due to distinct orientations of their plumblines. For example, the pair of folds in Fig. 8(a) and (d), with unflagged labels {12132434} and {12134234}, share the same strand graph topology shown in Fig. 8(b) and (e), but distinct polarised strand graphs (Fig. 8(c,f)).

**Figure 7:**
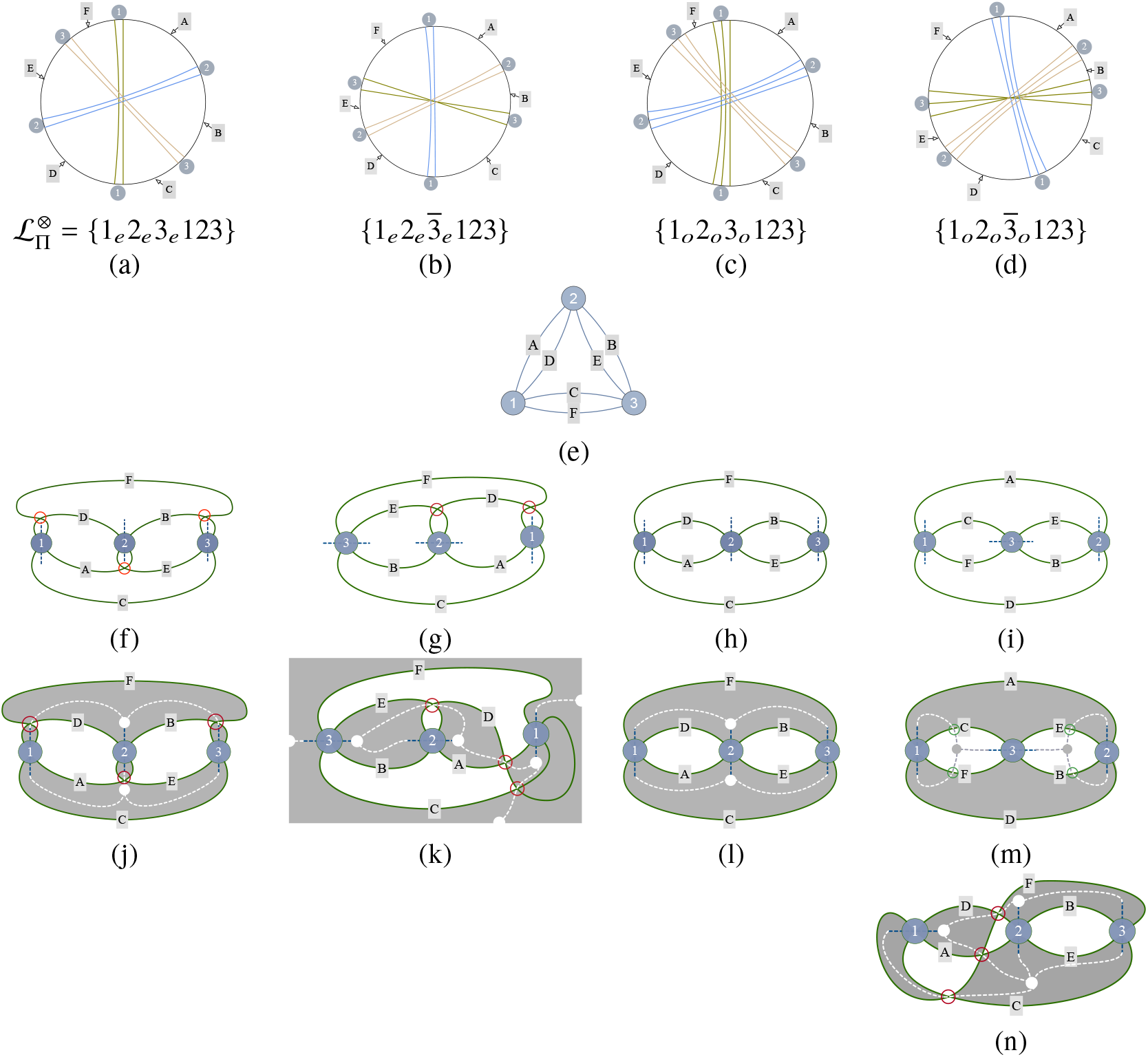
(a-d) Some three-ribbon folds with common unpolarised strand graph shown in (e). (f-i) The polarised strand graphs for (a-d) respectively. (j-l) Checkered coloring and one-colored skeletons of the one-colored polarised strand graphs from (f-h) respectively. The one-colored skeletons for these polarised strand graphs are marked by dashed arcs connecting vertex plumblines along skeletal edges, and filled circles at skeletal vertices. (Partial circles at the quadrilateral edges in (k) form a single vertex. Edge-crossings are marked by red circles.) (m) Checkered coloring of the crossing-free strand graph embedding in (i): plumblines though vertices 1, 2 are black, since they point into black domains, whereas the plumbline through 2 is white. The skeleton of this embedding is two-colored, some of whose edges cross from black to white domains (at sites circled in green). (n) An alternative embedding of the fold in (d), giving a one-colored skeleton with edge-crossings.

**Figure 8:**
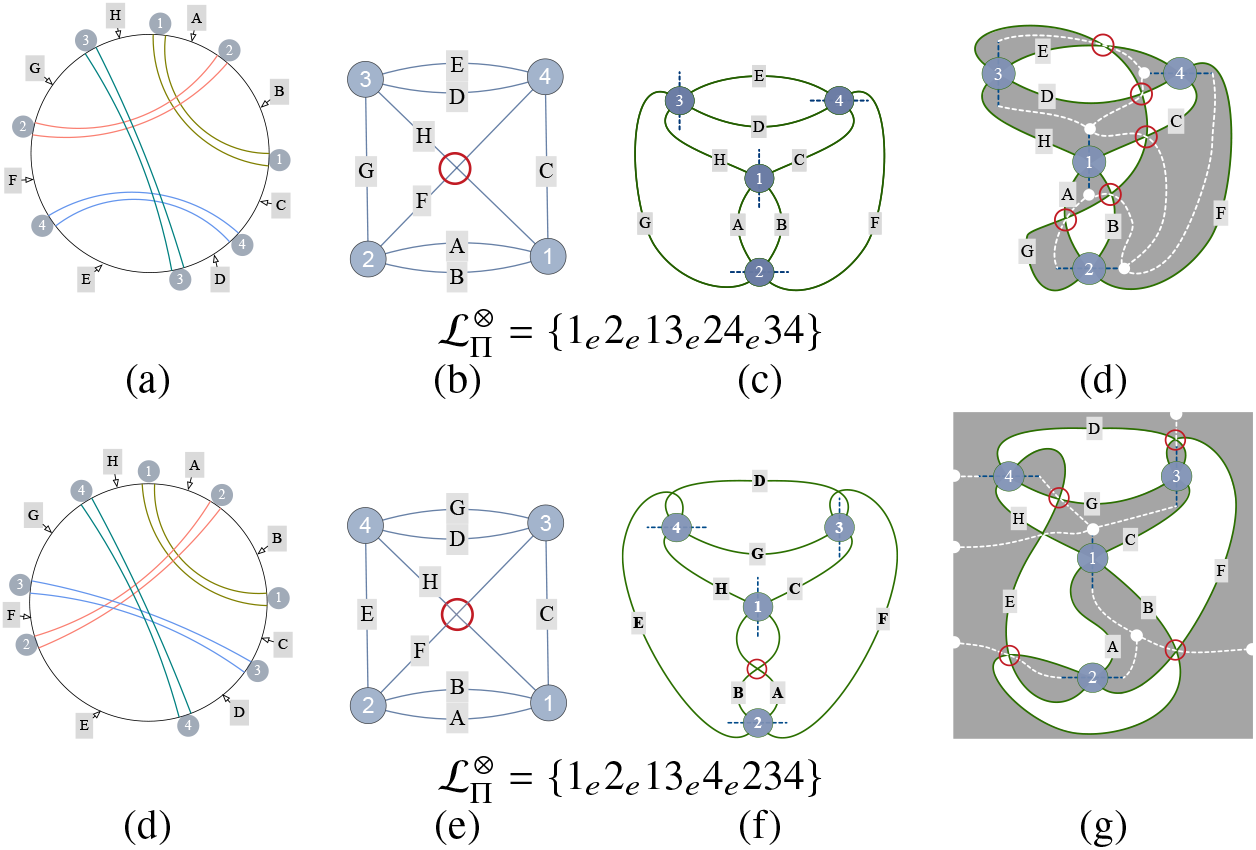
Circular diagrams (a) and (b) give isomorphic strand graphs, (b,e). However, their oriented rigid-vertex embeddings (c,f) are different. (d,g) The simplest one-colored embeddings of their polarised strand graphs give genus-four skeletons (dashed edges).

These classifications and sub-classifications of folds allow us to draw up a useful taxonomy of folds.

### One and two-colored skeletons of a fold embedding

The embeddings of polarised strand graphs in Figs. 7(f-i) and Figs. 8(c,f) can be projected onto the page, giving ‘shadow diagrams’ which are planar graphs with degree-4 vertices, induced by both strand graph vertices and edge-crossings. Those diagrams divide the page into closed inner domains, as well as a single external domain, extending to the edges of the page and bounded by the outermost edges of the graph. Alternate domains can be shaded, giving a checkered black and white pattern, as in a chessboard. Two complementary colorings are possible, depending on the initial black domain: choose that domain to include a plumbline. For example, the strand graph embeddings of folds 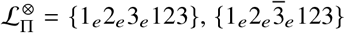 and {1_*o*_2_*o*_3_*o*_123} in Figs. 7(f), (g) and (h) give the checkered patterns in Figs. 7(j), (k) and (l) respectively. Since the plumbline of vertex 3 in Figs. 7(g) points into the external domain, that domain is black. We next construct a branched graph-like ‘skeleton’, characterising the topology of the black domains, by joining all plumblines, forming a network whose edges lie within black domains only. The resulting skeletons are drawn in Figs. 7(j), (k) and (l) as white dotted edges, meeting at degree-3 Y-junctions (indicated by white dots). The skeletons in Figs. 7(j) and (k) are topologically equivalent: both contain a pair of Y-junctions, joined by three edges, describing the so-called *θ*-graph. The skeleton in Fig. 7(k) includes three open edges radiating into the outer domain, terminated at the (arbitrary) rectangular boundary of the image. The distinction between the inner and outer domains is an artefact of the planar drawing: we remove that inequivalence by wrapping the drawing on to a sphere, such that the rectangular boundary is contracted to a single point. That operation ‘glues’ all radiating edges to a common (degree-3) vertex, so that the skeleton of this fold embedding is a graph with 4 degree-3 vertices, joined by 6 edges. Since the skeletal edges are confined to black domains in all three of the fold embeddings, they are classified as *one-colored* skeletons and their associated strand graph and fold embeddings as ‘one-colored embeddings’. Those embeddings can be realised by sequential winding of the strand around each edge of the one-colored skeleton, passing from one edge to the next according to the sequence labelled by 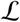, for appropriate labelling of the edges *i* = 1,…, *n*. In the completed winding, each skeletal edge hosts a duplex. Those one-colored fold embeddings will therefore be referred to as *fully-duplexed* folds. A detailed description of one-colored embeddings of a fully-duplexed fold, can be found in Supporting Information.

One-colored embeddings of folds typically include edge-crossings, visible in the embeddings of 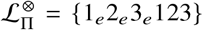, 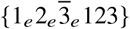 and 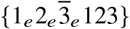 in Figs. 7(j) and (k). Indeed, generic embeddings include some edge-crossings. In some cases, folds can be embedded without any edge-crossings, such as the 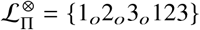 fold in Figs. 7(l). We call folds allowing uncrossed one-colored embeddings *relaxed*. More commonly, embeddings of the polarised strand graphs of folds which contain no edge-crossings result in *two-colored* embeddings, whose plumblines are located in both black and white domains. For example, the crossing-free embedding in Figs. 7(i) is two-colored, as shown in Fig. 7(m). However, that fold does have one-colored embeddings, such as that shown in Fig. 7(n), with three edge-crossings. Further, all one-colored embeddings of that fold 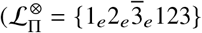 include edge-crossings. More generally, folds have multiple embeddings, due to the inherent flexibility of embedding polarised strand graphs. For example, the four-ribbon folds 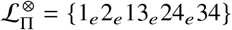 and 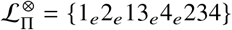 have two-colored embeddings, as shown in Figs. 8(c) and (f) respectively, or one-colored embeddings such as those in Figs. 8(d) and (g). In both cases, the number of edge-crossings changes between their one- and two-colored embeddings.

In fact, any fold has both one-colored and two-colored embeddings and one form can be switched to the other, introducing edge-crossings. The switch can be most simply effected by moving a single edge of the polarised graph embedding, illustrated in Fig. 9 for generic one- and two-colored regions of a fold embedding containing three-way junctions. Therefore switching between one- and two-colorings changes the number of edge-crossings in the planar strand graph embedding. The one- and two-colored embeddings in Figs. 9(a,c) respectively are uncrossed, whereas the two- and one-colored embeddings of the same fragment of the polarised grand graph in Figs. 9(b,d) have three edge-crossings. Further, that switch also affects the fold skeleton, since a new cycle is introduced into the skeleton to accommodate the new crossings (illustrated in Figs. 9(a,b)). In general, switching between one- and two-colored fold embeddings changes the number of edge-crossings and skeleton topology.

**Figure 9:**
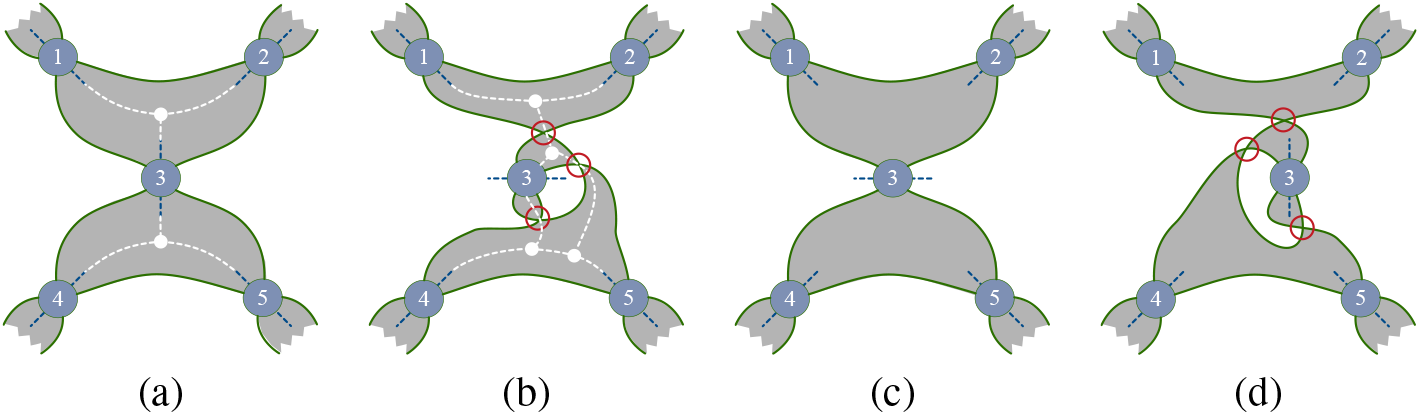
Switching between one- and two-colored embeddings around a generic vertex (3) separating a pair of Y-junctions (marked by white dots in (a,b)). (a,b) A crossing-free one-colored embedding and (b) two-colored embedding with three edge-crossings of the same polarised strand graph. (c) An uncrossed two-colored embedding and (d) one-colored embedding with three edge-crossings. (a,b) The fold skeletons are indicated by white dashed lines in (a,b); the skeleton in (b) contains an extra loop compared with that in (a), incrementing the genus by one.

There is one exception to that rule, specific to uncontracted folds containing hairpin bends. In those cases, the fold is uncontracted since its circular diagram includes isolated annular ribbons, which are deleted during contraction (via rule 3 above). Those cases are readily detected from their semi-flagged label, 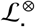, which includes repeated ribbon indices side-by-side, *viz*. 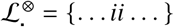. Such folds allow both one- and two-colored embeddings of their polarised strand graph without changing the number of edge-crossings (Fig. 10). That exception does not apply to contracted folds.

**Figure 10:**
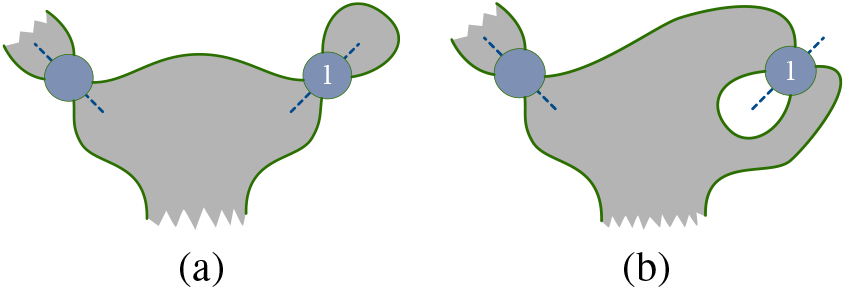
(a) A crossing-free, one-colored polarised strand graph embedding of a circular diagram containing a single - contractible - isolated annular ribbon (vertex 1). (b) A two-colored embedding of the same fold region, also uncrossed.

Recall that one-colored embeddings can be realised by winding the strand around the edges of the appropriate (one-colored) skeleton, such that every skeletal edge hosts local pair of duplexed strands. If the skeleton is two-colored, some of its edges are black at one end and white at the other, while others are monochromatic. For example, the two-colored embedding of the fold 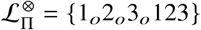 in Fig. 7(m) induces a two-colored ‘handcuff-graph’ skeleton, with two degree-3 vertices, both located in white domains, and three edges, two of which traverse both black and white domains while the third is monochromatic. Windings of the fold on two-colored skeletons differ from those on one-colored skeletons, since they necessarily include both ‘single-stranded’ (ss) and ‘double-stranded’, duplexed (ds) sections along the length of the closed strand. If a fold embedding is two-colored, only the monochromatic edges of the skeleton host ds regions; bichromatic edges contain a single unduplexed local fragment of the strand. Those embeddings are therefore classifed as *under-duplexed*.

These examples reveal the distinction between a *fold* and its *embedding*. We define a fold by its *topology*, characterised by fold label, circular diagram or polarised strand graph topology. The particular *geometry* of a fold is very flexible, and depends on the planar embedding of its polarised strand graph (though not its topology). Neither crossing-number of edges in planar projections of the polarised strand graph, skeleton topology, nor (one- or two-)coloring are fixed by the fold.

## RESULTS

Our study of duplexed folds deals with the following issues, dealt with in turn. First, we construct a catalogue of contracted folds of increasing topological complexity via one-colored embeddings. Alternative embeddings of those folds are explored, distinguishing between *crossed* and *uncrossed* embeddings. We then analyse entanglements of catalogued folds. Lastly, we apply those results to deduce the simplest knots (or pseudoknots) expected for simpler all-antiparallel folds made of a single loop of RNA, giving the simplest fold designs likely to form knotted single-stranded (ss)RNA.

### Constructing folds via planar skeletons

The previous section outlined the construction of a skeleton from a one- or two-colored fold embedding. Here, we construct a fold embedding from an arbitrary skeleton. One-colored embeddings of folds can be constructed on any (topologically planar) graph-like skeleton reticulating the sphere, containing *f* faces and *n* edges (*i* = 1, 2,…), as follows. Place a single-stranded loop in the interior of each face of the skeleton (e.g. Fig. 11(a)). Since each skeletal edge bounds two adjacent faces, there are a pair of strands along each edge, and *f* disjoint strand loops. Join adjacent loops straddling an edge *a* with a bridging strand, and replace the resulting H-shaped strand fragment in the neighbourhood of that bridge by an X-shaped strand crossing, reducing the number of disjoint strand loops to *f* – 1. Add additional bridges crossing arbitrary edges, until a single continuous strand results by bridging edges 1, 2,… (Figs. 11(b,c)). Deform the strand so that it forms pairs running along either side of each edge and intercalate a degree-4 vertex along each edge between the pairs of strands, corresponding to a double-helix along each edge. If an edge *i* is crossed by a bridge, it hosts a duplex with twist *t_i_* = ±1, if it is unbridged, the uncrossed parallel strands describe a duplex of twist *t_i_* = 0. The embedding of the polarised strand graph is unchanged if all edges with unit twist are replaced by duplexes whose twists are of odd parity, i.e. *t_i_* = o, where *o* denotes any odd integer and edges with zero twist are replaced by duplexes of even parity, *t_i_* = *e* (Figs. 11(d,e)). The orientation flags for each vertex (annular or moebius) can be determined by orienting the strand, giving locally antiparallel or parallel strand pairs along each edge. (Edges 1 - 6 in the skeleton in Fig. 11(d) and (e), have twist parities *oeeeoo* and *oeeooo* respectively.) The orientation and parity flags determine the circular diagrams and (canonical) fully-flagged fold labels 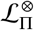 (Figs. 11 (f,g)) from which the polarised strand graphs can be reconstructed, as explained in detail above. Since the edges of the skeleton are coaxial with the plumblines of all duplexes, the construction generates one-colored embeddings.

**Figure 11:**
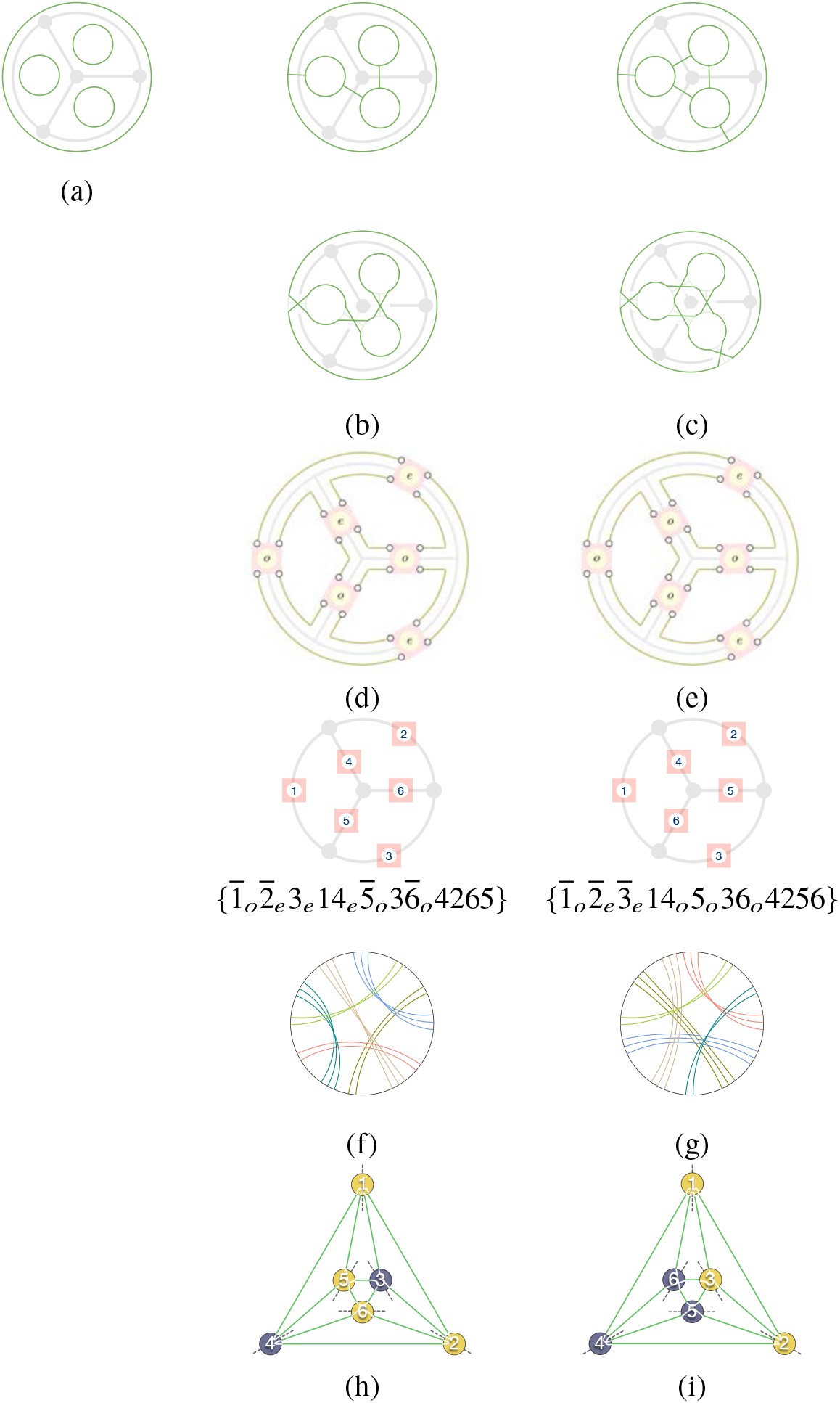
Construction of one-colored embeddings of fully-flagged folds (labelled 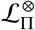) on the genus-3 wheel-like skeleton. (a) (Green) strand loops in each face of the graph. (b,c) Two possible sets of bridges giving one-component graphs, redrawn with bridges replaced by crossed edges to form one-component strands. (d,e) Schematic strand configurations, with duplexed strands on each edge of the wheel, whose parities are odd (*o*) along bridged edges, and even (e) otherwise. (h,i) Skeleton edge labelling corresponding to canonical fold labels and circular diagrams. (f,g) Circular diagrams for those uncrossed folds. (h,i) Crossing-free polarised strand graph embeddings.

In general, different arrangements of bridges on the skeleton can lead to a single continuous strand, giving more than one relaxed fold for any skeleton. For example, the bridging patterns on the wheel-like skeleton in Fig. 11(b,c) induce a pair of folds whose circular diagrams in Fig. 11(f,g) have distinct canonical labels, so those folds are different. In fact, the same skeleton hosts a third relaxed fold (with twist parities *π*_1_ – *π*_6_ = *oeeoeo*): a specific example of that fold is illustrated in Fig. 3. That multiplicity depends on the planar skeleton; some allow just one distinct fold.

That construction can be generalised to any planar skeleton, allowing folds to be enumerated via one-colored windings on different skeletons. Folds can also be sorted by the topological complexity of their underlying one-colored skeleton. We characterise skeleton topology by its cyclomatic number, 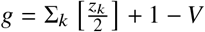, where the skeleton contains *V* vertices and each vertex *k* has degree *z_k_*. The average degree of a skeleton, < *z* >, defined by 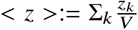, will also be useful. The number of vertices (*V*) and edges (*E*) in a generic skeleton is related to the genus *g* and < *z* > by

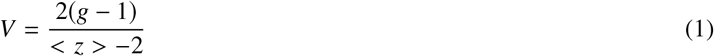

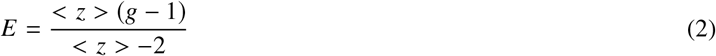

That number is the same as the *genus* of the closed (oriented and boundary-free) surface formed by ‘tubifying the skeleton’. (Tubification can be realised by inflating each edge in the skeleton to form a cylindrical tubule, then merging tubules whose underlying skeletal edges share a common vertex.) From here on, we equate the skeleton with its tubified surface and refer to the cyclomatic number of the skeleton and its associated fold embedding as its *genus*.

A genus-0 skeleton with just one edge bounded by a pair of (*z* = 1) vertices can host a closed loop with a bridge across the skeletal edge, or remain unbridged, admitting a single antiparallel duplex with odd or even parity respectively with labels {1_*o*_ 1} or {1_*e*_ 1} (Fig. 12(a,b)). Both {1_*o*_ 1} and {1_*e*_ 1} folds are uncontracted; they contract to the ‘unfold’, denoted {0} (Fig. 5(c)). The genus-1 skeleton has a single (degree-2) vertex and a single edge. Just one contracted fold is possible on this skeleton: 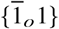 (Fig. 12(c)). (This fold includes, for example, the contracted 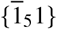 fold illustrated in Fig. 5(f)). Those genus-0 and −1 skeletons give junction-free folds, with at most one contracted duplex, since the skeletons have vertices of degree-1 or −2 only.

**Figure 12:**
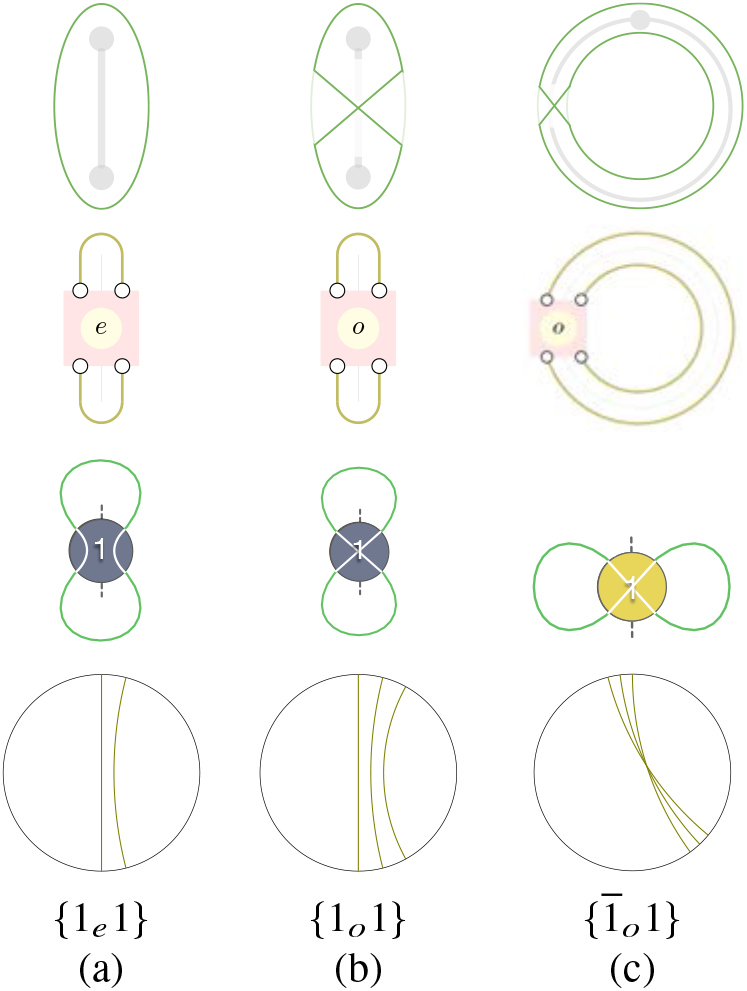
Constructions of one-colored embeddings of fully-flagged folds on (a,b) genus-0 and (c) genus-1 skeletons (*cf*. Fig. 11). (Top row) (a,b) Genus-0 and (c) genus-1 skeletons drawn in grey, overlayed by green loops which show all admissible parities of single-strand windings around the skeletons: (a) even twist parity and (b,c) odd twist parity. (An even parity winding on the skeleton in (c) induces two loops, rather than a single strand.). (Second row) A schematic illustration of all folds consistent with those parities. The strand (shown in green) enters and exits a central ‘switch box’ which twists the local strand pair an arbitrary even (*e*) or odd (*o*) number of half-twists, inducing a double helix with *t* crossings. (Third row) The corresponding crossing-free polarised strand graphs: blue and yellow vertices represent antiparallel and parallel duplexes respectively. Even parity twists are marked by uncrossed (white) strands within the vertex, odd parities by crossed strands. (Bottom row) Circular diagrams and fully-flagged fold labels.

The simplest higher-genus skeletons are 3-regular graphs (with trivalent vertices), such as the genus-3 wheel skeleton in Figs. 3 and 11. There are seven 3-regular graphs or genus-2 and −3 (27), each hosting one-colored skeletons and fully-duplexed folds with three-way (Y-) junctions. Analogous constructions to those in Figs. 11 and 12 on each of those seven skeletons leads to 21 distinct semi-flagged folds (with distinct labels 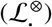) on genus-0, −1, −2 and −3 skeletons, tabulated in Table 2. Each entry in the Table describes a fold which can be made by winding a (possibly knotted) loop around the edges of its underyling skeleton which induces double-helices along each edge.

The catalogue of 21 distinct contracted one-colored semi-flagged folds 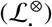 conflates a larger number of fully-flagged folds, formed by appending arbitrary twist parities to the label 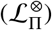. Some combinations of parity flags Π = *π*_1_ *π*_2_… *π_n_* induce relaxed folds, defined above. In some cases, just one set of parity flags (*π*_1_ *π*_2_…) allow relaxed fully-flagged folds; all other flags result in edge-crossings in the polarised strand graph. For example among all folds sharing the semi-flagged label 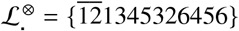, just one fully-flagged fold is relaxed: 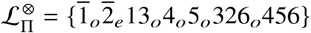, with parity flags Π = *oeoooo* (see Fig. 13(a)). In other cases, a number of fully-flagged folds sharing common semi-flagged label 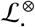 are relaxed (e.g. Fig. 13(b)).

**Figure 13:**
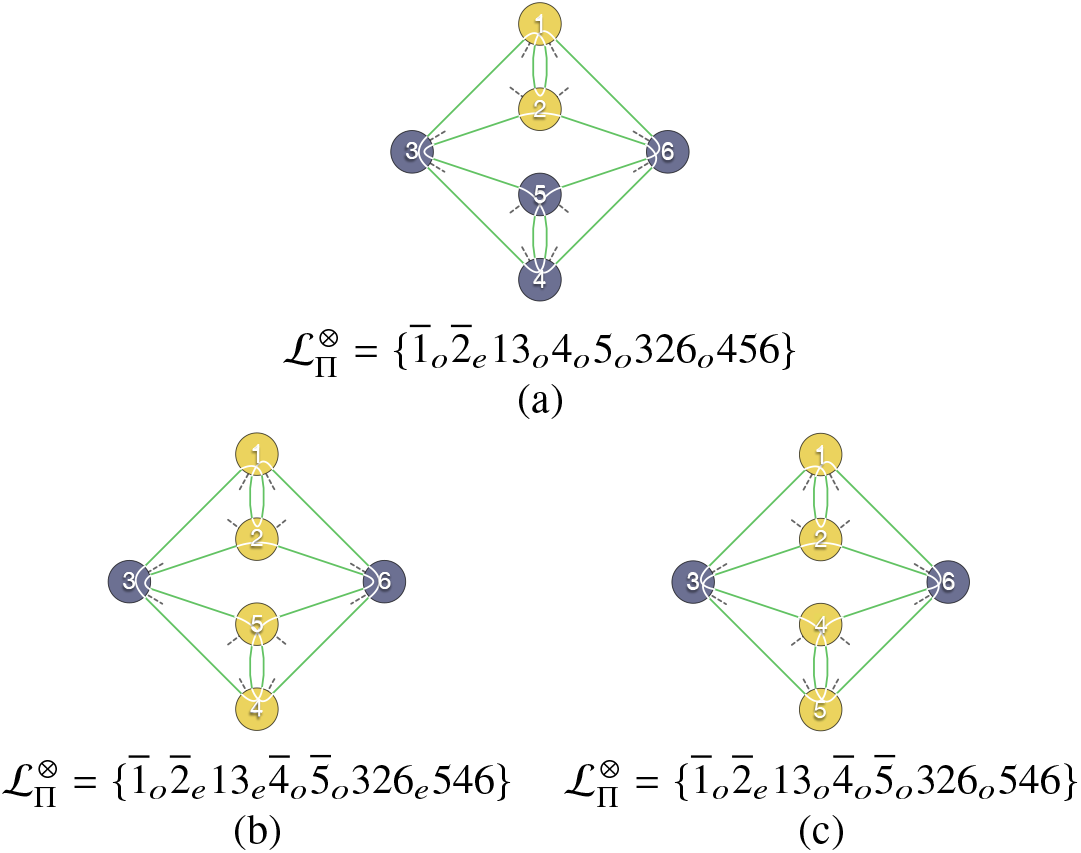
The genus-3 semi-flagged fold 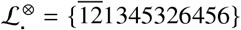 (fold 12 in Table 2) has just one suite of relaxed parities, *π*_1_ – *π*_6_ = *oeeooo*. (a) Schematic view of the uncrossed fold, showing the closed loop composed of green arcs between graph vertices and white segments within graph vertices. The genus-3 semi-flagged fold 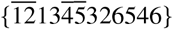 (fold 13 in Table 2) has relaxed parities, *π*_1_ – *π*_6_ = *oeeooe* or oeoooo, giving the pair of uncrossed folds whose polarised strand graphs are shown in (b) and (c) respectively.

In total, 41 distinct fully-flagged contracted folds are relaxed. (The folds are collected in Table 1 in Supporting Information.) Recall that relaxed folds can be embedded particularly simply as skeletal windings on one-colored skeletons (whose junctions are, et most, degree-3), forming fully-duplexed folds, devoid of edge-crossings except within those duplexes. All fold junctions, located at vertices of the skeleton, are conventional Y-junctions (28), so they can be realised such that every nucleotide is duplexed. In practice, they may be interrupted by bulges of unduplexed strands, but their contracted forms have uncrossed one-colored embeddings.

3-regular skeletons whose genus exceeds 3 include nonplanar graphs, such as the genus-4 ‘utility’ graph (the bipartite graph *K*_3,3_). Those cases too can be analysed (and will be explored more generally later in the paper). To allow a manageable search, we restrict our enumeration of one-colored folds with *g* > 3 to those wound on skeletons corresponding to the four planar 3-regular genus-4 graphs only. More than 150 uncrossed one-colored Y-junction folds are formed from those genus-4 skeletons, including five distinct semi-flagged folds, whose duplexes are exclusively antiparallel, collected in Table 3^3^. (These five cases lead to eight fully-flagged folds, collected in Table *S*2, Supporting Information.)

#### Prime and composite folds

Some of the folds in Table 2 are ‘composite’ (21), since their circular diagrams can be formed by joining simpler (‘prime’) diagrams. Conversely, if an arc can be drawn within a circular diagram which does not intersect ribbons, splitting a composite diagram along this arc ‘factorises’ the fold into composite folds, analogous to composite knots *K*_1_#*K*_2_ formed as connected sums of prime knots *K*_1_ and *K*_2_ (29). Composite folds with Y-junctions are wound on (3-regular) skeletons with at least one cut-edge, whose deletion leads to a two-component skeleton. For example, the contracted fold #6 in Table 2 is a composite of five prime folds, as shown in Fig. 14. Folds #3, 6, 7, 8, 9, 10 in Table 2 are composite.

**Figure 14:**
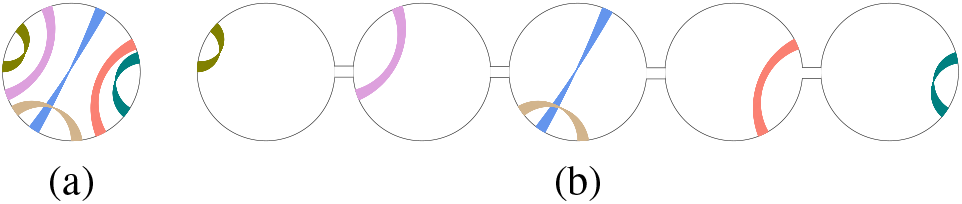
Reducing the composite (or reducible) semi-flagged fold (a) 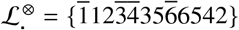 (fold #6, Table 2), to its five prime semi-flagged folds: (b) 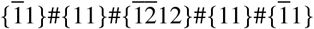.

#### Beyond Y-junctions

A new suite of folds can be constructed by allowing windings on skeletons which include higher-degree (*z*) vertices, where *z* > 3. As before, the simplest windings are those which induce double-helices around each edge of the skeleton (i.e. fully-duplexed). The resulting folds include four-way X-junctions (and possibly higher-order junctions also). All such folds are related to the Y-junction folds, since higher-degree vertex can be split into multiple Y-junctions intercalated with new edges. For example, any four-way (X-) junction in the original skeleton can be split into a pair of Y-junctions linked by a new edge (shown in Fig. 15(a,b)), 5-way junctions into three Y-junctions with two new edges, and so on. Although vertex-splitting changes the skeleton type, its genus remains unchanged. This vertex-splitting operation transforms a fully-duplexed winding on the higher-degree skeleton (*S*_X_) to an under-duplexed winding on the related skeleton with Y-junctions (*S*_Y_), with double-helices wound around all edges except those new edges created by splitting higher-degree vertices (see Figs. 15(a,b)). A pair of parallel and unduplexed single-strands run along those new edges. Since those parallel strands can be duplexed without affecting the winding, the same skeleton *S*_Y_ also hosts a fully-duplexed winding, whose double-helices along the new edges are untwisted (*t_i_* = 0), illustrated in Fig. 15(c). Accordingly, those duplexes must have even parities: if it is odd, the inverse construction, building an X-junction by shrinking a skeletal edge and merging adjacent Y-vertices, is blocked by an edge-crossing (see Figs. 15(d,e)).

**Figure 15:**
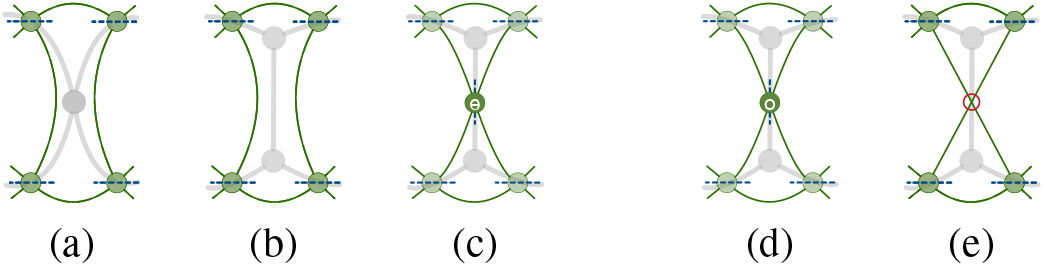
(a) shows the polarised strand graph of a fully-duplexed fold in the vicinity of an X-junction in a fold skeleton (drawn in grey in (a)). (b,c) The X-junction can be split into a pair of Y-junctions, joined by a new vertical edge in the fold skeleton. (b) The new edge hosts an unduplexed pair of parallel strands. (c) Duplexing those strands gives a fully-duplexed fold on the new skeleton. (d) A fully-duplexed fold on a skeleton with Y-junctions, whose edge hosts a double-helix with odd parity. (e) Unduplexing that double-helix leaves the polarised strand graph embedding with a single edge-crossing (at the least; any odd number of crossings is possible).

Provided the genus of *S_X_* is less than 4, the new fully-duplexed winding on *S*_Y_ must be listed in Table 2, since the genus of *S*_Y_ is equal to that of *S*_X_. It follows that all circular diagrams of folds on higher-order skeletons (*S*_X_) with *g* ≤ 3 can be found by deleting selected ribbons from a circular diagram in Table 2. Recall that each ribbon in the circular diagrams of fully-duplexed windings in Table 2 encodes a double-helical winding on some edge in the skeleton S_Y_. The deleted ribbons are those which encode (even parity!) duplexes on precisely those edges which are lost in creating the higher-order skeleton *S*_X_ from *S*_Y_, realised by shrinking edges and fusing vertices at either end. In summary, deletion of any subset of even-parity ribbons within the original diagram gives an uncrossed, double-stranded higher-order junction fold. Since even parity flags are required on specific ribbons, the catalog of fully-duplexed folds on higher-degree skeletons is therefore generated from subdiagrams of the expanded catalog of *fully*-flagged relaxed folds (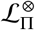, Table *S*1, Supporting Information), rather than the catalog (of semi-flagged folds 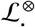) in Table 2. The catalog of all semi-flagged folds which can be realised as fully-duplexed windings on higher-degree skeletons is formed by stripping all parity flags from the higher-order fully-flagged folds. A simple example, giving a fold with a single X-junction, is shown in Fig. 16. (A worked example, giving all fully-duplexed folds on higher-degree skeletons from a single fold in Table 2 is given in Tables *S*3 and *S*4, Supporting Information.)

**Figure 16:**
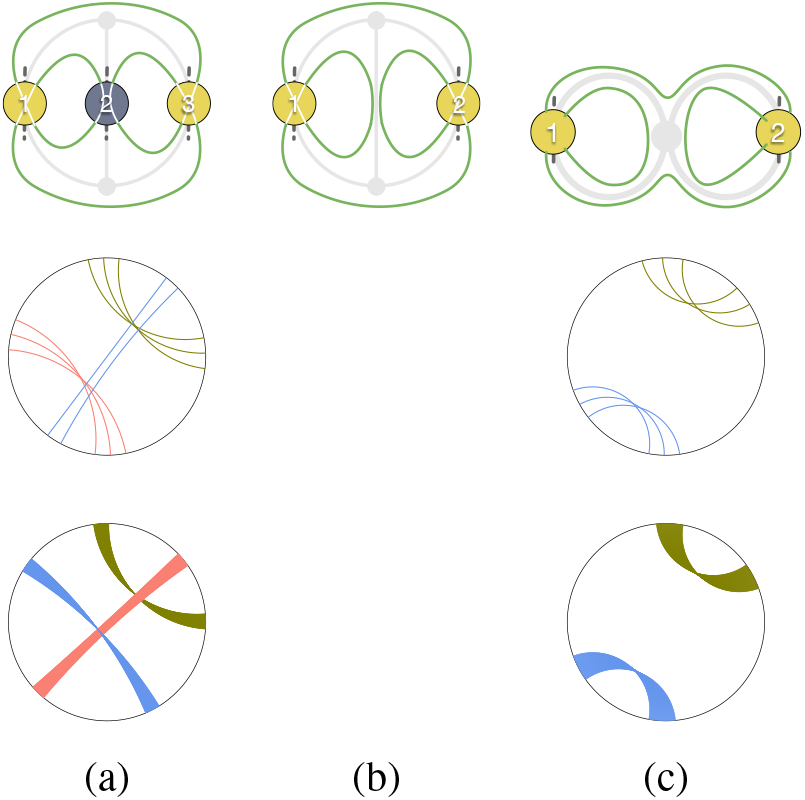
(Upper row) (a) A relaxed fully-flagged fold 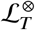 with fully-flagged label 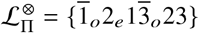, formed by assigning explicit twist *t*_2_ = 0 to the even-parity ribbon 2. (White edges within each graph vertex describe the simplest strand crossings in the vertex.) (b) The under-duplexed 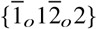 fold formed by denaturing the untwisted strand pair aligned with the central vertical edge of the skeleton. (c) A fully-duplexed fold induced by shrinking the central edge and fusing the pair of vertices at either end, forming a single X-junction. (Middle row) (a,c) Circular diagram for a relaxed fully-flagged fold 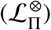 wound (a) on the *θ* graph and (c) on the bouquet graph *B*_2_, with a single X-junction. (Lower row) (a,c) Circular diagram for the semi-flagged folds 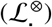.

Admissible circular diagrams of these contracted fully-duplexed folds on higher-degree skeletons, *S*_Y_ and their related skeletons *S*_Y_, are listed exhaustively in Table *S*5, Supporting Information. In some cases, the circular diagrams which result by ribbon deletions are uncontracted and contraction leads to relaxed folds whose genus is less than that of the precursor relaxed Y-junction fold. In all, 23 new contracted semi-flagged folds emerge. All of those can be formed by winding a loop around edges of (*g* ≤ 3) skeletons with higher-order junctions, listed in Table 4. Those 23 semi-flagged folds lead to 40 relaxed fully-flagged folds (listed in detail in Table *S*6, Supporting Information).

#### Under-duplexed folds

The folds in Tables 2, 3 and 4 are constrained to be fully-duplexed, with a single contracted duplex on each edge of their one-colored skeletons, giving 3(*g* – 1) double-helices. Those folds maximise the number of duplexes on each skeleton, since no more than one contracted duplex per edge is possible. In contrast, if an odd-parity duplex (with *t_i_* = 1,3,… twists) within an uncrossed embedding of a Y-junction fold is unduplexed, at least one edge-crossing must remain on that edge and the skeleton remains unchanged. A new ‘under-duplexed’ fold results, with a deficit of duplexes (< 3(*g* – 1)). Under-duplexed folds can therefore be wound on the same skeleton as their parent saturated fold, traversing all edges of the skeleton twice, according to the unflagged integer sequence of the parent fold 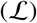. However, since strand pairs along some edges of the skeleton are not duplexed in the under-duplexed fold, it is no longer exclusively double-stranded, and includes a pair of single strands for every ‘denatured’ odd-parity duplex.

It follows that the complete catalog of circular subdiagrams given by deleting any subset of ribbons from any circular diagram in Table 2 includes under-duplexed folds, as well as the fully-duplexed folds listed in Table 4. The latter result by deleting even-parity ribbons, the former by deleting odd-parity ribbons. Consider for example the folds formed by deleting a single ribbon from any one of the three-ribbon folds listed in Table 2 (fold numbers 3,4,5), namely 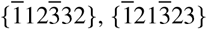 and {123123}. Provisionally label the unduplexed fold by an edited parent fold label, with ribbon *i* crossed-out (i) if that ribbon is deleted in the circular diagram. The two-ribbon folds 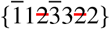 (ex fold 3) and 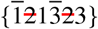 (ex fold 4) both give canonical label 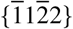. Similarly, the fold 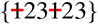 (ex fold 5) has canonical semiflagged label {1212} and 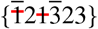 (ex fold 4) has canonical label 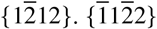 is listed in Table 4 as a fully-duplexed fold with a single X-junction. The remaining contracted folds, {1212} and 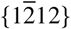, are under-duplexed, therefore necessarily include both double-stranded and single-stranded sections.

In some cases, under-duplexed folds due to ribbon deletions are uncontracted and subsequent contraction results in a fully-duplexed fold. For example, the fold formed by deleting a single ribbon from the genus-2 three-ribbon fold 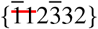 contracts to the one-ribbon fold 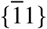, rather than a two-ribbon fold. Likewise, folds formed by deleting four ribbons from the genus-3 six-ribbon folds 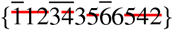 (ex fold 6) and 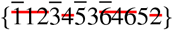 (ex fold 9) with the same uncontracted label 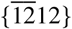, also contract to the one-ribbon fully-duplexed fold 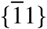.

### Fold embeddings and edge-crossings

Any contracted fold containing up to six distinct ribbons (*i* = 1,…, *n*, where *n* ≤ 6) whose semi-flagged label 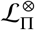 is missing from Table 2 can be embedded as an under-duplexed fold with Y-junctions. Some of those, listed in Table 4, can be embedded as fully-duplexed folds containing higher-order X- (etc.) junctions; most cannot. For example, the semi-flagged fold 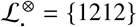 is not included in Tables 2 or 4. Since its circular diagram consists of a pair of crossed annular ribbons, any fold - of arbitrary genus - whose circular diagram includes the circular diagram characteristic of {1212} (a pair of crossed annular ribbons) can be used as a template for a fold embedding. The resulting embedding is realised by first embedding the fold with additional ribbons as usual and then unduplexing the double-helices corresponding to those extraneous ribbons in the more complex circular diagram which are not one of the the pair of crossed ribbons, retaining only two duplexes corresponding to the pair of crossed annular ribbons. Unduplexed helcies of even parity give uncrossed parallel single-strands, whereas unduplexed helices of odd parities leave a pair of crossed single-strands (i.e. equivalent to duplexes of twists *t_i_* = 0 and *t_i_* = ±1 respectively). Circular diagrams of the genus-2 fold {123123} (fold 5) and the genus-3 folds 8, 11, 12, 14, 15 and 16 contain that crossed annular ribbon motif (*cf*. Table 2). Some one-colored embeddings of the {1212} fold formed by unduplexing those folds are shown in Fig. 17. The number of skeletal edges on those skeletons is (3(*g* - 1)): since *i* = 1, 2 in the {1212} fold only two are duplexed, so generic higher-genus folds can be wound on skeletons such that 2 skeletal edges host duplexes and the remaining 3*g* – 5 host pairs of strands, which are both ‘single-stranded’ (as opposed to duplexed, or ‘double-stranded’).

**Figure 17:**
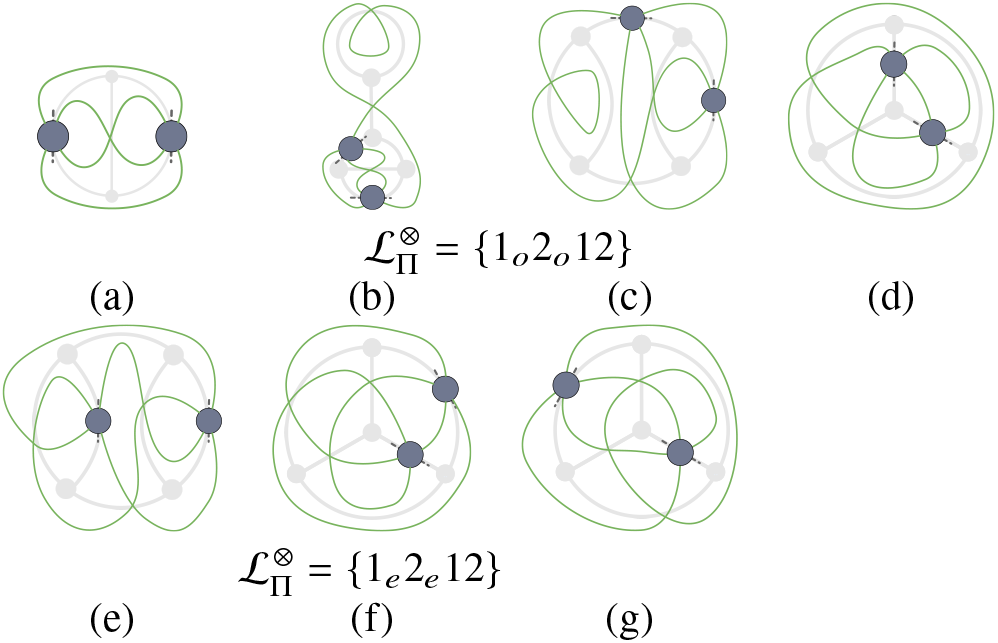
Some one-colored polarised strand graph embeddings on different skeletons of folds sharing semiflagged label 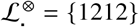. All fold skeletons are traced faintly in grey between pairs of green edges of the polarised strand graphs. (a-c) Embeddings of the fully-flagged 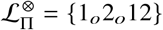 fold (a) ex fold 5 in Table 2, (b) ex fold 8, (c) ex fold 12 and (d) ex fold 15. (e-g) Embeddings of the fully-flagged 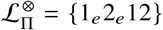 fold (e) ex fold 11, (f) ex fold 14, and (g) ex fold 16. Since the fold includes just 2 duplexes (labelled 1 and 2 all of these fold embeddings are under-duplexed.

Note also that the folds in Tables 2 and 4 also have under-duplexed embeddings on higher-genus one-colored skeletons, provided the circular diagrams for those higher-genus folds include the ribbon motifs found in the lower-genus diagram in Tables 2 and 4.

#### Crossing number of the polarised strand graph: one-colored embeddings

All of the one-colored embeddings of the {1212} folds in Fig. 17 contain edge-crossings, ranging from a single edge-crossing (*N*_×_ = 1) in (a) to *N*_×_ = 3 in (b,c,e,f,g). Most of the unduplexed strand pairs running between the Y-junctions are crossed, although the embeddings in Figs. 17(c-h) also contain some *un*crossed strand pairs, arrowed in Fig. 19.

**Figure 18:**
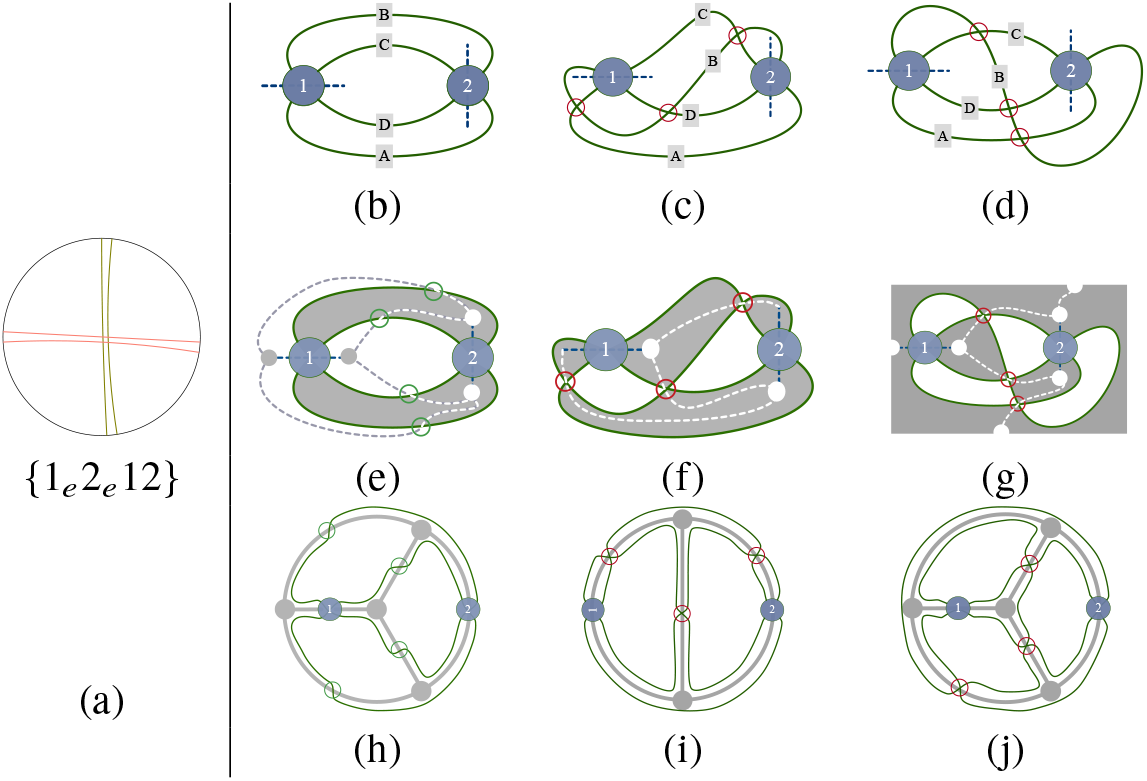
(a) Circular diagram of the (fully-flagged) fold 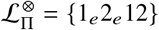. (b-d) Embeddings of the polarised strand graph with 0, 3 and 3 edge-crossings respectively (circled in red). (e-g) Checkerboard coloring of the graph embeddings with skeletons marked by dotted lines. (e) A two-colored skeleton whose edges cross the strand graph edges (circled in green); (f-g) Alternative one-colored skeletons. (h) The fold as a winding of a two-colored skeleton. Single strands wind around some of the edges (marked with green circles) and a pair of parallel strands around the others. (i,j) Rebuilding the same fold by winding a pair of parallel strands around each edge of the (i) genus-2 and (j) genus-3 one-colored skeletons (with additional crossings).

**Figure 19:**
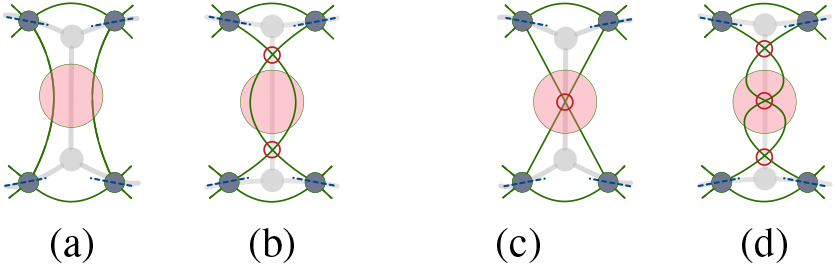
(a-e) One-colored embeddings of the polarised strand graphs in Fig. 17(c-g), with unduplexed strands from their parent folds highlighted in pink. Unduplexed strand pairs which are uncrossed are arrowed.

Any *even* number of crossings can be intercalated into those unduplexed edges to give a new embedding of the polarised strand graph, as shown in Fig. 20. Similarly, the (un-arrowed) crossed edges in Fig. 19 allow any odd number of edge-crossings.

**Figure 20:**
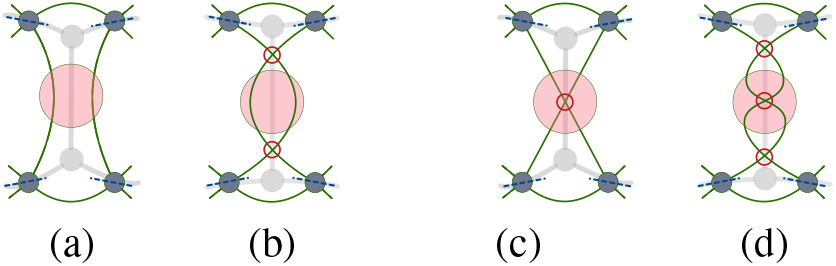
Alternative one-colored embeddings of a fragment of a polarised strand graph with different edge-crossings (circled in red, *N*_×_) along an unduplexed edge of the skeleton (highlighted in pink). (a) an uncrossed edge-pair (*N*_×_ = 0) and (b) the pair with two edge-crossings (*N*_×_ = 2). Embeddings of a strands graph with (c) a crossed edge-pair (*N*_×_ = 1) and (d) the same edge-pair with three crossings (*N*_×_ = 3).

Simpler embeddings of a fold have smaller numbers of edge-crossings in their polarised strand graph; the minimum crossing number (*N*_x_) among all possible one-colored embeddings of the polarised strand graph of a fold is an important index. (Later, we will determine this minimal number for all possible embeddings, including two-colored embeddings.) If a semi-flagged label 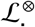 is included in Tables 2, 3 and 4, the associated folds can be embedded as a fully-duplexed genus-*g* fold with Y-junctions, such that *N*_×_ lies between 0 and 3(*g* – 1). More generally, semi-flagged (and fully-duplexed) folds in Table 4, whose average vertex-degree < *z* > is larger than 3 have *N*_×_ between 0 and 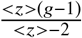 (from eq. 2). The lower bound, *N*_x_ = 0, is realised if all twist parities (*π_i_*) are tuned to allow a relaxed embedding. Since a single edge-crossing is sufficient to retune any detuned parity, at most *N*_×_ = *E* crossings are needed, where 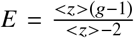. For example, the semi-flagged fold 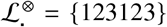 is listed in Table 2 as a genus-2 fold, whose tuned parities are all odd (*π*_1_ = *π*_2_ = *π*_3_ = *o*). Therefore the fully-flagged fold 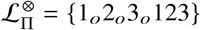 has *N*_×_ = 0. In contrast, the folds {1_*e*_2_*o*_3_*o*_ 123}, {1_*e*_2_*e*_3_*o*_ 123} and {1_*e*_2_*e*_3_*e*_ 123} have minimal crossing numbers *N*_×_ = 1, 2, 3 respectively.

Minimal edge-crossing numbers for embeddings of under-duplexed folds 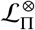 with up to six ribbons, whose semi-flagged label 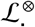 is not in Tables 2 or 4, can be determined as follows. First, find higher-genus (fully-duplexed) ‘parent’ folds, which we label 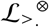, whose circular diagrams include the circular diagram of the fold in question, as for the example in Fig. 17. For these cases too, *N*_×_ is dependent on the tuned parities of the parent fold(s) 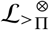, which give crossing-free embeddings (listed in Tables *S*1 and *S*6, in Supporting Information.) We split ribbons in the circular diagram of 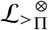 into those which are also present in the circular diagram of 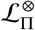 and those which are extraneous. First, each duplex induced by a ribbon found in both 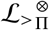 and 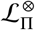 contributes either *N*_×_ = 0 or *N*_×_ = 1, depending on whether the relevant parity is tuned. In addition, each duplex in 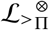 whose corresponding ribbon is absent in the circular diagram of 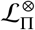 contributes *N*_×_ = 0 or 1, if its tuned parity is even or odd respectively. Thus, the minimal number of edge-crossings *N*_×_ in a one-colored embedding is at most 3(*g*_>_ – 1) where the fold 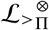 has genus *g*_>_ (realised by a fold wound on a 3-regular skeleton, all of whose parity flags are mistuned). Consider, for example, folds sharing semi-flagged label 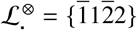. Those can be embedded as under-duplexed windings of the fully-duplexed fold, *viz*. 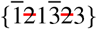. The relaxed parent fold has label 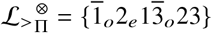(Fig. 16(a)). In this case, ribbon 2 can be unduplexed to give an uncrossed edge-pair, since its tuned parity is even (Fig. 16(b)). The remaining duplexes in the parent fold, (1,3) are relaxed provided both parities are odd. Therefore the simplest one-colored embedding of the fold 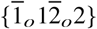 is uncrossed (*N*_×_ = 0), whereas that of 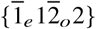 has *N*_×_ = 1, etc. (Fig. 21).

**Figure 21:**
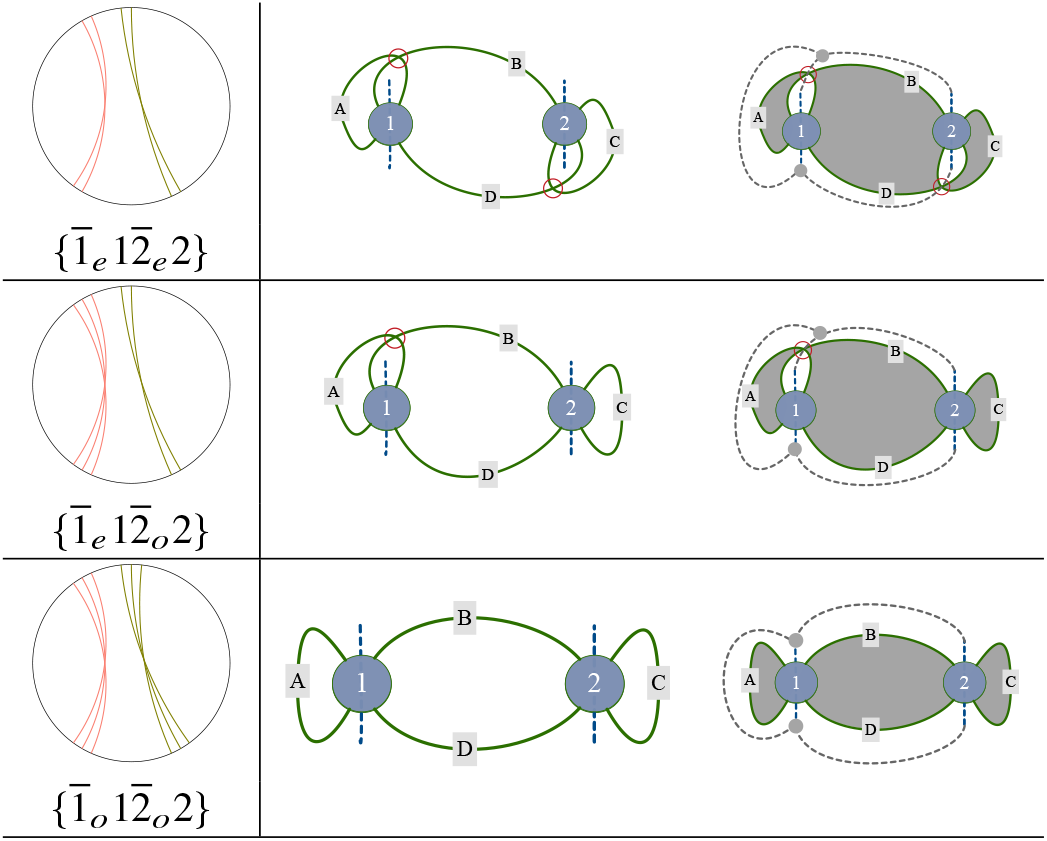
Embedding folds with common semiflagged label 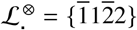. (Left column) Circular diagrams for fully-flagged folds 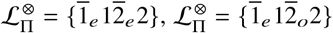 and 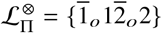. (Middle row) Embeddings of the fully-flagged folds as polarised strand graphs. The 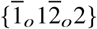 fold allows an embedding of the strand graph with uncrossed strands; 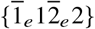 and 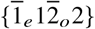 contain edge-crossings (circled in red).(Right row) Fold skeletons, shown as dashed lines. All three skeletons are one-colored.

In summary, one-colored embeddings of the polarised strand graph of a contracted fully-flagged fold have a minimal number of crossings, *N*_×_ though that number can be incremented by any even integer. Folds in Tables 2, 3 and 4 allow fully-duplexed embeddings without any edge-crossings (*N*_×_ = 0), provided all twist parities are tuned. Fully-duplexed one-colored embeddings of other folds with up to six contracted duplexes result in edge-crossings (*N*_×_ > 0). Lastly, one-colored embeddings of under-duplexed folds are either crossed or uncrossed, depending on their parity flags compared with those of parent folds.

#### Crossing number of the polarised strand graph: one or two-colored embeddings

Any contracted fully-flagged contracted fold can be swapped from a one- to two-colored embedding and vice versa, via the local moves illustrated in Fig. 9. Those moves introduce edge-crossings, so different minimum crossing numbers (*N*_×_) are realised for one- and two-colored embeddings. A crossing-free embedding of a contracted fold may therefore be one- or two-colored, but not both: a crossing-free one-colored embedding of a fold does not allow an uncrossed two-colored embedding and vice versa. For example, the fold {1_*e*_2_*e*_12} can be embedded via various one- or two-colored skeletons of genus-2 and −3, shown in Fig. 18. Both one-colored embeddings (on genus-2 and −3 skeletons respectively, see Fig. 18(c,f,i) and (d,g,j)) have *N*_×_ = 3, whereas the two-colored embedding is crossing-free (*N*_×_ = 0), wound on a genus-3 skeleton, see Fig. 18(b,e,h). Likewise, the {1_*o*_2_*e*_12} fold can be wound on a two-colored (genus-3) skeleton with *N*_×_ = 1 (Fig. 22(b,e,h)), or one-colored (genus-2 or −3) skeletons with *N*_×_ = 2 (Fig. 22(c,f,i)) or 3 (Fig. 22(d,g,j)).

**Figure 22:**
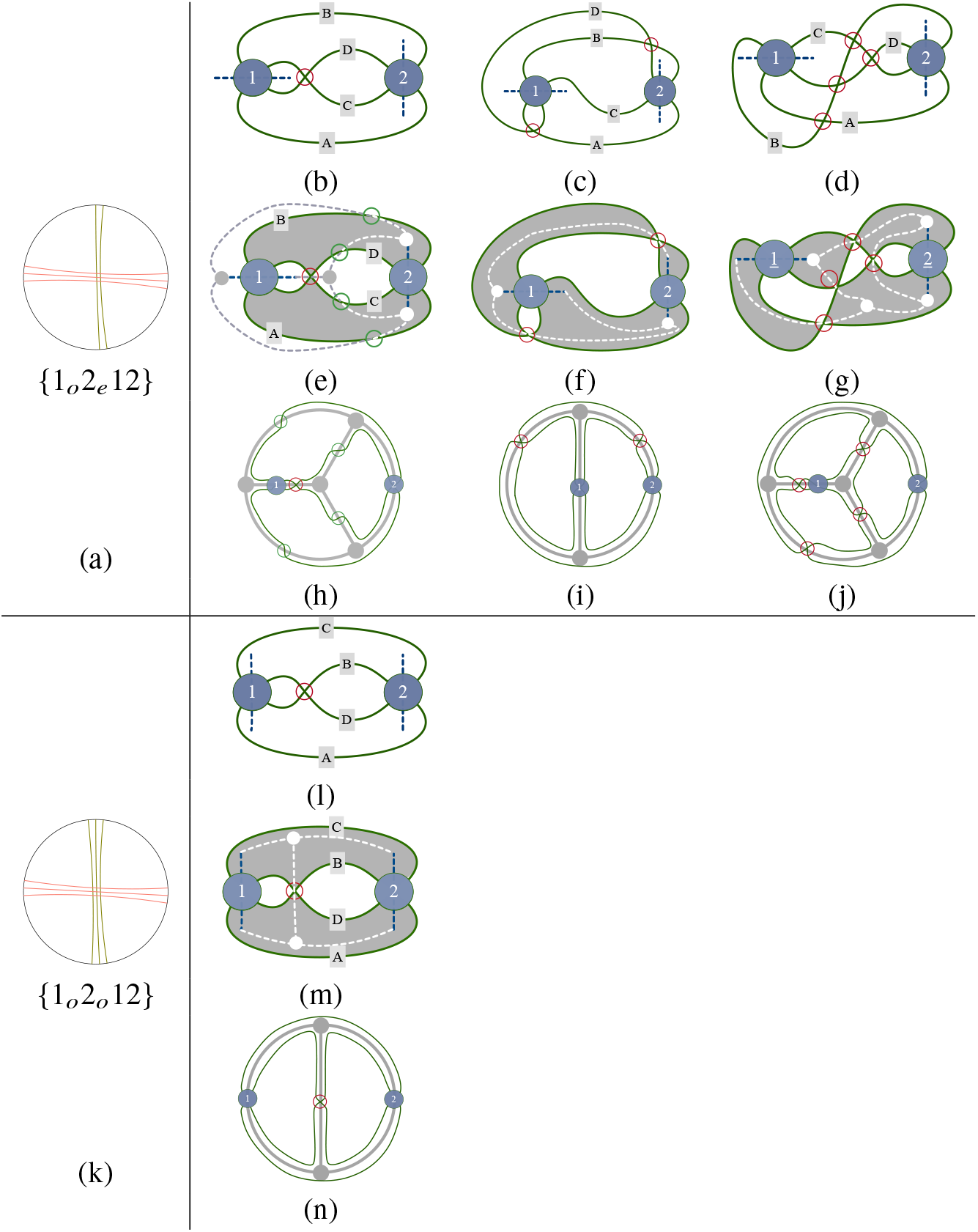
Embedding the fully-flagged folds 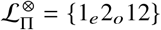 and 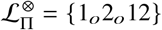, whose circular diagrams are shown in (a) and (k). Embeddings of {1_*e*_2_*o*_12} formed by windings on (b,e,h) a two-colored genus-3 skeleton, (c,f,i) a one-colored genus-2 skeleton and (d,g,j) a one-colored genus-3 skeleton. (l-n) Crossing-free embedding of {1_*o*_2_*o*_12} wound on a one-colored (genus-2) skeleton.

Notice that both one- and two-colored embeddings of the {1_*o*_2_*e*_12} fold contain edge-crossings (*N*_×_ > 0), irrespective of the embedding. Similarly, all embeddings of the polarised strand graph of the {1_*e*_2_*o*_12} fold contains edge-crossings. One-colored embeddings with *N*_×_ = 2 are possible. One crossing is due to the detuned odd parity duplex in the parent 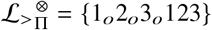 fold, a second crossing is required to retune the even-parity duplex 1_*e*_ to 1_*o*_(Fig. 22(c,f,i)). (More complex one-colored embeddings induce extra crossings, (Fig. 22(d,g,j)). Two-colored embeddings have *N*_×_ ≥ 1 (Fig. 22(b,e,h)).

It is not difficult to find larger families of folds whose embeddings necessarily include edge-crossings. For example, all fully-flagged folds sharing the semi-flagged label 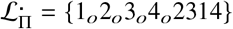 are embedded with edge-crossings, regardless of the orientation flags associated with each duplex, ⊗_*i*_. These include 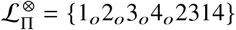, 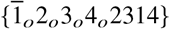, etc. More broadly still, families of folds sharing common *unflagged* labels can be found, whose embeddings include edge-crossings, regardless of their orientation or parity flags. Unflagged fold labels (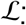, or 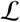) determine the pure topology of the (unpolarised) strand graph, stripped of rigid-vertex edge ordering and plumblines. If 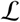 induces a topologically nonplanar graph, all folds sharing that bare label must have edge-crossings. The simplest relevant example is the so-called complete graph with five vertices, *K_5_* (30), whose edges join each vertex (*i* = 1,… 5) to the other four vertices (*i*’). That graph is realised by the unflagged label 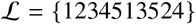 or any (reverse) cyclic permutation, since each vertex *i* is adjacent to *i*’ (e.g. reading from left-to-right, 1 is adjacent to 2, 5, 3. Since the string is cyclic, vertex 4 is also adjacent to 1). A large number of other unflagged labels result by swapping pairs of vertex labels *i* → *j* (e.g. 1 → 3 gives {1234513524} → {3214531524}).

In short, folds lie in three classes, distinguished by the minimum number of edge-crossings possible for one- and two-colored embeddings of their polarised strand graphs, listed in Table 5.

#### Relaxed or (a)-type folds

Fully-duplexed folds which allow one-colored embeddings of their polarised strand graphs without edge-crossings, are described as relaxed, since a crossing-free, one-colored embedding allows the strand to be duplexed along the entire length of each edge, interrupted only by fold junctions located at vertices of its fold skeleton. The architecture of those junctions corresponds to so-called ‘conventional’ junctions, as classified by Seeman *et. al* or ‘anti-junctions’, defined in (28). Conventional junctions are found, for example in transient four-way (X) Holliday junctions (31) and G-quadriplex motifs, observed in DNA *in vivo* (32) and DNA and RNA *in vitro* (33). Two examples of relaxed genus-3 folds, namely folds 12 and 14 (following the indexing in Table 2) flagged with natural parities, are highlighted in Fig. 23. Fig. 23e shows a fully-duplexed string realised by fold 14, whose duplexing is illustrated by (an arbitrary number of) red rungs. The fold contains four conventional Y-junctions, located at vertices of the genus-3 (‘white’) skeleton in Fig. 23(a). The vertices of the complementary one-colored (‘black’) skeleton Fig. 23(b), genus-12) mark anti-junctions: there are three seven-way anti-junctions in the interior of the fold. A fourth nine-way anti-junction, indicated by the semi-discs scattered around the boundary in Fig. 23(b), is formed if the planar fold in Fig. 23(f) wraps onto a sphere, so that those semi-discs glue to a single disc and the unbounded outermost region is gathered into a finite domain, centred by that anti-junction. A relaxed version of fold 12 is shown in Fig. 23(c,d,g,h,i). Here too, the pair of complementary (genus-3 and −5) skeletons define junctions and anti-junctions of the relaxed fold. We note in passing that in this example of the relaxed fold, we have assigned zero twist to ribbon 2, whose natural parity is even, i.e. *t*_2_ = 0. In order to highlight the duplexing of that pair of strands, we represent the untwisted duplex by a pair of unit twists of opposite sign. That schema is a useful one for untwisted duplexes, discussed in more detail below.

**Figure 23:**
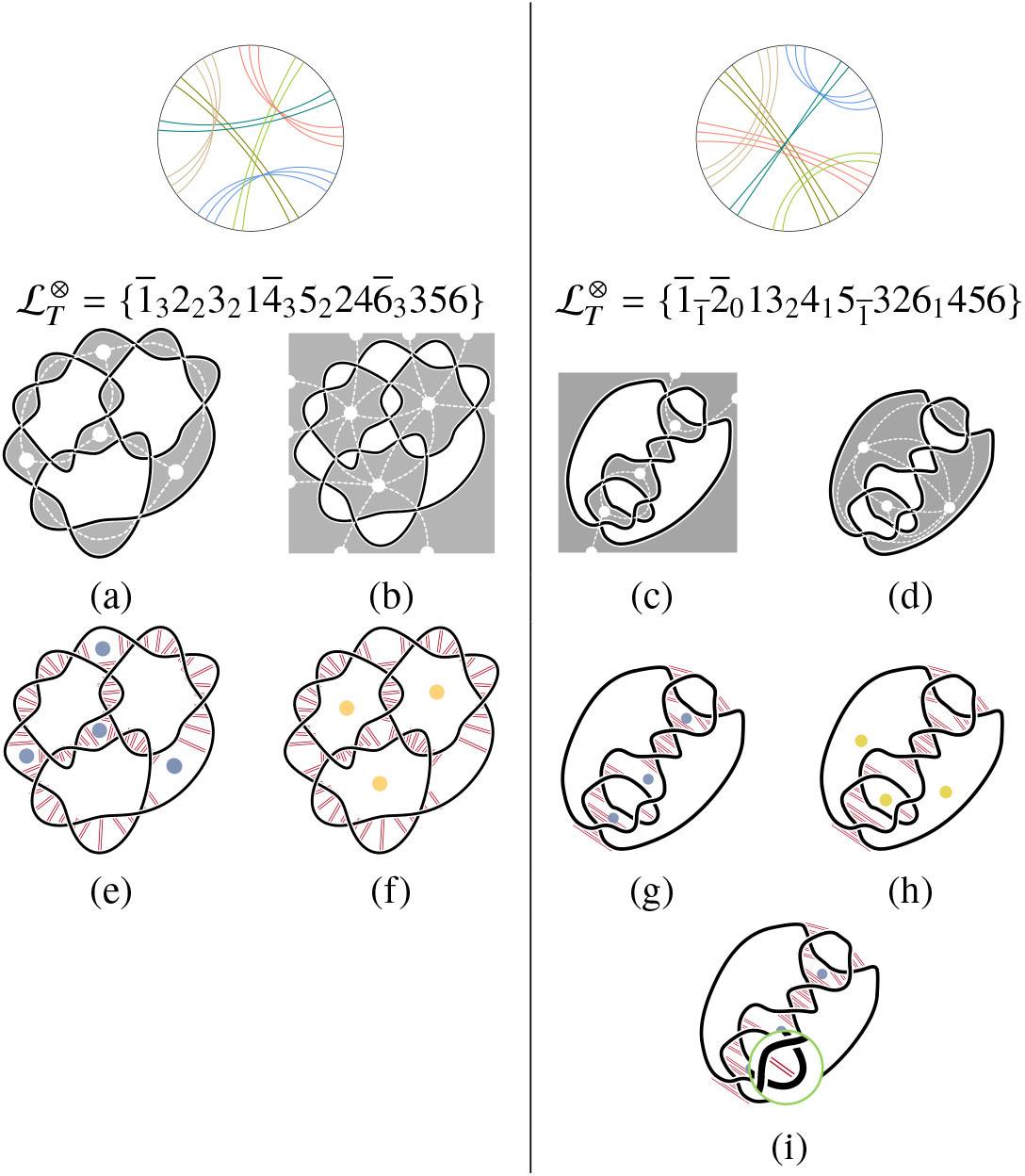
(Left, right) Relaxed genus-3 fully-flagged folds 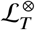. (a,b) Checkered colorings and one-colored skeletons of a fully-flagged fold corresponding to fold 14 in Table 2 (*cf*. Fig. 3). (c,d) A fully-flagged fold corresponding to fold 12 in Table 2 (*cf*. Fig. 13).(e-h) Three-way conventional Y-junctions (grey dots) and (d) (seven-way) anti-junctions (yellow dots) for those folds. (i) the diagram in (g), with (green inset) a magnified view of the untwisted duplex (*i* = 2, *t*_2_ = 0), represented by a pair of unit twists of opposite sign.

Embeddings of relaxed folds need not be fully-duplexed and may include more complex junction architectures than conventional junctions. Indeed, the degree of duplexing can vary from the minimum number of complementary nucleotides required to induce the requisite twists to all nucleotides in the loop. Further, any embedding of the polarised strand graph is possible, with multiple strand crossings, provided the rigid-vertex ordering and plumbline orientation relative to that vertex ordering is conserved. Such crossings disrupt the ideal junction/anti-junction architecture. For example, the ±1 pair of crossings adjacent to the untwisted duplex, shown in Fig. 23(i), disturb the pair of adjacent three-way junctions (grey dots in Fig. 23(g)). Those junctions are recovered by untwisting the ± 1 twist pair. Nevertheless, relaxed folds *may* be embedded to give very rigid structures, stabilised by bonding interactions across each complementary base-pair, rendering the entire fold double-stranded.

#### Uncrossed two-colored (or (b)-type) folds

Like (a)-type folds, fully-flagged folds in this second class allow embeddings of their polarised strand graph without edge-crossings. However, in contrast to (a)-type folds, those embeddings are two-colored. Duplexing is controlled by the plumbline axes, and since those axes are two-colored, duplexing flips along strand joining black and white plumblines, inducing junction architecture unlike that of conventional (or anti-) junctions of (a)-type folds, classified as meso-junctions in (28). For example, the crossing-free embedding of the 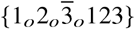 fold in Fig. 7(i) is two-colored^4^. The resulting flipped arrangement of H-bonded ‘rungs’ along the fold, linking complementary nucleotides, is illustrated in Fig. 24.

**Figure 24:**
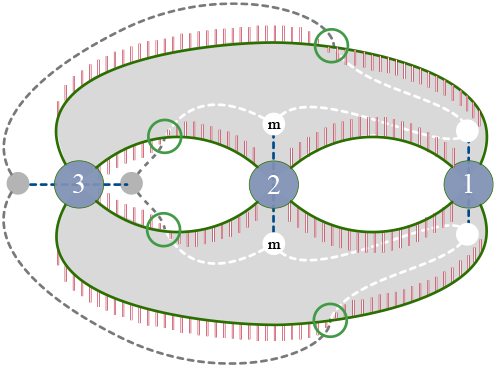
Schematic representation of a two-colored embedding of the fully-flagged fold 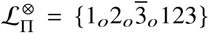 with no edge-crossings (*N*_×_ = 0) (*cf*. Fig. 7(d,i)). Black and white dots indicate vertices of the two-colored fold skeleton. Green circles indicate sites where skeletal edges change from black to white domains. The orientation of base-pairing is determined by plumblines of the polarised strand graph embedding; since those plumblines induce a two-colored (genus-4) skeleton, fold junctions (marked *m*) are meso-junctions. The red rungs indicate base-pairing orientation, which flips from one side of the strand to the other at the green circles.

Unlike (a)-type folds, (b)-type folds necessarily include single-stranded regions around those meso-junctions.

#### Crossed (or (c)-type) folds

The third class includes all fully-flagged folds whose polarised strand graphs embed with edge-crossings, regardless of the embedding. Clearly, these (c)-type folds cannot be fully duplexed, since the crossed edges induced by non-natural twist parities disrupt duplexing and induce unpaired single-strand regions in the fold. Edge-crossing also disrupts junctions. Those folds are expected to be more labile than either (a) or (b)-type folds, since they necessarily include significant stretches of single-stranded regions in the vicinity of edge-crossings.

The classification of fully-flagged folds as either (a), (b) or (c)-type folds (Table 5) is convenient, describing their simplest embeddings and junction architectures. There is an ideal ranking of these folds, from potentially ‘rigid’ (a)-type folds, to very floppy (c)-type folds, though that ranking assumes simplest embeddings (minimising *N*_×_).

### Crossing-free embeddings of the strand graph (*N*_×_ = 0)

All folds listed in Tables 2, 3 and 4 are (a)-type folds, other cases are either (b)- or (c)-type folds. To distinguish those, other techniques are required. Since (b)- and (c)-type folds have different *N*_×_ determination of the minimal number of edge-crossings, *N*_×_, is needed for arbitrary fold embeddings, whether one- or two-colored. First we outline the construction of semi-flagged folds whose duplexes have odd parity only. Lastly, we construct simplest embeddings of arbitrary folds from their unflagged label 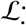, from which *N*_×_ can be determined for any combination of orientation and parity flags.

#### Odd-parity contracted folds

Consider for now, semi-flagged folds 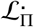 whose parity flags *π_i_*, are exclusively odd, *π*_1_*π*_2_ … = *oo* …, which we denote 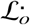. In those cases, the graph topology is conserved by replacing graph vertices by a single shadow-crossing (X), introduced above (e.g. Fig. 1). Kauffman has introduced ‘virtual’ knots, which allow both ‘classical’ (over-under) crossings, and ‘virtual’ crossings, whose topmost strand is unspecified, analogous to edge-crossings (34). Here we adapt that concept and label crossings of the strand which are subsumed within the strand graph vertices as ‘real’ crossings, and edge-crossings - visible in the strand graph embedding - as ‘virtual’. Real crossings are therefore those induced by helical winding in duplexes (whose twists *t_i_* induce *t_i_* crossings), whereas virtual crossings are unduplexed edge-crossings. The shadow diagram of any polarised or unpolarised strand graph embedding of a fold with *n* ribbons - whether one- or two-colored - therefore includes *n* real crossings and *N*_×_ virtual crossings. The unflagged fold label 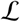 encodes a closed walk around the strand loop, marking each vertex-crossing and passing through unlabelled over edge-crossings. If the original polarised strand graph can be embedded without edge-crossings (*N*_×_ = 0), 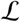 represents the shadow diagram of a classical knot. Therefore any such fold 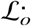, regardless of its orientation flags 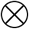, can be embedded such that *N*_×_ = 0. On the other hand, if 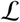 encodes a *virtual* knot, *N*_×_ > 0 and the knot contains *N*_×_ virtual crossings, folds 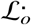 have *N*_×_ > 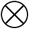 for all possible flags.

Two criteria suffice to ensure 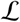 represents a real rather than virtual knot in the semi-flagged fold 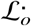, due to Dehn (35). First, there must be an even number of integers between any pairs of like integers (a constraint first recognised by Gauss). Next a modified cycle 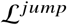 is constructed from the code, which effectively encodes an Euler walk around the knot (traversing all edges once), where the walker turns right or left at each crossing, thereby jumping between over- and under-strands.

The label 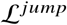 is generated from 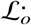 by iteratively reversing the order of all digit strings lying between 1 … 1, 2 … 2, etc. For example, the label 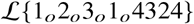 (Fig. 25(a)) gives a jump cycle 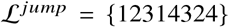 as follows: {1<u>23</u>14324} → {132<u>143</u>24} → {13<u>2</u>34124} → {13234<u>12</u>4} → {13234214}. A new circular diagram is constructed from the label 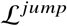 (Fig. 25(b)). If some subset of the arcs can be reflected from the interior of the bounding perimeter to its exterior to give a diagram with no edge-crossing, *N*_×_ = 0 (Fig. 25(c)), the original fold 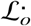 allows an embedding with *N*_×_ = 0. (That process can be computed, allowing an arbitrary code 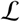 to be identified as relevant to real or virtual knots, see (36) for a succinct summary. In practice, for small *n* it is readily done by hand.) The segregation of inner from outer arcs allows explicit construction of the uncrossed embedding of the shadow diagram of 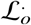 (Fig. 25(d)), giving a crossing-free embedding of the strand graph (which is unpolarised, since there are no orientation flags). Insertion of suitably oriented plumblines generates the polarised strand graph embedding (Fig. 25(e)), leading to an explicit planar drawing of the simplest embedding of any fully-flagged (all-odd parity) fold 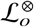 (e.g. Fig. 25(f)).

**Figure 25:**
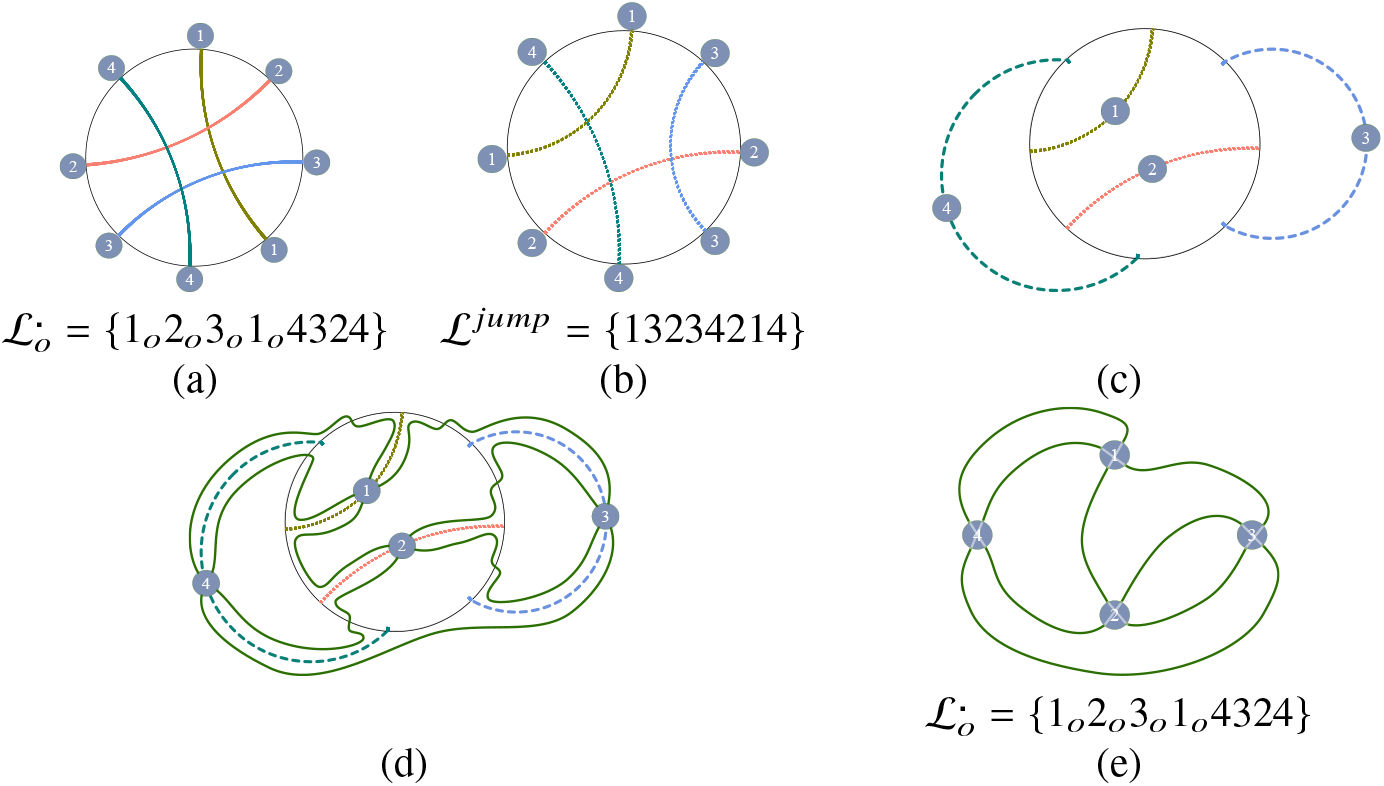
(a) A semiflagged fold 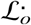; since the ribbons have unflagged orientation (i.e. either annular or moebius), they are drawn as zero-width arcs, rather than ribbons. The Dehn diagram is constructed by (b) drawing the reconfigured ‘jump’ label 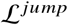 and (c) reflecting selected arcs to the outside in order to remove all arc-crossings. (d) An embedding of the fold 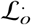 is found by building a closed cycle with label 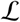 and crossing the perimeter when passing from an inner to outer arc, or vice versa. (e) The embedding of the unpolarised strand graph of 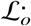 without edge-crossings.

Dehn’s criteria determine whether the polarised strand graph of a semi-flagged fold whose parities are all odd (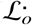) can be embedded in the plane without crossings *N*_×_ = 0 for an arbitrary label 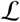. All distinct labels with *n* ≤ 5 (summarised in Table 1 and listed in Table *S*7, Supporting Information) have been checked against Dehn’s criteria, so all semi-flagged folds 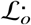 can be classified according to their edge-crossings, distinguishing those allowing *N*_×_ = 0 from others which require *N*_×_ > 0 regardless of their embeddings. The resulting classification is listed in Table 6.

Folds with no edge-crossings can be flagged with any combination of ribbon orientations to give fully-flagged folds 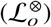 with uncrossed embeddings, listed in Table 7. Flagging those folds with any combination of annular and moebius orientation flags leads to all folds 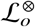, with *N*_×_ = 0, which are *a priori* uncontracted. Some of those contract to new fold labels with even parities, also listed in Table 7. The suite of resulting folds (with up to four ribbons in the label 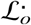) are described in detail in Table 7. This procedure can be extended at will: all crossing-free contracted odd-parity folds with *n* ribbons are found by checking all distinct labels 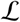 with up to *n* ribbons. However, crossing-free contracted folds with *n* ribbons including even parities require searches of uncontracted semi-flagged folds of all labels 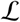 with up to 2*n* ribbons, since each contracted ribbon with even parity is equivalent to an uncontracted pair of ribbons (nested or crossed, depending on the parity of the pair of like-parity ribbons).

### Simplest embeddings of generic fully-flagged folds

Previous constructions led to (i) one-colored fully-flagged folds 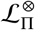 and (ii) one- or two-colored odd parity fully-flagged folds 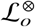 whose polarised strand graphs can be embedded without edge-crossings. More generally, for any unflagged label, 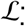, we can find all possible combinations of orientation and parity flags which allow crossing-free embeddings of the resulting fully-flagged folds 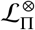. The construction relies on the possibility of forming an uncrossed planar embedding (*N*_×_ = 0) of the unpolarised graph 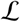 which we assume (for now) is topologically planar. In many cases, uncrossed embeddings can be produced from standard mathematical graph drawing packages which produce Tutte-like embeddings (e.g. *Mathematica*). Such embeddings rely on Tutte’s Spring Theorem, which guarantees an uncrossed planar embedding of a graph, provided it is topologically planar and 3-vertex-connected (37) (a simplified account can be found online at (38)). We call these embeddings ‘Tutte-like’, since they extend Tutte’s spring-like embeddings, to produce embeddings of planar, 3-connected multigraphs by curving multigraph edges joining the same pair of vertices to avoid edge overlap. If the graph is 3-connected that embedding is unique, otherwise there is some flexibility (readily detected for simpler folds.) We then add internal chords within each vertex, joining graph edges on the vertex perimeter in pairs, to form an (Euler) cycle which traverses each edge, tracing out a path with label 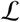. (The process is a generalisation of the construction described above, illustrated in Fig. 4.) That cycle passes through each vertex twice, and the pair of internal chords are uncrossed or crossed. Consider fro example a fold with bare label 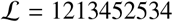. That label allows a planar Tutte-like embedding of the 3-connected (multi-)graph, shown in Fig. 26(a). Next, strands are extended through all 5 vertices to give the Euler cycle 1213452354; one possible construction is shown in Fig. 26(b), which gives the marked strands in (c).

**Figure 26:**
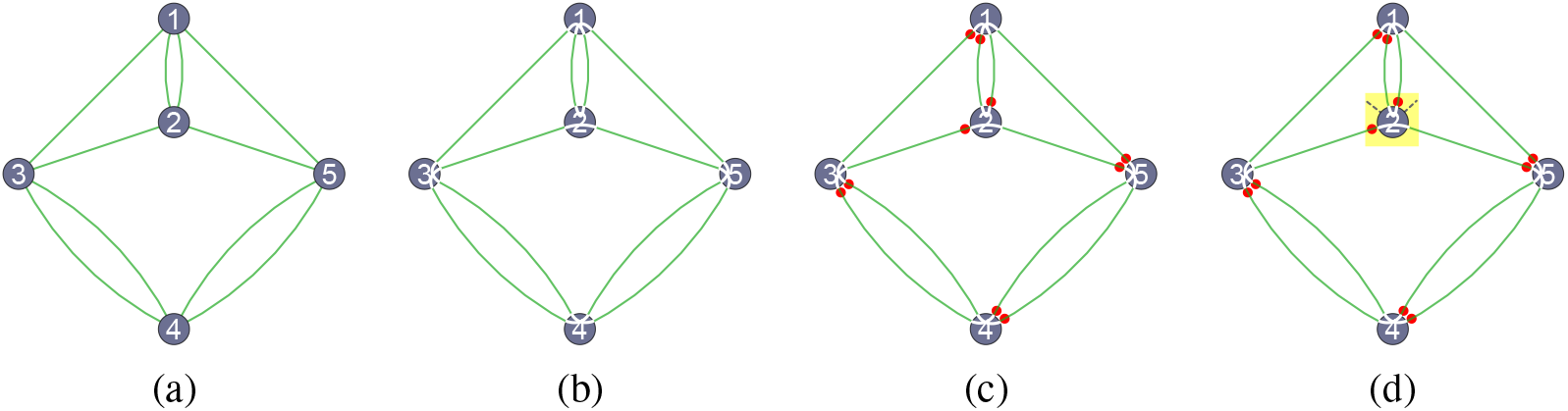
(a) An unpolarised strand graph common to all folds with bare label 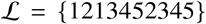, found by standard graph-drawing algorithm enforcing Tutte-like embeddings (assuming 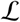 encodes a topologically planar graph). (b) One possible closure of the graph edges into a cycle with label 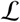 by connecting edge pairs via white chords within each graph vertex, forming uncrossed (even parity, *π_i_*, = *e*) and crossed (odd parity, *π_i_*, = *o*) motifs at vertices 2 and 1,3,4,5 respectively, giving a semiflagged fold 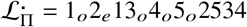. (c) Directing the cycle in (b) and marking each exit strand from vertices 1,…, 5 by red dots. (d) Since vertex 2 has even parity its plumbline is necessarily oriented left↔right rather than top↔bottom, inducing an antiparallel duplex at 2.

An uncrossed Tutte embedding describes embeddings of polarised strand graphs with *N*_×_ = 0, representing specific fully-flagged folds 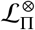, whose flags Π and 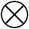 are found as follows. The pattern(s) of crossed or parallel internal chords in each vertex determines the twist parity: crossed chords within vertex *i* signal a duplex of odd parity (*π_i_*, = *o*), whereas uncrossed chords encode an even parity vertex (*π_i_*, = *o*). For example, the label 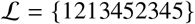 can be decorated with vertex chords as shown in Fig. 26(b), leading to a semi-flagged fold with label 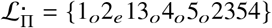.

To determine admissible orientation flags at each vertex, they must first be ‘marked’ by directing the Euler cycle and marking each exit strand from a vertex (*cf*. Fig. 4). Marking of the cycle 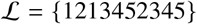 leads to the pattern in Fig. 26(c). Finally, the marked graph embedding must be polarised by inserting plumblines at each vertex. Assuming a local *NS* (equivalent to *SN*) plumbline at each vertex, its polar meridian partitions the four edges of the strand graph incident to that vertex into a pair of strands, two in each (*N* and *S*) hemisphere. If the vertex has odd parity, the plumbline has two allowed orientations, corresponding to both possible pairings. If it has even parity, the plumbline passes between the uncrossed internal chords, allowing just one pairing of edges. For example, the Euler cycle in Fig. 26(b) has just one even vertex (vertex 2) so that plumbline is constrained as shown in Fig. 26(d). All other vertices in the graph have two alternative plumbline axes.

Possible annular/moebius flags depend on the even/odd parity flag at that vertex and its plumbline, illustrated in Fig. 27. The pair of markings associated with vertex *i* both lie within the *N* or *S* hemispheres (sharing a common latitude), or both within *E* or *W* hemispheres (with common longtitude). In the former case, the vertex *i* hosts a parallel duplex, and the associated ribbon has moebius orientation ⊗*_i_*, denoted *ī*); alternatively, the vertex hosts an antiparallel duplex with an associated annular ribbon.

**Figure 27:**
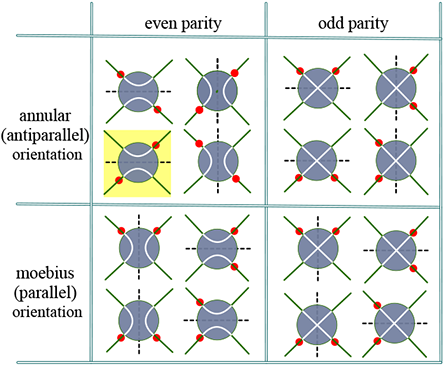
Possible strand trajectories (white paths) and directions through a single vertex of a polarised strand graph, marked by red dots indicating strand direction denoting exit arrows of a directed folded loop. Plumblines are drawn as dashed lines though the vertex. Uncrossed and crossed strands characterise polarised strand graph vertices hosting even and odd parity duplexes respectively. The duplex orientation (parallel or antiparallel) is set by the dot configuration with respect to the plumbline. For example, vertex 2 in Fig. 26(d) corresponds to the motif highlighted in yellow.

The marked graph in Fig. 26(c) has a fixed plumbline through vertex 2 (shown in Fig. 26(d)), since that vertex has even parity, so that vertex 2 hosts an antiparallel duplex (and its associated fold diagram contains an annular ribbon). The other (odd parity) vertices can be fully-flagged by inserting a ‘vertical’ or ‘horizontal’ plumbline at those vertices, so their orientation flags can be either parallel (moebius) or antiparallel (annular), depending on that choice. The continuous strand configuration drawn in Fig. 26(b) therefore induces 2^4^ = 16 different fully-flagged folds with labels 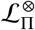. Those fully-flagged labels a 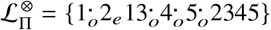, where vertices denoted *i*· have either orientation flag, i.e. *i* or *ī*.)

In general, a label 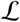 allows more than one combination of even and odd parities, corresponding to different Euler cycles traced though the graph embedding. For example, the Euler cycle 1213452534 in Fig. 26(b) is one of eight different strand paths through the vertices of the unpolarised graph are possible, sorted in Fig. 28. The resulting fully-flagged folds are shown in Fig. 29.

**Figure 28:**
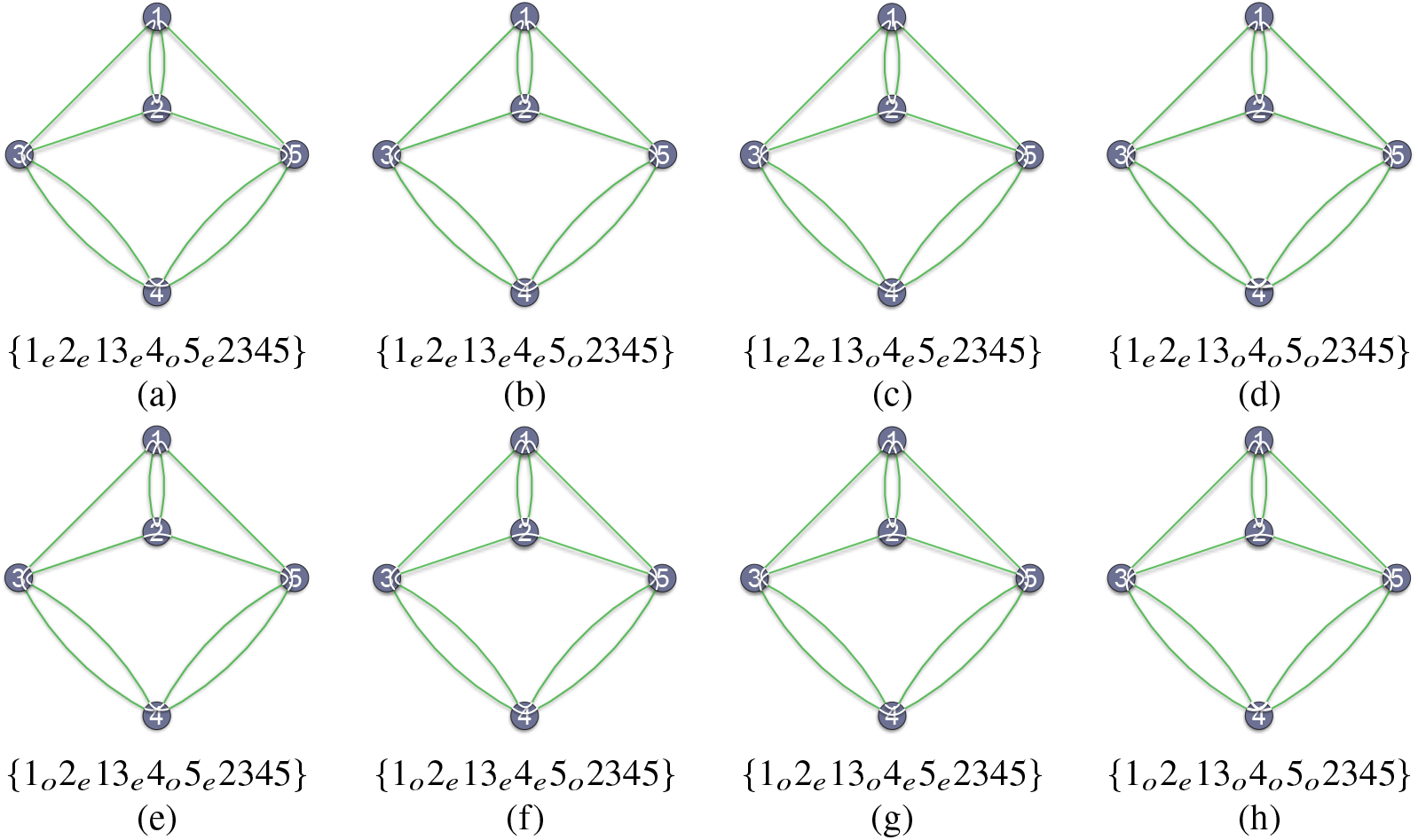
All semi-flagged folds 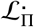 allowing crossing-free embeddings (*N*_×_ = 0), with bare label 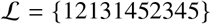.

**Figure 29:**
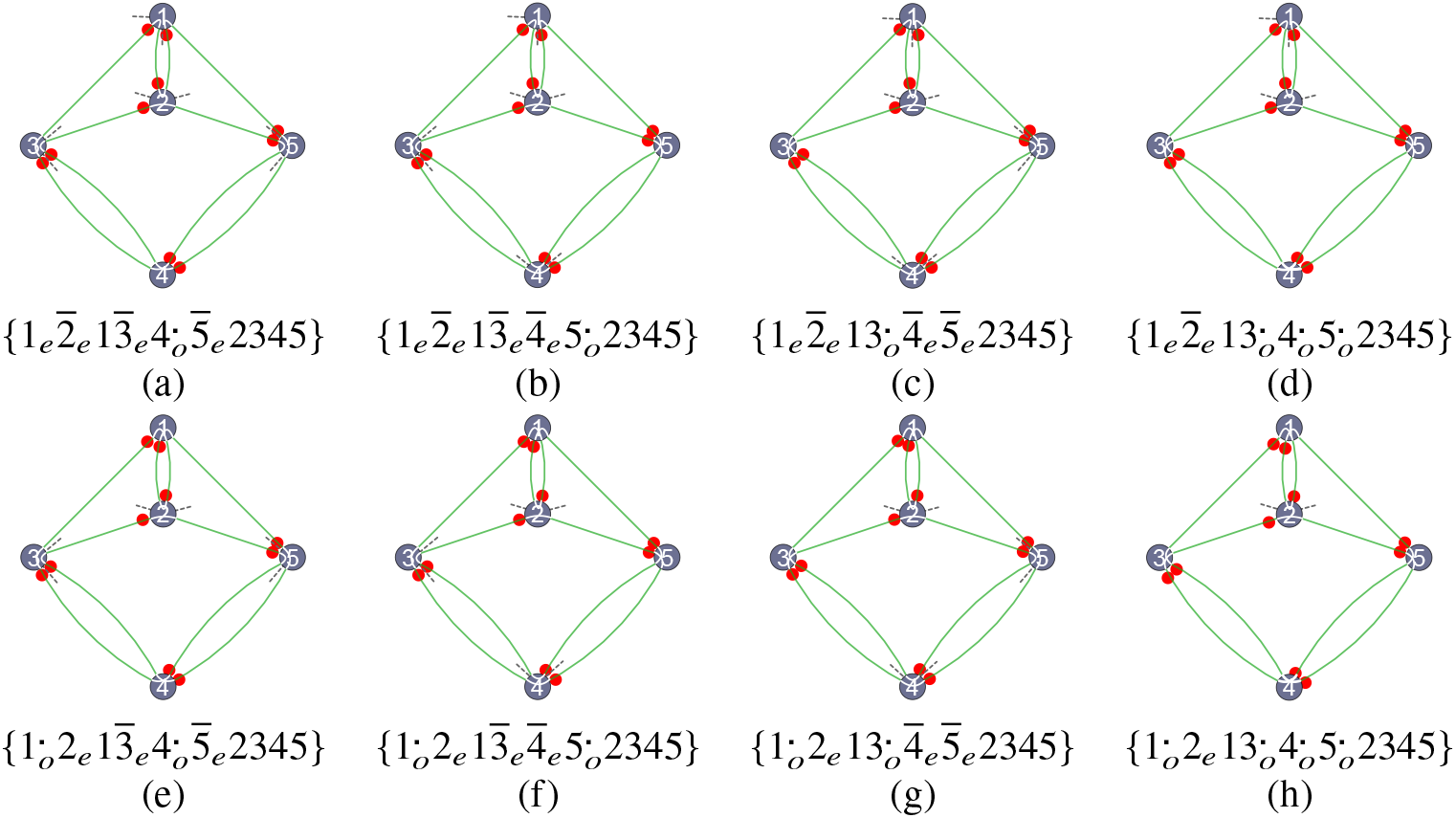
All fully-flagged folds 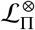 allowing crossing-free embeddings, with bare label 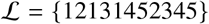. Ribbons with superscript *i*· may be labelled *i* or *ī*, e.g. 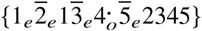 in (a) denotes folds 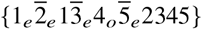 and 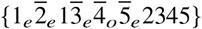.

In total, 42 fully-flagged labels result. That estimate over-counts the number of distinct folds, since the canonical label 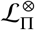 is conserved if vertices 3 ↔ 5 are exchanged. (Exchanging 3 ↔ 5 gives exchanged labels 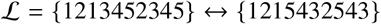, whose canonical form is {1213452345}). The number of distinct folds is therefore reduced by a half, so that the bare fold label 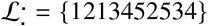 embedded via the unpolarised strand graph shown in Fig. 26(a) admits 21 distinct fully-flagged folds. Since the strand graph embedding is uncrossed (*N*_×_ = 0), all of the resulting fully-flagged folds are either (a)- or (b)-type folds, depending on plumbline orientations within odd parity vertices. For example, the folds in Fig. 29(h) include a fold with annular ribbons whose contracted duplexes are all-antiparallel, {1_*o*_2_*e*_13_*o*_4_*o*_5_*o*_2345} (Fig. 30(a)). Since that embedding is two-colored, it is a (b)-type fold. If the orientation flags associated with each odd parity vertex are switched from annular to moebius, the 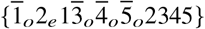 fold results (Fig. 30(b)), which is an (a)-type (relaxed) fold. Since each duplex has two possible orientation and parity flags, there are 4^5^/2 = 512 distinct fully-flagged folds with bare label {1213452534} in total. It follows that the vast majority are (c)-type folds, whose planar embeddings contain edge-crossings, regardless of the embedding. These (c)-type folds can be formed from *any* (a)- or (b)-type fold by detuning previously tuned orientation and/or parity flags. Since switching flags in contracted folds necessarily inserts edge-crossings, all fully-flagged fold whose labels are not among those in Fig. 30 must be (c)-type folds, with *N*_×_ > 0. For example, the one- and two-colored embeddings of uncrossed folds in Fig. 30(a,b) give the one- and two-colored embeddings of the fold {1_*e*_2_*e*_13_*e*_4_*e*_5_*e*_2345} shown in Fig. 30(c,d).

**Figure 30:**
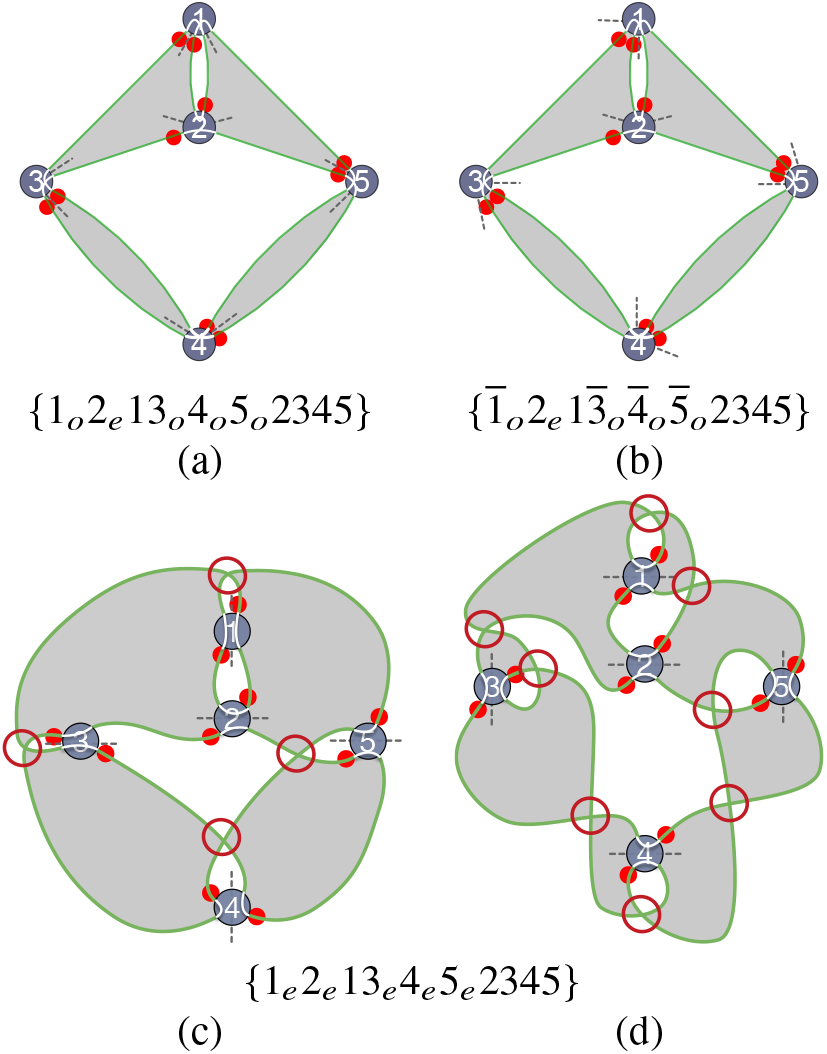
(a,b) Two of the 16 fully-flagged uncrossed folds in Fig. 29(h): (a) An uncrossed one-colored (a)-type (relaxed) fold. (b) An uncrossed two-colored ((b)-type) fold. (c,d) Alternative one and two-colored embeddings of the fold {1_*e*_2_*e*_13_*e*_4_*e*_5_*e*_2345} fold by inserting edge-crossings neighbouring vertices 1, 3, 4, 5 in the embeddings in (a) and (b) respectively to detune selected orientation and parity flags as in Fig. 31. The fully-flagged fold in (a) is transformed to the fully-flagged fold in (c) by switching *π*_1_, *π*_3_, *π*_4_ and *π*_5_, giving *N*_×_ = 4. Fold (b) is transformed to the fully-flagged fold in (d) by switching the same 4 parity flags, as well as orientation flags ⊗_1_, ⊗_3_, ⊗_4_ and ⊗_5_ leading to *N*_×_ = 8.

More generally, orientation (⊗_*i*_) and/or parity flags (*π_i_*) can be switched at any vertex by inserting a single edge-crossing into the graph embedding, in ‘parallel’ or ‘series’, as shown in Fig. 31.

**Figure 31:**
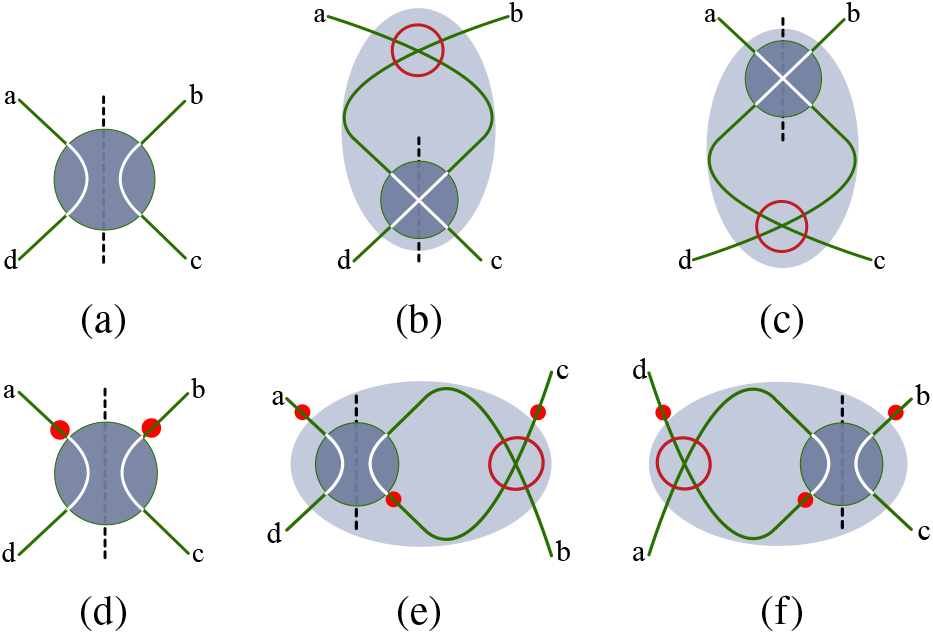
Switching parity and orientation switches at a vertex whose plumbline is aligned *NS* by inserting an edge-crossing (circled in red). (a-c) The parity flag *i_π_* is switched from (a) even to (b,c) odd by crossing the pair of edges facing *N* (or *S*).(d-f) Crossing the pair of edges facing *E* (or *W*) switches the orientation flag *i*^⊗^ from (d) parallel to (e,f) antiparallel.

The minimal number of edge-crossings, *N*_×_ can be estimated for an arbitrary fully-flagged fold 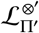, as follows. First, generate all possible fully-flagged folds allowing embeddings with *N*_×_ = 0, whose flag-free labels are identical to that of the target fold 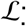. Denote those folds as 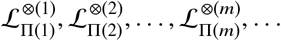. Clearly, if one of those coincides with 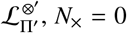, *N*_×_ = 0. Otherwise, the number of flag switches (*N*(*i*)_×_) required to detune the orientation and parity flags ⊗(*m*) and Π(*m*) to match those of the target fold (⊗’ and Π’) is found for all *m*. The simplest embedding of 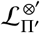 has *N*_×_ = *min*(*N*(*i*)_×_), the minimum value of *N*(*i*)_×_.

The construction is not limited to topologically planar graphs. If the label 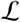 encodes a nonplanar graph, *N*_×_ includes the minimum number of edge-crossings in the embedding of the bare graph encoded by the cycle 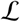 (for planar graphs, this number os 0). For example, the nonplanar *K*_5_ graph can be embedded with *N*_×_ = 1 so all folds whose bare label encodes that graph have *N*_×_ ≥ 1.

#### Valid fold labels: constructing all folds from a bare label 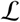

The process described above affords a simple constructive proof that any label 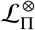 is an admissible label for a fully-flagged duplexed fold, assuming 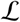 contains just two occurrences of each digit. First, it is evident that any such string 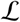 encodes a 4-regular graph. Any embedding of that graph can be used to initiate the construction of a fold, irrespective of *N*_×_ in that embedding. A directed Euler cycle encoding the path 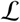 is traced out in the graph, constraining parity flags. The associated marked cycle, plus allowed plumbline axes, determine the suite of parity and orientation flags associated with that embedding. Judicious switching of flags gives an embedding of the polarised strand graph for the label 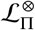. In general, the resulting embedding will be far from simplest, with more edge-crossings than necessary. Nevertheless, it is an admissible embedding of the polarised strand graph and therefore a duplexed winding corresponding to 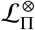. Simplest embeddings are found by first embedding the undecorated graph 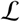 with as few edge crossings as possible.

#### All-antiparallel and all-parallel relaxed folds of arbitrary even and odd genus

Table 1 suggests the rapid growth of the number of folds sorted by bare fold label 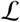 with ribbon number *n* - ca. 800 distinct labels are found for *n* = 6. Since arbitrary orientation flags can be appended to those fold labels, most of those induce ‘mixed parallelism’ folds, with both parallel and antiparallel double-helices. If all ribbons are assumed to be annular, forming semi-flagged labels 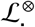, the number of different contracted ‘all-antiparallel’ folds are a small subset of generic folds, listed in Table 8.

**Table 8:** Distinct semiflagged, contracted canonical fold labels, 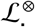, for folds containing up to 5 duplexes, assuming all ribbons are annular.

| $n$ | *nmbr of labels* | *smallest $\mathcal{L}$* | *largest $\mathcal{L}$* |
| --- | --- | --- | --- |
| 0 | 1 | {0} | {0} |
| 1 | 0 | — | — |
| 2 | 1 | {1212} | {1212} |
| 3 | 1 | {123123} | {123123} |
| 4 | 4 | {12123434} | {12341234} |
| 5 | 19 | {1212343545} | {1234512345} |

The most arresting examples of ‘all-parallel’ and ‘all-antiparallel’ folds, whose duplexes are exclusively parallel or antiparallel, have one-colored embeddings on particularly simple (and symmetric) skeletons known as generalised *θ*-graphs. Like the conventional *θ*-graph, they include just two vertices of the same degree linked by meridional edges; generalised *θ_z_* skeletons versions include *z* meridians. The simplest member, with a single edge joining end-vertices (*θ*_1_) is a genus-0 skeleton and the usual *θ* graph is *θ*_3_. For arbitrary integer *z*, the *θ_z_*-skeleton has genus *z* – 1.

The all-antiparallel fully-flagged folds 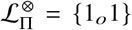 and {1_*o*_2_*o*_3_*o*_123} can be wound on one-colored genus-0 and −2 skeletons (*θ*_1_ and *θ*_3_ respectively) to give relaxed folds. Those two cases are the simplest examples of an infinite family of *all-antiparallel* folds of arbitrary *even* genus (2*k*, where *k* is any positive integer): a generic member has fully-flagged label 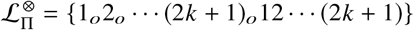. Like the genus-0 and −2 folds, all members can be wound on generalised-*θ*_z_ one-colored skeletons, *θ*_2*k*+1_, to give relaxed folds. The first three members of the family are shown in Fig. 32(a-c). These all-antiparallel folds have particularly simple circular diagrams, containing 2*k* + 1 crossed annular ribbons (see Fig. 32).

**Figure 32:**
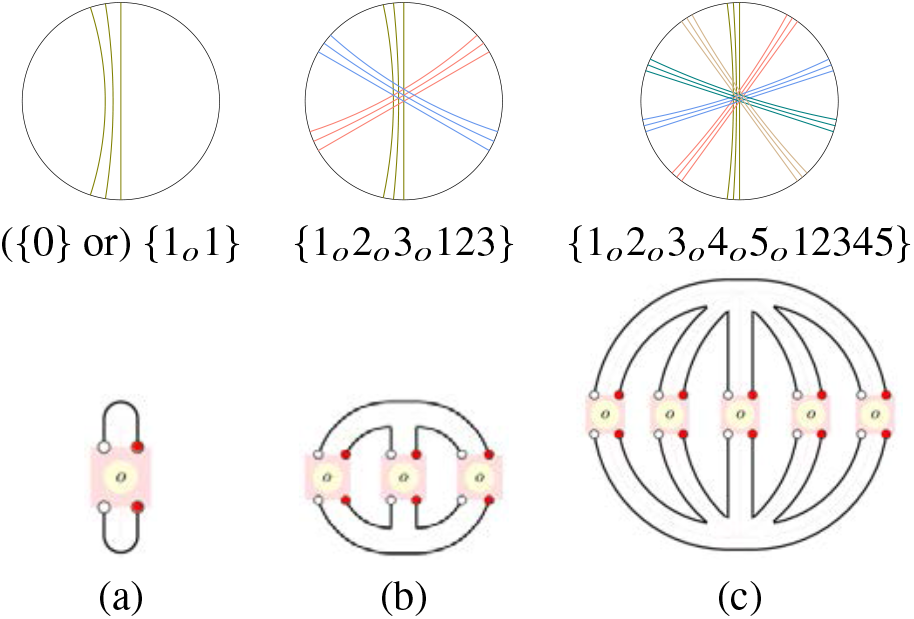
(Top) All-antiparallel natural-parity fully-flagged folds (labelled by 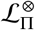) drawn as two-track railways with 1, 3, 5 and 6 odd-parity duplexes respectively. (Middle) Canonical fold labels and (bottom) (all-annular) circular diagrams. (a) The genus-0 fold {1_*o*_1} (which contracts to the trivial unfold {0}). (b) Genus-2 fold on the conventional *θ* (*θ*_3_) skeleton, with a pair of conventional degree-3 (Y) junctions. (c) Genus-4 fold on the ‘lunar’ skeleton of the *θ*_5_ graph, with a pair of degree-5 junctions. (The orientation of each duplex can be found by orienting the fold, marked by the pair of red dots denoting exit strands from each vertex; parallel and antiparallel duplexes have dot-pairs located at the same or antipodal poles.)

Junctions of *θ*_2*k*+1_ skeletons can be split in a number of ways to give new (even-genus) skeletons with lower-order junctions. Since the *θ*_2*k*+1_ skeletons are one-colored, those new edges host unduplexed and unwound pairs of strands. Those strands can be duplexed (with twist *t_i_* = 0) and replaced by duplexes of arbitrary even-parity twist. Further, the new duplexes are also antiparallel, inducing a number of new all-antiparallel folds on even-genus skeletons with more than two vertices. Conversely, the higher-degree {1_*o*_2_*o*_ … (2*k* + 1)_*o*_ 12 … (2*k* + 1)} folds are under-duplexed embeddings of many other folds of lower degree. If the skeletal edges are split until all vertices are degree-3, all-antiparallel folds wound on skeletons with Y-junctions result, whose circular diagrams contain the characteristic crossed (annular) ribbon motif of the original higher-order even-genus *θ*_2*k*+1_ fold. The genus-4 folds with indices 18 – 21 in Table 3 are simpler examples of those Y-junction folds, whose skeletons can be contracted to *θ*_5_ skeletons.

A related infinite family of *all-parallel* folds are found on generalised-*θ*_2*k*_ skeletons of *odd* genus, 2*k* – 1. Those folds are generalisations of the fully-flagged folds 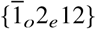 and 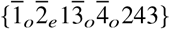 on genus-1 and 3: generic genus-(2*k* – 1) folds have canonical label 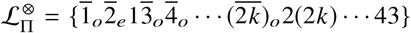, whose simplest members are shown in Fig. 33. (In contrast to all-antiparallel folds, if the vertices in these *θ*_2*k*+2_ folds are split to form windings on lower-degree skeletons, the pair of unwound strands or even-parity duplexes traversing the new exposed edges are antiparallel, so those new folds include both parallel and antiparallel duplexes.)

**Figure 33:**
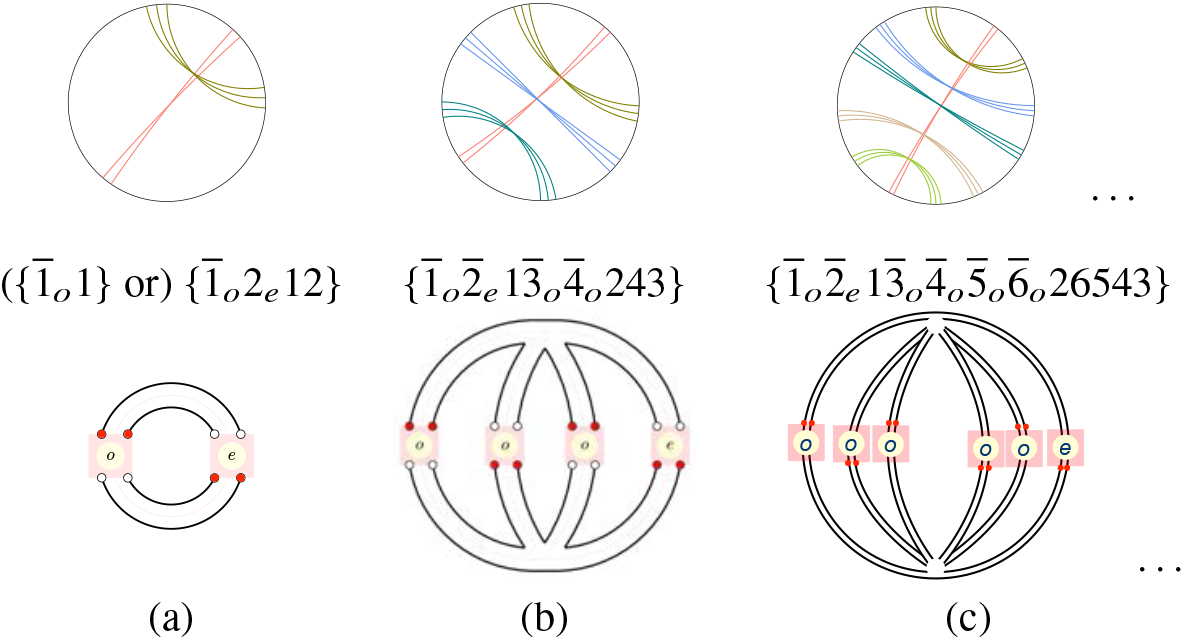
A family of relaxed all-parallel fully-flagged folds 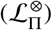 wound on skeletons of odd genus. (a-c) Genus-1, −3 and −5 folds on (a) the genus-1 torus, (b) the genus-3 *θ*_4_ skeleton and (c) the genus-5 *θ*_6_ skeleton. (The {1_*o*_2_*e*_12} fold is uncontracted; it contracts to {1_*o*_1}.) The location of the even-parity duplex (marked *e*) is arbitrary.

These all-antiparallel folds on even-genus skeletons are relaxed provided all parities are odd, whereas the odd-genus all-parallel folds are relaxed provided only one of the 2*k* duplexes has even parity and all others have odd parity. Those (relaxed) folds and their related circular diagrams lead to numerous relaxed folds on higher-genus skeletons, since they can be glued to give all-parallel and all-antiparallel composite folds with various junction orders and duplex counts by gluing appropriate *θ*_z_ skeletons. Three examples are shown in Fig. 34.

**Figure 34:**
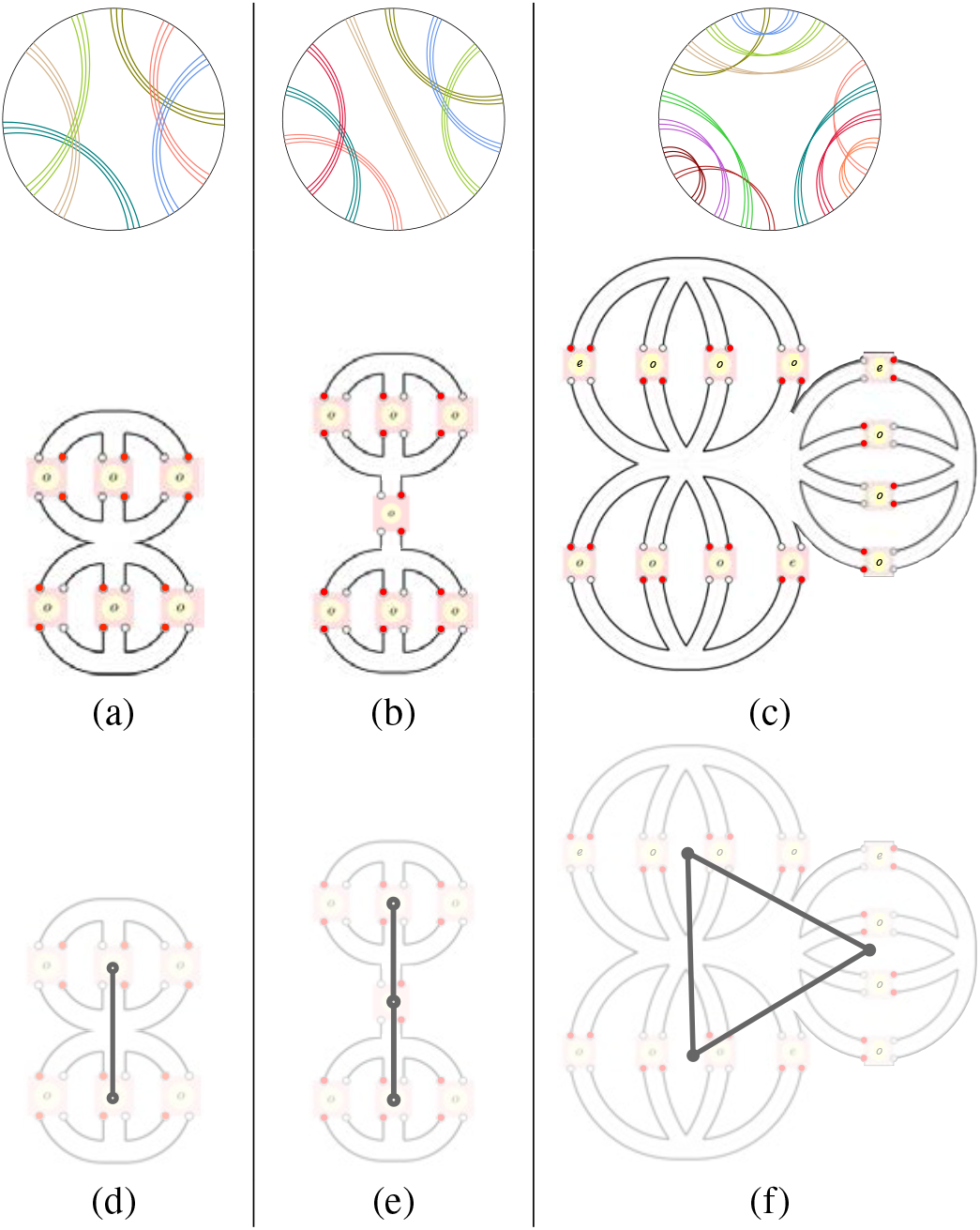
Collections of either all-parallel or all-antiparallel folds on generalised *θ* skeletons in Figs. 32 and 33 (of odd and even genus respectively) can be glued to form composite all-parallel or all-antiparallel folds. (a) An all-antiparallel fold on a new genus-4 skeleton formed by merging a pair of all-antiparallel folds, giving a fold skeleton with a pair of Y-junctions and a (central) degree-6 junction. (b) A glued stack of (from top to bottom) genus-2, genus-0, genus-2 all-antiparallel folds, forming a new genus-4 all-antiparallel fold. (c) Gluing three radially arranged all-parallel genus-3 fold modules gives a new all-parallel genus-9 fold.The circular diagrams (top row) describe composite - rather than prime - folds, due to the gluings. (d-f) The ‘spines’ of these glued folds formed by replacing each generalised *θ* skeleton by a vertex, joined by edges joining glued skeletons.

Generic gluings can be very complex. Each gluing has an associated ‘spine’, whose vertices denote any even/odd genus *θ*_z_ fold and whose edges describe pairs of glued folds. (The spines for the gluings in Fig. 34(a-c) are shown in Fig. 34(d-f).) All-parallel folds are possible for arbitrary spine graphs decorated with even-genus *θ*_2*k*+2_ folds, including those with cycles, such as the fold in Fig. 34(c). All-antiparallel folds are realised provided the spine is a (cycle-free) tree, decorated with odd-genus *θ*_2*k*+1_ folds. The spine topology is reflected in the circular diagram for the composite fold, evident in the circular diagrams in Fig. 34(a-c).

### Folds and antifolds

The contraction operation - merging nested annular ribbons and closing fans of moebius ribbons - can be reversed, to give an expanded (uncontracted) circular diagram for any fold, formed by splitting a double-helix (with total twist *t_i_*) into a series of helices of arbitrary twists *t_ij_*, where ∑_*j*_*t_ij_* = *t_i_*. A convenient expansion is formed by splitting double-helices with twist *t_i_* ≠ 0 into |*t_j_*| identical unit-twist helices *t_ij_*, where 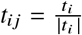. If the contracted double-helix is untwisted (*t_i_* = 0), we split it into a pair of unit twists of opposite sign, ± 1. The resulting circular diagram encodes an expanded fold, denoted 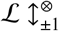. For example, the 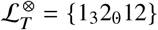 fold expands to 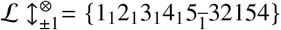 (Fig. 35). The polarised graph of the expanded fold is formed by replacing each vertex in the graph of the contracted fold by multiple vertices connected ‘in series’ along the plumbline of the original graph.

**Figure 35:**
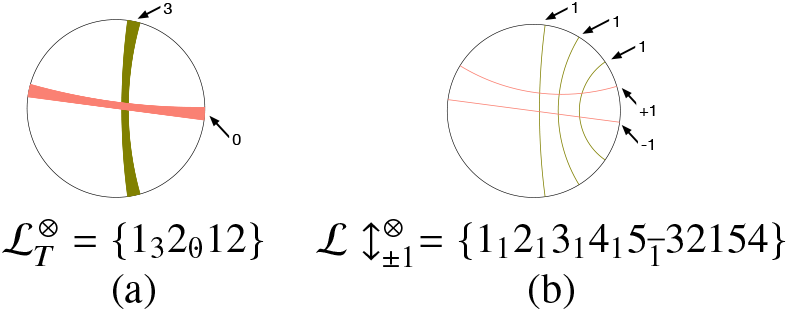
(a) A contracted fully-flagged fold whose ribbons have twists 3 and 0. (b) Circular diagram of the expanded fully-flagged fold 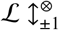, containing duplexed with unit twists only.

If all plumblines in the polarised strand graph of the expanded fold are switched from *NS* to *EW* orientations and the signs of those unit twists are reversed, the fold is converted to its ‘antifold’. A relaxed fold, whose complementary black and white skeletons define conventional junctions and anti-junctions, has a relaxed antifold, with conventional junctions and anti-junctions at vertices of its white and black skeleton respectively. Expanded fold/antifold pairs have the same bare (unflagged) fold label 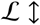. Orientation flags of all unit-twist ribbons are switched between moebius and annular (and vice versa) in transforming the fold into its anti-fold, whereas parity flags are conserved, since the twists |*t_i_*| = 1 are switched between 1 and 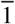 or vice versa, retaining odd parity. In summary, switching from the expanded fold to its expanded antifold effects three simultaneous structural permutations: 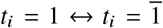, moebius ↔ annular ribbons and conventional junction ↔ anti-junctions. (Meso-junctions, present in necessarily unrelaxed two-colored fold embeddings are unchanged.) Therefore fold/antifold pairs have the following characteristics: (i) all-parallel folds have all-antiparallel antifolds; (ii) relaxed folds have relaxed antifolds; (iii) *homochiral* folds, whose constituent double-helices are all left- or all right-handed, have (right- or left-handed) homochiral antifolds. For example, a fold 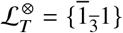 illustrated in Fig. 36(a,b,d), has expanded label 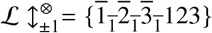, so the expanded antifold has label 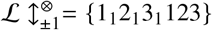. The expanded antifold is already contracted, since it contains a fan of annular ribbons. In this example, both the fold and its antifold are low-genus relaxed folds (with generic labels 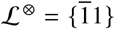 and {123123}, listed in Table 2). However, low-genus folds need not induce low-genus antifolds. Consider the (relaxed) fold-antifold pair due to the 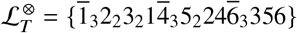 fold, shown in Fig. 37, whose circular diagram is redrawn in Fig. 37(a). The circular diagram of the expanded fold, with fifteen (unit-twist) ribbons shown in Fig. 37(b), gives the polarised strand graph and simplest strand embedding in Figs. 37(c-e). The circular diagram of the expanded antifold also contains fifteen ribbons, whose orientation flags are switched, as in Fig. 37(f). The corresponding polarised strand graph is shown in Fig. 37(g). Figs. 37(e) and (i) demonstrate a generic feature of relaxed fold-antifold pairs: their one-colored skeletons are (two-dimensional) dual graphs of each other. That duality leads to the simple relation between the genus of the simplest embedding of the expanded fully-flagged fold *g*(*F*), where 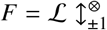 denotes the fold, and the genus of its expanded antifold, 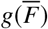:

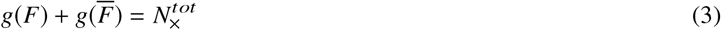

**Figure 36:**
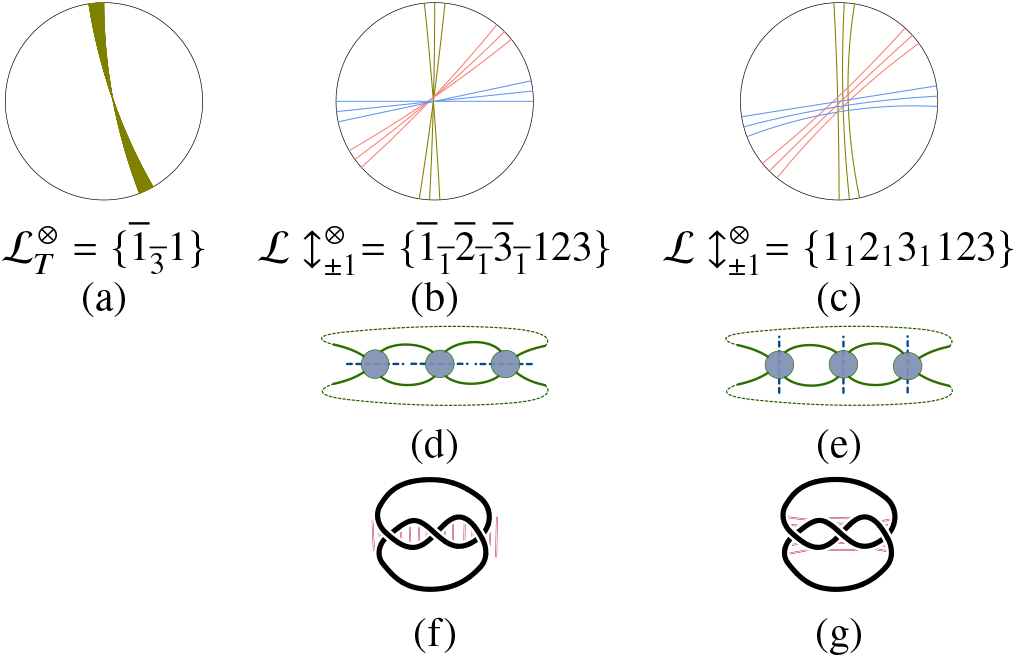
(a) Circular diagram of a contracted fold. (b) Its expanded diagram: a fan of three moebius ribbons, each with twist 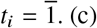. (c) Its antifold: a fan of three annular ribbons and twists *t*_1_ = 1. (d,e) Embeddings of the polarised strand graphs of the realxed fold and its relaxed antifold. (f,g) Strand windings corresponding to the embeddings in (d,e). Base-pairing between complementary nucleotides is indicated by red rungs.

**Figure 37:**
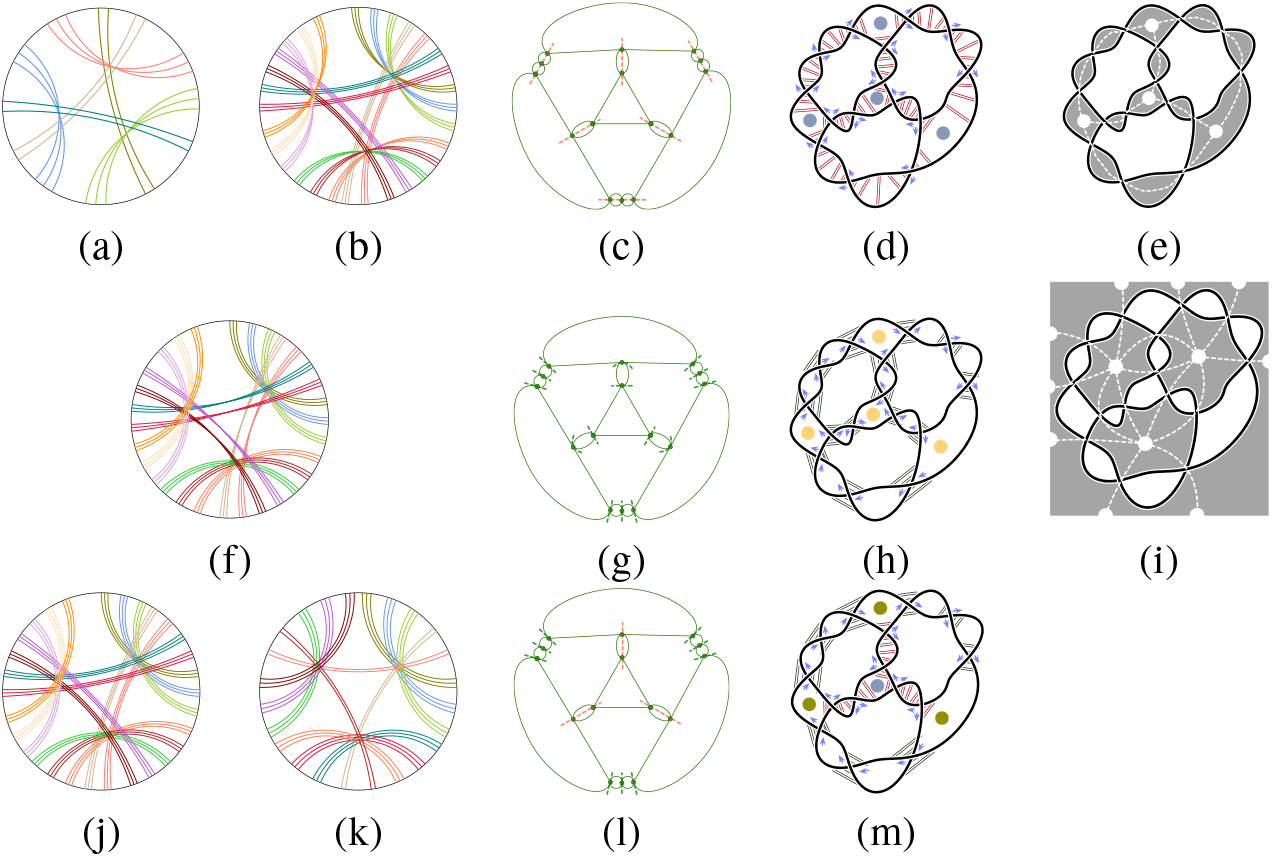
(a) Circular diagram of a contracted fold 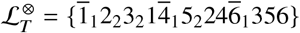 which shares the fully-flagged label of the fold in Fig. 3. (b) The circular diagram of the expanded fold, with fifteen unit twist ribbons. (c) Its polarised strand graph. (d) Simplest embedding of the duplexed strand winding, with arrows marking one polar orientation of the closed strand, revealing parallel and antiparallel double-helices. Grey dots denote three-way conventional junctions. (e) The skeleton of this relaxed fold (*cf*. Fig. 3(d)). (f) The circular diagram of its antifold, which cannot be further contracted. (g) The repolarised strand graph of fold (g). (h) The resulting duplexed fold, with three-way anti-junctions marked by yellow dots whose skeleton is shown in (i) (drawn as a Schlegel diagram, whose partial white vertices on the boundary glue to form a single vertex). (j) An isomorphic fold, whose expanded circular diagram contains fifteen annular ribbons. (k) Contraction of the circular diagram in (j). (l) The polarised strand graph. (m) Simplest embedding of the resulting all-parallel strand winding, with a conventional three-way junction (grey dot in the centre) and three-way meso-junctions (green dots).

Here 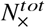 is the total number of crossings in the actual fold winding, rather than its polarised strand graph. If *F* is an (a)- or (b)-type fold, 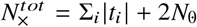, where twists *t_i_* are summed over all contracted double-helices, and *N*_0_ is the number of untwisted (but duplexed) helices (with *t_i_* = 0). If *F* is a (c)-type fold, an additional *N*_×_ edge-crossings are present, visible in the polarised strand graph embedding. In those cases, 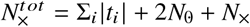. It follows from equation 3 that the genus of an antifold 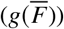 can be smaller, equal or larger than that of the original fold (*g*(*F*)), depending on the sign of 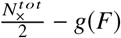.

Though the skeletons of fold-antifold pairs may have very different genus, the pair nevertheless have identical strand tangling, ignoring base-paring. For example, strand windings in Figs. 36(f,g) are identical. (However, base-pairings are very different.) That tangling equivalence of fold-antifold pairs rests on an inherent ambiguity in the direction of a helical axis within a double-helix containing just a single half-twist (*t* = ±1), evident in Fig. 38. The helical axis can be interpreted as *NS* or *EW*, provided the sign of the unit twist is reversed if the axis is reversed. Therefore the fold 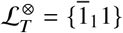 and antifold 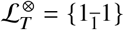 are equivalent.

**Figure 38:**
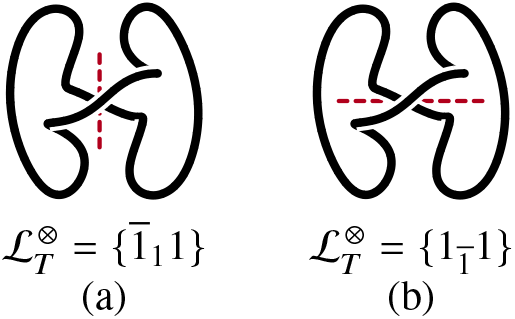
A double-helix with a single half-twist can be interpreted as (a) a vertically-aligned helix (along the red axis) with twist *t* = 1 or (b) a horizontally-aligned helix with twist 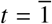.

It follows that if all plumblines are reoriented in an extended fold 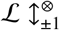, and the twist sign reversed, the tangling or *knotting* of the helical strands - determined by the over-under crossings of strands - is unchanged. Folds and antifolds therefore share identical strand tangling.

#### Isomorphic folds

Fold-antifold pairs belongs to a much larger suite of *isomorphic* folds 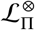, whose expanded folds 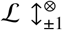, have identical *un*polarised strand graphs. Whereas all plumbines are swapped in going from a fold to its antifold, generic isomorphic folds are formed by swapping a subset of plumblines in the polarised strand graph of the expanded fold only. Since these isomorphic folds share the same helical winding and differ only in the arrangements of complementary base-pairs, expanded labels of all isomorphic folds share identical unflagged label, 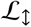 (a string of length 2Σ_i_ |*t*_i_|). Since any subset of plumblines in the expanded polarised strand graph can be flipped, 2^∑_i_|*t*_i_^ isomorphic folds are possible, though duplications are common, due to equivalent folds (with identical fully-flagged canonical fold labels). For example, the one-ribbon fold/antifold pair in Fig. 36, for which Σ_i_|*t*_i_| = 3, belongs to the family of isomorphic folds, including four homochiral folds shown in Fig. 39 with common unflagged label 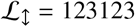.

**Figure 39:**
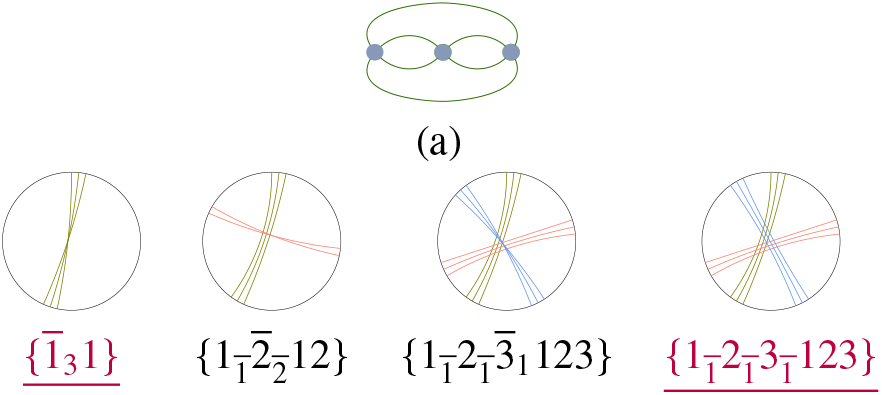
Contracted circular diagrams and labels 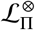 for four isomorphic folds whose expanded folds 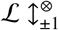 share common unflagged label L ↕= 123123 as well as the unpolarised strand graph in (a), where each vertex hosts a single-twist double-helix. The pair of underlined fold labels colored purple are relaxed (i.e. one-colored) and homochiral; 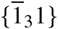 is also all-parallel and 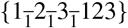 is all-antiparallel.

There are seven isomorphic homochiral folds with common unflagged label for their extended diagrams 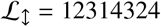, shown in Fig. 40. One of those folds, 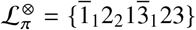, has an uncrossed one-colored embedding of its polarised strand graph and is therefore relaxed. This family has just one relaxed fold, 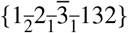, since its antifold is identical.

**Figure 40:**
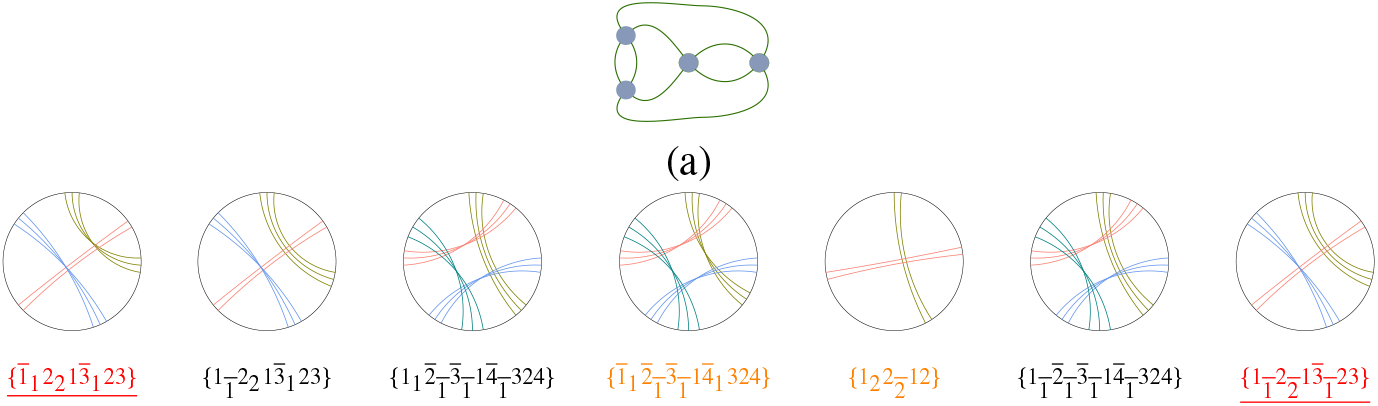
Circular diagrams and labels 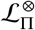 for all contracted homochiral isomorphic folds sharing the expanded unpolarised strand graph 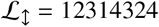 and unpolarised strand graph in (a), where each vertex hosts a single-twist double-helix. The pair of underlined fold labels colored red are relaxed (i.e. one-colored) and homochiral. All-parallel and all-antiparallel folds are colored orange.

In contrast, the two-ribbon fan fold 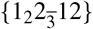 with Σ_i_|*t*_i_| = 5 belongs to a family of isomorphic folds with unflagged label 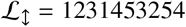, including those whose contracted circular diagrams are illustrated in Fig. 41.

**Figure 41:**
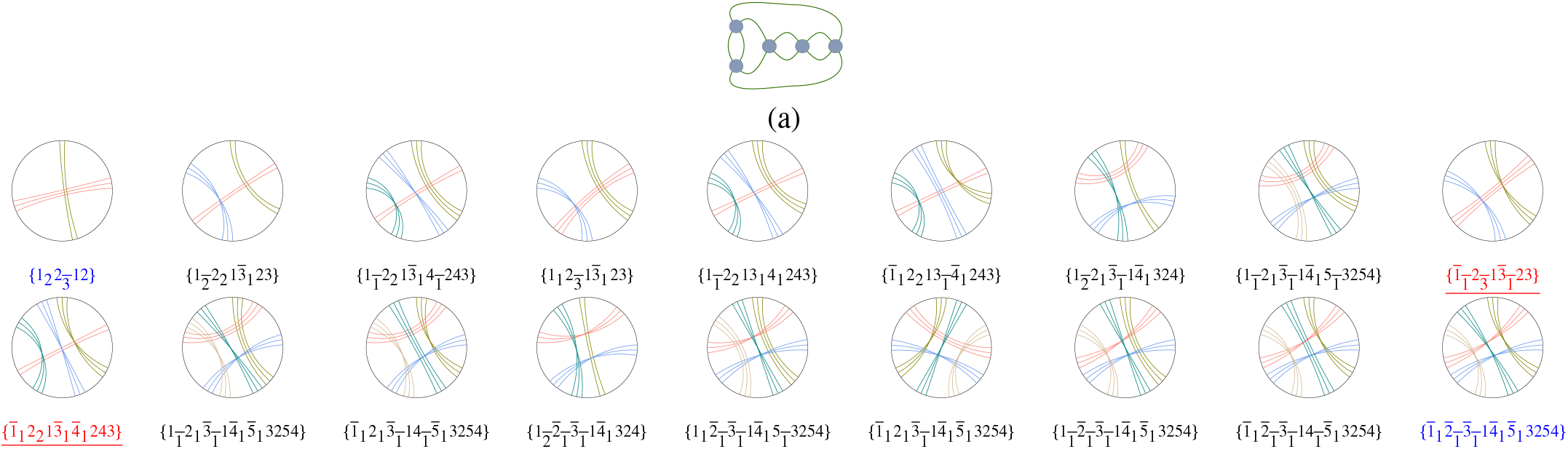
Circular diagrams and labels 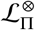 for contracted isomorphic folds sharing the expanded unflagged label 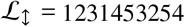 and unpolarised strand graph in (a). The pair of underlined fold labels colored red are homochiral and relaxed; the all-parallel fold and its all-antiparallel antifold are colored blue.

Any family of isomorphic (a)-type folds includes all three fold/antifold pairs introduced above: relaxed (with junctions or anti-junctions only), homochiral (left- or right-handed) and all-parallel or all-antiparallel. These often coincide for simpler folds. For example, the isomorphic folds in Fig. 41 include a fold/antifold pair of homochiral and relaxed folds (marked in red in the figure) and all-parallel/all-antiparallel folds (marked in blue). In contrast, two of the four isomorphic folds in (Fig. 39) describe the homochiral, all-(anti)parallel and relaxed fold/antifold pairs (marked in purple in the figure.)

These examples confirm that identical knots can be realised by very different - though isomorphic - duplexed folds. Conversely, can a single fold generate different knots?

### Fold knotting

The knotting of an embedded fold induced by duplexing is *a priori* delicate, given the theoretical possibility that *any* knot can be realised by *any* fold, provided the non-duplexed strand fragments unwind the windings induced by strand duplexing and rewind the target knot. More technically, knotting is the result of all crossings in the embedding 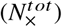 including ∑_*i*_|*t_i_*|) strand crossings within the duplex and *N*_×_ ‘shadow’ edge-crossings in the embedding of the strand graph. While the duplex crossings are dictated by its twist *t_i_*, edge-crossings can adopt either embedding as shown in Fig. 2. Our ‘simplest’ embeddings minimise *N*_×_: in practice a fold can be embedded with *N*_×_ + 2*z* crossings, where *z* is any integer (*cf*. Fig. 20). As that number grows, so does the variety of possible strand knots in the fold, since each shadow edge-crossing (labelled by some index *k*, whose twists we denote *τ_k_* in contrast to twists within duplexes, denoted *t_i_*) can be assigned an effective twist of *τ_k_* = 1 or 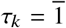, retuning all net twists at will. Nevertheless, the spectrum of knots realised by helical duplexing is likely to be restricted, since some specific combinations of twists at edge-crossings, *τ_k_*, are more likely than others. For example, in the absence of any interactions within the string beyond secondary duplexing, the distribution of various edge-crossings τ_k_, is likely to be random. Our aim is to uncover the simplest strand tangling that can result by strand duplexing. The ‘simplest’ fold embeddings, which are induced by secondary interactions (assumed here to be due to Watson-Crick complementarity), are those resulting from the embedding(s) of the polarised strand graph for the fold which minimise the number of edge-crossings, *N*_×(*i*)_.

Within that scenario, all (a)- and (b)-type folds, with *N*_×_ = 0, lead to unique knots. However, other embeddings of those same folds, with *N*_×_ = 2*z* are also possible, which generate more or less complex knots. For example, the fold 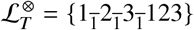 has more general label 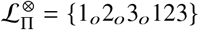. That fold is relaxed (e.g. Fig. 7(c,h)) so *N*_×_ = 0. (Fig. 39) allows embeddings with *N*_×_ + 2*z* = 2*z* realised for (a) *z* = 0 (Fig. 42(a) and *z* = 2 (Figs. 42(b,c,d). The simplest embedding, with *z* = 0 induces the trefoil knot, 3_1_. Further, simplest embeddings of all folds which are isomorphic to 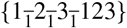, shown in Fig. 39, are also knotted in the same way. More complex embeddings, shown in Figs. 42(b,c,d), induce the unknot 0_1_, trefoil or the cinquefoil knot, 5_1_.

**Figure 42:**
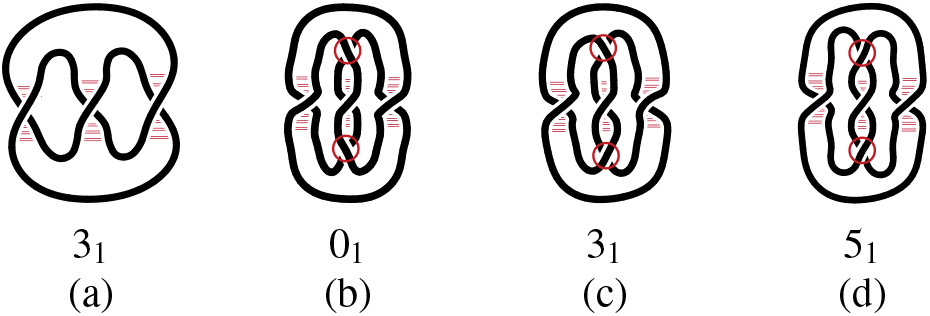
(a-d) Four examples of tangles induced by the 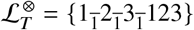 fold with different numbers of edge-crossings in the central duplex: (a) *N*_×(*i*)_ = 0 (*i* = 1, 2, 3), (b-d)*N*_×(1)_ = *N*_×(3)_ = 0, *N*_×(2)_ = 2. The pair of edge-crossings on either side of the duplexed crossing (circled in red) have twists (from top to bottom) (b) 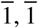, (c) 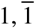 and (d) 1,1. The ‘simplest’ embedding minimises *N*_×_(*i*), namely embedding (a), which forms the trefoil knot (3_1_). The 0_1_ case is unknotted and 5_1_ is the cinquefoil knot.

Strand knotting in (c)-type folds - even assuming their simplest embeddings, for which *N*_×_ > 0 - is less straightforward. For example, 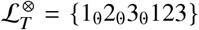 has *N*_×_ = 3, since all three (even) flags are detuned (the relaxed version of this fold has fully-flagged label 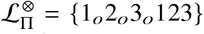, as noted in the previous example). Each edge-crossing *τ_k_* can have arbitrary sign, allowing three possible fold embeddings: the unknot, 0_1_, the ‘right-handed’ handed trefoil, 3_1_, or its enantiomer, 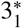, as shown in Fig. 43.

**Figure 43:**
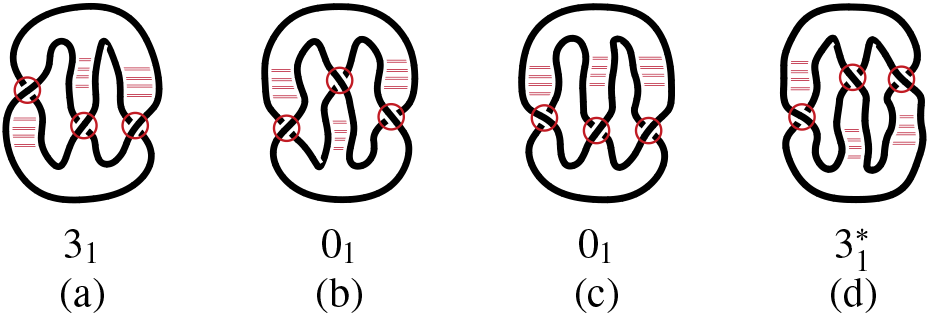
The simplest embeddings of the 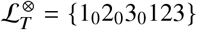 strand winding, with *N*_×_ = 3 (circled in red). 3_1_ and 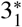 are trefoil enantiomers.

Since each of the *N*_×_ edge-crossings has two alternative twists (*τ_k_* = ±1), simplest embeddings of generic (c)-type folds can realise up to 2^*N*_×_^ different knots.

#### Knots realised by simpler (a)- and (b)-type all-antiparallel and all-parallel contracted folds

Table 7 enumerates all (a)- and (b)-type folds in order of increasing crossing index 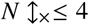 for their extended folds, where 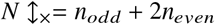, and *n_even_* and *n_odd_* denote the numbers of even- and odd-parity ribbons in the extended fold. 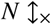 is a useful index of fold complexity, since the tools introduced above allow exhaustive enumeration of uncrossed folds (*N*_×_ = 0) up to any specified value. There are many additions to Table 7 if *N*_×_ = 5. However, among those, none are all −antiparallel and just one of is all-antiparallel, {1_*o*_2_*o*_3_*o*_4_*o*_5_*o*_ 12345} (*cf*. Fig. 32(c)). A useful catalog of all-parallel and all-antiparallel folds with *N*_×_ ≤ 5 is given in Table 9.

**Table 9:**
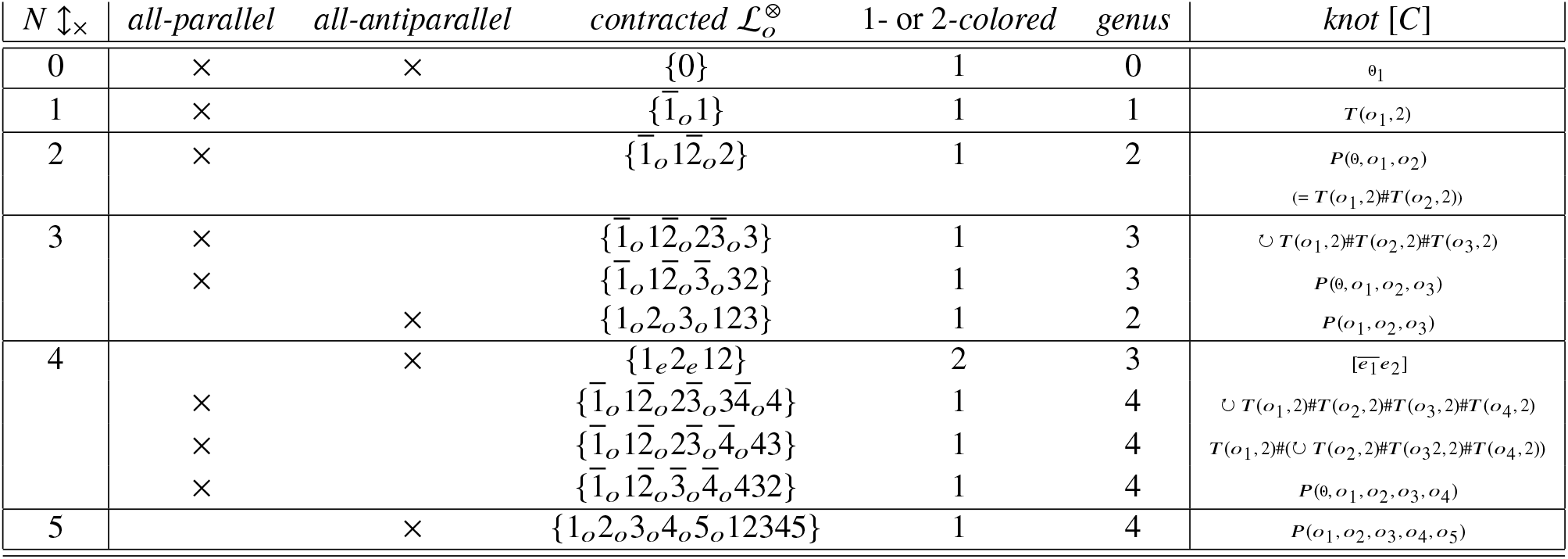
Contracted (a)- and (b)-type fully-flagged folds whose constituent double-helices are either all-parallel, or all-antiparallel. Folds are ranked by 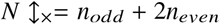 (the number of odd- and even-parity helices), and classified as (a)- or (b)-types (see Table 5). Knots are specified as follows. Twists on ribbon *i* are labelled *e_i_* or *o_i_*, depending on their parity. The trivial unknot is denoted 0_1_. *T*(*o*_1_, 2) denotes the (*p, q*) torus knot (where *o*_1_ is the (odd) twist in helix 1). *P*(*a, b, c*,…) denotes the generalised pretzel knot with twists *a, b, c*,…. Composite knots of *K*_1_, *K*_2_, etc. are denoted *K*_1_#*K*_2_…. If those are formed by cyclic gluing, they are prefixed by a circular arrow, 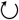. Knots labelled 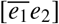 correspond to Conway rational links.

| $N \uparrow_{\times}$ | *all-parallel* | *all-antiparallel* | *contracted* $\mathcal{L}_o^{\otimes}$ | 1- or 2-colored | genus | knot $[C]$ |
| --- | --- | --- | --- | --- | --- | --- |
| 0 | × | × | $\{0\}$ | 1 | 0 | $0_1$ |
| 1 | × | | $\{\overline{1}_o 1\}$ | 1 | 1 | $T(o_1, 2)$ |
| 2 | × | | $\{\overline{1}_o \overline{12}_o 2\}$ | 1 | 2 | $P(0, o_1, o_2)$<br>$(= T(o_1, 2) \# T(o_2, 2))$ |
| 3 | × | | $\{\overline{1}_o \overline{12}_o \overline{23}_o 3\}$ | 1 | 3 | $\cup T(o_1, 2) \# T(o_2, 2) \# T(o_3, 2)$ |
| | × | | $\{\overline{1}_o \overline{12}_o \overline{3}_o 32\}$ | 1 | 3 | $P(0, o_1, o_2, o_3)$ |
| | | × | $\{1_o 2_o 3_o 123\}$ | 1 | 2 | $P(o_1, o_2, o_3)$ |
| 4 | | × | $\{1_e 2_e 12\}$ | 2 | 3 | $[\overline{e_1 e_2}]$ |
| | × | | $\{\overline{1}_o \overline{12}_o \overline{23}_o \overline{34}_o 4\}$ | 1 | 4 | $\cup T(o_1, 2) \# T(o_2, 2) \# T(o_3, 2) \# T(o_4, 2)$ |
| | × | | $\{\overline{1}_o \overline{12}_o \overline{23}_o \overline{4}_o 43\}$ | 1 | 4 | $T(o_1, 2) \# (\cup T(o_2, 2) \# T(o_3, 2) \# T(o_4, 2))$ |
| | × | | $\{\overline{1}_o \overline{12}_o \overline{3}_o \overline{4}_o 432\}$ | 1 | 4 | $P(0, o_1, o_2, o_3, o_4)$ |
| 5 | | × | $\{1_o 2_o 3_o 4_o 5_o 12345\}$ | 1 | 4 | $P(o_1, o_2, o_3, o_4, o_5)$ |

Recall that the simplest embeddings of all the folds in Table 9 describe unique strand knots since *N*_×_ = 0, whose specific type depends on specific twist values, *t_i_*. In fact, the knots belong to reasonably well-known knot families, including prime and composite torus knots, pretzel knots and algebraic tangles and closures of rational tangles, listed accordingly in the Table. Specific knots can be determined via tables of simpler members of those families, described in part in (39); see also (40, 41). (Specific knots induced by all-antiparallel folds with small (positive) twists will be explicitly identified later (Table 14).)

**Table 10:**
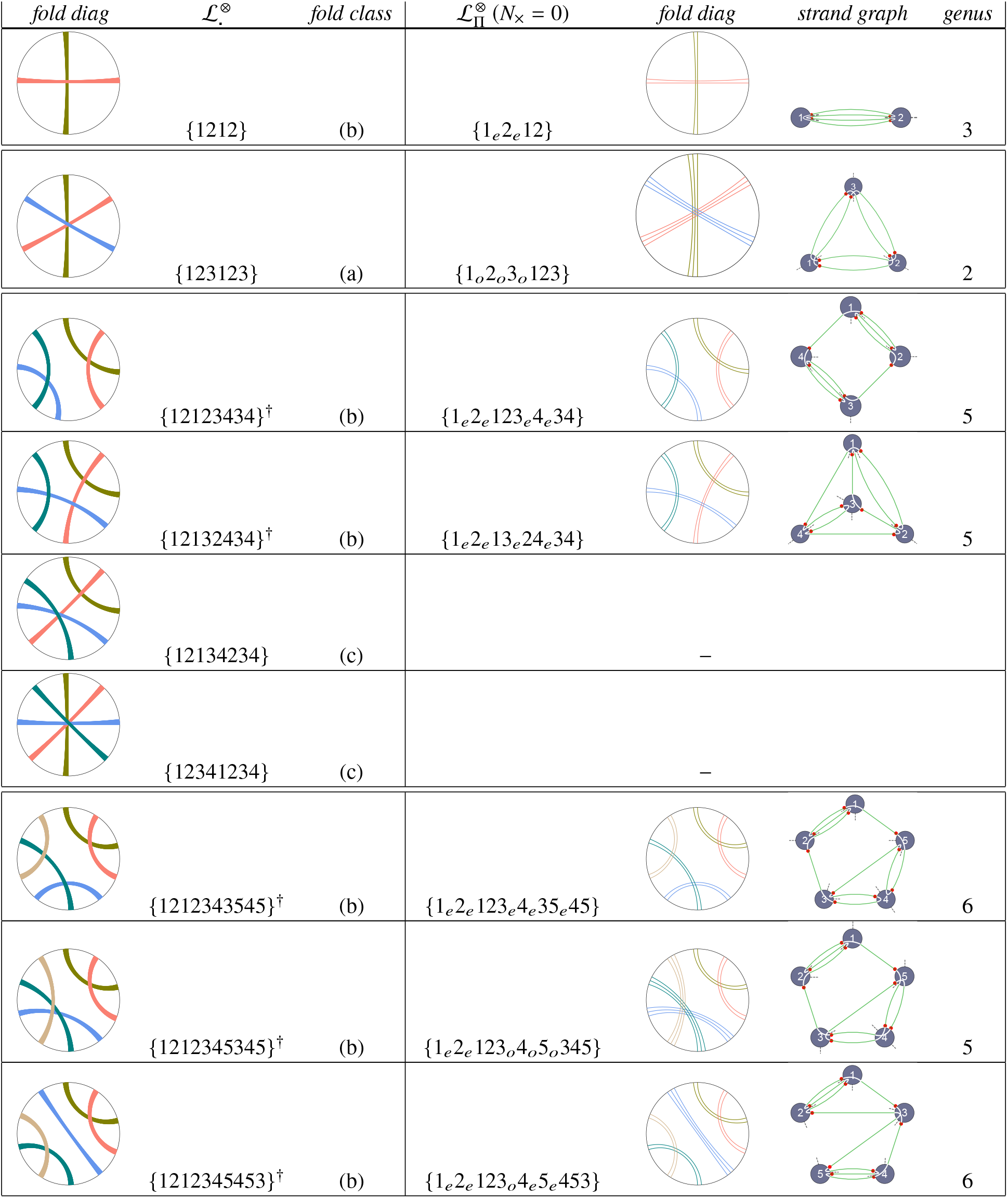
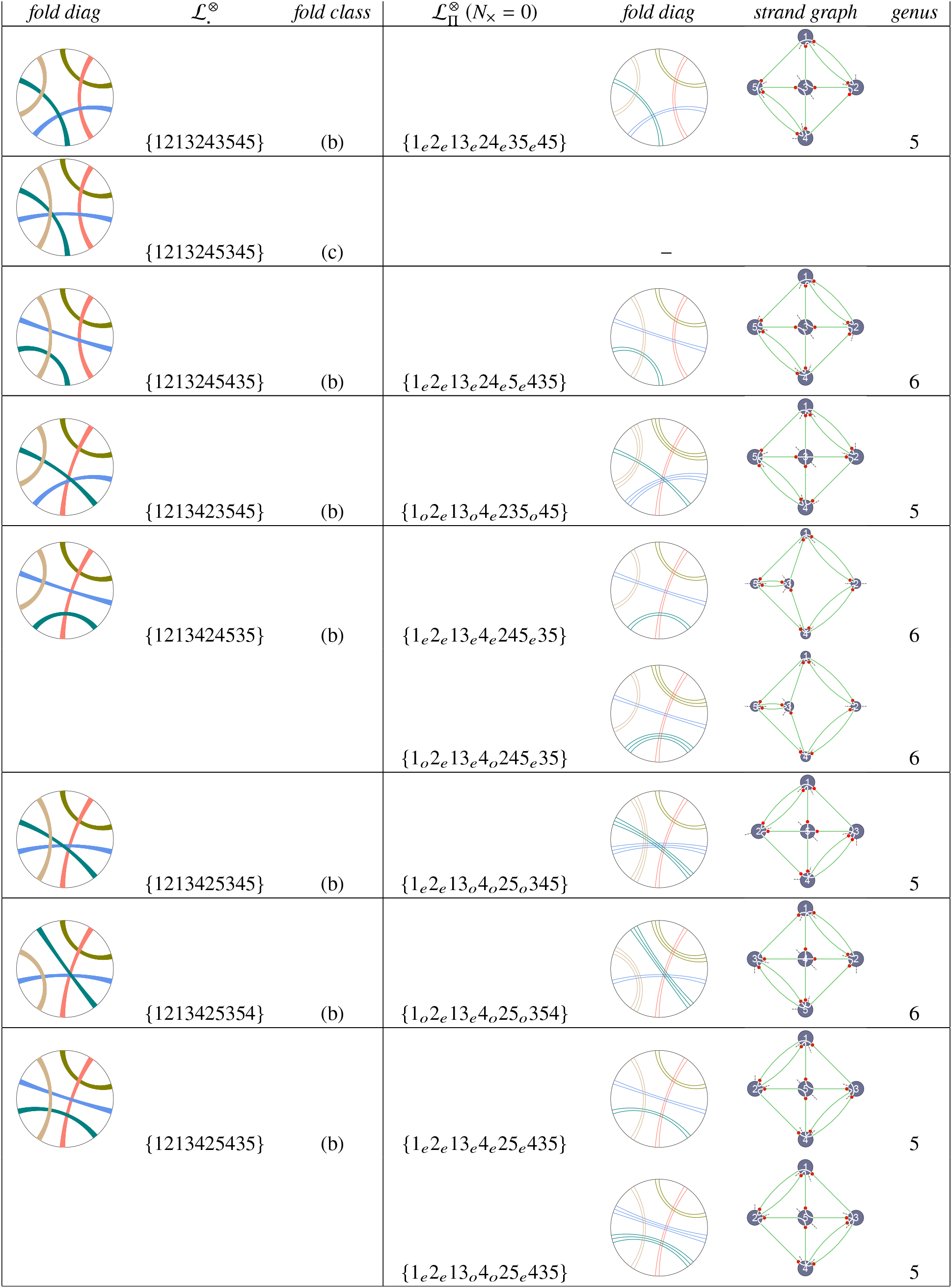
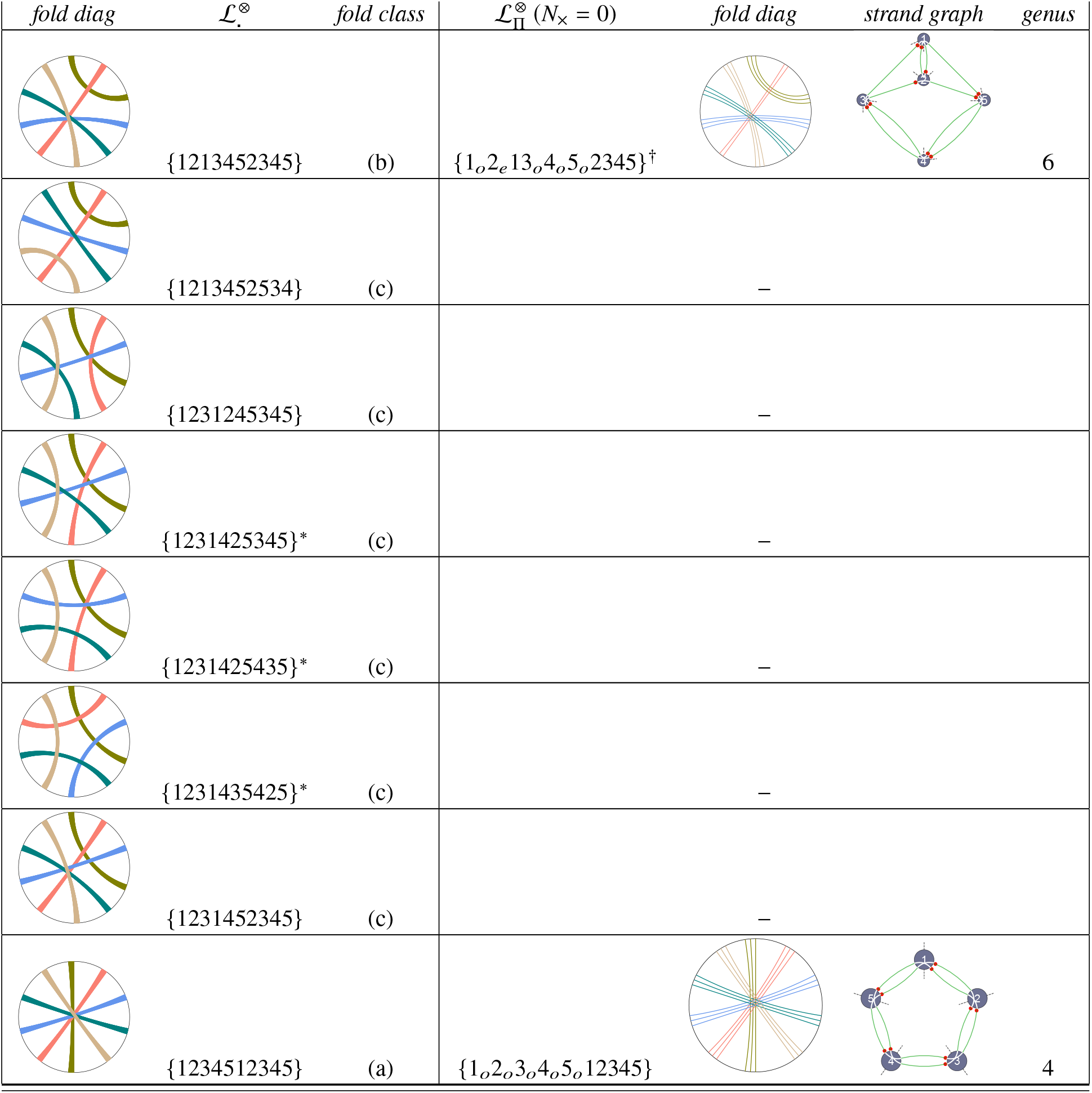
All contracted semi-flagged fold labels 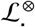 containing up to five antiparallel double-helices. Most folds are prime; composite folds are marked with the superscript^†^. Specific parity flags (*π_i_* = *o* or *e*) can be appended to (a)- and (b)-type folds to give fully-flagged folds 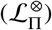 with *N*_×_ = 0. Those crossing-free embeddings of the polarised strand graphs are shown with crossed or uncrossed strands drawn in white within each graph vertex, representing odd or even parity double-helices. Red dots confirm that all double-helices are antiparallel (*cf*. Fig. 4). The three semi-flagged folds whose labels are appended with * superscripts induce the non-planar graph *K*_5_. (The fold marked^†^ is discussed earlier, see Fig. 30(a).)

| fold diag | $\mathcal{L}_\Pi^\otimes$ | fold class | $\mathcal{L}_\Pi^\otimes (N_\times = 0)$ | fold diag | strand graph | genus |
| --- | --- | --- | --- | --- | --- | --- |
|  | {1212} | (b) | {1<sub>e</sub>2<sub>e</sub>12} |  |  | 3 |
|  | {123123} | (a) | {1<sub>o</sub>2<sub>o</sub>3<sub>o</sub>123} |  |  | 2 |
|  | {12123434}<sup>†</sup> | (b) | {1<sub>e</sub>2<sub>e</sub>123<sub>e</sub>4<sub>e</sub>34} |  |  | 5 |
|  | {12132434}<sup>†</sup> | (b) | {1<sub>e</sub>2<sub>e</sub>13<sub>e</sub>24<sub>e</sub>34} |  |  | 5 |
|  | {12134234} | (c) | — | — | — | — |
|  | {12341234} | (c) | — | — | — | — |
|  | {1212343545}<sup>†</sup> | (b) | {1<sub>e</sub>2<sub>e</sub>123<sub>e</sub>4<sub>e</sub>35<sub>e</sub>45} |  |  | 6 |
|  | {1212345345}<sup>†</sup> | (b) | {1<sub>e</sub>2<sub>e</sub>123<sub>o</sub>4<sub>o</sub>5<sub>o</sub>345} |  |  | 5 |
|  | {1212345453}<sup>†</sup> | (b) | {1<sub>e</sub>2<sub>e</sub>123<sub>o</sub>4<sub>e</sub>5<sub>e</sub>453} |  |  | 6 |

| *fold diag* | $\mathcal{L}_\cdot^\otimes$ | *fold class* | $\mathcal{L}_\Pi^\otimes (N_\times = 0)$ | *fold diag* | *strand graph* | *genus* |
| --- | --- | --- | --- | --- | --- | --- |
| | $\{1213243545\}$ | (b) | $\{1_e 2_e 13_e 24_e 35_e 45\}$ | | | 5 |
| | $\{1213245345\}$ | (c) | — | — | — | — |
| | $\{1213245435\}$ | (b) | $\{1_e 2_e 13_e 24_e 5_e 435\}$ | | | 6 |
| | $\{1213423545\}$ | (b) | $\{1_o 2_e 13_o 4_e 235_o 45\}$ | | | 5 |
| | $\{1213424535\}$ | (b) | $\{1_e 2_e 13_e 4_e 245_e 35\}$ | | | 6 |
| | | | $\{1_o 2_e 13_e 4_o 245_e 35\}$ | | | 6 |
| | $\{1213425345\}$ | (b) | $\{1_e 2_e 13_o 4_o 25_o 345\}$ | | | 5 |
| | $\{1213425354\}$ | (b) | $\{1_o 2_e 13_e 4_o 25_o 354\}$ | | | 6 |
| | $\{1213425435\}$ | (b) | $\{1_e 2_e 13_e 4_e 25_e 435\}$ | | | 5 |
| | | | $\{1_e 2_e 13_o 4_o 25_e 435\}$ | | | 5 |

| *fold diag* | $\mathcal{L}_\cdot^\otimes$ | *fold class* | $\mathcal{L}_\Pi^\otimes (N_\times = 0)$ | *fold diag* | *strand graph* | *genus* |
| --- | --- | --- | --- | --- | --- | --- |
|  | {1213452345} | (b) | {1<sub>o</sub>2<sub>e</sub>13<sub>o</sub>4<sub>o</sub>5<sub>o</sub>2345}<sup>†</sup> |  |  | 6 |
|  | {1213452534} | (c) |  | — |  |  |
|  | {1231245345} | (c) |  | — |  |  |
|  | {1231425345}<sup>*</sup> | (c) |  | — |  |  |
|  | {1231425435}<sup>*</sup> | (c) |  | — |  |  |
|  | {1231435425}<sup>*</sup> | (c) |  | — |  |  |
|  | {1231452345} | (c) |  | — |  |  |
|  | {1234512345} | (a) | {1<sub>o</sub>2<sub>o</sub>3<sub>o</sub>4<sub>o</sub>5<sub>o</sub>12345} |  |  | 4 |

**Table 11:** All nontrivial fully-flagged linear folds of an open single strand 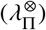 containing up to four contracted antiparallel double-helices and fold diagrams whose strand closure yields contracted fully-flagged (a)- or (b)-type folds (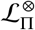, among the first four rows in Table 10. (The simplest fold, with a single antiparallel double-helix or series of double-helices interrupted by bulges, closes to give the circular fold {11}, which contracts to the trivial fold {0}.) The folds are indexed 1 – 15 (where applicable, appended with the pseudofold name, *H, K, L* or *M* from (21)). (^†^ This fold is a composite linear fold; all others are prime folds.)

| *fold ID* | $\lambda_{\Pi}^{\otimes}$ | *lin diagram* | *circ diagram* | $\mathcal{L}_{\Pi}^{\otimes}$ | genus | fold type |
| --- | --- | --- | --- | --- | --- | --- |
| 1 ($H$) | $[1_e 2_e 12]$ | | | $\{1_e 2_e 12\}$ | 3 | (b) |
| 2 ($K$) | $[1_e 2_e 13_e 23]$ | | | $\{1_e 2_e 12\}$ | 3 | (b) |
| 3 ($K$) | $[1_o 2_e 13_o 23]$ | | | $\{1_e 2_e 12\}$ | 3 | (b) |
| 4 ($L$) | $[1_o 2_o 3_o 123]$ | | | $\{1_o 2_o 3_o 123\}$ | 2 | (a) |
| 5 | $[1_e 2_e 123_e 4_e 34]^{\dagger}$ | | | $\{1_e 2_e 123_e 4_e 34\}$ | 5 | (b) |
| 6 | $[1_e 2_e 13_e 24_e 34]$ | | | $\{1_e 2_e 13_e 24_e 34\}$ | 5 | (b) |
| 7 | $[1_e 2_e 13_e 4_e 342]$ | | | $\{1_e 2_e 123_e 4_e 34\}$ | 5 | (b) |
| 8 ($M$) | $[1_e 2_o 3_o 14_o 234]$ | | | $\{1_o 2_o 3_o 123\}$ | 2 | (a) |
| 9 ($M$) | $[1_o 2_o 3_o 14_e 234]$ | | | $\{1_o 2_o 3_o 123\}$ | 2 | (a) |
| 10 | $[1_e 2_e 3_e 14_e 342]$ | | | $\{1_e 2_e 13_e 24_e 34\}$ | 5 | (b) |
| 11 | $[1_e 2_e 3_e 234_e 14]$ | | | $\{1_e 2_e 123_e 4_e 34\}$ | 5 | (b) |
| 12 | $[1_e 2_e 3_e 24_e 143]$ | | | $\{1_e 2_e 13_e 24_e 34\}$ | 5 | (b) |
| 13 | $[1_e 2_e 3_e 24_e 314]$ | | | $\{1_e 2_e 13_e 24_e 34\}$ | 5 | (b) |
| 14 | $[1_e 2_e 3_e 4_e 2413]$ | | | $\{1_e 2_e 13_e 24_e 34\}$ | 5 | (b) |
| 15 | $[1_e 2_e 3_e 4_e 3142]$ | | | $\{1_e 2_e 13_e 24_e 34\}$ | 5 | (b) |

**Table 12:**
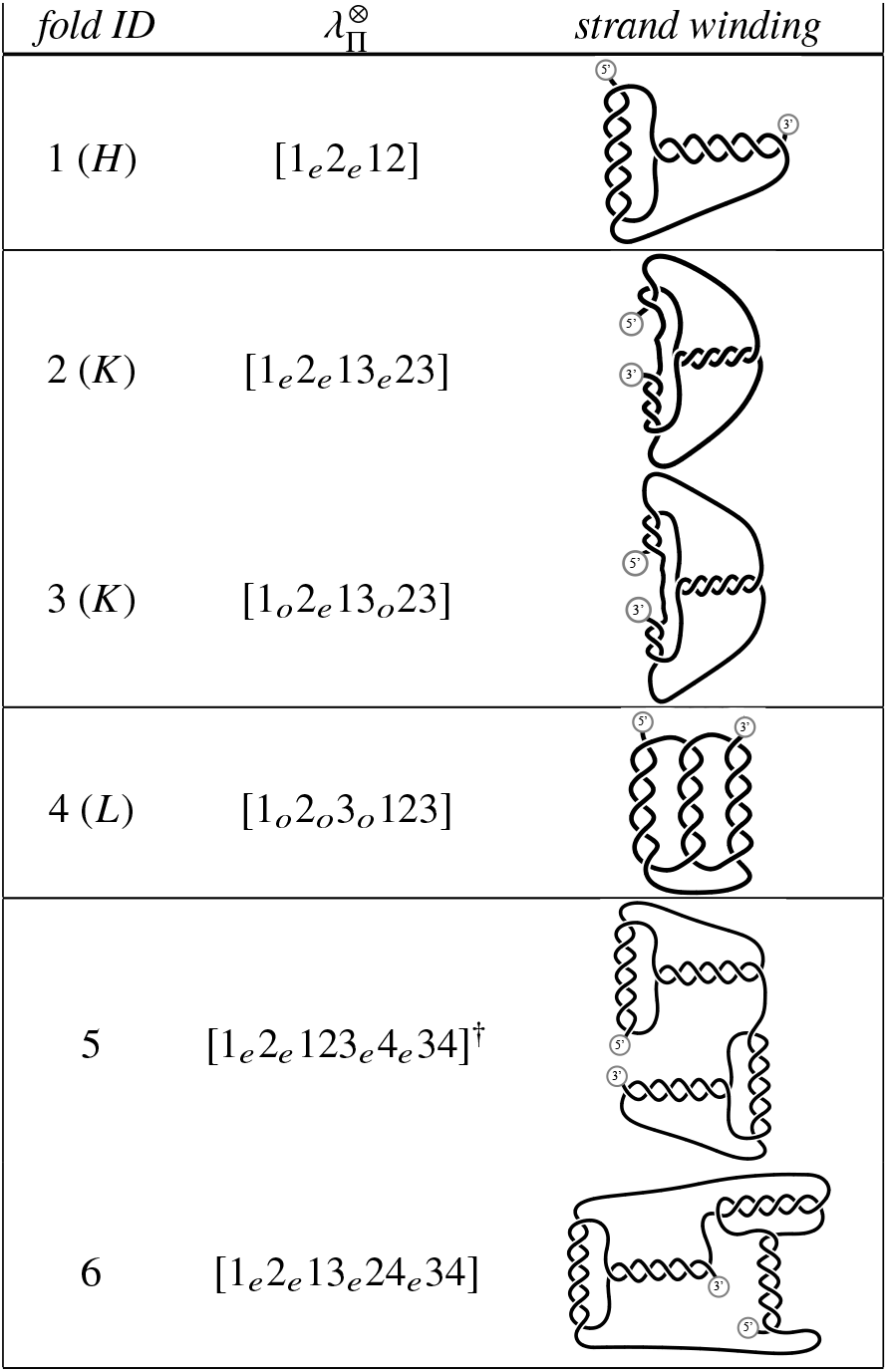
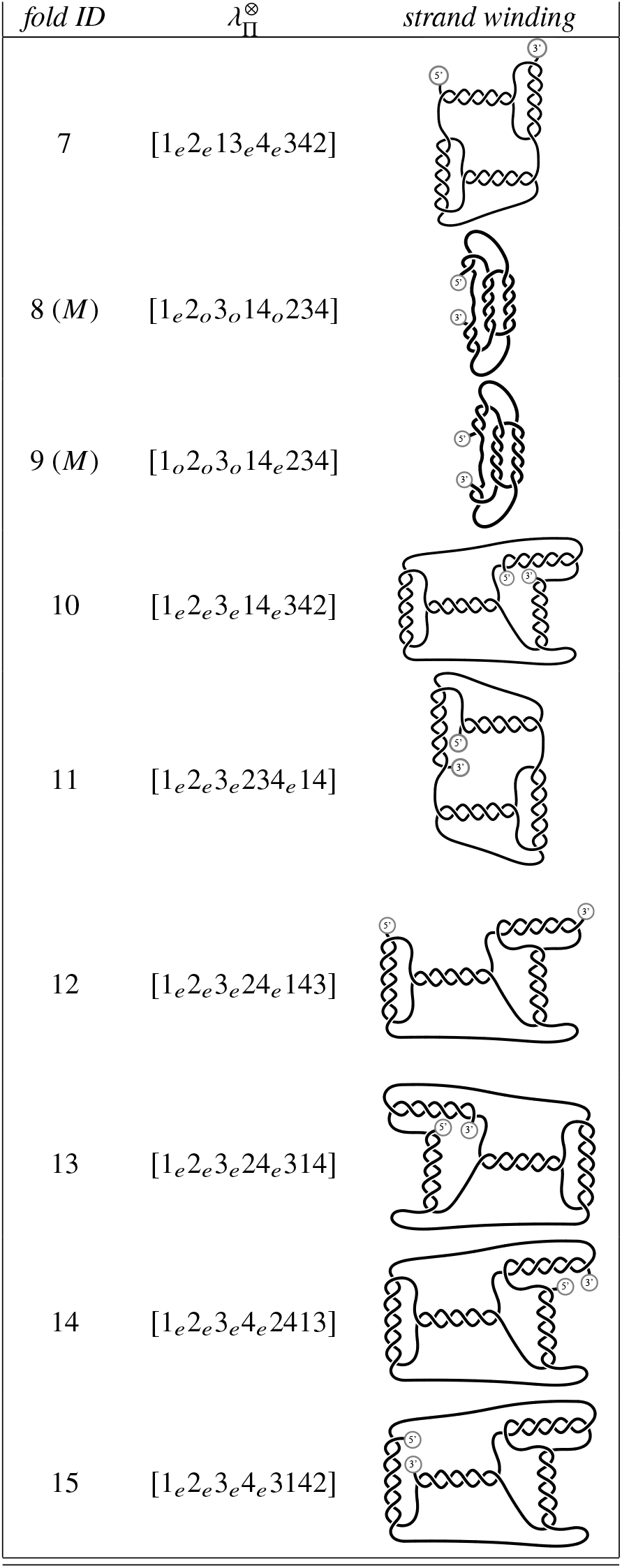
Simple embeddings without edge-crossings (*N*_×_ = 0) of strand windings of all 4-helix contracted linear folds in Table 11. The windings contain duplexed (double-stranded) domains and connecting single-stranded regions. Duplexes of odd or even parity are represented by right-handed double-helices with 5 and 6 half-twists (*t_i_*) respectively. They can be replaced by duplexes with arbitrary (right- or left-handed) twists (including *t_i_* = 0), provided the parity is conserved. The embeddings are chosen for schematic clarity rather than adherence to actual ssRNA structures. However, any structure must be topologically equivalent to those embeddings (admitting modified twists), in the sense that its polarised strand graph is identical. (That allows, for example, more complex embeddings, produced by phantom moves, which deforming the winding such that single-stranded edges pass through each other.)

| fold ID | $\lambda_\Pi^\otimes$ | strand winding |
| --- | --- | --- |
| 1 ($H$) | $[1_e 2_e 12]$ | |
| 2 ($K$) | $[1_e 2_e 13_e 23]$ | |
| 3 ($K$) | $[1_o 2_e 13_o 23]$ | |
| 4 ($L$) | $[1_o 2_o 3_o 123]$ | |
| 5 | $[1_e 2_e 123_e 4_e 34]^\dagger$ | |
| 6 | $[1_e 2_e 13_e 24_e 34]$ | |

| *fold ID* | $\lambda_{\Pi}^{\otimes}$ | *strand winding* |
| --- | --- | --- |
| 7 | $[1_e 2_e 13_e 4_e 342]$ | |
| 8 (*M*) | $[1_e 2_o 3_o 14_o 234]$ | |
| 9 (*M*) | $[1_o 2_o 3_o 14_e 234]$ | |
| 10 | $[1_e 2_e 3_e 14_e 342]$ | |
| 11 | $[1_e 2_e 3_e 234_e 14]$ | |
| 12 | $[1_e 2_e 3_e 24_e 143]$ | |
| 13 | $[1_e 2_e 3_e 24_e 314]$ | |
| 14 | $[1_e 2_e 3_e 4_e 2413]$ | |
| 15 | $[1_e 2_e 3_e 4_e 3142]$ | |

**Table 13:**
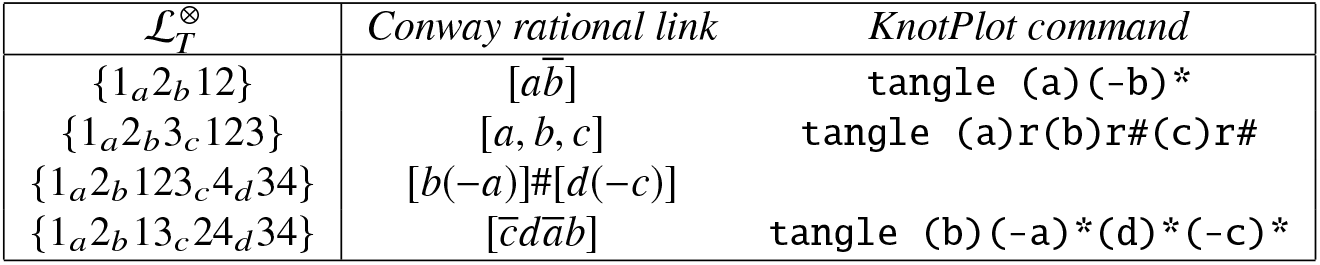
Classification of all distinct circular folds with up to four duplexes in Table 10 as Conway rational links. The total number of strand crossings associated with each duplex are denoted *a, b, c*,….

**Table 14:**
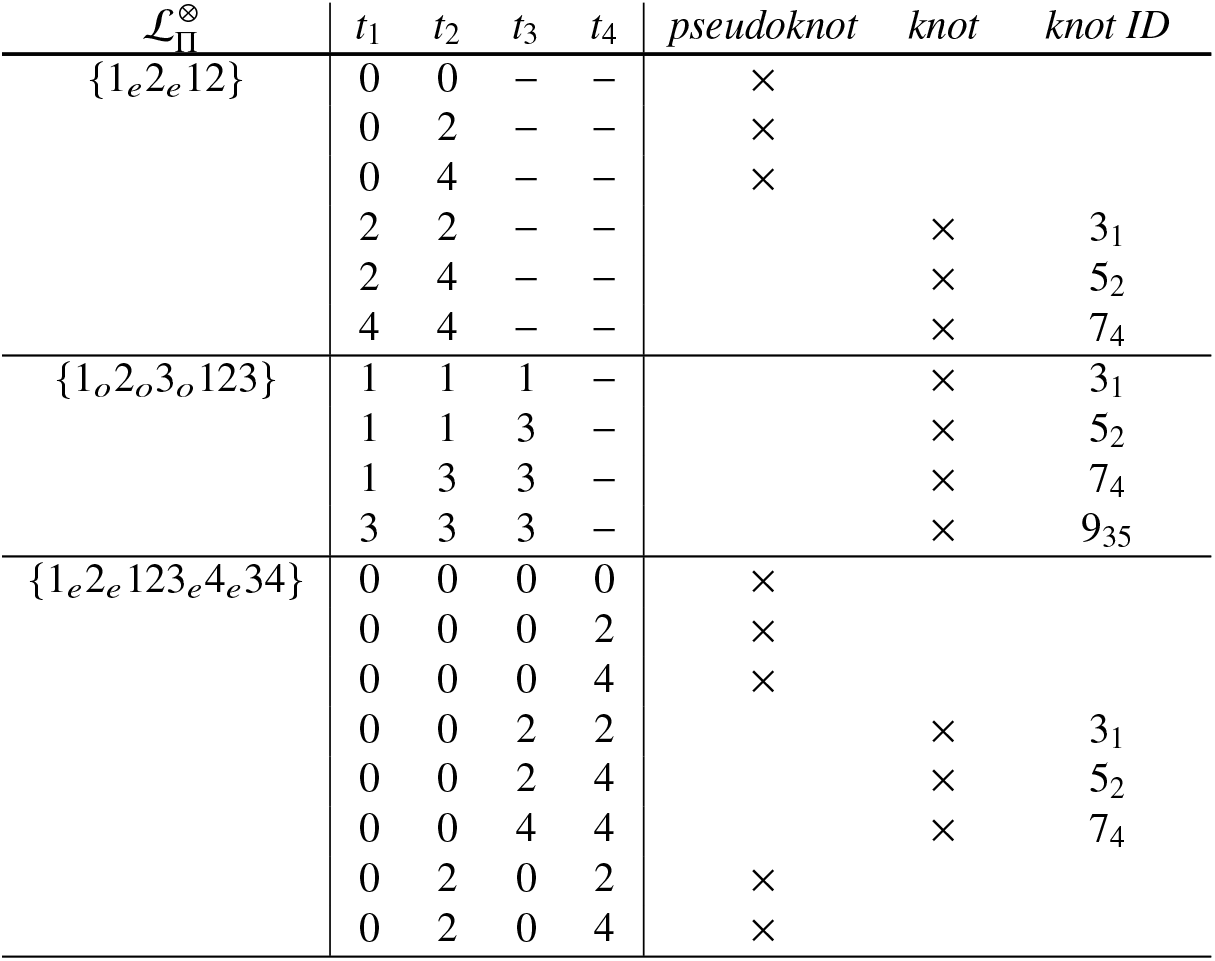
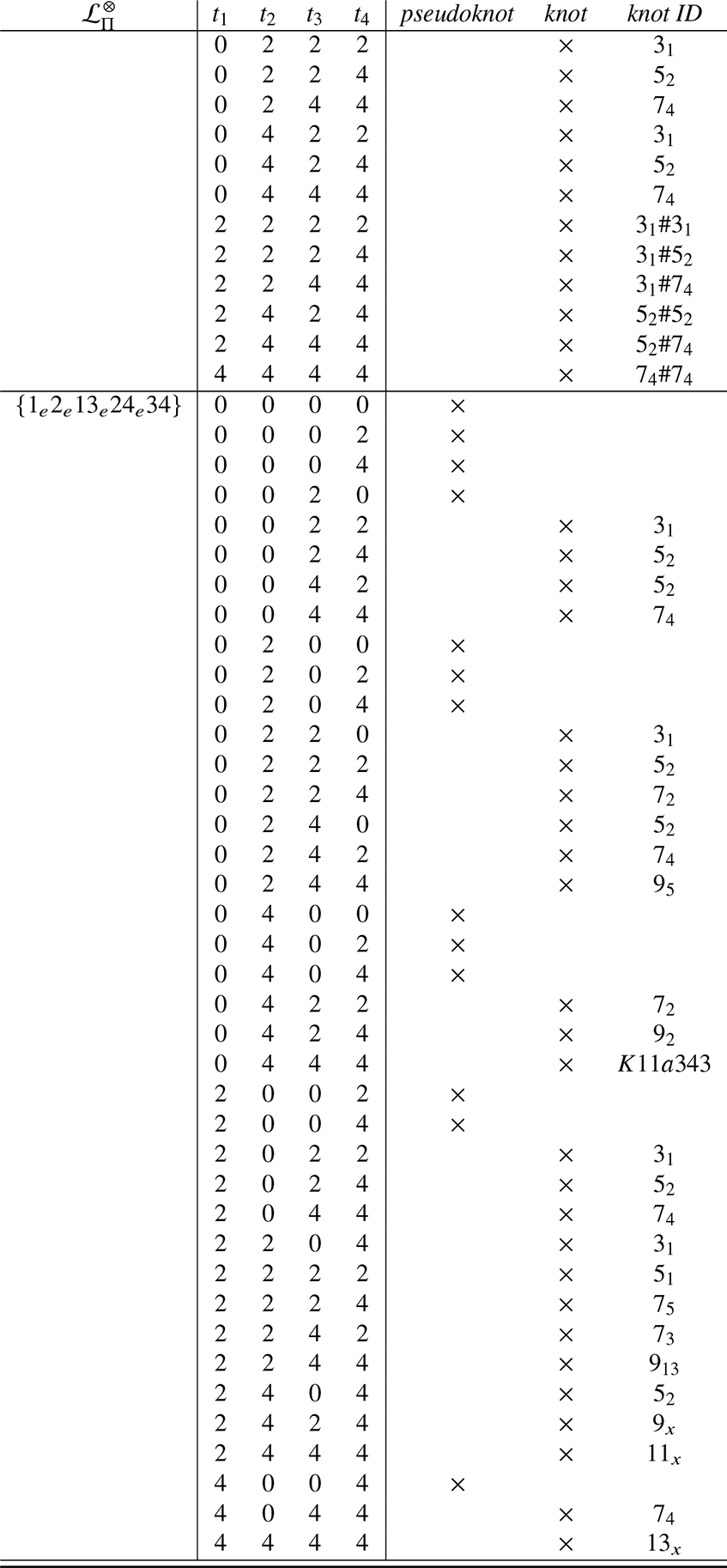
Classification of circular folds whose parities are tuned to allow uncrossed embeddings (*N*_×_ = 0) as pseudoknots, forming the trivial (un)knot 01, or genuine knots. Knots with less than 10 crossings are labelled by their conventional (Alexander-Briggs) symbol, more complex knots by their Hoste-Thistlethwaite label (48). (The knots 9_×_, 11_×_ and 13_×_, labelled by their minimal crossing number *N* as *N*_×_ are unidentified.)

### ssRNA folds

Canonical contracted fold labels, as well as their associated circular diagrams and polarised strand graphs offer a useful framing of the folding of an ssRNA string. Duplexing of the ssRNA string is dictated by the nucleotide sequence. From our perspective, the fold is determined by the relative arrangements of complementary sub-sequences along the string, which sets the bare label 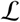 and orientation flags 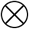, and the number of nucleotides in each sub-sequence, which determines twist *t_i_* and therefore parity flags Π. The tangling associated with that fold topology can be realised by any number of possible embeddings of the fold, which generally include stretches of unpaired single-strands along the string and duplexed double-strands. Here we assume that duplexes are induced by complementary nucleotides (though other interactions which associate or ‘bond’ nucleotide pairs are equally valid, provided those associations induce single- or double-helices only). If every pair of strand fragments of contiguous nucleotides which contributes one component of the double-helix is labelled *i*, where *i* = 1, 2…, *n*, the 2*n* integer string corresponding to the ordering of helical regions from the 5′ to 3′ ends of the ssRNA encodes an unflagged (and likely non-canonical) *n*-ribbon label for the (likely uncontracted) fold, 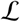. The corresponding semi-flagged fold label 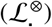 is formed by appending orientation flags for each ribbon (*i*), depending on the relative orientations of the corresponding complementary regions (i.e. *i* or 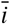 for antiparallel and parallel duplexes respectively). A generic ssRNA fold within our schema includes both parallel and antiparallel duplexes. Though parallel orientation in RNA folds is found *in vitro* or *in vivo*, via, for example, Hoogsteen base-pairing (42), anti-parallel double-helices are more likely. We therefore confine our analysis from hereon to all-antiparallel folds (whose circular diagrams include annular ribbons only). Further, we assume that all double-stranded regions adopt the A-form double-helical geometry, constraining all double-helices to be right-handed (i.e. *t_i_* ≥ 0) with close to 33° twist per nucleotide. In that conformation the number of (half-)twists per duplex in given by 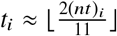, where (*nt*)_*i*_ is the number of nucleotides in each helical strand of the duplex and ⌊⌋ denotes the integer floor operator. (Other estimates of the number of half-twists in each duplex could be made, accounting for different helical structures, non-ideal interactions, physico-chemical environment, etc.) If *t_i_* is an even or odd integer, the associated ribbon parity *π_i_* is *e* or *o* respectively, giving the parity flag Π = *π*_1_*π*_2_… *π_n_*. A fully-flagged label and circular diagram 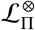 results. The canonical label of the corresponding contracted fold is deduced from 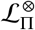 by contracting nested annular ribbons, equivalent to deleting strand bulges separating duplexes, as described above.

Suitable canonical unflagged labels 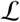 for ssRNA are culled from the compete list of flag-free labels (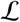, listed in Table *S*7, Supporting Information) by forming all-parallel semi-flagged folds 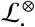 (where all orientation flags are annular) and deleting uncontracted cases, giving a suite of unflagged folds with up to 5 contracted duplexes, summarised in Table 8. All of the resulting contracted folds 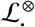 with 2 – 5 ribbons are listed in Table 10. (Folds with 0 – 1 ribbons contract to {0}). Some, though not all, of the folds can be assigned parity flagsΠ = *π*_1_*π*… *π_n_*) to form fully-flagged folds without edge-crossings (*N*_×_ = 0), forming (a)- and (b)-type folds. The remaining cases are (c)-type folds, regardless of their parity flags.

We find 26 contracted semi-flagged all-antiparallel folds, including the trivial fold {0}. Fold labels, circular diagrams and polarised strand graphs of all non-trivial cases are catalogued in Table 10. 9 of those are (c)-type folds, including 3 whose unpolarised strand graph is the nonplanar graph (*K*_5_). The remainder can be flagged to give 20 different contracted (a)- and (b)-type folds, including two (a)-type folds and 18 (b)-type folds.

### From (all-antiparallel) circular to (all-antiparallel) linear fold diagrams

Table 10 implicitly assumes the ssRNA string is closed, giving a circular diagram of the fully-flagged fold 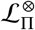, which can be rotated or flipped without changing the fold. In that case, the numbering of duplexes *i* = 1,…, *n* is arbitrary, though its canonical label, assigns labels to each ribbon, modulo symmetries of the diagram. Frequently ssRNA is linear rather than closed, with well defined 5′ and 3′ ends. Cutting the perimeter circle somewhere along its perimeter and straightening the perimeter arc gives the linear diagram. The cut leads to a natural duplex numbering, with duplexes 1 and *n* adjacent to the 5′ and 3′ ends respectively, leading to a linear ribbon diagram (which we denote 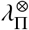). A number of linear fold diagrams can be formed from a single circular diagram, depending on the location of the cut and the string direction (left-to-right, or right-to-left, corresponding to 5′ - 3′ or 3′ - 5′ strings). The cut need not be confined to unduplexed regions of the fold; cutting within a duplex is also possible, splitting the duplex into a pair adjacent to the 5′ and 3′ ends. Since the parities of those terminal duplexes must sum to that of the original contracted duplex, odd-parity (*o*) parent duplexes split to give *oe* or *eo* pairs, and even-parity duplexes (*e*) form *ee* or *oo* pairs. For example, the fold 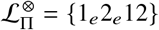 shown in Fig. 44(a) expands to three distinct expanded folds (Fig. 44(b-d)). Like their precursor circular diagrams, linear diagrams can be (un)contracted, depending on the presence or absence of nested annular (or crossed moebius) ribbons.

**Figure 44:**
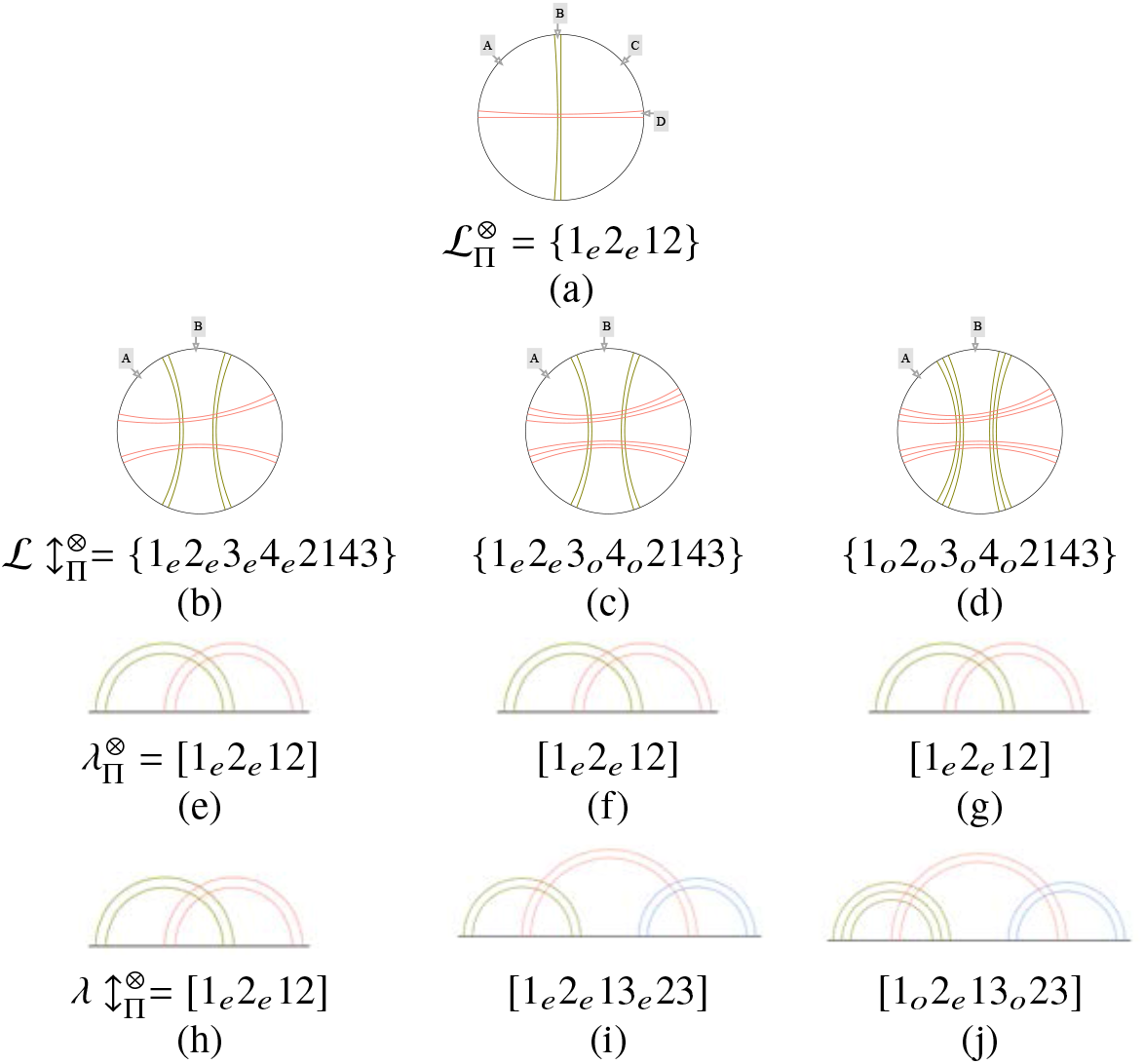
From the fully-flagged contracted fold of (a) circular ssRNA 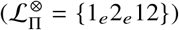 to (e-j) fully-flagged, contracted linear ssRNA folds 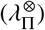. The (even parity) contracted duplexes in (a) are split into two uncontracted ribbons in (b,c,d). Linear folds form by cutting the perimeter and opening the circle into a line. If the diagram is cut at site *A*, the contracted linear diagrams and labels shown in (e,f,g) result. (Even and odd ribbon parities in the linear ribbon diagrams are specified by two or three parallel rulings within a ribbon, as for the circular diagrams.) Alternatively, the circular contracted folds in (a,b,c) can be cut at site *B*, giving contracted linear folds in (h,i,j) respectively. Three distinct linear contracted linear folds result (b,i,j), all sharing the same contracted circular fold (a).

All possible (contracted) linear diagrams from a circular fold 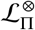 are found by first forming the circular diagram of the expanded fold 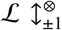 (defined above), cutting that expanded diagram at some site on the perimeter, then contracting the resulting linear diagram.The maximum number of different contracted linear diagrams which can result from a circular diagram with *n* contracted ribbons is 2*n*, since the cut may be on one side of each ribbon or within each ribbon. In practice, that count is usually smaller due to the symmetry of the circular diagram. For example, the circular fold {1_*e*_2_*e*_12}, with 4 cut sites *A, B, C* and *D*, has 2 symmetrically distinct sites, e.g. *A* and *B* (Fig. 44(a)). Cutting at *A* gives the uncontracted linear folds 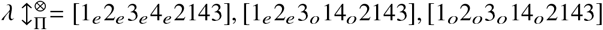, all of which contact to 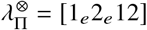 (Fig. 44(e-g)). Cutting at *B* splits one ribbon into a pair of even parity ribbons, giving the linear uncontracted folds 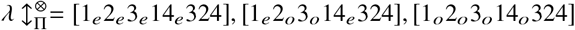, which contract to [1_*e*_2_*e*_12], [1_*e*_2_*e*_ 13_*e*_23], [1_*o*_2_*e*_ 13_*o*_23] respectively (Fig. 44(h-j)). The symmetry of the circular diagram ensures that no new linear folds result from the reversed strings. Therefore the three contracted linear folds 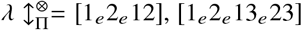 and [1_*o*_2_*e*_13_*o*_23] all share common contracted circular fold {1_*e*_2_*e*_ 12}. Since the parent circular fold is a (b)-type fold, all three linear folds are also uncrossed (b)-type folds (*N*_×_ = 0).

Cutting a contracted circular fold with *n* duplexes induces contracted linear folds with either *n* or *n* + 1 duplexes. The catalog of folds in Table 10 therefore leads to an exhaustive catalog of distinct contracted, uncrossed, all antiparallel linear folds with up to 5 duplexes (tabulated in Table *S*8, Supporting Information), plus an incomplete list of linear folds with 6 duplexes. The fully-flagged uncrossed, contracted, all-antiparallel folds with up to four ribbons in Table 10 result in the extended catalog of uncrossed, contracted, all-antiparallel linear folds listed in Table 11.

Contracted linear folds with up to four duplexes *not* included in Table 11 cannot be embedded in the plane without edge-crossings, i.e. *N*_×_ > 0.

### Pseudoknotted all-antiparallel folds

The suite of linear diagrams in Table 11 are very similar to arc diagrams commonly used to characterise ssRNA folds. There are, however, differences between our circular diagrams and conventional descriptions. First, note that our circular diagrams allow both moebius and annular ribbons, encoding parallel and antiparallel duplexes respectively. That distinction is an essential one if we admit parallel as well as antiparallel ssRNA duplexes, beyond our analysis in this section. Second, ssRNA folds typically include stem loops and bulges, which are ignored in the current analysis, since they do not contribute to contracted folds. Kissing loops are however included, since they may induce strand duplexing. Most known ssRNA folds contract to the trivial unfold, {0}, since their linear diagrams contain uncrossed arcs. Our focus here is on more complex folds, in a topological sense.

Nontrivially folded ssRNA includes all so-called pseudoknotted folds in ssRNA (43), identified in a variety of viruses, ribozymes etc. (44, 45). These folds are characterised by a circular diagram with non-local duplexing interactions (sometimes classified as tertiary interactions (43)) pairing distant strand regions, leading to circular diagrams with crossed ribbons (43,45). That characterisation includes folds which may encode knots or ‘pseudoknots’, assumed to give unknotted tangles (isotopic to the unknot 0_1_). That issue was acknowledged in the pioneering paper of Pleij *et. al*, who recognised that the formation of knots vs. pseudoknots depends on the twists *t_i_* (43). A relatively recent survey of ssRNA folds argued “pseudoknotted structures containing multiple helices of at least ten basepairs would be prime candidates for knotting” (46). Our approach allows clarification of that argument, leading to simpler ‘pseudoknotted’ folds (whose circular diagrams contain crossed ribbons). Further, we derive specific combinations of twists which induce knots rather than pseudoknots for those folds.

Previous analyses have proposed four simplest pseudoknotted folds, known as *H, K, L* and *M* folds (21). H-pseudoknots are relatively prevalent *in vivo*, though *K*, and (to a lesser extent) *L* folds are also known, though *M* folds are currently unknown (23). Those folds correspond to semi-flagged linear contracted folds 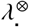, since they are classified by arc diagrams without twists:

- 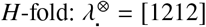
- 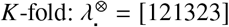
- 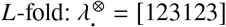
- 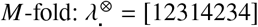

It is interesting - and not totally unexpected - that the relative abundance of known variants of these folds in ssRNA decreases from top to bottom in Table 10. In contrast, the topological indexing of circular forms of these folds developed here is uncorrelated with abundance: closed forms of the *H, K* and *L* pseudoknotted folds wind on (two-colored) genus-3, (two-colored) genus-3 and (two-colored) genus-2 skeletons respectively (*cf*. Table 10). The *K* fold, as yet undetected, is a linear variant of the (rare) *L* fold; both close to give the same one-colored fold on a genus-2 skeleton. Those data suggest that the skeletal genus of the circular fold related to a linear fold is uncorrelated with the frequency of observed pseudo-knotted ssRNA folds, in contrast to an earlier analysis, which found a correlation between occurrence and a distinct measure of fold topology, also denoted ‘genus’. Note that our ‘genus’ refers to the genus of an underlying skeleton (more properly, the genus of the inflated handlebody formed from that skeleton) which can be wound by the RNA strand to give the fold, whereas the ‘genus’ in the classification of (21) refers to the topology of a surface which can accommodate the *circular arc diagram* without intersecting arcs, rather than the fold itself. We do find however, a correlation between the ordering of folds from top to bottom in Table 10 and abundance of known ssRNA folds, although it is difficult to gauge if currently known folds represent a random selection of actual folds *in vivo*. Given the experimental limitations, that data is likely skewed to simpler folds, however such a correlation is a useful working hypothesis. If valid, it implies that there are many other linear folds which are as likely to form as the (currently unknown!) *M* fold: we find 11 distinct 4-duplex linear pseudoknotted folds, including the pair of *M* folds. Our hypothetical ranking of folds therefore differs from that of (21),

### (Pseudo-)Knotting of simpler all-antiparallel fold embeddings

The presence of knots in closed ssRNA strings is not *a priori* determined by the precise fold 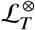, since there is an infinite variety of embeddings of the fold, with different numbers of extra-duplex edge-crossings (*N*_×_), as discussed above. As before, we assume folds adopt their simplest embedding, minimising *N*_×_, and the twist at all edge-crossings *τ_k_* is of unit magnitude (*τ_k_* = ±1). Since (a)- and (b)-type folds listed in Table 10 allow *N*_×_ = 0 embeddings and pseudoknotting or knotting of the folded ssRNA loop is uniquely determined by the twists *t_i_*. The *H, K, L* and *M* pseudoknotted folds can be realised as crossing-free (*N*_×_ = 0) fully-flagged folds for specific tuned parity flags (*cf*. Table 11). Just one parity class of the *H* and *L* folds allows uncrossed ((b)- and (a)-type) embeddings, namely ee and ooo respectively, so that the linear folds *λ* = [1_*e*_2_*e*_ 12] and *λ* = [1_*o*_2_*o*_3_*o*_ 123] induce unique (pseudo)knots (listed in Table 11). The *K* and *M* folds admit two classes with uncrossed embeddings, *eee* and *oeo* (*λ* = [1_*e*_2_*e*_ 13_*e*_23], [1_*o*_2_*e*_‘13_*o*_23]) and *eooo* and *oooe* (*λ* = [1_*e*_2_*o*_3_*o*_ 14_*o*_234], [1_*o*_2_*o*_3_*o*_ 14_*e*_234], listed in Table 11). Those named folds account for 6 of the 15 uncrossed folds of similar complexity in Table 11. We analyse the (pseudo)knotting of all 15 linear folds, via their circular folds which result by joining their 5^0^ and 3^0^ ends. Those have semi-flagged labels 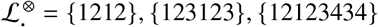 and {12132434}.

The resulting (pseudo)knots for each linear fold [λ], appended with specific duplex twists, *t_i_*, can be found by constructing a planar, crossing-free winding of the ssRNA loop from the polarised strand graphs shown in Table 10. The winding is formed by replacing each polarised vertex in the strand graph by a double-helix with *t_i_* half-twists, whose axis is oriented parallel to the plumbline in the polarised strand graph. The resulting suite of planar embeddings of strand windings is collected in Figs. 12. Some care is needed to ‘read’ these windings, since entry and exit strand around double-helices depend on their twists. To highlight the various duplexed and single-stranded regions, all duplexes are drawn with twists *t_i_* = 5 or *t_i_* = 6, where duplex i has odd or even parity respectively.

Strictly speaking, none of these folds are knotted, due to their loose ends. We assume those ends are joined without adding any edge-crossings (so that *N*_×_ remains 0), in which case the resulting embedding is isotopic (equivalently tangled) to the unknot, 0_1_, or some non-trivial knot, labelled by its conventional knot name. Generally, the duplexed winding differs from the conventional embedding of the knot. In order to identify the induced knot, most windings need to be rewound by some sequence of (Reidemeister) moves which conserve the knotting. (We find the freely-available *KnotPlot* program very useful for these purposes (47).) In fact, all of the (prime) knots can be classified as algebraic links (i.e. numerator closures of generalisations of Conway tangles to rational tangles) (39), listed in Table 13, which simplifies their construction and identification.

If the parity of *t_i_* matches that of the uncrossed fold (*cf*. Table 11), the number of strand crossings associated with each duplex within a fold (*a, b, c*,… in Table 13) is given by *t_i_*, governed solely by the number of nucleotides within the duplex. Alternatively, the twist parity is detuned, in which case the number of strand crossings is equal to *t_i_* + *t*_×(*i*)_ where *t*_×(*i*)_ = ± 1. In the latter cases, different knots emerge from a single fold, depending on the sign of the crossing, *t*_×(*i*)_.

#### Simplest knotting of linear folds with *N*_×_ = 0

We analyse first unique (pseudo)knots formed by uncrossed (i.e. (a) or (b)-type) fully-flagged all-antiparallel folds 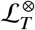, with all twists *t_i_* ≤ 4 tuned to allow crossing-free embeddings of their polarised strand graphs. Those cases are listed in Table 14. The possible symmetries of the oriented strand graphs reduce the number of distinct twists *t_i_* for these folds as follows.

- {1_*e*_ 2_*e*_ 12} folds are symmetric with respect to exchange of ribbon labels 1 ↔ 2
- {1_*o*_2_*o*_3_*o*_ 123} folds are symmetric with respect to cyclic or reverse-cyclic permutation of labels (i.e. 123 ↔ 231 ↔ 312 ↔ 321 etc.)
- {1_*e*_ 2_*e*_ 123_*e*_4_*e*_ 34} folds are symmetric with respect to the permutations 1234 ↔ 2143 ↔ 3412 ↔ 4321
- {1_*e*_ 2_*e*_ 13_*e*_24_*e*_ 34} folds are symmetric with respect to the permutations 1234 ↔ 4321

The data in Tables 11 and 14 determine whether a linear fold 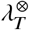 with natural parities allowing an uncrossed embedding is a *bona fide* knot, or a pseudoknot. The linear fold is first translated into the equivalent canonical circular fold formed by closing 5′ and 3′ ends, according to Table 11. Twist parities (*π_i_*) in the circular label 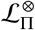 are then replaced by twists *t_i_*, forming the label 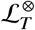, plus its equivalent labels according to allowed ribbon permutations listed above. In practice, choose the permutation which gives the smallest number formed by concatenating all twists *t_i_*. For example, the fold 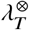 has twists *t*_1_*t*_2_*t*_3_*t*_4_ = 2402. That fold has fully-flagged label [1_*e*_2_*e*_3_*e*_234_e_ 14] (fold 11 in Table 11), whose circular canonical fold is {1_*e*_2_*e*_ 123_*e*_4_*e*_34}. Equivalent twist labels are therefore *t*_1_*t*_2_*t*_3_*t*_4_ = 2402 ↔ 4220 ↔ 0224 ↔ 2042, minimised for *t*_1_, *t*_2_, *t*_3_, *t*_4_ = 0,2,2,4. Table 14 identified the fold {1_*e*_2_*e*_ 123_*e*_4_*e*_34} with *t*_1_, *t*_2_, *t*_3_, *t*_4_ = 0,2,2,4 as knotted, forming the 5_2_ knot.

#### Simplest knotting of linear folds with *N*_×_ > 0

The analysis in the previous section assumes all twist parities are tuned to allow an uncrossed embedding of the characteristic polarised strand graph of the (circular equivalent of the linear) fold. If that is not the case, all embeddings of the strand graph include edge crossings, discussed in some detail above. The simpler examples are the (c)-type folds in Table 10, which contain edge-crossings regardless of their parity flags. In addition, (a) and (b)-type folds with detuned parity flags also contain edge-crossings. Analysis of the strand (pseudo)knotting of folds with edge-crossings follows that developed in the previous section, with the significant addendum that all edge crossings can be embedded in (at least) two distinct ways. The compounds the multiplicity of allowed solutions, leading to lengthy catalogs for folds. We therefore spell out the process for simpler cases only, without exhaustive enumeration.

Consider first folds whose edge-crossings are due to detuned parity flags of related (a) and (b)-type folds. In those cases, a single edge-crossing, with *τ_i_* = ±1, is associated with each detuned parity flag, *π_i_*. That crossing retunes the flag, as shown in Fig. 31(a-c), incrementing or decrementing the net twist from *t_i_* to *t_i_* ± 1. A detuned parity flag allows the fold to either unwind somewhat, resetting its net twist to a lower value (*t_i_* - 1), or wind up (*t_i_* + 1) (assuming all twists *t_i_* are non-negative). Since each detuned parity increments the net twist by ±1 to retune the net twist, (pseudo)knots induced by detuned (a) and (b)-type folds are among the list in Table 14 - they are, however ‘phase-shifted’ up or down by one row in each column of Table 14. If the duplex has *t_i_* = 0 and the crossing is assigned negative twist, 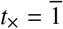, the associated net twist is negative, allowing distinct enantiomers of knots to form compared to those induced by positive net twists.

The following detuned linear *H* and *K* folds (*cf*. folds 1 - 3 in Table 11) 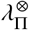, close to give canonical circular folds 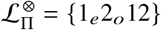:

- [1_*e*_2_*o*_ 12] (*H* fold)
- [1_*o*_2_*e*_ 12] (*H* fold)
- [1_*e*_2_*e*_ 13_*o*_23] (*K* fold)
- [1_*e*_2_*o*_ 13_*e*_23] (*K* fold)
- [1_*o*_2_*e*_ 13_*e*_23] (*K* fold)

Since the twist parity of *π*_2_ = *o* must be retuned to give the crossing-free circular fold {1_*e*_2_*o*_ 12}, the simplest embeddings of these linear folds require a single edge-crossing, *N*_×_ = 1. The resulting (pseudo)knots for *τ*_1_ = ±1 are listed in the upper rows of Table 15.

**Table 15:** (Pseudo)knotting of detuned circular folds with canonical labels {1_*e*_2_*o*_ 12} and {1_*o*_2_*o*_12}, whose simpest embeddings have *N*_×_ = 1 and 2 respectively.

| $\mathcal{L}_\Pi^\otimes$ | $\tau_i$ | $t_1$ | $t_2$ | $t_3$ | $t_4$ | *pseudoknot* | *knot* | *knot ID* |
| --- | --- | --- | --- | --- | --- | --- | --- | --- |
| $\{1_e 2_o 12\}$ | $\overline{1}$ | 0 | 1 | – | – | × | | |
|  |  | 0 | 3 | – | – | × |  |  |
|  |  | 2 | 1 | – | – | × |  |  |
| | | 2 | 3 | – | – | | × | $3_1$ |
|  |  | 4 | 1 | – | – | × |  |  |
| | | 4 | 3 | – | – | | × | $5_2$ |
|  | 1 | 0 | 1 | – | – | × |  |  |
|  |  | 0 | 3 | – | – | × |  |  |
| | | 2 | 1 | – | – | | × | $3_1$ |
| | | 2 | 3 | – | – | | × | $5_2$ |
| | | 4 | 1 | – | – | | × | $5_2$ |
| | | 4 | 3 | – | – | | × | $7_4$ |
| $\{1_o 2_o 12\}$ | $\overline{1}, \overline{1}$ | 1 | 1 | – | – | × | | |
|  |  | 1 | 3 | – | – | × |  |  |
|  |  | 3 | 1 | – | – | × |  |  |
| | | 3 | 3 | – | – | | × | $3_1$ |
| | $\overline{1}, 1$ | 1 | 1 | – | – | × | | |
|  |  | 1 | 3 | – | – | × |  |  |
| | | 3 | 1 | – | – | | × | $3_1$ |
| | | 3 | 3 | – | – | | × | $5_2$ |
| | 1, 1 | 1 | 1 | – | – | | × | $3_1$ |
| | | 1 | 3 | – | – | | × | $5_2$ |
| | | 3 | 1 | – | – | | × | $5_2$ |
| | | 3 | 3 | – | – | | × | $7_4$ |

Likewise, the following linear folds close to give canonical circular folds 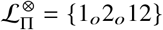, which have *N*_×_ = 2:

- [1_*o*_2_*o*_ 12] (*H* fold)
- [1_*e*_2_*o*_ 13_*o*_23] (*K* fold)
- [1_*o*_2_*o*_ 13_*e*_23] (*K* fold)

The resulting (pseudo)knots for 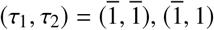 and (1,1) are listed in the lower rows of Table 15.

Detuned linear *L* and *M* folds (*cf*. folds 4,8,9 in Table 11) 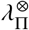, close to give canonical circular folds 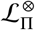 with a single edge-crossing (*τ*_1_) required to retune the parity of ribbon *π*_1_:

- [1_*e*_2_*o*_3_*o*_123] (*L* fold)
- [1_*o*_2_*e*_3_*o*_123] (*L* fold)
- [1_*o*_2_*o*_3_*e*_123] (*L* fold)
- [1_*e*_2_*o*_3_*o*_14_*e*_234] (*M* fold)
- [1_*o*_2_*o*_3_*o*_14_*o*_234] (*M* fold)
- [1_*e*_2_*e*_3_*o*_14_*o*_234] (*M* fold)
- [1_*e*_2_*o*_3_*e*_14_*o*_234] (*M* fold)
- [1_*o*_2_*e*_3_*o*_14_*e*_234] (*M* fold)
- [1_*o*_2_*o*_3_*e*_14_*e*_234] (*M* fold)

The resulting tangles for *τ*_1_ = ±1 are listed in the upper rows of Table 16.

**Table 16:**
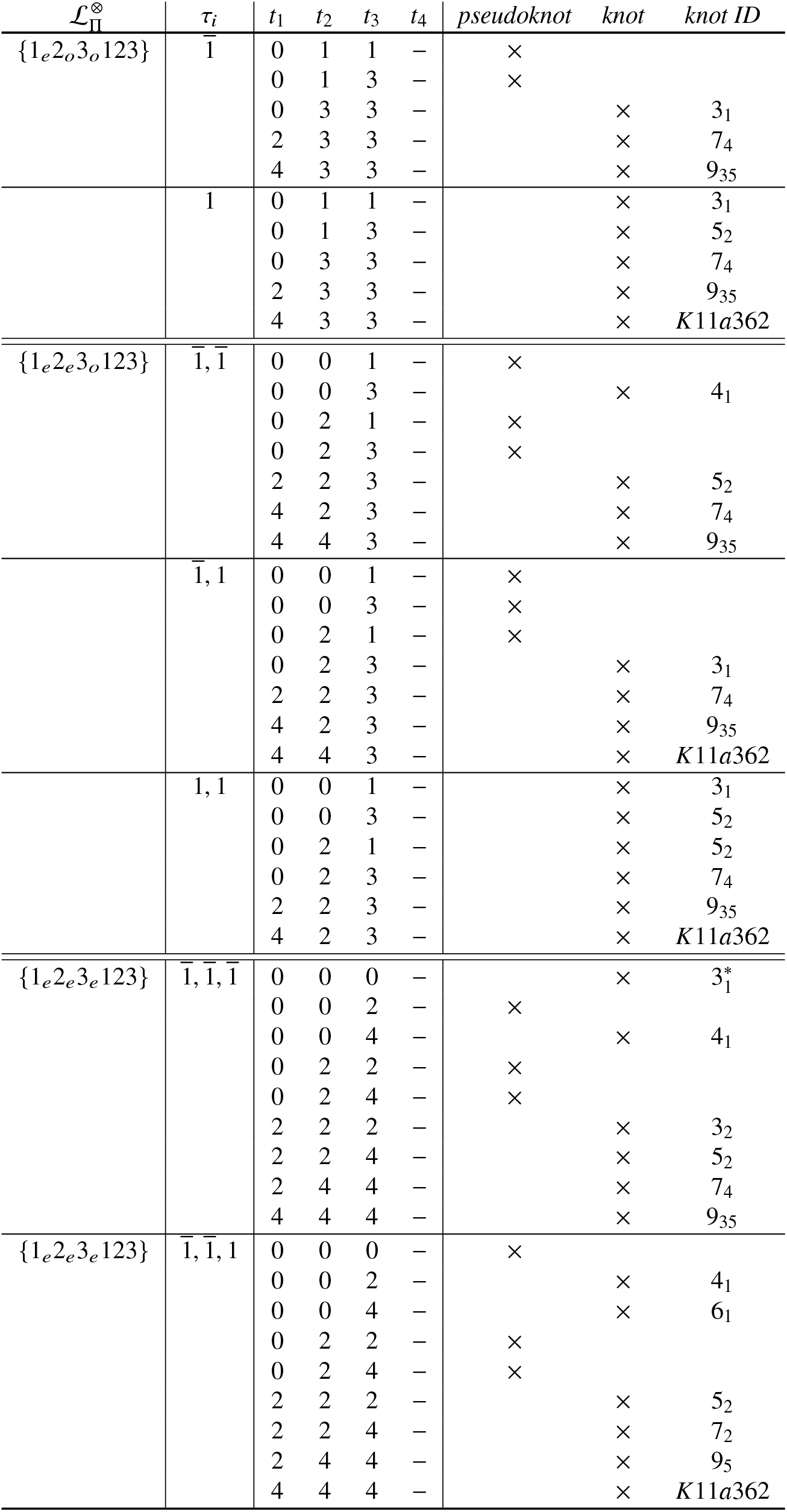
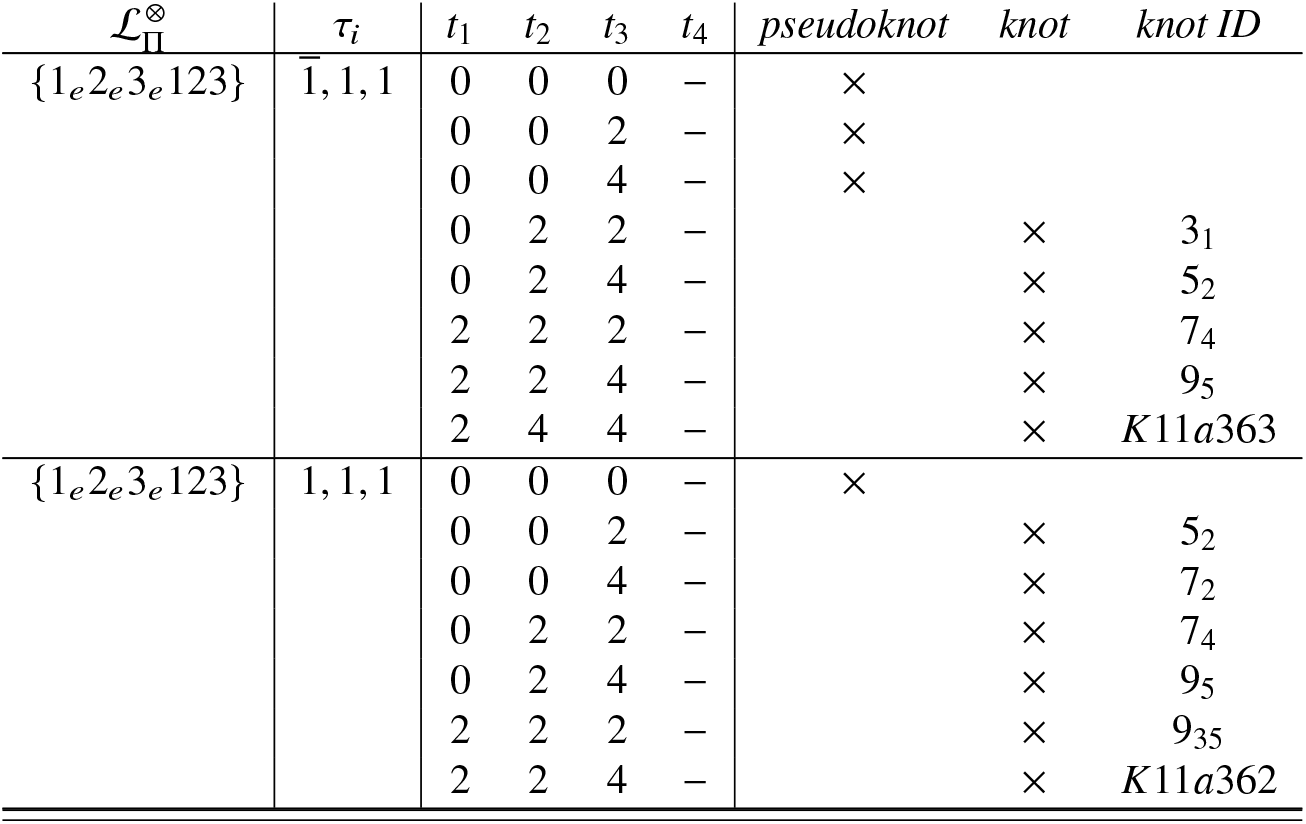
(Pseudo)knotting of detuned circular folds with canonical labels {1_*e*_2_*o*_3_*o*_ 123}, {1_*e*_2_*e*_3_*o*_ 123} and {1_*e*_2_*e*_3_*e*_123}, whose simpest embeddings have *N*_×_ = 1, 2 and 3 respectively. (The trefoils 3_1_ and 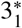 are enantiomers, *cf*. Fig. 43.)

The same analysis, though tedious, is readily done for detuned cases of the remaining linear folds in Table 11, which close to form (detuned) circular folds with labels 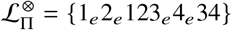 and {1_*e*_2_*e*_13_*e*_24_*e*_34}. Since those folds have four contracted duplexes, *N*_×_ = 1, 2, 3 or 4 edge-crossings for 1 – 4 detuned parities respectively are required for the simplest embeddings of these folds. Each of those crossings can have twist *τ_i_*, = ±1, allowing considerable unwinding or winding compared with their relaxed embeddings in cases where all four ribbons are detuned and edge-crossings have equal twists 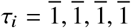 and *τ_i_* = 1,1,1,1 respectively.

Finally, consider embeddings of (c)-type folds, which contain edge-crossings, regardless of the parity flags. Two distinct phenomena are possible. In one scenario, (c)-type folds have topologically planar unpolarised strand graphs, but crossing-free embeddings of the polarised graph include moebius flags, leading to folds with parallel duplexes. That class includes the semiflagged folds in Table 10 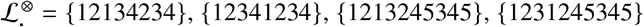 and {1231452345}. Whereas crossed (a) and (b)-type folds considered in the previous section have detuned parity flags, these (c)-type folds have detuned orientation flags, regardless of their parity flags. For example, the bare fold label 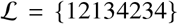 allows four different fully-flagged folds with *N*_×_ = 0, shown in Fig. 45, all with both parallel and antiparallel double-helices.

**Figure 45:**
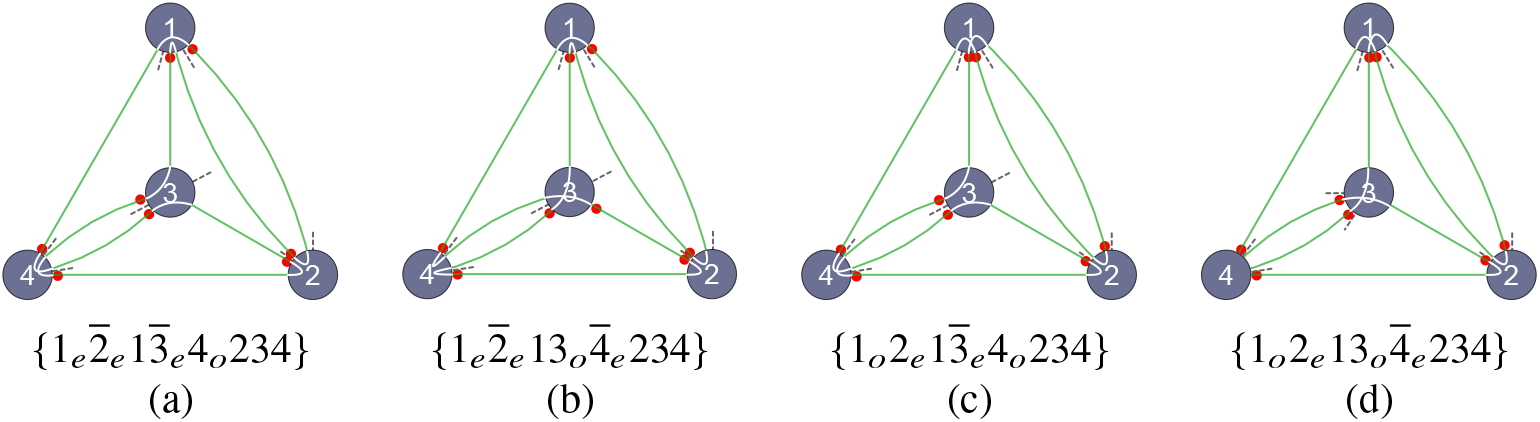
All fully-flagged folds with bare label 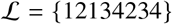. None are all-antiparallel.

Recall that any detuned orientation flag ⊗_*i*_, inducing a parallel duplex, can be retuned to induce an antiparallel duplex by adding a single edge-crossing to the polarised strand graph, as shown in Fig. 31(d-f). Each mistuned orientation flag contributes a single edge-crossing to a minimally crossed embedding of the fold. Thus, for example, the folds in Figs. 45(a,b) can be retuned to give all-antiparallel folds provided *N*_×_ = 2. Those retuned folds have labels {1_*e*_2_*e*_ 13_*e*_4_*o*_234} and {1_*e*_2_*e*_ 13_*o*_4_*e*_234} respectively. Similarly, the folds in Figs. 45(c,d) can be retuned to the folds {1_*o*_2_*e*_ 13_*e*_4_*o*_234} and {1_*o*_2_*e*_ 13_*o*_4_*e*_234} with the insertion of a single edge-crossing, *N*_×_ = 1. As for the crossed folds with detuned parity flags, the simplest embeddings of the crossed folds with detuned orientation flags have twists *t*_×(*i*)_ = ±1 associated with each detuned flag, leading to multiple possible (pseudo)knots. The simpler examples of these fold embeddings are also rational tangles, so their (pseudo)knots too can be constructed numerically via those tangle descriptions (e.g. analogous to the commands in *KnotPlot* listed in Table 13).

A second scenario plays out when the strand graph is topologically nonplanar, regardless of orientation of parity flags. For example, the bare folds 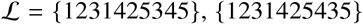 and {1231435425} (Table 10) induce the nonplanar graph *K*_5_, introduced above. Since *K*_5_ can be embedded with just one crossing (and no less), the bare fold embedding has *N*_×_ = 1. Appending parity and orientation flags to those bare folds adds a further number of crossings, equal to the number of detuned flags. As before, strand (pseudo)knotting can be deduced by assigning ±1 twists to all crossings, including the single crossing characteristic of the bare fold.

### Alternating knots by design

The paper is primarily motivated by the characterisation of fold topology resulting from a sequence of nucleotides along a ssRNA chain. We close with a brief comment on the converse problem, namely the design of (possibly artificial) nucleotide sequences to induce folding into some target topology, underlying DNA and RNA nanotechnology. In particular, the preceding analyses suggest particularly robust topological designs. Such preferred designs include antiparallel duplexes only, since nucleotide sequence design for parallel duplexes is more delicate. Further, right-handed helices are more readily realised by ssRNA than left-handed versions. Among the three broad classes of all-antiparallel folds, (a)-type folds are particularly attractive targets since they can be designed to build assemblies which are strongly enthalpically favoured due to the possibility of duplexing virtually all nucleotides, except those confined to the (conventional) junctions. We therefore target knots which can be realised as all-antiparallel, homochiral (*t_i_* ≥ 0) (a)-type (one-colored) folds, since they are likely to have few competing folds of comparable stability, constraining strand geometries and preventing uncontrolled strand crossings.

Consider, for example, nucleotide sequences likely to induce folding of ssRNA into strictly alternating knots, with minimal crossing numbers *N_min_* between 0 – 10 crossings. Further, target the simplest embeddings of those knots, which realise those minimal crossing numbers, so that the total number of crossings in the winding, 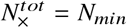. Other embeddings of the same knot are possible with 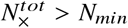. However, if 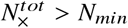 all embeddings differ only by ‘flypes’ (29). That considerably simplifies the analysis, since flyping leaves both the parallel/antiparallel and homo/heterochiral characteristics of a winding unchanged. That allows us to analyse just a single one-colored embedding of each alternating knot (any one will do, provided with 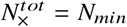).

A circular fold diagram for any knot embedding is easily constructed by reversing the construction of a fold coloring introduced earlier. First color faces of the shadow embedding onto a plane alternately black and white around each vertex, forming a checkered drawing. Next, align all white and black plumblines through graph vertices such that they pass between adjoining white and black domains respectively. A pair of one-colored fold skeletons are then formed by joining all black or white skeletons through black or white domains. Number each crossing of the knot by distinct integers, *i* = 1,…, *N_min_*, and replace crossings by a strand graph vertex *i* - with appropriate plumbline polarisation - and twists *t_i_* = ± 1, depending on the sign of the unit twist (*cf*. Fig. 38). A continuous path along the knot defines the label 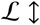 of the expanded polarised strand graph of the knot. The orientation flags of the (all-white) fold or its (all-black) antifold are determined by marking the directed the cycle 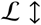, as in Figs. 28 and 29. Since all crossings of the knot give graph vertices (so that *N*_×_ = 0) and the twists are measured with respect to monochromatic (all-white or all-black) vertices, 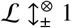 describes an (a)-type (relaxed) fold.

Generic knots lead to a pair of circular diagrams of the fold-antifold pair which are heterochiral, with both positive and negative helical twists, *t_i_*), and mixed orientation flags, combining parallel and antiparallel duplexes. However, in some cases, the relaxed fold-antifold pair is homochiral, one of which is all-parallel and the other all-antiparallel. All-antiparallel, right-handed double-helical folds with *N*_×_ crossings in the expanded fold are therefore possible for selected alternating knots. (If the knot is topologically chiral just one enantiomer is suitable.) Among the 196 alternating knots with *N*_×_ ≤ 10, 34 can be realised as uncrossed, homochiral all-antiparallel folds, listed in Table 17. Each of those may be realised by duplexed folds of suitably encoded ssRNA strings.

**Table 17:** Alternating knots described by relaxed, homochiral, all-antiparallel duplexed folds. * The 10-crossing knots were analysed by Neave Taylor.

| *crossing #* | *knots* | *# (total #)* |
| --- | --- | --- |
| 0 | $0_1$ | 1 (1) |
| 1 | — | — |
| 2 | — | — |
| 3 | $3_1$ | 1 (1) |
| 4 | — | 0 (1) |
| 5 | $5_1, 5_2$ | 2 (2) |
| 6 | — | 0 (3) |
| 7 | $7_1, \dots, 7_7$ | 7 (7) |
| 8 | — | 0 (18) |
| 9 | $9_1, \dots, 9_7, 9_9, 9_{10}, 9_{16}, 9_{18},$<br>$9_{23}, 9_{28}, 9_{35}, 9_{38}, 9_{39}$ | 16 (42) |
| 10* | $10_{49}, 10_{53}, 10_{55}, 10_{63}, 10_{66},$<br>$10_{80}, 10_{101}, 10_{120}$ | 8 (123) |

Fully-flagged, contracted fold labels can be derived for these all-antiparallel *N*_×_-crossing knots, defining the combinatorics of sequences and complementary sequences along the ssRNA strand. For example, the conventional embedding of the alternating knot 10_120_ illustrated in Fig 46(a) induces a homochiral, all-antiparallel fold, so an ssRNA sequence can be designed to form that knot by the following steps. First the alternating knot embedding (Fig. 46(a)) is directed, and marked as shown in Fig. 46(b). Plumblines are positioned to enforce antiparallel duplexes (Fig. 46(c)), setting the twist in each crossing to *t_i_* = 1 or 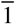. This alternating embedding gives a fold whose twists are all equal to 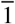 and whose plumblines are all monochromatic. The resulting fold is a therefore a homochiral all-antiparallel (a)-type fold, with four conventional Y-junctions and a single conventional X-junction, illustrated in Fig. 46(c). However, this embedding of the 10_120_ knot gives (disfavoured) negative twists. The simpler design target is therefore its enantiomer 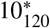. The polarised strand graph embedding of the expanded fold characterising 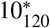, 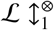 is shown in (Fig. 46(d)), which contracts to the all-parallel, homochiral relaxed fold circular fold shown in Fig. 46(e), whose circular diagram is given in Fig. 46(f). Linear folds are formed by cutting the circular fold at various sites; the symmetry of the circular diagram limits the number of distinct linear folds. Ten distinct linear folds result, including six labelled in (Fig. 46(d-i)), and four additional folds formed by reversing the folds in (e-h).

**Figure 46:**
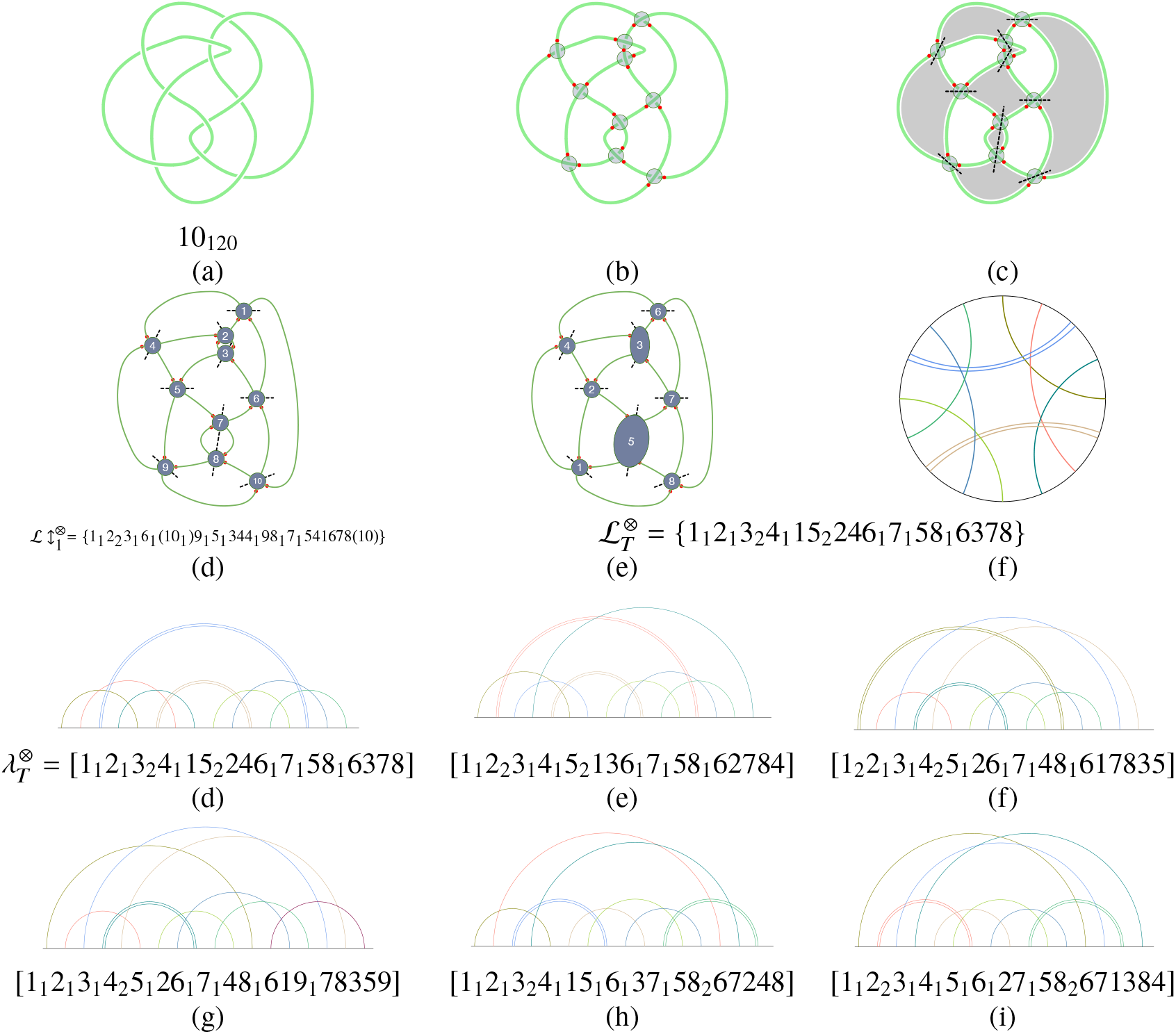
Construction of linear ssRNA sequences designed to wind into the 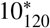 alternating knot embedding in (a). (This is the enantiomer of the conventional drawing of 10_120_.) The fold label is formed from the one-colored oriented strand graph, constructed in (b-c). (c) Arbitrary labelling of the expanded fold 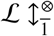 with crossings 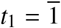 at all graph vertices, in order to induce 10 antiparallel duplexes. (d,e) The contracted fold with canonical label 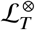 containing 8 duplexes, with *t*_3_ = *t*_5_ = 2 due to contraction of pairs of crossings. (d-i) Six distinct linear folds 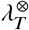 formed by opening the circular fold at symmetrically distinct sites on the perimeter. Four additional linear folds exist, formed by reversing the folds in (e),(f),(g) and (h), swapping 5′ ↔ 3′ ends. (Reversing folds (d) and (i) give equivalent folds due to the symmetry of their linear fold diagrams.)

Those linear fold diagrams describe the ordering of self-complementary domains distributed along the length of the ssRNA strand from 5′ to 3′ ends. The designs in folds (d,e,f,h,i) (and the reversed folds from (e) and (f)) include eight self-complementary domains (numbers 1 - 8 in the labels 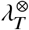), while fold (g) (and its reverse) includes nine. The number of nucleotides within each domain is between 1 – 5 for domains with *t* = 1 and 6 – 11 for domains with twists *t_i_* = 2, so that *ca*. 20 - 90 nucleotides are required to form the requisite duplexes, separated by single-stranded regions. The number of unpaired nucleotides within those ss regions is not fixed by the fold diagram. The circular diagram is (bilaterally) symmetric if all single-stranded regions between duplexes are of equal length. Judicious design of the ssRNA string may allow embeddings of 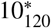 which exhibit similar symmetry. Those geometric details - though interesting - are beyond the scope of this paper; we note only that symmetric forms of circular diagrams lead at once to symmetric fold embeddings.

## DISCUSSION

This paper has been inspired in part by an analysis of known ssRNA folds, which suggested that knotting is rare at best, and possibly non-existent *in vivo* (11). Those authors closed their paper by urging the development of structure-prediction tools to distinguish ssRNA knots from pseudoknots. While the question of knotting of ssRNA *in vivo* remains open, the constructions in this paper suggest an explicit path towards that goal. Those constructions require substantial groundwork, which, though lengthy, is in our view useful since they allow comprehensive enumeration of folds to limited complexities. In particular, we have enumerated all possible one-colored duplexed (semi-flagged) circular folds (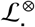) containing up to five contracted duplexes. A subset of those folds include all relaxed folds wound on skeletons of genus *g* ≤ 3. An overlapping set of contracted, all-antiparallel and uncrossed circular folds - either one- or two-colored - with up to five contracted duplexes has also been catalogued.

Analysis of duplexed knots which may result by folding requires the development of oriented strand graphs, with a number of explicit constructions of those graphs from their associated circular diagrams. Oriented strand graphs (or, equivalently, their associated circular diagrams or canonical fold labels) are the central concept of this analysis, allowing us to rigorously sort folds into equivalence classes. That construction rests on the notion that a ‘fold’ is a topological rather than geometric property, since oriented strand graphs can have an infinite variety of geometrically distinct embeddings. Topological classification allows for explicit sorting of folds into manageable class sizes. Further, it allows us to link a fold with the concept of a knotted fold. That too requires some groundwork, via the new concepts of an antifold and isomorphic folds. Those additions are useful, in that they expose important differences between a fold and a knot.

In our view, the question of knotting of ssRNA *in vivo* remains open. However, one definitive conclusion can be made. If ssRNA folds are duplexed according to the most likely A-helical geometry, fold knotting is readily achieved. Table 14 reveals that the {123123} fold is necessarily knotted provided all three double-helices have 6 - 10 nucleotides per strand (*t*_1_ = *t*_2_ = *t*_3_ = 1) *in the absence of extra-duplex edge crossings* (*N*× = 0). Similarly, the simplest four-duplex folds, {12123434} and {12132434} are knotted as long as two of their duplexes have 1 - 5 nucleotides in each strand (*t*_1_ = *t*_2_ = 0) and the remaining duplexes have 12 - 15 nucleotides (*t*_3_ = *t*_4_ = 2,2), provided *N*× = 0. On the other hand, Table 15 reveals that the simple two-duplex fold {1212} is also knotted, provided both duplexes have 6 - 10 nucleotides per strand (*t*1 = *t*2 = 1) and the additional edge-crossings have the same twist (s_1_ = s_2_1). Clearly, strand knotting is possible without highly wound double-helices. However, those configurations rely on specific configurations of single-stranded regions in the ssRNA chain, dictating the number of single-strand crossings *N*_×_ and their type *τ_k_*. Certain secondary or tertiary structural features will surely favour specific values for *N*_×_ and *τ_k_* over others. The knot listings above have been deduced on the basis of minimal edge-crossings only. That constraint, though fit-for-purpose here, is artificial. It does however afford a useful middle ground. For if it does not hold, additional edge-crossings are present. Those additional crossings necessarily increment *N*_×_ by an even number, since the addition of an odd number will detune some orientation or parity flags, resulting in a different fold. If those new crossings are randomly distributed between *t*_×_ = 1 and 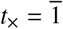, the knot tables above remain valid. Alternatively, if they are unbalanced, and favour positive rather than negative crossings, a net increase in twists results, further enhancing the likelihood of a knotted fold. Conversely, an excess of edge-crossings with negative twists will reduce the knotting provided the twists associated with strand duplexing *t*_i_ are large compared with the added negative twists. If *t*_i_ is small however, those edge crossings can rewind the fold to form a knot enantiomer of the opposite hand. (A simple example is the {123123} fold, where *t*_1_ = *t*_2_ = *t*_3_ = 0, containing three edge-crossings, 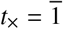, in Table 16). Given these complications, definitive conclusions are impossible to draw in the absence of specific data for *N*_×_ and *t*_×_. That finding is useful. While the magnitude of crossings in a duplex (*t*_i_) is driven largely by secondary interactions, the presence of a pseudoknot or a genuine knot is critically dependent on the sign of the *N*_×_ unduplexed edge-crossings, which - in the absence of tertiary interactions - is statistical, leading to a distribution of knots and pseudoknots for the same ssRNA. On the other hand, tertiary interactions are likely to steer those crossings, favouring one form over the other. Presumably, those interactions are finely tuned *in vivo* to favour pseudoknots over knots. If they can be modified to switch edge-crossings, viral efficiency is likely to be compromised.

These ideas can be applied for example to an ssRNA fold of extreme current relevance: the pseudoknotted region of the RNA genome of SARS-Cov-2, the virus responsible for the current COVID pandemic. The relevant tangled domain lies between nucleotides *nt*(13418) and *nt*(13488) (between the ORF1a and ORF1b regions (49)), buried within the very large 30 000 ssRNA genome. Disruption of this pseudoknotted fold is a potential target for inactivation of the virus (19), affecting the multiple proteins encoded by this programmed-1 ribosomal frameshifting region (16). Modelling has led to the likely secondary structure of this psedoknotted fold which contains three duplexed domains, published in (18) (Figure 1.) That can be recast into the linear diagram shown in Fig. 47(a) where the *x*-axis indicates the nt index in the genome (−13418, the nt index of the first nt in this region). The contracted diagram (which ignores duplex S3) has semi-flagged linear fold label 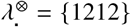, where ribbons 1 and 2 correspond to the stem regions denoted *S*_1_ and *S*_2_ in (18). The fold therefore corresponds to the H-pseudoknot, indexed as fold 1 in Table 11 and can be abstracted to the simplified embedding shown in Table 12. (In practice, the fold embedding aligns both helices, embedded orthogonal to each other in our Figure.) If we idealise those stem regions to A-form double-helices, the number of nucleotides in ribbons 1 and 2, ca. 10 and 6 respectively, translate to *t*_1_ ≈ 1.8 and *t*_2_ ≈ 1.2, giving a fully-flagged fold 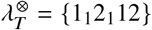 corresponding to 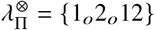. Both twists are therefore detuned, so that *N*_×_ = 2. The relevant string tangling can be read from the lower half of Table 15, since 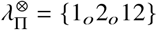. There are three possibilities: 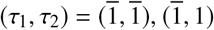 or (1,1), corresponding to 0_1_, 0_1_ and 3_1_ respectively.

**Figure 47:**
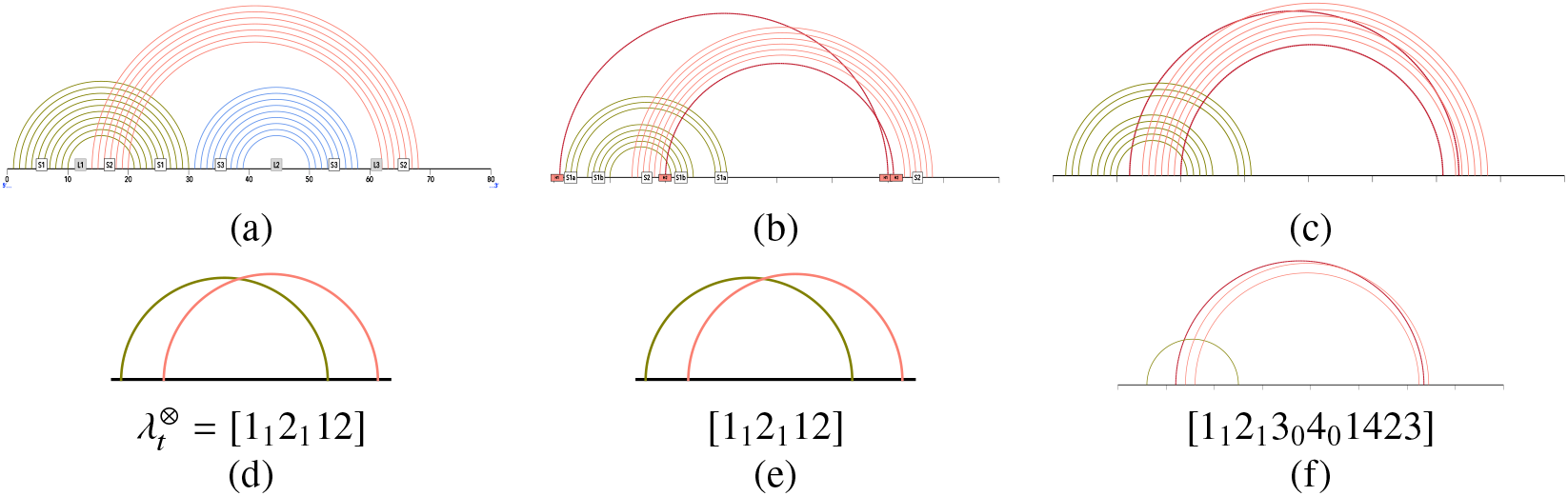
(a) Linear fold diagram of the pseudoknotted frameshifting stimulatory region in the RNA genome of SARS-Cov-2. Base-pairing of individual nucleotides is indicated by arcs, colored according to the relevant duplex (or stem region) labelled *S*1, *S*2 and *S*3 following the labelling in (18). The fold contracts to the 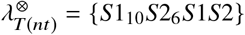, where *T*(*nt*) is the number of nt within the *S*1 and *S*2 annular ribbons, ignoring bulges etc. (b) Simplified fold diagram of a possible ‘L3 threaded’ structure given in (18) (omitting the *S*3 loop for clarity), including two tertiary H-bonded interactions (indicated by blood-red arcs). (c) Simplified fold diagram of a possible ‘5’ threaded’ structure given in (18) (omitting the *S*3 loop for clarity), including two tertiary H-bonded interactions (indicated by blood-red arcs). (d-f) Contracted fold diagrams of the folds in (a-c) respectively, with linear fold labels.

If the fold is driven by (idealised *A*-form) secondary structure alone, all three combinations of (*τ*_1_, *τ*_2_) are equally likely. In other words, there is a substantial probability (33%) of (trefoil) knotting within this region. Given the virulence of this virus, that is likely to be a substantial over-estimate, assuming viral reproduction requires unknotted RNA. In order to suppress knotting, specific interactions, disrupting the ideal *A*-helical geometry (thereby retuning the effective twists *t*_i_) may be active. Those include helical distortions due to specific nucleotides within the duplexes and tertiary interactions. Since tertiary interactions between *nt*(*i*) and *nt*(*j*) in the string effectively glue those nucleotides, the interaction can be modelled by a semi-flagged arc spanning locations *i* and *j*, with zero effective twist, *t*_i_ = 0 (i.e. *π*_i_ = *e*). (The ribbon is semi-flagged since it has no preferred - annular or moebius - orientation). For example, simulations by Woodside *et. al* (18) suggest a number of tertiary interactions in this pseudoknotted region, including H-bonding between *H*1 = *nt*(1) - *nt*(60) and *H*2 = *nt*(20 - *nt*(61) (their ‘L3 threading’ model), giving the modified (uncontracted) linear fold diagram in Fig. 47(b). Since that diagram contracts to the same diagram in (a), those interactions do not alter the fold topology and are unlikely to ensure unknotting.

Alternative structures are also proposed in (18), including ‘5’ threading’ models’, with slightly different tertiary (H-bonding) interactions. In those cases, one of those tertiary interactions interrupts the *S*2 duplex, effectively splitting it into a pair of duplexes and resulting in a new fold topology. A simplified linear diagram of the fold is shown in Fig. 47(c), which indeed contracts to a new linear fold topology, with label 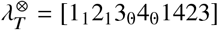. Closure of the 5’ and 3’ ends gives a circular fold diagram with contracted label 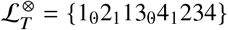, with semi-flagged label, 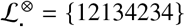. Table 10 indicates that this semi-flagged fold is a (c)-type fold, regardless of twist parities. The construction algorithm for a fold with arbitrary fold label, described earlier in the paper, leads to a simplest embedding of the fold 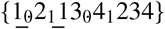 with *N*_×_ = 3. That embedding can be formed, for example, by first building the fully-flagged fold 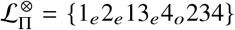 (for which *N*_×_ = 0), then switching the parity flag in ribbon 2 (π_2_ → *o*) and orientation flags in ribbons 2 and 3 (⊗_2_, ⊗_3_ → annular) with a single edge-crossing resulting from each switch. Assuming the three edge-crossings have equal probability of either crossing-type (i.e. *τ*_k_ = 1 or 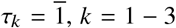), the knotting statistics are different to those derived above for idealised structures induced by secondary duplexing only, or those involving the ‘L3 threading’ model. Those structures allow a 33% chance of trefoil knotting, whereas the new topology of the 5’-threading model structures reduce that knotting probability to 25%. Although the analysis is far from conclusive at this stage, the point is clear: fold knotting is dependent on topology, and tertiary interactions can reset knotting probabilities of folds. From a theoretical perspective, that conclusion is interesting and (to us) surprising, since we have assumed those tertiary interactions simply pull disparate strand regions together, and leave all helical twists unchanged. That finding has clear practical relevance, since it demonstrates that viral RNA can be fundamentally edited by switching on or off specific tertiary interactions.

Lastly, we point out that the analysis here has been developed assuming the simplest scenario, where folds are induced by Watson-Crick duplexing. The approach is more generally applicable, allowing a quantitative picture of entanglement even in the absence of double-helices. For example, generic attractive interactions between distant regions of an extended polymeric molecule can be considered to induce an untwisted double-helix, so that *t*_i_ = 0, and the analysis above remains valid, since that configuration is a special case of an even-parity helix. Further, extension of the analysis to include triple-helices, etc. is conceptually straightforward. Circular diagrams for triple-helical folds include triangular (etc.) ribbon domains, whose three edges characterise the three possible pairings of strands, each with characteristic orientation flags.

## ACKNOWLEDGMENTS

I am very grateful to Neave Taylor (Sydney) whose data allowed me to complete Tables 3 and 17, Damian Lin (Sydney) for discussions related to Dehn’s construction, Casey Duckering (Chicago) for his open source “Hyperbolic” software (50) and Rob Scharein (Vancouver) for assistance with his “KnotPlot” software (47). I also thank Robert K. Thomas (Oxford) and Myfanwy Evans (Potsdam) for encouragement.

## SUPPORTING INFORMATION

### NAMING AND SORTING FOLDS

Circular fold diagrams can be labelled by a sequence of positive and negative digits, encoding the sequence of ribbons meeting the perimeter along a cyclic traverse of the perimeter from an arbitrary origin. Each ribbon, *i*, intersects the perimeter at two pairs of footings and has even or odd parity, set by its associated twist, *t*_i_. The first encounter with ribbon i is tagged with the parity of that ribbon, denoted *π*_i_ giving a ribbon label, {i_o_} or {i_e_}. If the ribbon is moebius, that first entry is further flagged by a negative sign, ī. For example, the simplest nontrivial fold has label 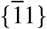.

A nested sequence of annular ribbons is labelled {123 … 321. The nested annular ribbons can be merged to a single reduced ribbon by sliding their footings together, providing the gaps between adjacent ribbon footings are free with no other intercalated ribbons. The merging process gives the relation between fold diagrams:

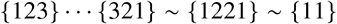

so that adjacent repeated (positive) entries can be deleted. The twist associated with the resulting merged annular ribbons is 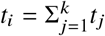.)

Similarly, a fan of crossed moebius ribbons can merged by closing the fan into a single moebius ribbon, provided there are no other ribbons in the interstices between footings:

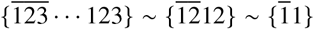

Reduced folds merge all nested annular ribbons and all fanned moebius ribbons. Conversely, any reduced label can be expanded via intercalation of nested or star-shaped ribbons.

The reduced labels are not unique, since a circular walk around the perimeter may head off from any location on the perimeter, and proceed clockwise or anticlockwise, so a cyclic or reversed cyclic permutation of the label, renumbering each ribbon so that the first encounter is 1, gives the same fold.

1. from some origin, walk clockwise or anticlockwise around the perimeter once, and label each new ribbon terminal by successive digits, 1, 2,…, so that every digit is listed twice: one entry for each terminal. All closed tours around around the perimeter are therefore encoded by a label made of a 2*k*-digit string. The labels (may) depend on the starting terminal and the direction of the cyclic tour.
2. append an orientation flag to each ribbon in the label, identifying it as annular or moebius. If the ribbon *i* is annular, it is labelled *i* with associated twist index ⊗_*i*_ = 0; if it is moebius, it is labelled ī with associated orientation index *O*_i_ = 1
3. append a parity flag to each ribbon in the label, identifying it as odd or even, i.e. if a ribbon *i* is even, it is labelled *i*_e_ and *i_o_* respectively with associated parity indices *P_i_* = 0 and *P_i_* = 1 respectively.
4. any ‘bare’ label is a string of positive digits, stripped of its orientation flag, leaving an integer. The canonical bare label, denoted 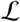, is the smallest integer. This means that the walk sets out from a ribbon with smallest span (and bare label 1), and proceeds in the direction that arrives at the other end of the ribbon faster.
5. if more than one walk encodes the same minimal bare label, 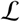, distinguished from each other by distinct orientation flags, we weight the orientation flags as follows. Any label has associated orientation index, 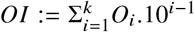: the canonical label minimises the orientation index. Denote this canonical label 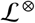. The first rule allows us to choose a unique origin and direction if the fold contains multiple ribbons with shortest, second-shortest,…, spans. If all spans are equal, annular ribbons are leftmost in the flagged canonical label.

Those rules are sufficient to determine unique canonical labels for folds sharing a common semi-flagged ribbon diagram 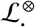.

Examples of non-canonical and canonical labels 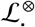 are shown in Fig. 49. The six-ribbon fold diagram in Fig. 49(c) has smallest digit string 121345326456, so its canonical label is 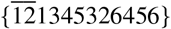, corresponding to a clockwise or anticlockwise circuit around the closed strand.

**Figure 48:**
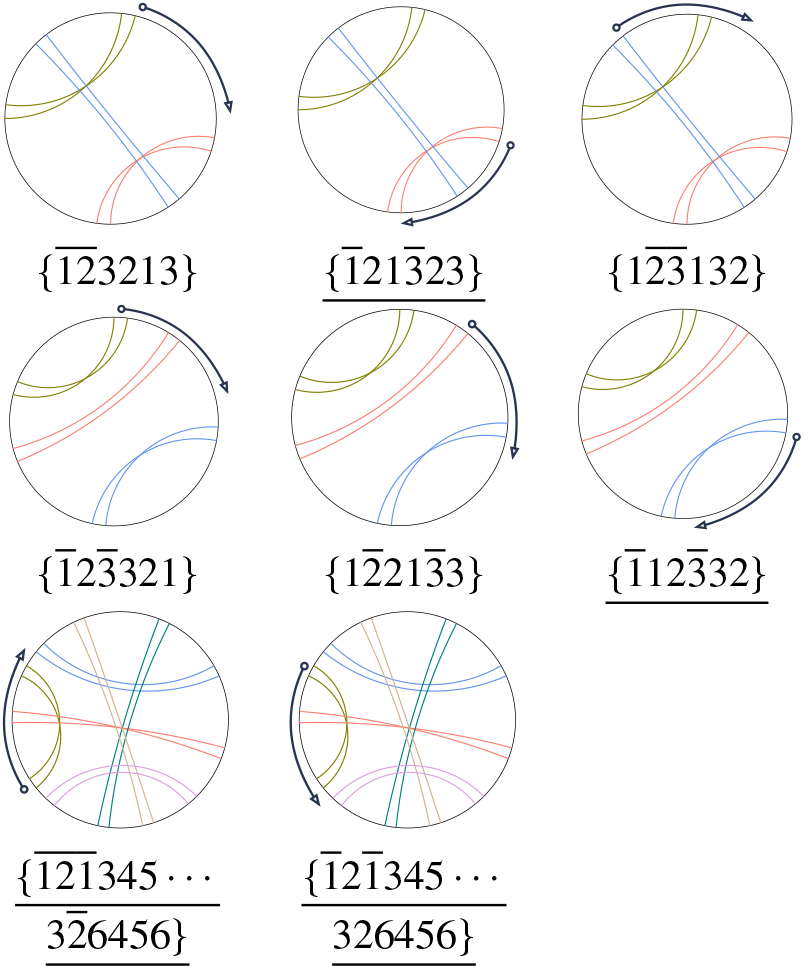
Possible fold labels 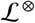. Canonical labels are <u>underlined</u> for each fold.

**Figure 49:**
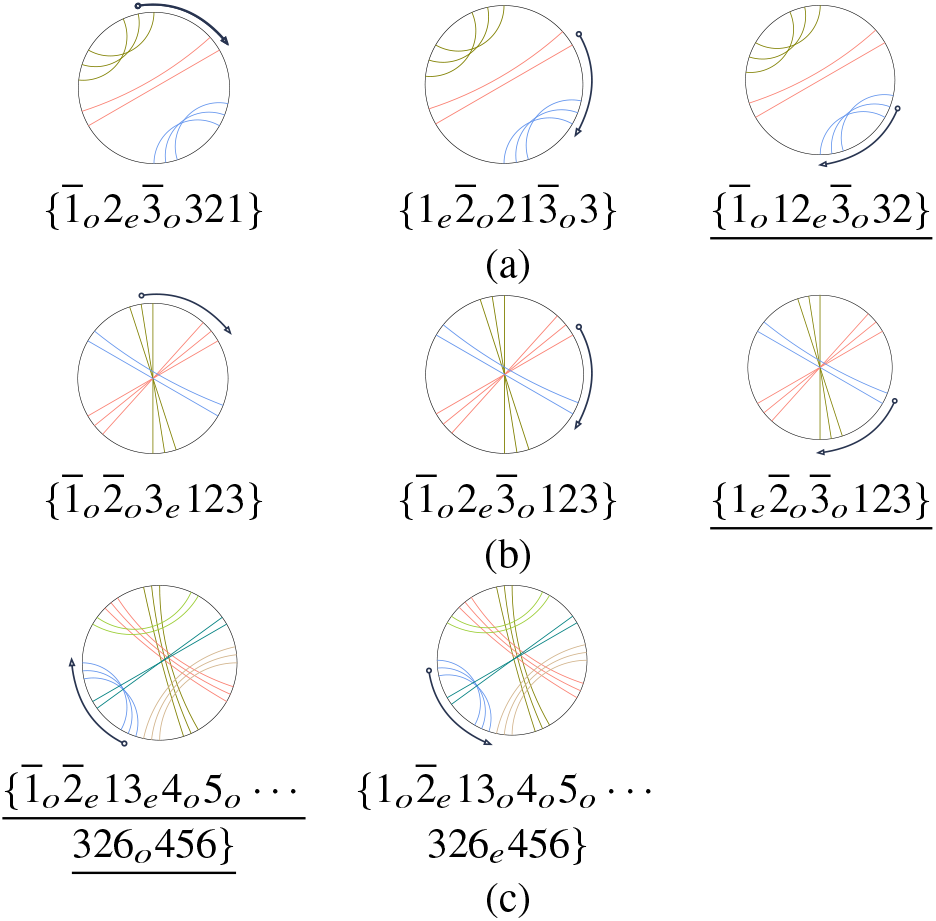
Possible fold labels for (a-c) three different circular diagrams. Canonical flagged labels are <u>underlined</u> for each fold.

#### Fully-flagged canonical fold labels

In order to distinguish the simplest all-duplexed folds from others, lengthier canonical flagged labels are required, which include twist parity flags, in addition to the ribbon orientation flags. Canonical fully-flagged labels are deduced by adding an additional flag index for twist parity, giving the complete algorithm:

1. from some origin, walk clockwise or anticlockwise around the perimeter once, and label each new ribbon terminal by successive digits, 1, 2,…, so that every digit is listed twice: one entry for each terminal. All closed tours around around the perimeter are therefore encoded by a label made of a 2*k*-digit string. The labels (may) depend on the starting terminal and the direction of the cyclic tour.
2. append an orientation flag to each ribbon in the label, identifying it as annular or moebius, i.e. if a ribbon *i* is annular, the ribbon has positive index *i* and associated orientation index ⊗_i_ = 0; if it is moebius, the ribbon has negative index, ī and associated orientation index ⊗_i_ = 1
3. append a parity flag subscript to each digit, *P*_i_ = 0,1 if *t*_i_ is even/odd, giving a parity index 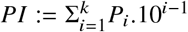.
4. a ‘fully flagged’ label for a walk is the 2*k*-digit string, with parity and orientation flags appended to each digit. (Since each digit *i* appears twice, to save space, we omit the flags on the second, rightmost entry to form the ‘flagged label’, e.g. 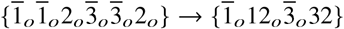
5. any ‘bare’ label is a string of positive digits, stripped of orientation and parity flags, which is an integer. The canonical bare label, denoted 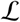, is the smallest integer.
6. a label has associated orientation and parity indices, 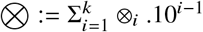 and 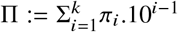: the canonical flagged label minimises the orientation weighting, 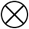. Denote the canonical fully flagged label 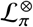 and (partially flagged) labels denoting the ribbon orientations only 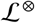.
7. if the rules above fail to produce a unique canonical flagged label, a tie-breaker is applied: the canonical flagged label minimises the parity weighting, Π, promoting even-parity ribbons over others.

Those rules are sufficient to determine unique canonical dressed labels for all folds.

They also allow sorting of folds by first (i) increasing 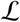, then (ii) increasing *OI* and lastly (iii) increasing *PI*.

Consider, for example the fold in Fig. 49(a). Since the fold diagram has bilateral symmetry, there are three flagged labels, dependent on the origin, and independent of the direction of a cyclic walk around the perimeter, as shown in Fig. 49. Just one label has smallest bare digit string (112332), therefore the canonical flagged label for this fold is 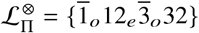. In contrast, the fold in Fig. 49(b) has bare label 123123 for all circuits, with possible flagged labels:

- 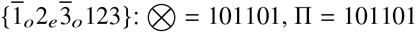
- 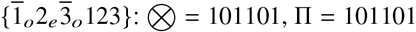
- 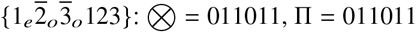

Since 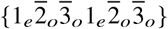 minimises 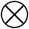, the canonical label for this fold is 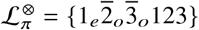.

The fold in Fig. 49(c) has smallest digit string 121345326456, with alternative flagged labels:

- 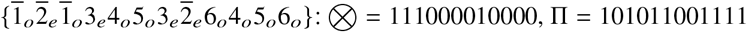
- 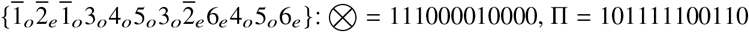

Both flagged labels have equal 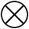, but 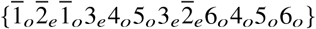 minimises Π, so the canonical flagged label for fold (xviii) is 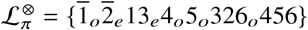.

#### Canonical labels for specific members of fold families

Specific folds, with integer twists *t*_i_ are denoted by replacing twist parity subscripts, *o* or *e* by *t_i_*. For example, the canonical fold label

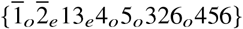

includes folds

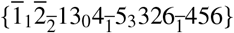

and

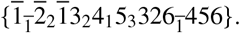

These are canonical labels, since both folds lie within the fold (family) 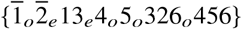.

For some fold families, the canonical flagged label needs a further criterion to give a canonical label for specific folds. For example, a specific fold in the family with fully flagged fold label

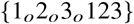

with three odd-parity twists 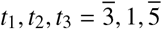 has possible labels:

1. 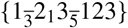
2. 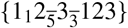
3. 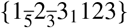
4. 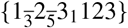
5. 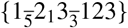
6. 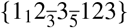

The canonical label for a specific fold minimises the twist parity weighting, Π, followed by (ii) the minimiser of the absolute unsigned twist ordinal, 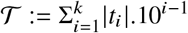. Thus, for the six alternative labels above,

1. 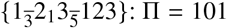
2. 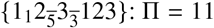
3. 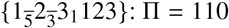
4. 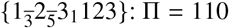
5. 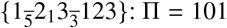
6. 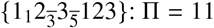

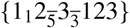 and { 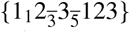 minimise Π (= 11). Those two labels have 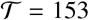 and 135 respectively. Therefore the canonical flagged label for this specific fold is 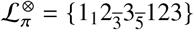.

### A DETAILED EXAMPLE OF A FOLD LABEL, ITS CIRCULAR DIAGRAM AND POSSIBLE EMBEDDINGS

Fig. 50 illustrates in detail the relationship between (a) the circular diagram of the fully-flagged fold 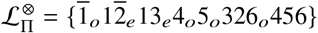, which is a relaxed fold, fully-duplexed on a genus-3 skeleton. Two simple embeddings of that skeleton are possible: one resembling a four-rung ladder and the other a pair of concentric circles joined by horizontal edges. The corresponding relaxed fold embeddings are drawn schematically in (b,c), with parallel pairs of duplexed strands along each skeletal edge. Each edge contains a switch box colored blue or yellow, representing parallel and antiparallel duplexes respectively. The boxes are labelled *i_e_* or *i_o_*, where *i* (= 1,…, 6) is the label of that skeletal edge inherited from the canonical label 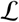 in (a). Within each box, strand pairs are subjected to an even *e* or odd *o* number of half-twists depending on the relevant parity flag Π = *oeeooo*, forming double-helices. Those twist parities allow an infinite family of relaxed strand windings, provided *t*_i_ is tuned to the parity *π*_i_. One example, with 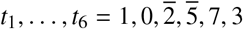 (generating a fold label 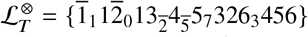) is shown in Figs. 50(d-g). Double-helices are labelled by their corresponding ribbon label *i* in (d) and twist *t*_i_ in (f). The directed winding shown in (g) reveals parallel duplexes in *i* = 1 - 2 and antiparallel duplexes in *i* = 3 - 6, consistent with the semi-flagged fold label 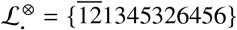.

**Figure 50:**
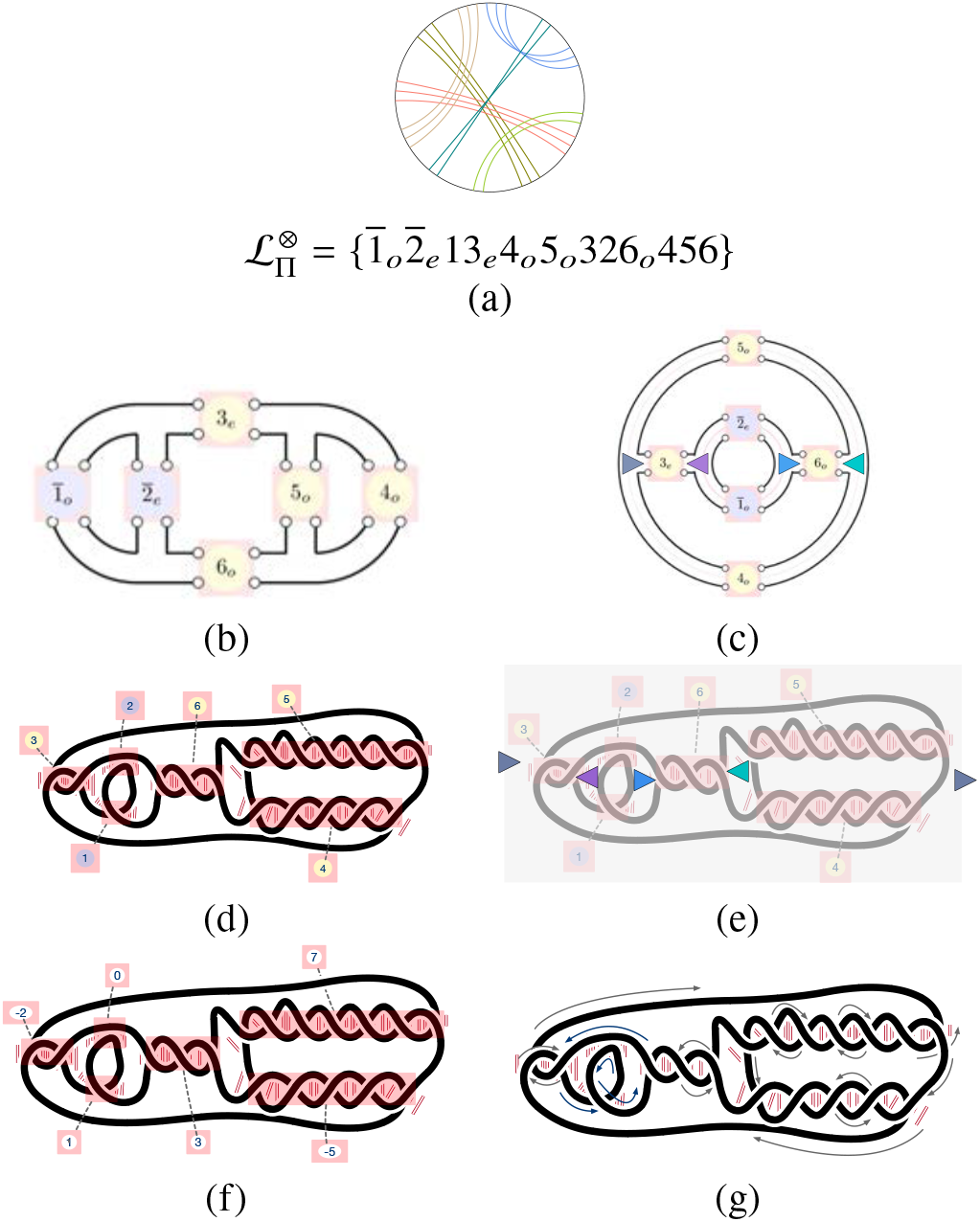
A fully-flagged fold derived from the semi-flagged fold 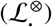 listed as fold 12 in Table 2. (An embedding of the oriented stand graph is shown in the Table.) (b) An embedding of the genus-3 fold, with (blue) parallel and (yellow) anti-parallel duplexes marked by swtich boxes (as in Fig. 12), labelled by ribbon number and orientation and twist parity. (c) A second embedding, with the conventional Y-junctions marked as colored triangles. (d) A specific strand winding sharing fully-flagged label 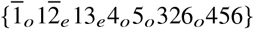, namely 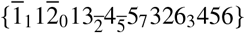. The fold is similar to the embedding in (b), with duplexes 4 and 5 dragged (and rotated) to the far right of the fold. (e) The same strand winding, with Y-junctions colored according to the coloring in (b). (The grey triangle appears at either side of the diagram. It is realised as a three-way junction in the fold by wrapping the LHS and RHS ends to glue the two copies into a single copy.) (f) The fold, labelled with numbers of half-twists twists, *t*_i_, for each duplex. (f) The directed strand winding. (Blue) duplexes corresponding to moebius ribbons are parallel windings of the parallel strands and (yellow) duplexes corresponding to annular ribbons are antiparallel.

### RELAXED CONTRACTED FULLY-FLAGGED FOLDS

**Table 18:**
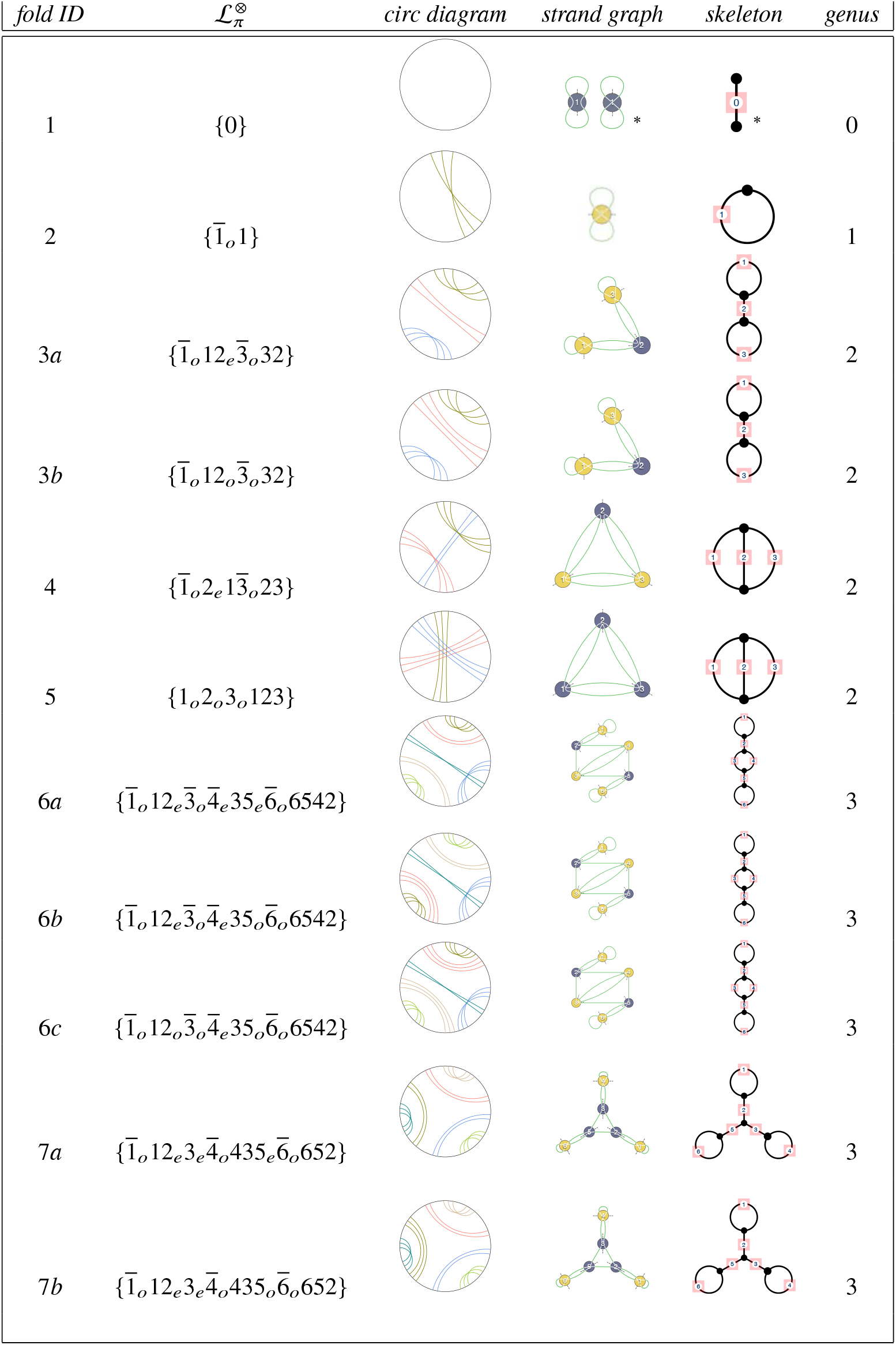
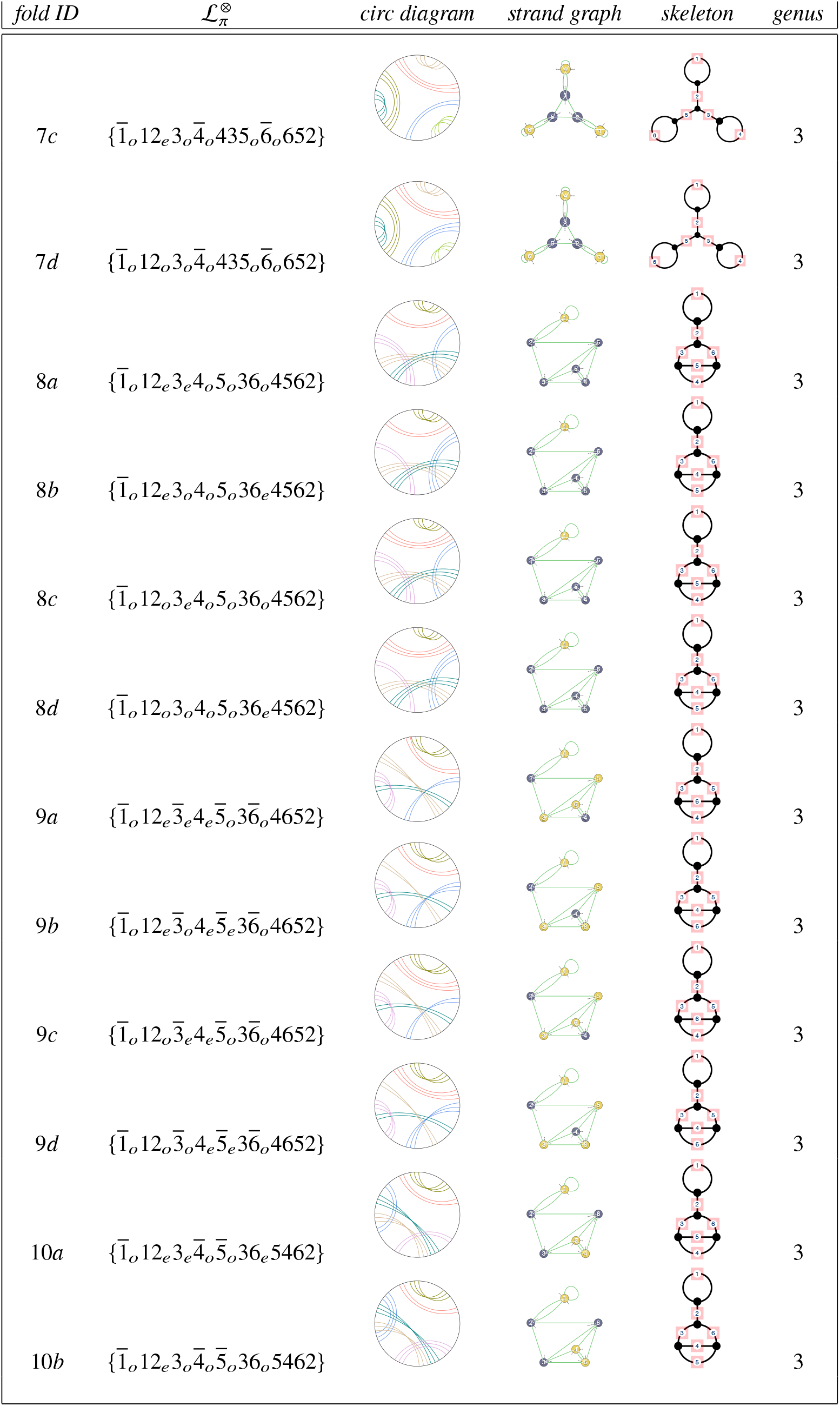
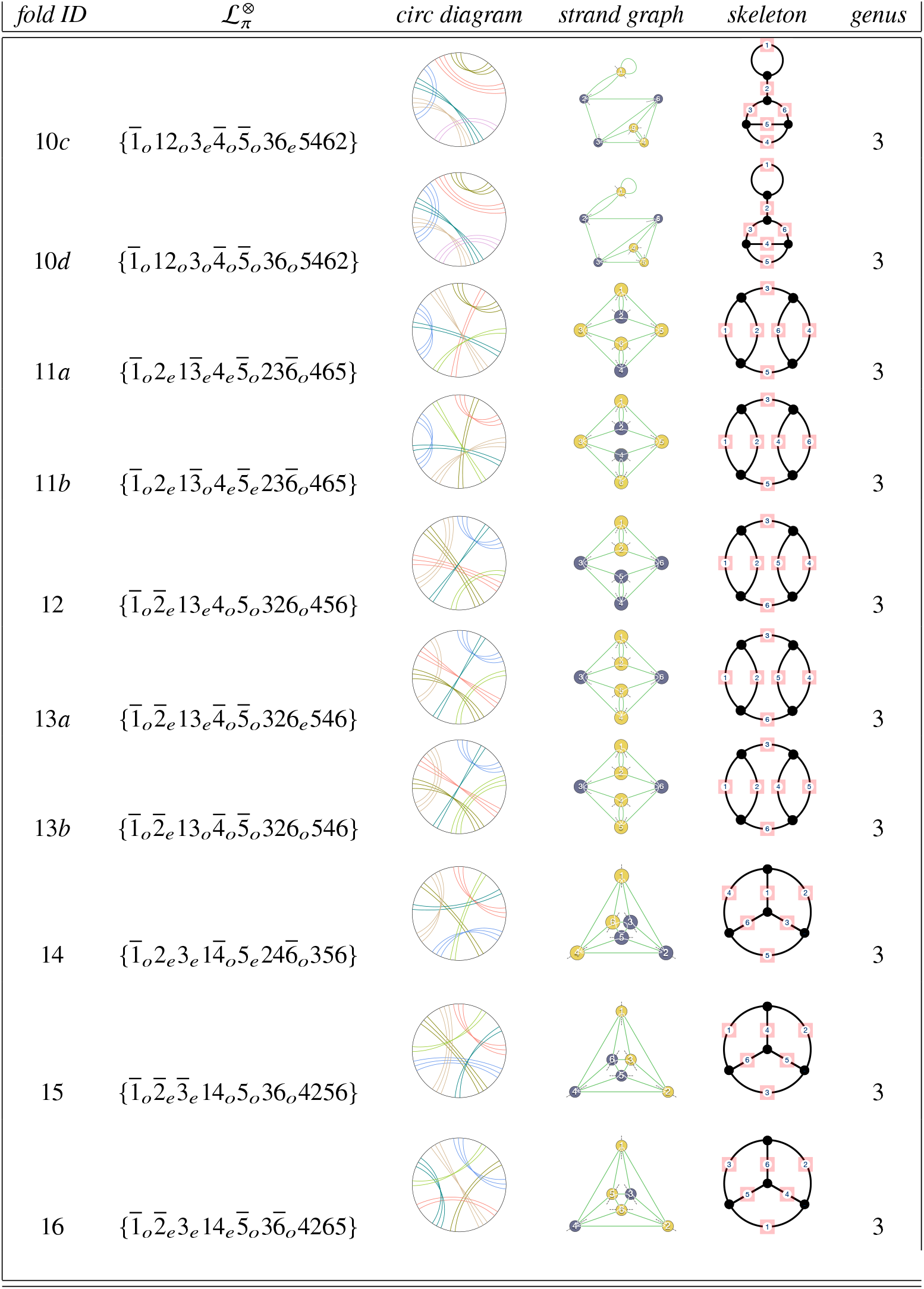
Fully-flagged labels, skeletons and circular diagrams for relaxed genus-0 - 3 folds found by appending tuned parity flags to the semiflagged folds in Table 3 in the main text. The fold ID refers to the fold reference in Tables 1 – 3 in the main text of the paper. Plumblines in oriented strand graphs are marked by dotted lines through vertices. Graph vertices are colored blue-gray and mustard for vertices hosting antiparallel and parallel double-helices respectively. (* The strand graphs and skeleton of the (unfold) {0} are uncontracted; the contracted fold is empty.)

| *fold ID* | $\mathcal{L}_\pi^\otimes$ | *circ diagram* | *strand graph* | *skeleton* | *genus* |
| --- | --- | --- | --- | --- | --- |
| 1 | {0} |  |  |  | 0 |
| 2 | $\{\bar{1}_o 1\}$ | | | | 1 |
| 3a | $\{\bar{1}_o 12_e \bar{3}_o 32\}$ | | | | 2 |
| 3b | $\{\bar{1}_o 12_o \bar{3}_o 32\}$ | | | | 2 |
| 4 | $\{\bar{1}_o 2_e 1\bar{3}_o 23\}$ | | | | 2 |
| 5 | $\{1_o 2_o 3_o 123\}$ | | | | 2 |
| 6a | $\{\bar{1}_o 12_e \bar{3}_o \bar{4}_e 35_e \bar{6}_o 6542\}$ | | | | 3 |
| 6b | $\{\bar{1}_o 12_e \bar{3}_o \bar{4}_e 35_o \bar{6}_o 6542\}$ | | | | 3 |
| 6c | $\{\bar{1}_o 12_o \bar{3}_o \bar{4}_e 35_o \bar{6}_o 6542\}$ | | | | 3 |
| 7a | $\{\bar{1}_o 12_e 3_e \bar{4}_o 435_e \bar{6}_o 652\}$ | | | | 3 |
| 7b | $\{\bar{1}_o 12_e 3_e \bar{4}_o 435_o \bar{6}_o 652\}$ | | | | 3 |
| 7c | $\{\bar{1}_o 12_e 3_o \bar{4}_o 435_o \bar{6}_o 652\}$ | | | | 3 |
| 7d | $\{\bar{1}_o 12_o 3_o \bar{4}_o 435_o \bar{6}_o 652\}$ | | | | 3 |
| 8a | $\{\bar{1}_o 12_e 3_e 4_o 5_o 36_o 4562\}$ | | | | 3 |
| 8b | $\{\bar{1}_o 12_e 3_o 4_o 5_o 36_e 4562\}$ | | | | 3 |
| 8c | $\{\bar{1}_o 12_o 3_e 4_o 5_o 36_o 4562\}$ | | | | 3 |
| 8d | $\{\bar{1}_o 12_o 3_o 4_o 5_o 36_e 4562\}$ | | | | 3 |
| 9a | $\{\bar{1}_o 12_e \bar{3}_e 4_e \bar{5}_o 3\bar{6}_o 4652\}$ | | | | 3 |
| 9b | $\{\bar{1}_o 12_e \bar{3}_o 4_e \bar{5}_e 3\bar{6}_o 4652\}$ | | | | 3 |
| 9c | $\{\bar{1}_o 12_o \bar{3}_e 4_e \bar{5}_o 3\bar{6}_o 4652\}$ | | | | 3 |
| 9d | $\{\bar{1}_o 12_o \bar{3}_o 4_e \bar{5}_e 3\bar{6}_o 4652\}$ | | | | 3 |
| 10a | $\{\bar{1}_o 12_e 3_e \bar{4}_o \bar{5}_o 36_e 5462\}$ | | | | 3 |
| 10b | $\{\bar{1}_o 12_e 3_o \bar{4}_o \bar{5}_o 36_o 5462\}$ | | | | 3 |
| 10c | $\{\bar{1}_o 12_o 3_e \bar{4}_o \bar{5}_o 36_e 5462\}$ | | | | 3 |
| 10d | $\{\bar{1}_o 12_o 3_o \bar{4}_o \bar{5}_o 36_o 5462\}$ | | | | 3 |
| 11a | $\{\bar{1}_o 2_e 1\bar{3}_e 4_e \bar{5}_o 23\bar{6}_o 465\}$ | | | | 3 |
| 11b | $\{\bar{1}_o 2_e 1\bar{3}_o 4_e \bar{5}_e 23\bar{6}_o 465\}$ | | | | 3 |
| 12 | $\{\bar{1}_o \bar{2}_e 13_e 4_o 5_o 32\bar{6}_o 456\}$ | | | | 3 |
| 13a | $\{\bar{1}_o \bar{2}_e 13_e \bar{4}_o \bar{5}_o 32\bar{6}_e 546\}$ | | | | 3 |
| 13b | $\{\bar{1}_o \bar{2}_e 13_o \bar{4}_o \bar{5}_o 32\bar{6}_o 546\}$ | | | | 3 |
| 14 | $\{\bar{1}_o 2_e 3_e 1\bar{4}_o 5_e 24\bar{6}_o 356\}$ | | | | 3 |
| 15 | $\{\bar{1}_o \bar{2}_e \bar{3}_e 14_o 5_o 36_o 4256\}$ | | | | 3 |
| 16 | $\{\bar{1}_o \bar{2}_e 3_e 14_e \bar{5}_o 3\bar{6}_o 4265\}$ | | | | 3 |

**Table 19:**
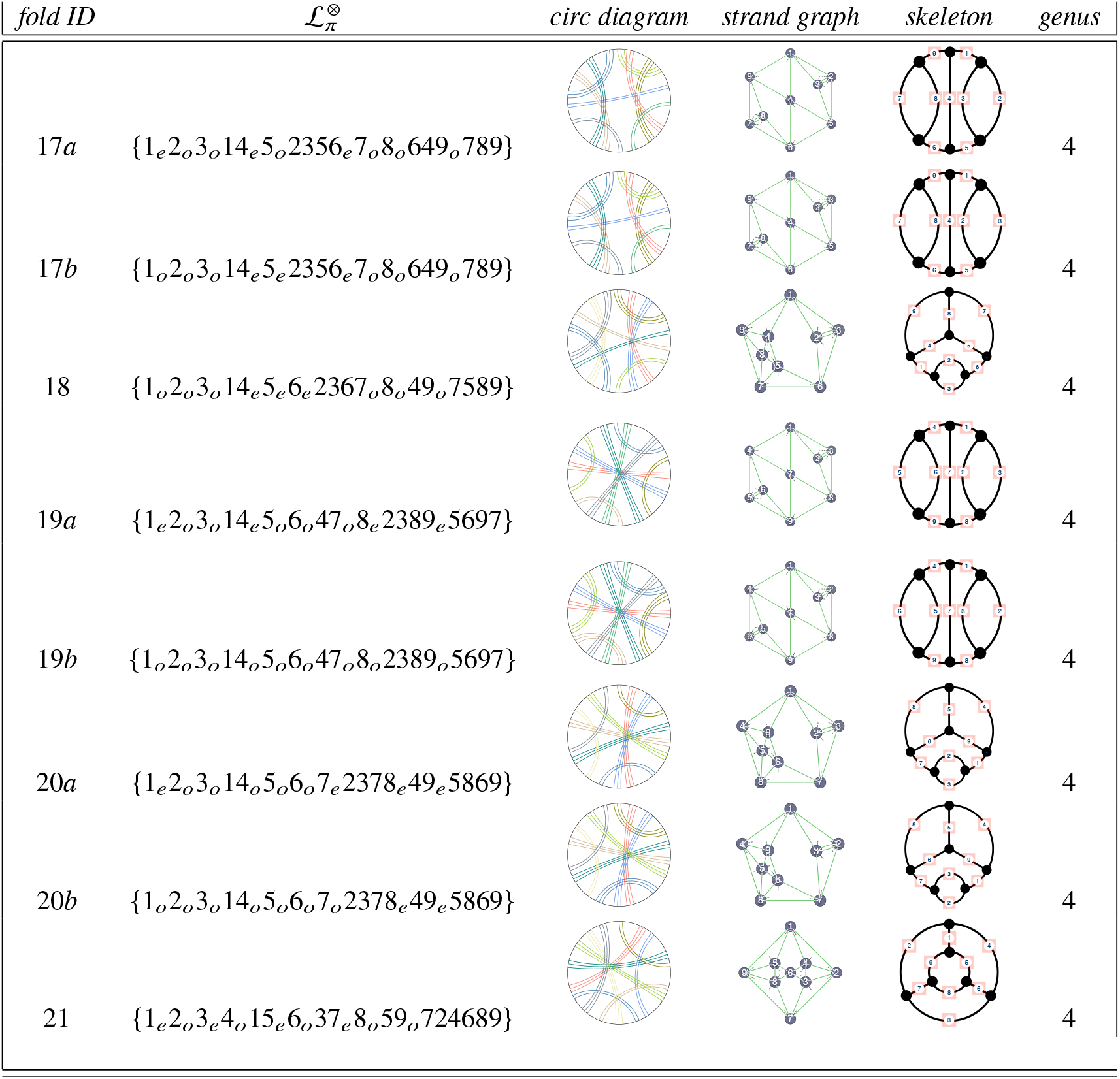
Fully-flagged labels, skeletons and circular diagrams for relaxed genus-4 folds found by appending tuned parity flags to the semiflagged folds in Table 4 in the main text, *cf*. Table 18. These parity flag data are courtesy of Neave Taylor.

### FROM A Y-JUNCTION FOLD TO ALL FOLDS CONTAINING X (FOUR-WAY) AND HIGHER JUNCTIONS

**Table 20:**
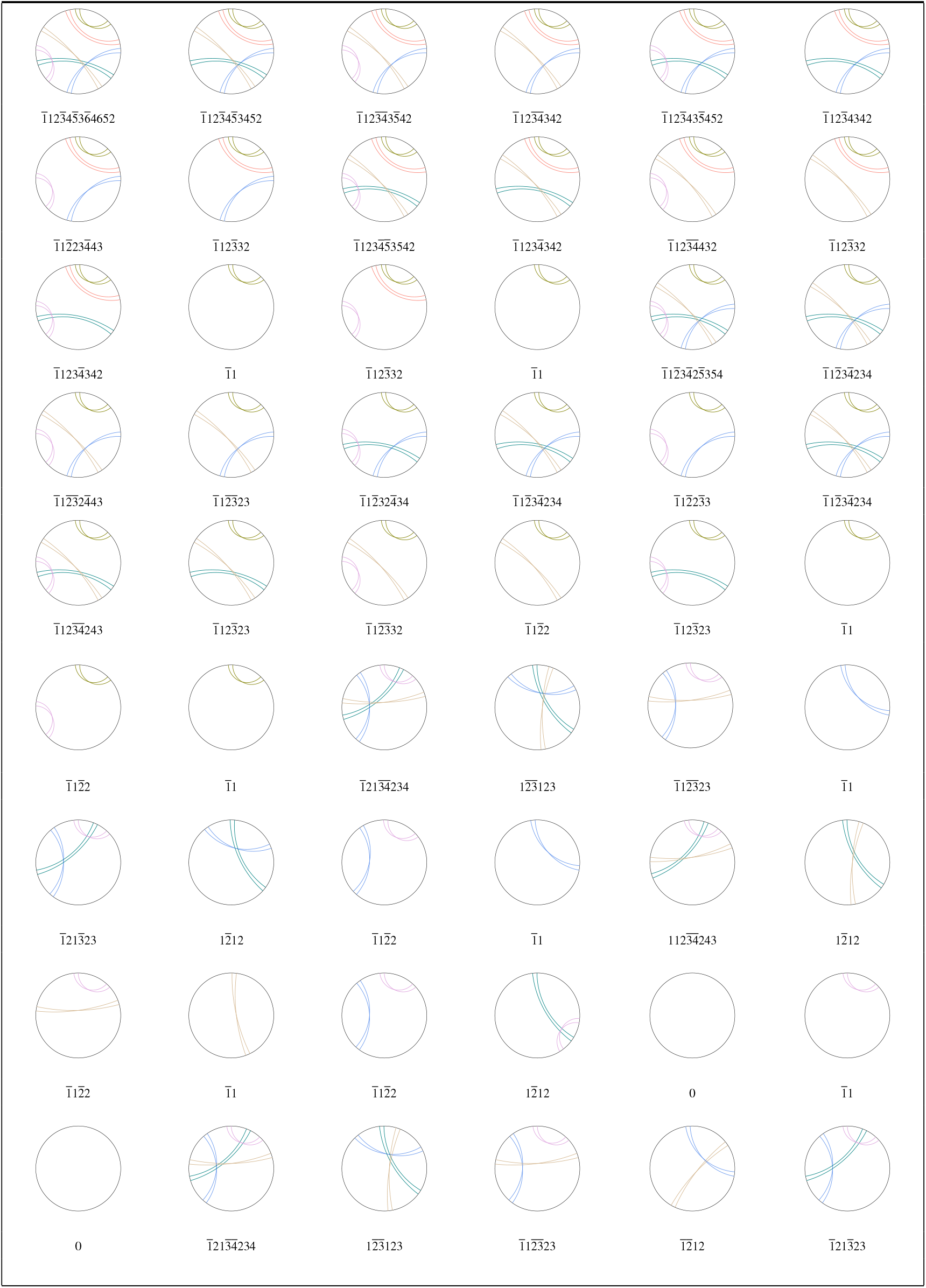
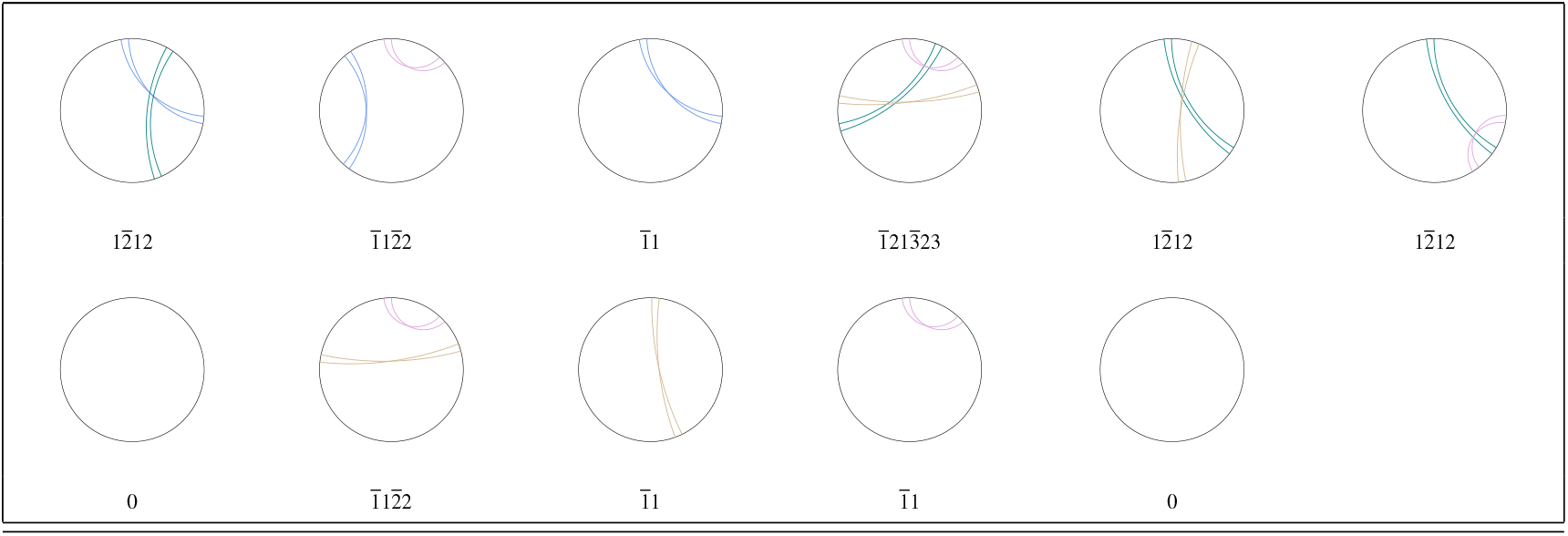
Contracted ribbon diagrams and their canonical semiflagged labels 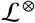 from the 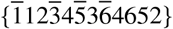 fold by ribbon deletions. (Ribbons are drawn with 2 rulings for clarity; twist parities are arbitrary since the folds are semiflagged.)

**Table 21:**
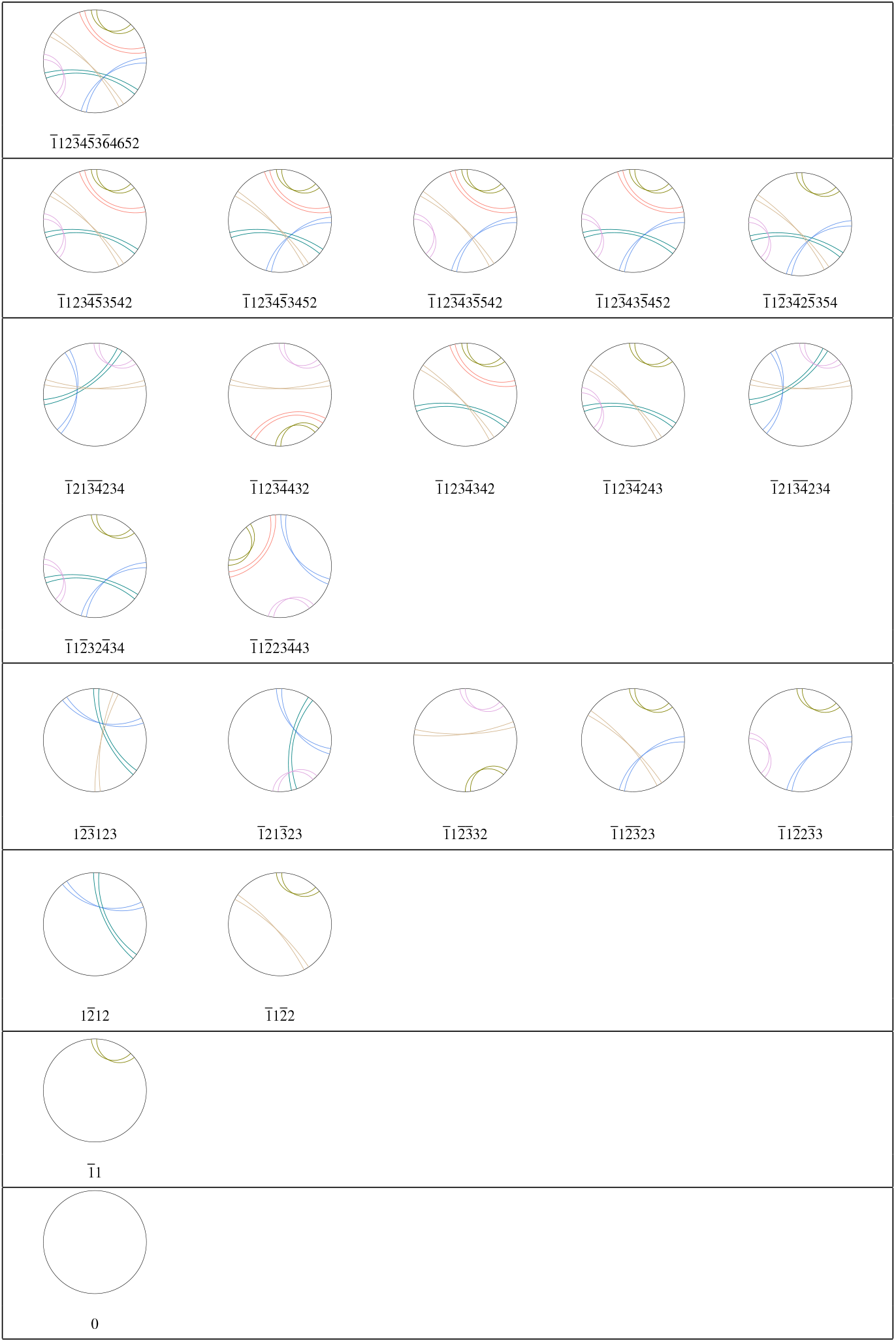
Distinct (contracted) ribbon diagrams derived from (and including) the 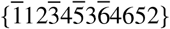 fold and semiflagged labels 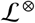. (Ribbons are drawn with 2 rulings for clarity; twist parities are arbitrary since the folds are semiflagged.)

The entries in Table 22 reduce to forty distinct folds, listed in Table 23, sorted by canonical fold label.

**Table 22:**
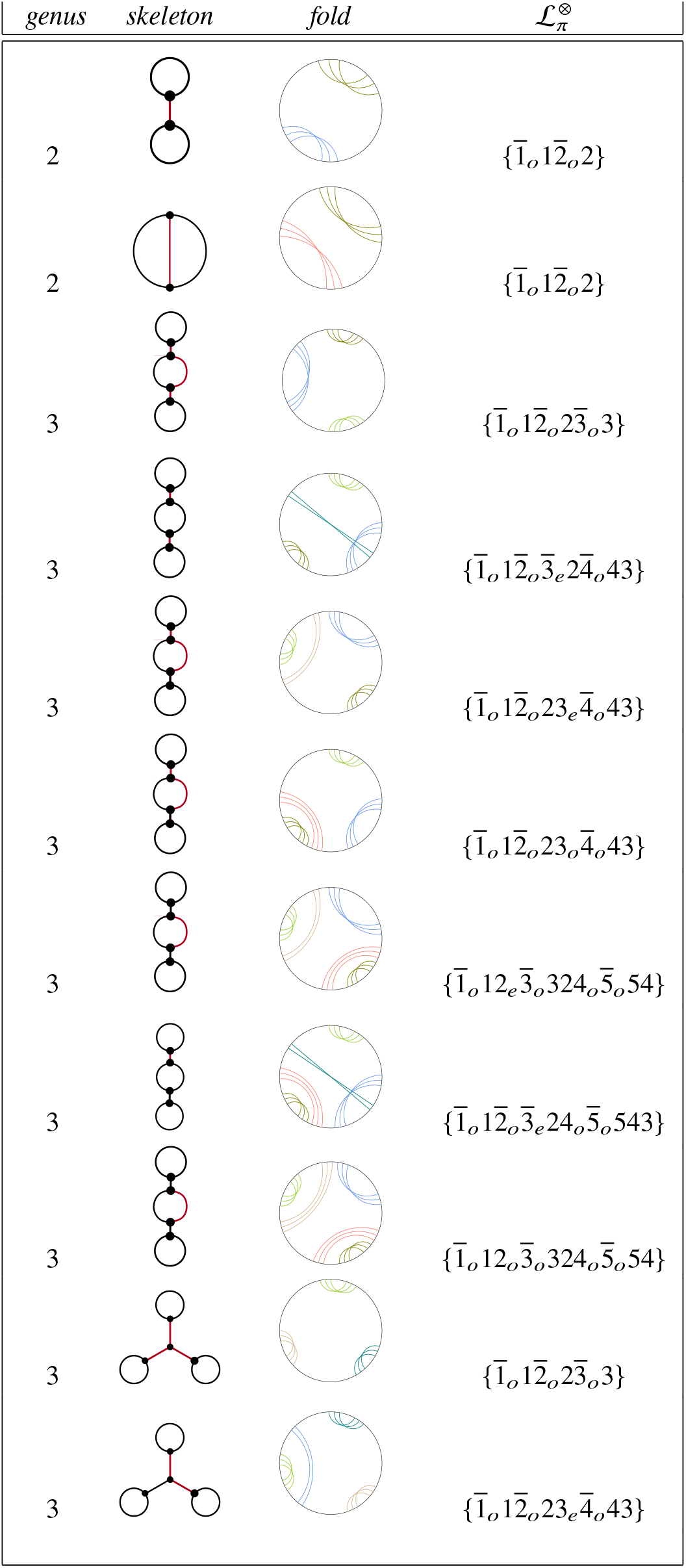
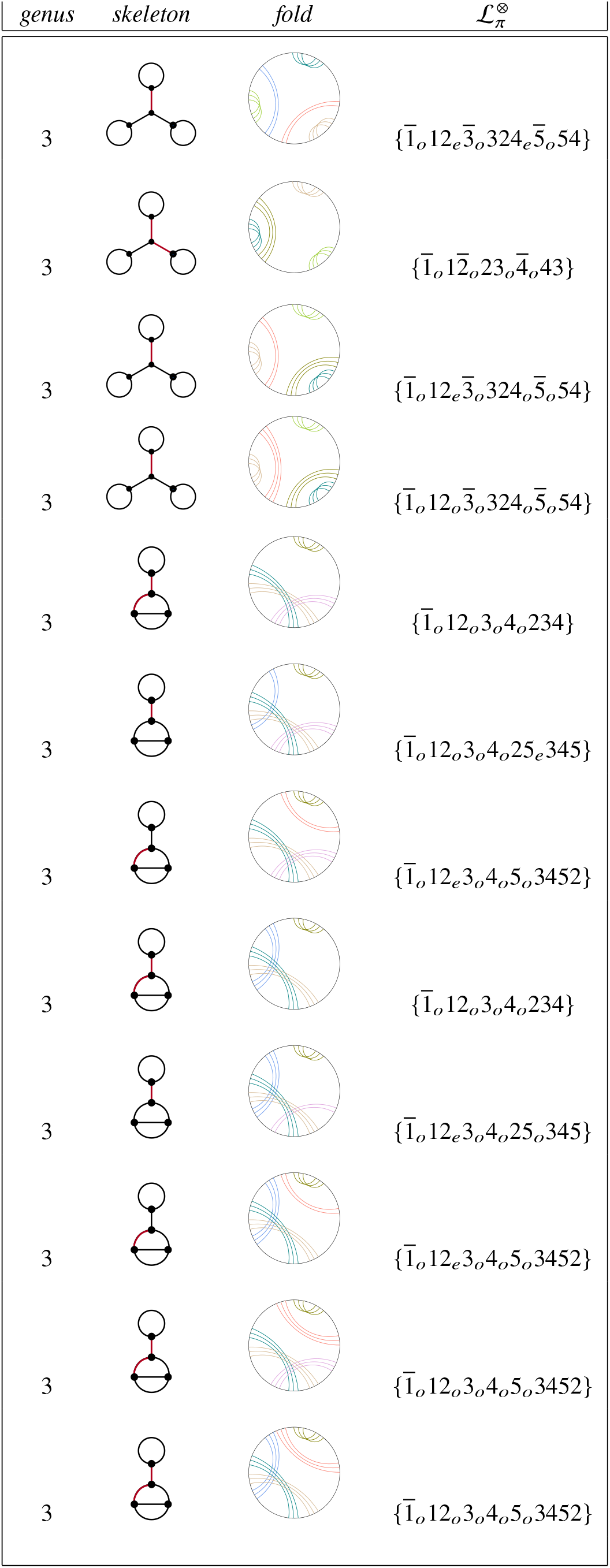
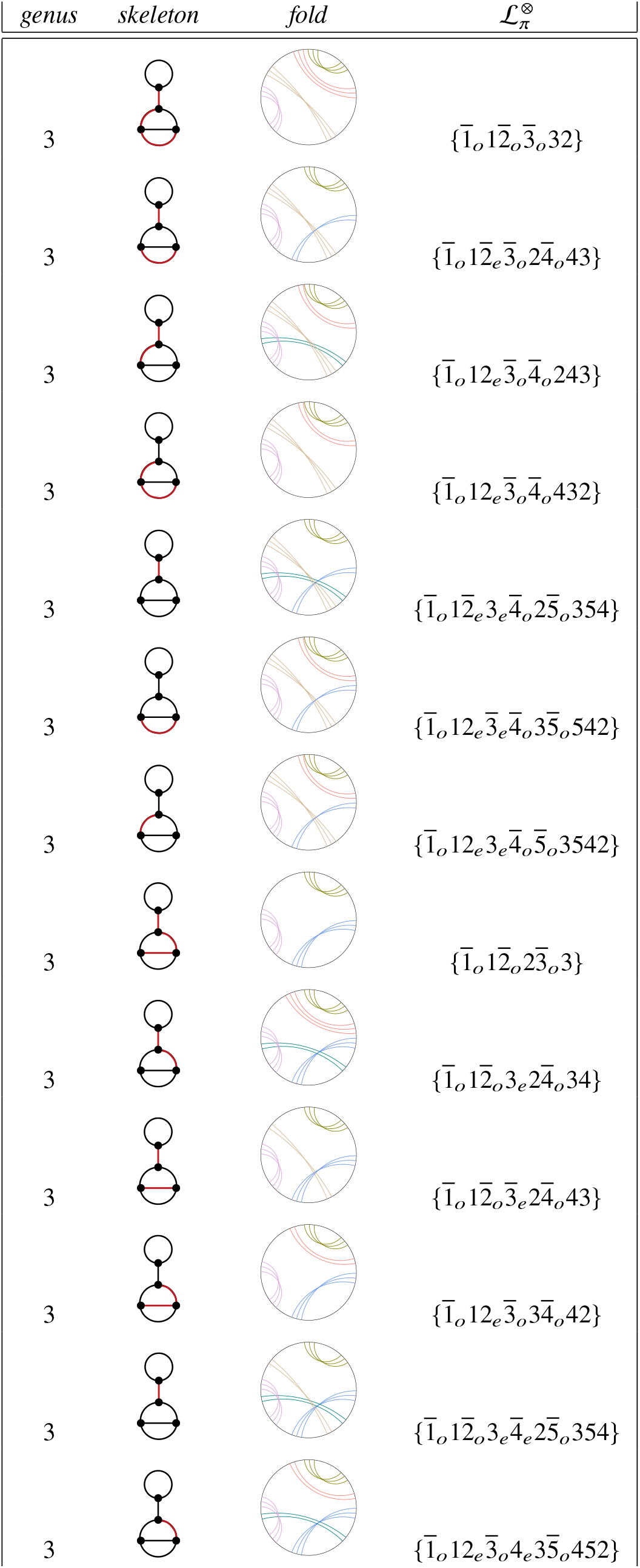
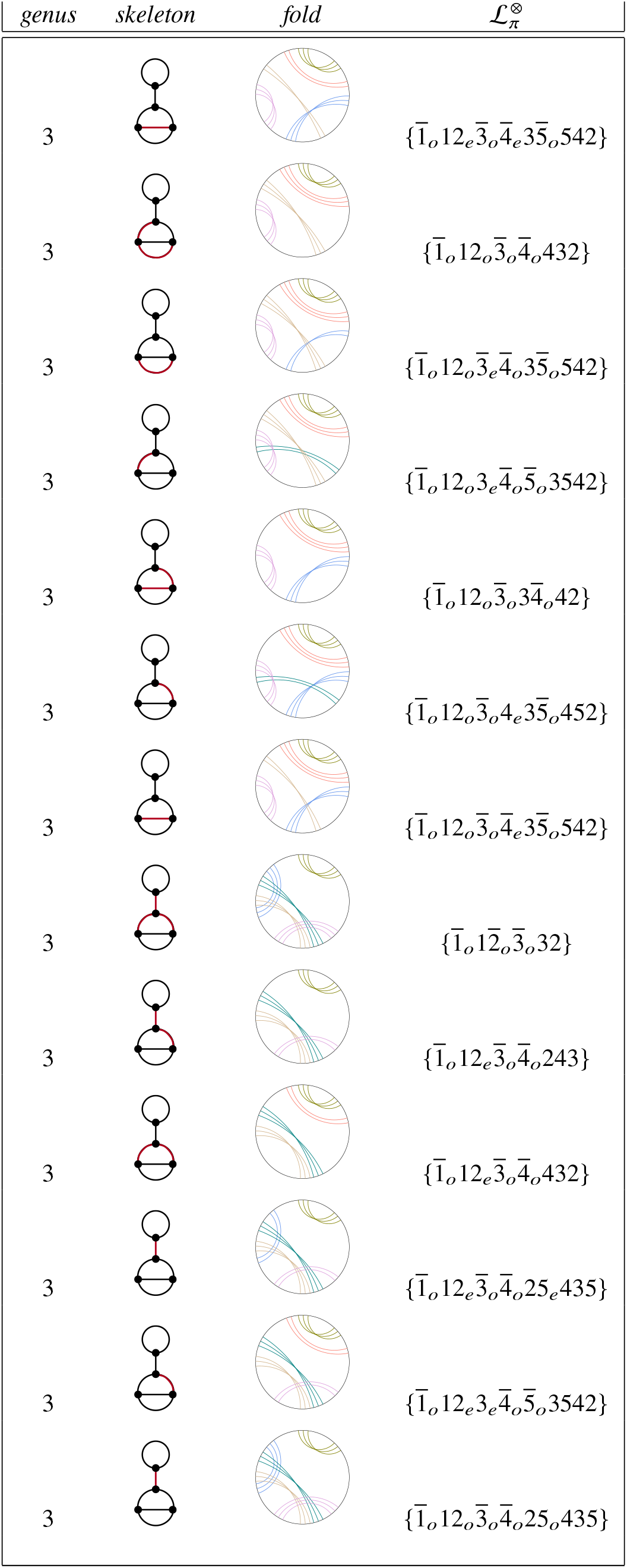
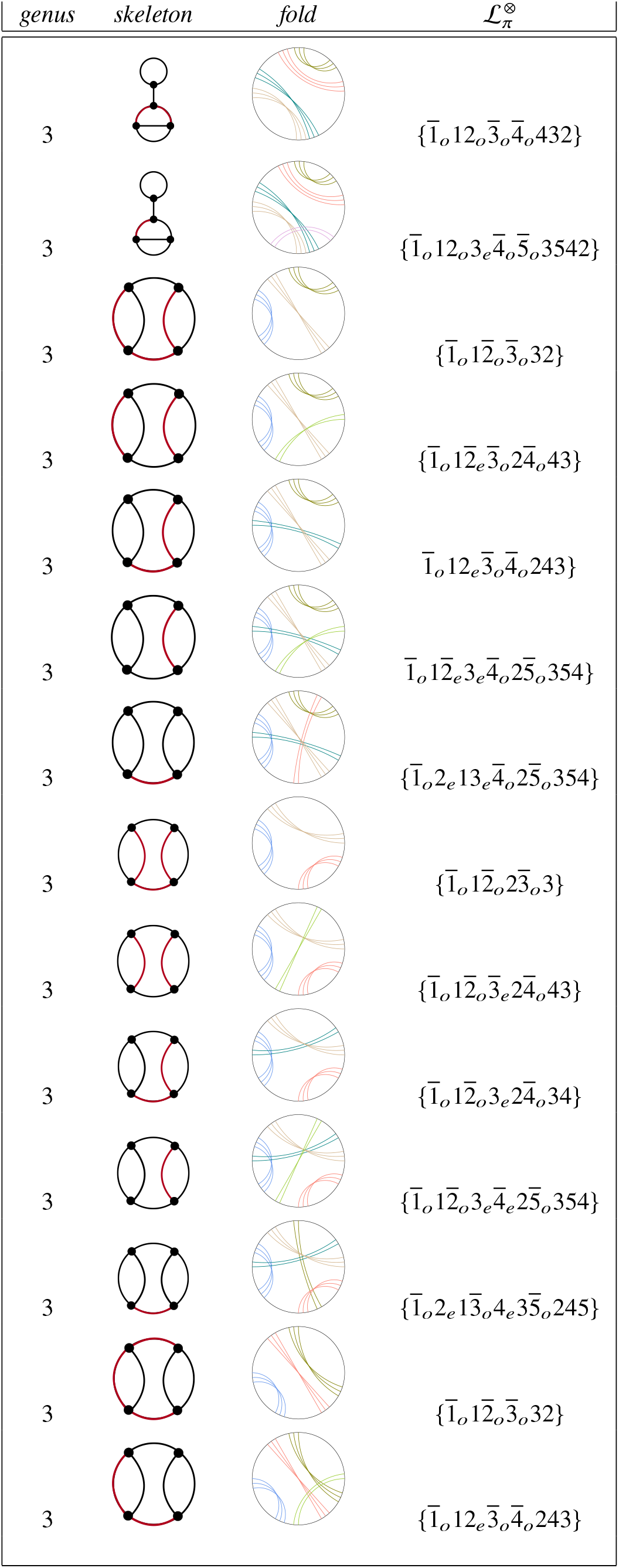
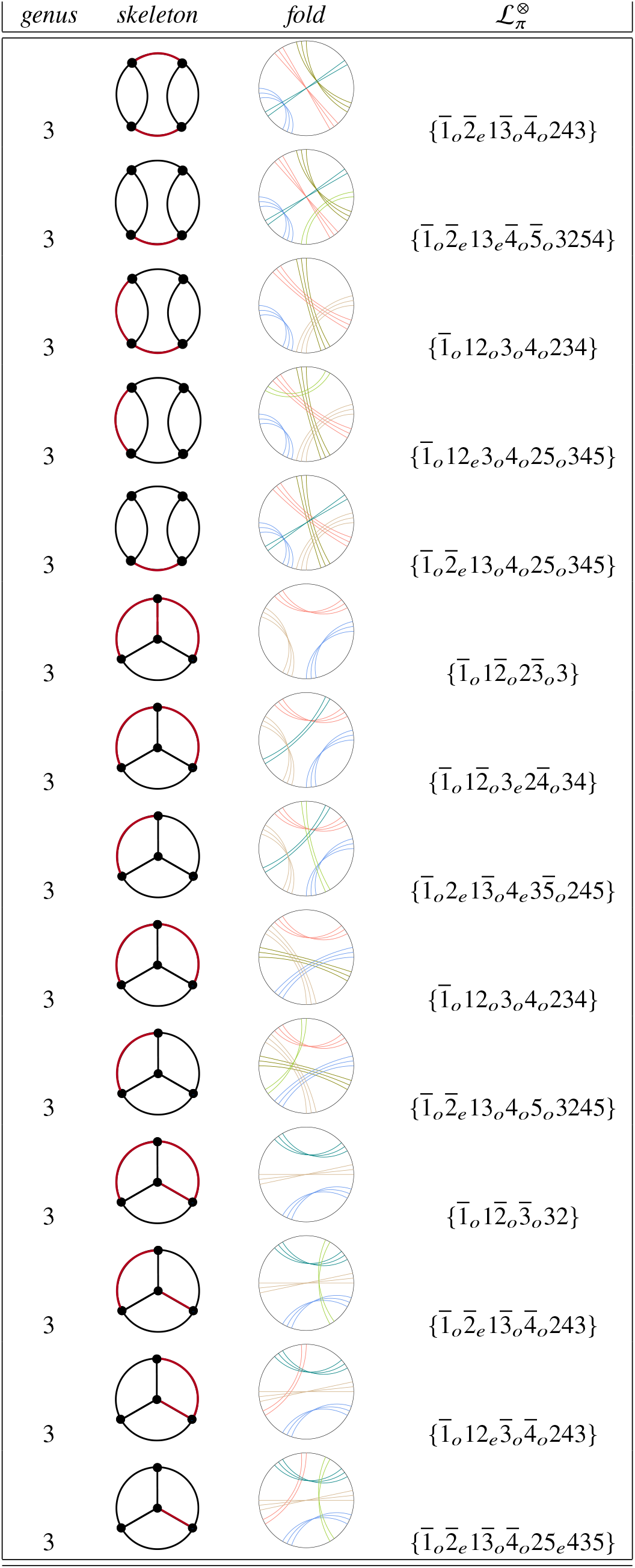
Relaxed folds with higher-order junctions, derived from the relaxed folds in Table 18 which contain exclusively Y-junctions, by deletion of some or all even-parity ribbons. Deleted edges are marked in red. Vertices at both ends of deleted edges are merged to form the skeleton of the handlebody hosting the fold.

**Table 23:**
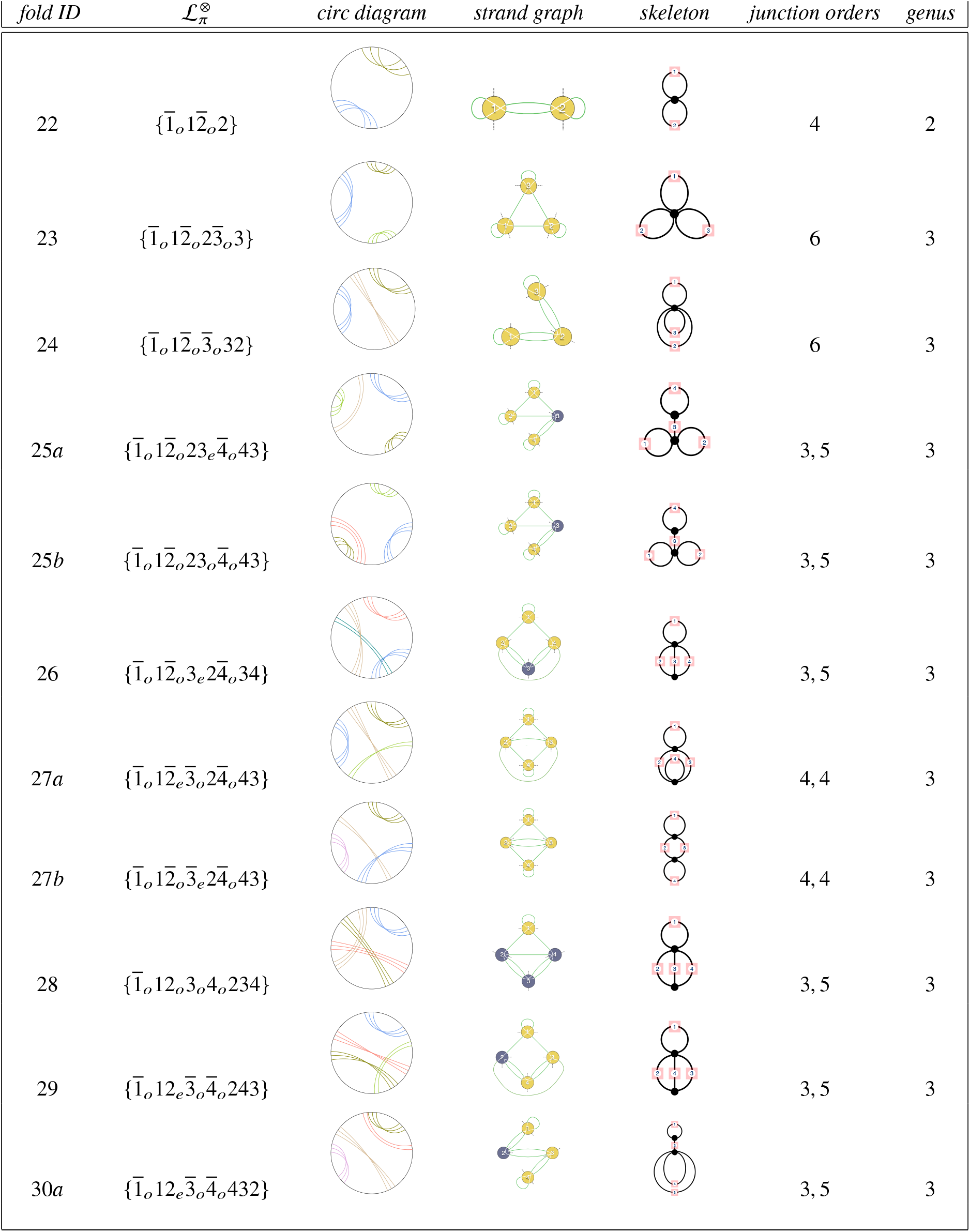
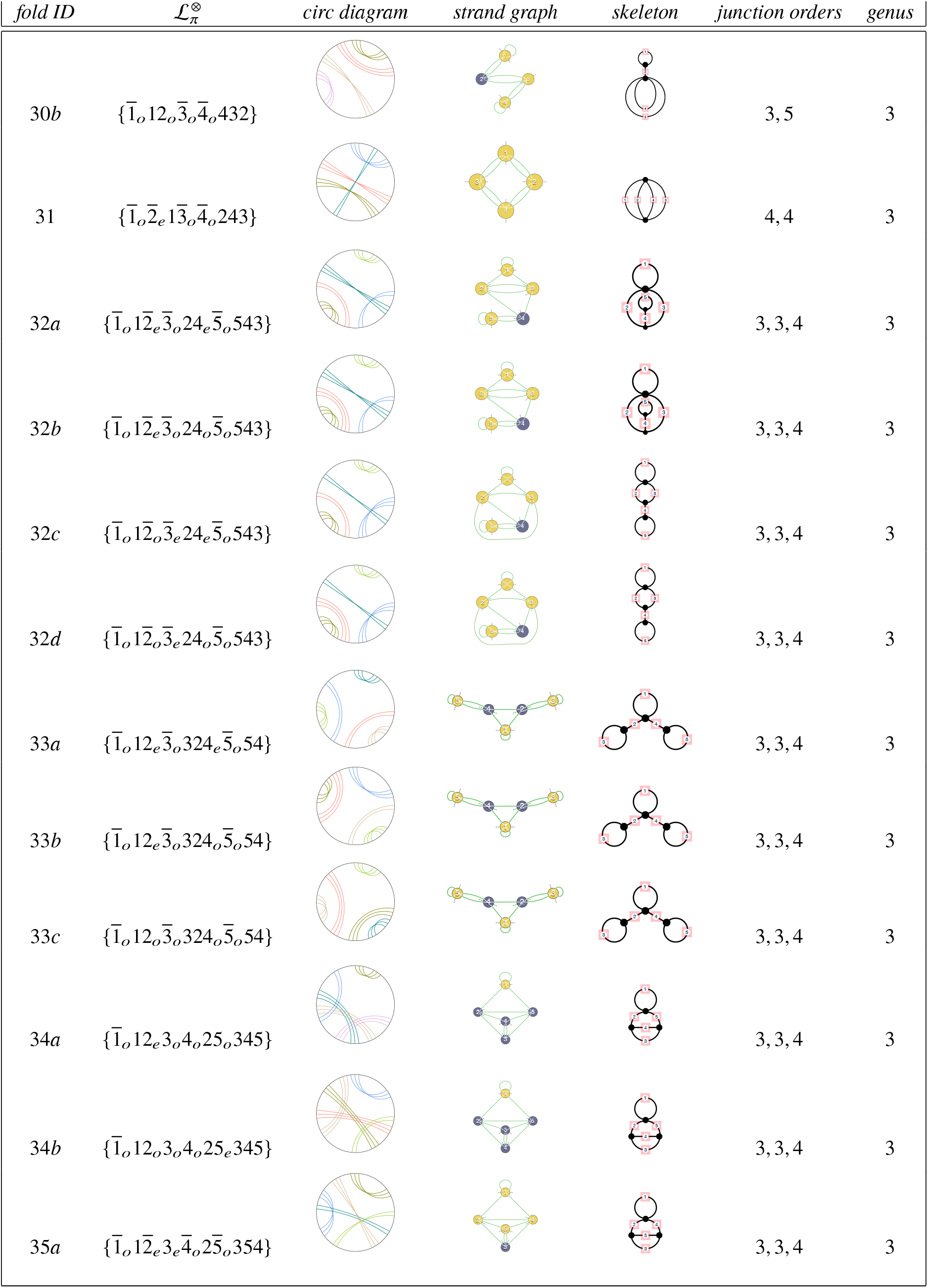
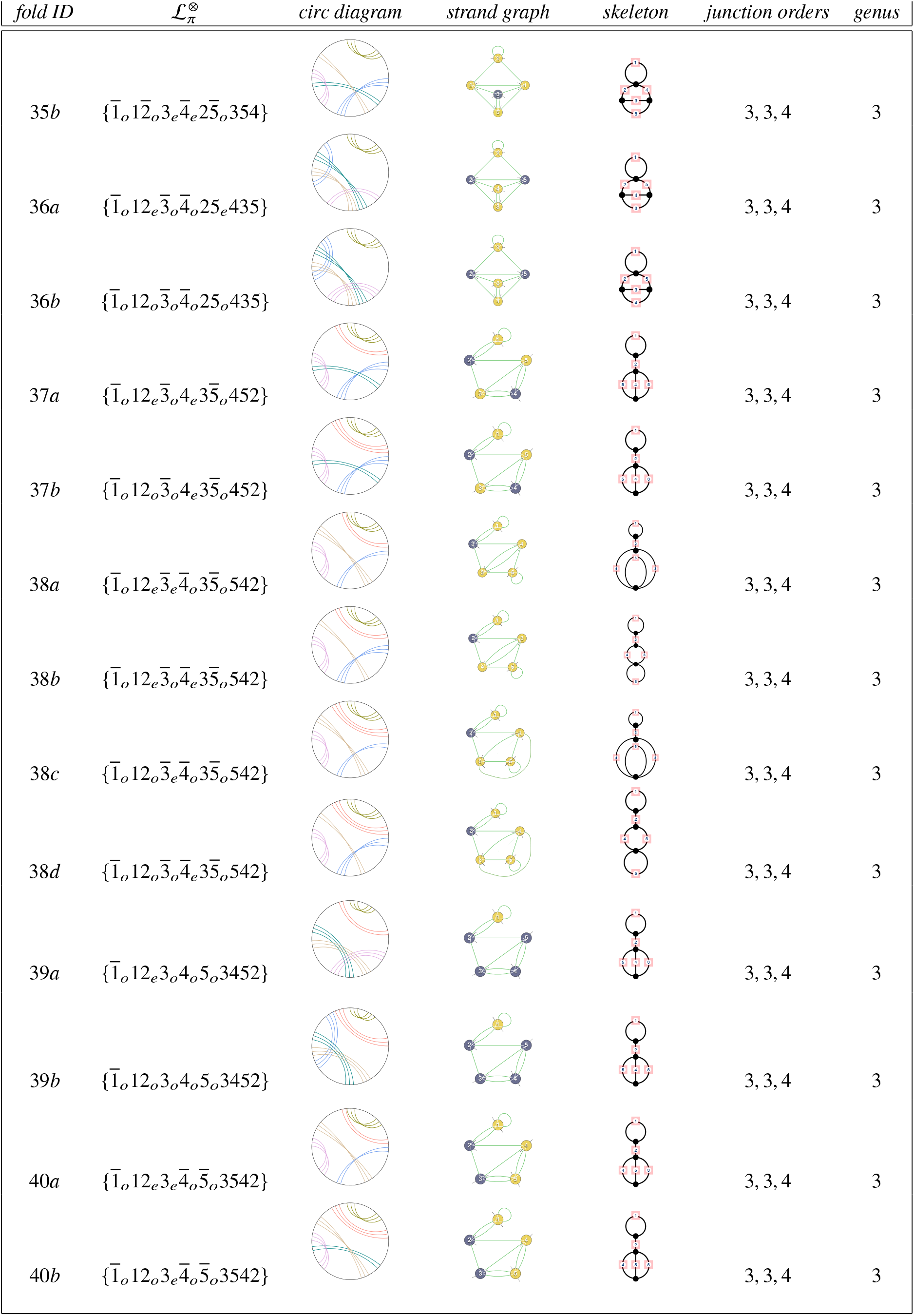
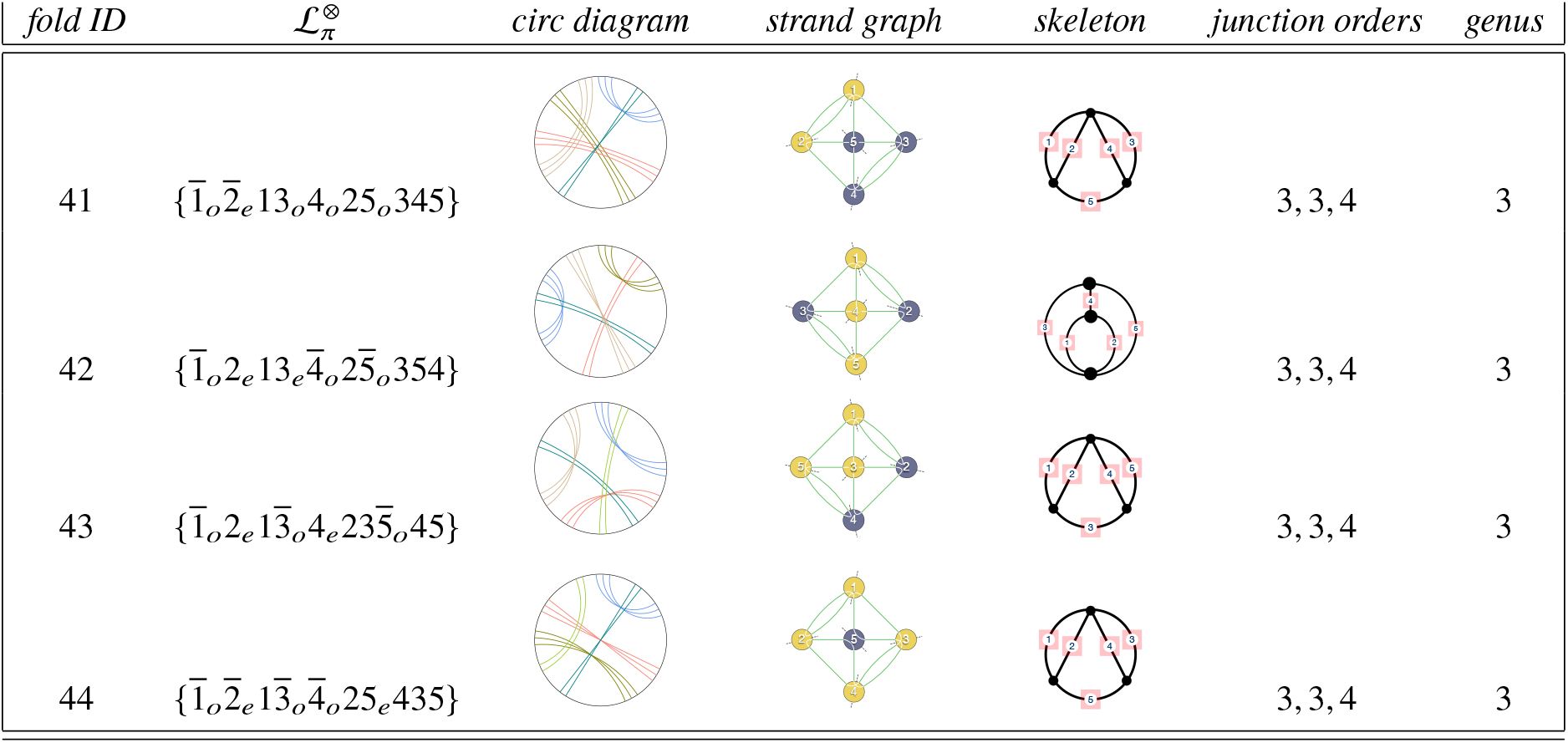
Catalog of distinct relaxed folds with (conventional) higher-order junctions (4-way, 5-way etc.), derived in Tables ??, ??, ?? and 22 sorted by canonical fold label. All folds with genus between 0-3 are included; there are no genus-0 or genus-1 relaxed folds with higher-order junctions. The Table lists the junction orders and the skeletons with twisted duplexed strands wound on each edge. Twist parities for each winding are equal to the parity subscripts associated with each ribbon in the fold label.

### CANONICAL UNFLAGGED LABELS WITH UP TO FIVE RIBBONS

**Table 24:**
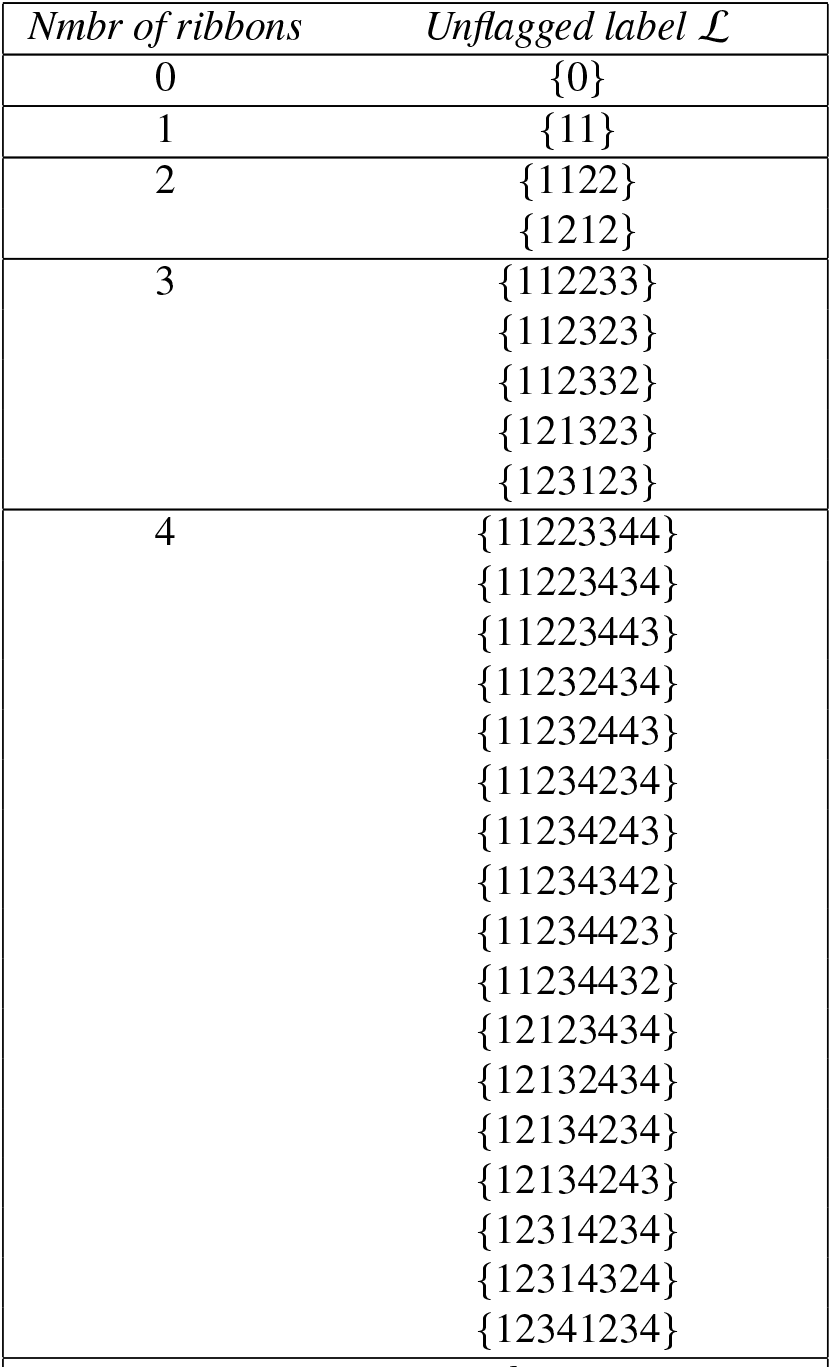
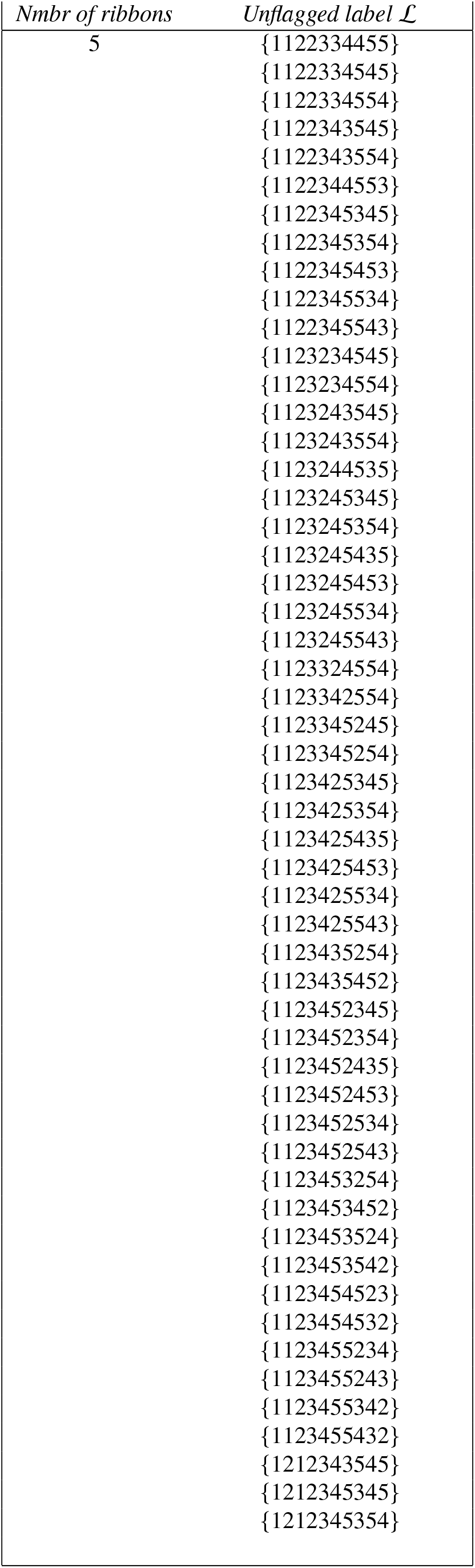
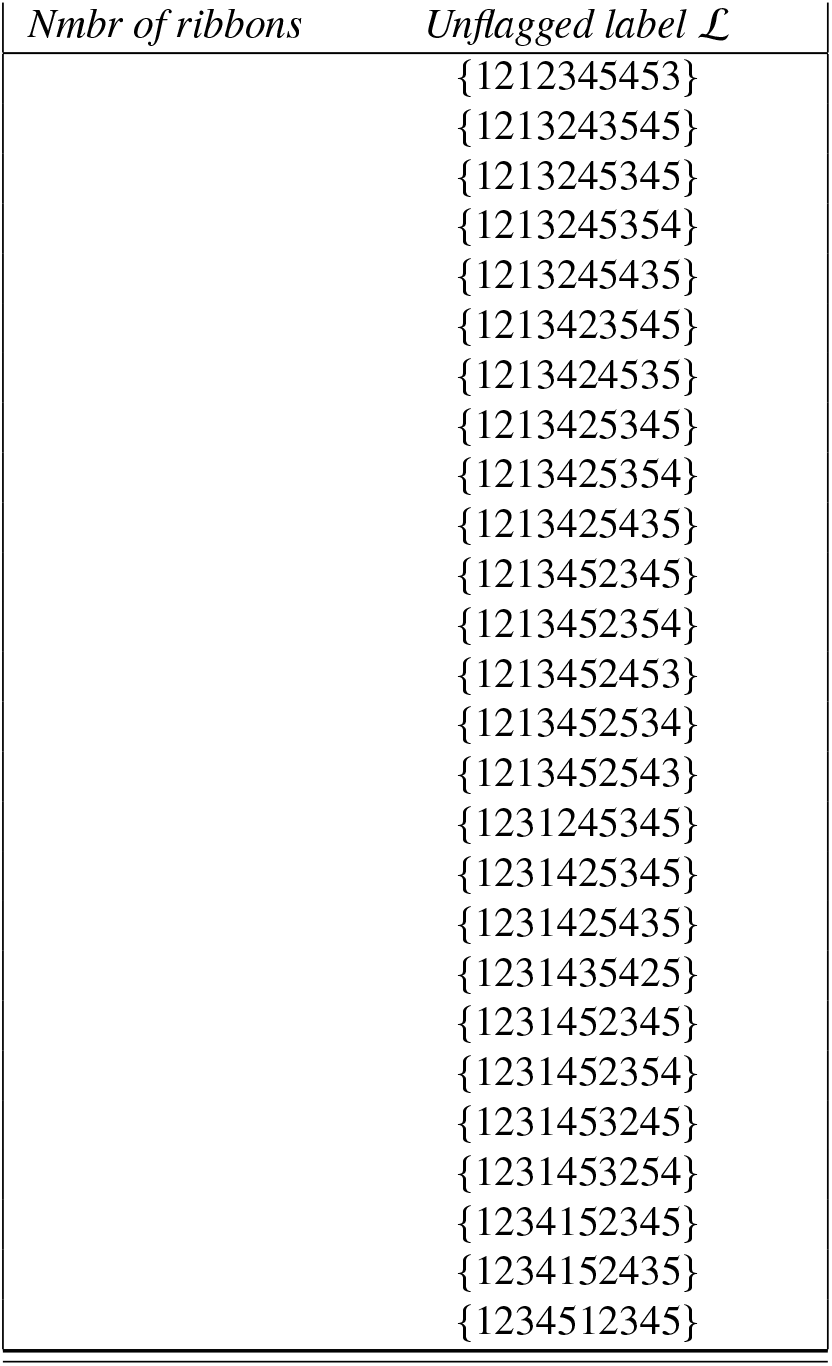
Ordered canonical unflagged labels for 0 - 5 ribbons. Labels are formed by enclosing digit strings between braces, thus 123123 → {123123} etc.

| *Nmbr of ribbons* | *Unflagged label $\mathcal{L}$* |
| --- | --- |
| 0 | $\{0\}$ |
| 1 | $\{11\}$ |
| 2 | $\{1122\}$<br>$\{1212\}$ |
| 3 | $\{112233\}$<br>$\{112323\}$<br>$\{112332\}$<br>$\{121323\}$<br>$\{123123\}$ |
| 4 | $\{11223344\}$<br>$\{11223434\}$<br>$\{11223443\}$<br>$\{11232434\}$<br>$\{11232443\}$<br>$\{11234234\}$<br>$\{11234243\}$<br>$\{11234342\}$<br>$\{11234423\}$<br>$\{11234432\}$<br>$\{12123434\}$<br>$\{12132434\}$<br>$\{12134234\}$<br>$\{12134243\}$<br>$\{12314234\}$<br>$\{12314324\}$<br>$\{12341234\}$ |
| *continued on next page . . .* |  |
| 5 | {1122334455} |
|  | {1122334545} |
|  | {1122334554} |
|  | {1122343545} |
|  | {1122343554} |
|  | {1122344553} |
|  | {1122345345} |
|  | {1122345354} |
|  | {1122345453} |
|  | {1122345534} |
|  | {1122345543} |
|  | {1123234545} |
|  | {1123234554} |
|  | {1123243545} |
|  | {1123243554} |
|  | {1123244535} |
|  | {1123245345} |
|  | {1123245354} |
|  | {1123245435} |
|  | {1123245453} |
|  | {1123245534} |
|  | {1123245543} |
|  | {1123324554} |
|  | {1123342554} |
|  | {1123345245} |
|  | {1123345254} |
|  | {1123425345} |
|  | {1123425354} |
|  | {1123425435} |
|  | {1123425453} |
|  | {1123425534} |
|  | {1123425543} |
|  | {1123435254} |
|  | {1123435452} |
|  | {1123452345} |
|  | {1123452354} |
|  | {1123452435} |
|  | {1123452453} |
|  | {1123452534} |
|  | {1123452543} |
|  | {1123453254} |
|  | {1123453452} |
|  | {1123453524} |
|  | {1123453542} |
|  | {1123454523} |
|  | {1123454532} |
|  | {1123455234} |
|  | {1123455243} |
|  | {1123455342} |
|  | {1123455432} |
|  | {1212343545} |
|  | {1212345345} |
|  | {1212345354} |
|  | {1212345453} |
|  | {1213243545} |
|  | {1213245345} |
|  | {1213245354} |
|  | {1213245435} |
|  | {1213423545} |
|  | {1213424535} |
|  | {1213425345} |
|  | {1213425354} |
|  | {1213425435} |
|  | {1213452345} |
|  | {1213452354} |
|  | {1213452453} |
|  | {1213452534} |
|  | {1213452543} |
|  | {1231245345} |
|  | {1231425345} |
|  | {1231425435} |
|  | {1231435425} |
|  | {1231452345} |
|  | {1231452354} |
|  | {1231453245} |
|  | {1231453254} |
|  | {1234152345} |
|  | {1234152435} |
|  | {1234512345} |

**Table 25:** Expanded version of Table 11 in the main text, including all nontrivial ‘uncrossed’ folds of an open single strand 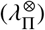 containing up to 5 contracted antiparallel double-helices and fold diagrams whose strand closure yields an uncrossed contracted fully-flagged fold 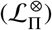. Fold diagrams for all linear folds with up to 4 contracted duplexes are shown in the main text.

| $\lambda_{\Pi}^{\otimes}$ | $\mathcal{L}_{\Pi}^{\otimes}$ |
| --- | --- |
| $[1_e 2_e 12]$ | $\{1_e 2_e 12\}$ |
| $[1_e 2_e 13_e 23]$ | $\{1_e 2_e 12\}$ |
| $[1_o 2_e 13_o 23]$ | $\{1_e 2_e 12\}$ |
| $[1_o 2_o 3_o 123]$ | $\{1_o 2_o 3_o 123\}$ |
| $[1_e 2_e 123_e 4_e 34]$ | $\{1_e 2_e 123_e 4_e 34\}$ |
| $[1_e 2_e 13_e 24_e 34]$ | $\{1_e 2_e 13_e 24_e 34\}$ |
| $[1_e 2_e 13_e 4_e 342]$ | $\{1_e 2_e 123_e 4_e 34\}$ |
| $[1_e 2_o 3_o 14_o 234]$ | $\{1_e 2_e 13_e 24_e 34\}$ |
| $[1_o 2_o 3_o 14_e 234]$ | $\{1_e 2_e 13_e 24_e 34\}$ |
| $[1_e 2_e 3_e 14_e 342]$ | $\{1_e 2_e 13_e 24_e 34\}$ |
| $[1_e 2_e 3_e 234_e 14]$ | $\{1_e 2_e 123_e 4_e 34\}$ |
| $[1_e 2_e 3_e 24_e 143]$ | $\{1_e 2_e 13_e 24_e 34\}$ |
| $[1_e 2_e 3_e 24_e 314]$ | $\{1_e 2_e 13_e 24_e 34\}$ |
| $[1_e 2_e 3_e 4_e 2413]$ | $\{1_e 2_e 13_e 24_e 34\}$ |
| $[1_e 2_e 3_e 4_e 3142]$ | $\{1_e 2_e 13_e 24_e 34\}$ |
| $[1_e 2_e 123_e 4_e 35_e 45]$ | $\{1_e 2_e 123_e 4_e 35_e 45\}$ |
| $[1_e 2_e 123_o 4_o 5_o 345]$ | $\{1_e 2_e 123_o 4_o 5_o 345\}$ |
| $[1_e 2_e 123_o 4_e 5_e 453]$ | $\{1_e 2_e 123_o 4_e 5_e 453\}$ |
| $[1_e 2_e 13_e 24_e 35_e 45]$ | $\{1_e 2_e 13_e 24_e 35_e 45\}$ |
| $[1_e 2_e 13_e 24_e 5_e 435]$ | $\{1_e 2_e 13_e 24_e 5_e 435\}$ |
| $[1_e 2_e 13_e 24_e 5_e 453]$ | $\{1_e 2_e 123_e 4_e 34\}$ |
| $[1_o 2_e 13_o 24_e 5_e 453]$ | $\{1_e 2_e 123_e 4_e 34\}$ |
| $[1_o 2_e 13_o 4_e 235_o 45]$ | $\{1_o 2_e 13_o 4_e 235_o 45\}$ |
| $[1_e 2_e 13_e 4_e 245_e 35]$ | $\{1_e 2_e 13_e 4_e 245_e 35\}$ |
| $[1_o 2_e 13_e 4_o 245_e 35]$ | $\{1_o 2_e 13_e 4_o 245_e 35\}$ |
| $[1_e 2_e 13_o 4_o 25_o 345]$ | $\{1_e 2_e 13_o 4_o 25_o 345\}$ |
| $[1_o 2_e 13_e 4_o 25_o 354]$ | $\{1_o 2_e 13_e 4_o 25_o 354\}$ |
| $[1_e 2_e 13_e 4_e 25_e 435]$ | $\{1_e 2_e 13_e 4_e 25_e 435\}$ |
| $[1_e 2_e 13_o 4_o 25_e 435]$ | $\{1_e 2_e 13_o 4_o 25_e 435\}$ |
| $[1_e 2_e 13_e 4_e 25_e 453]$ | $\{1_e 2_e 13_e 24_e 34\}$ |
| $[1_o 2_e 13_e 4_e 25_o 453]$ | $\{1_e 2_e 13_e 24_e 34\}$ |
| $[1_o 2_e 13_o 4_o 5_o 2345]$ | $\{1_o 2_e 13_o 4_o 5_o 2345\}$ |
| $[1_e 2_e 13_e 4_e 5_e 4253]$ | $\{1_e 2_e 13_e 24_e 34\}$ |
| $[1_o 2_e 13_o 4_e 5_e 4253]$ | $\{1_e 2_e 13_e 24_e 34\}$ |
| $[1_e 2_e 13_e 4_e 5_e 4523]$ | $\{1_e 2_e 123_e 4_e 34\}$ |
| $[1_o 2_e 13_o 4_e 5_e 4523]$ | $\{1_e 2_e 123_e 4_e 34\}$ |
| $[1_e 2_e 3_e 14_e 35_e 254]$ | $\{1_e 2_e 13_e 24_e 34\}$ |
| $[1_o 2_e 3_e 14_o 35_e 254]$ | $\{1_e 2_e 13_e 24_e 34\}$ |
| $[1_e 2_e 3_e 234_e 15_e 45]$ | $\{1_e 2_e 123_e 4_e 34\}$ |
| $[1_o 2_e 3_e 234_e 15_o 45]$ | $\{1_e 2_e 123_e 4_e 34\}$ |
| $[1_e 2_e 3_e 24_e 15_e 435]$ | $\{1_e 2_e 13_e 24_e 34\}$ |
| $[1_o 2_e 3_e 24_e 15_o 435]$ | $\{1_e 2_e 13_e 24_e 34\}$ |
| $[1_e 2_e 3_e 24_e 315_e 45]$ | $\{1_e 2_e 13_e 24_e 34\}$ |
| $[1_e 2_o 3_e 24_e 315_o 45]$ | $\{1_e 2_e 13_e 24_e 34\}$ |
| $[1_e 2_e 3_e 4_e 2415_e 35]$ | $\{1_e 2_e 13_e 24_e 34\}$ |
| $[1_o 2_e 3_e 4_e 2415_o 35]$ | $\{1_e 2_e 13_e 24_e 34\}$ |
| $[1_e 2_e 3_e 4_e 3415_e 25]$ | $\{1_e 2_e 123_e 4_e 34\}$ |
| $[1_o 2_e 3_e 4_e 3415_o 25]$ | $\{1_e 2_e 123_e 4_e 34\}$ |

| $\lambda_{\Pi}^{\otimes}$ | $\mathcal{L}_{\Pi}^{\otimes}$ |
| --- | --- |
| $[1_o 2_o 3_o 4_o 5_o 12345]$ | $\{1_o 2_o 3_o 4_o 5_o 12345\}$ |

## Footnotes

1 Each ribbon has its own pair of flags, e.g. orientation flags, denoted ⊗_*i*_. For typesetting convenience, we denote 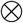 as the combined flag, combining all individual flags 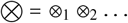.

2 Note that odd-parity twists are denoted *o*, whereas zero-twists are denoted 0

3 Analysis of these folds was done by Neave Taylor.

4 This is a genus-3 fold, since its circular diagram (Fig. 7(d)) is contained within the circular diagrams of the fully-duplexed genus-3 folds 11,12,14,15,16 in Table 2. One possible genus-3 embedding is shown in Fig. 7(m).

